# Subpopulations of soluble, misfolded proteins commonly bypass chaperones: How it happens at the molecular level

**DOI:** 10.1101/2021.08.18.456736

**Authors:** Ritaban Halder, Daniel A. Nissley, Ian Sitarik, Edward P. O’Brien

## Abstract

Subpopulations of soluble, misfolded proteins can bypass chaperones within cells. The scope of this phenomenon and the lifetimes of these states have not been experimentally quantified, and how such misfolding happens at the molecular level is poorly understood. We address the first issue through a meta-analysis of the experimental literature. We find that in all quantitative protein refolding-function studies, there is always a subpopulation of soluble but misfolded and less-functional protein that does not fold in the presence of one or more chaperones. This subpopulation ranges from 8% to 50% of the soluble protein molecules in solution. Fitting the experimental time traces to a kinetic model, we find these chaperone-bypassing misfolded states take months or longer to fold and function in the presence of different chaperones. We next addressed how, at the molecular level, some misfolded proteins can evade chaperones by simulating six different proteins interacting with *E. coli*’s GroEL and HtpG chaperones when those proteins are in folded, unfolded, or long-lived, soluble, misfolded states. We observe that both chaperones strongly bind the unfolded state and weakly bind the folded and misfolded states to a similar degree. Thus, these chaperones cannot distinguish between the folded and long-lived misfolded states of these proteins. A structural analysis reveals the misfolded states are highly similar to the native state – having a similar size, amount of exposed hydrophobic surface area, and level of tertiary structure formation. These results demonstrate that *in vitro* it is common for appreciable subpopulations of proteins to remain misfolded, soluble, and evade the refolding action of chaperones for very long times. Further, these results suggest that this happens because these misfolded subpopulations are near-native and therefore interact with chaperones to a similar extent as properly folded proteins. More broadly, these results indicate a mechanism in which long-time scale changes in protein structure and function can persist in cells because some protein’s non-native states can bypass components of the proteostasis machinery.

**TEASER:** Near-native, misfolded protein conformations explain why some soluble proteins fail to refold in the presence of chaperones.

## INTRODUCTION

Some soluble, misfolded proteins can bypass the refolding action of chaperones *in vivo* according to folding and functional assays^1-4^. Typically, in these assays the protein of interest is purified after it has been expressed either heterologously or constitutively from different synonymous messenger RNA (mRNA) variants. A synonymous mRNA variant is an mRNA molecule where one or more codons have been replaced by a synonymous codon, which does not alter the encoded protein’s primary structure but alters the mRNA’s nucleotide sequence.

For example, introducing synonymous mutations into the Chloramphenicol acetyltransferase (CATIII) enzyme decreased its specific activity by 20%^5^. Since the specific activity is an ensemble average over the soluble fraction of proteins it can be inferred that these synonymous mutations shifted a portion of the soluble protein molecules into a misfolded ensemble with decreased activity. Many other examples of this phenomenon exist. The ability of soluble FREQUENCY (FRQ) protein to bind to its partner protein ‘White Collar-2’ (WC-2) was decreased by half when a synonymous variant of FRQ was produced^6^. Since FRQ was expressed *in vivo* this is evidence that chaperones did not catalyze the proper folding of that portion of soluble, misfolded FRQ protein molecules that could not bind WC-2.

In some of these studies alternative explanations to the formation of soluble misfolded proteins have been ruled out. Most of these studies have characterized the properties only of soluble protein through the use of ultra-centrifugation, ruling out influences from insoluble aggregates. Many also controlled for changing expression levels, ruling out the possibility that it is changes in protein levels causing this phenomenon. Finally, in some studies, gels, size-exclusion chromatography, and mass spectrometry were run to rule out the possibility that higher order, non-native oligomers were present (see Supplementary File 1).

Three fundamental questions arise from these observations: How common is it for soluble, misfolded proteins to bypass chaperones? How long does it take for these misfolded states to fold? And, finally, how do some misfolded proteins avoid chaperones at the molecular level? These are biologically important questions because the answer to the first two questions could impact our understanding of how protein homeostasis is maintained in cells. The answer to the final question would help to explain how synonymous mutations can have long term impacts on protein structure and function *in vivo*.

To address these questions, we carried out a meta-analysis of the experimental literature focused specifically on *in vitro* studies where quantitative measurements can be carried out with appropriate controls (Figure. 1a). We find that subpopulations of soluble, misfolded proteins unaffected by the presence of chaperones are the norm rather than the exception, and that these misfolded states can take months or longer to fold. To answer the third question, we use coarse-grained and all-atom molecular dynamics to simulate the interactions of newly synthesized proteins with the post-translational chaperones GroEL and HtpG (Figures. 1b,c,d,e) and identify how some misfolded states can energetically and structurally bypass these chaperones.

**Figure 1.**
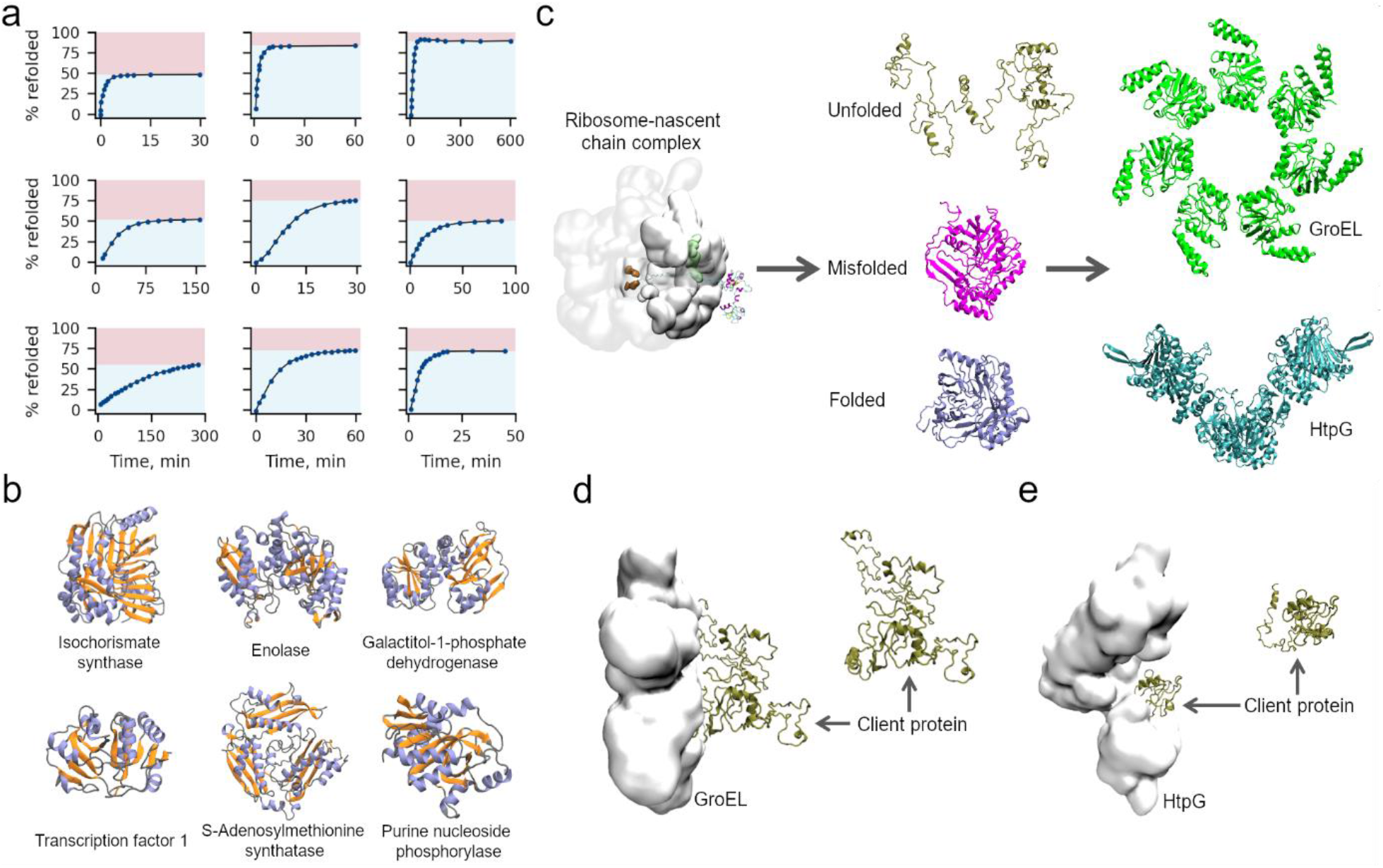
Meta-analysis of protein refolding and representations of GroEL, HtpG, and client proteins. (a) Through a meta-analysis of the experimental literature we show an appreciable fraction (indicated by the red shaded region of each subplot) of protein molecules bypass chaperones in vitro even though they are not folded, and take months or longer to reach their folded functional state. (b) Cartoon models of the native state reference structures for six proteins whose interactions with GroEL/HtpG we model. Helix, sheet, and loop regions are colored light purple, orange, and grey, respectively. (c) Unfolded, misfolded, and folded conformations were generated by synthesizing each protein using a coarse-grain ribosome-nascent chain complex. After ejection from ribosome, the nascent protein may remain unfolded, reach a misfolded state, or fold. These conformational states may then interact with several post-translational chaperones such as GroEL and HtpG. (d) Characteristic structures in both the bound and unbound states of GroEL (white space-filling model) and Isochorismate synthase (brown cartoon). (e) Same as (d) but for the bound and unbound states of HtpG and Purine nucleoside phosphorylase.

## METHODS

### Extrapolation of refolding timescales

Raw data were extracted from the published experimental papers listed in Table 1 using PlotDigitizer (http://plotdigitizer.sourceforge.net/). These raw values, which represent the percent refolded as a function of time, were then converted to the percent non-native as a function of time by taking %non-native = 1 −%refolded. The resulting %non-native versus time data series were then divided by 100%, giving the time-dependent probability of the protein being non-native, *P*_NN_(*t*), before being fit with the equation

**Table 1.**
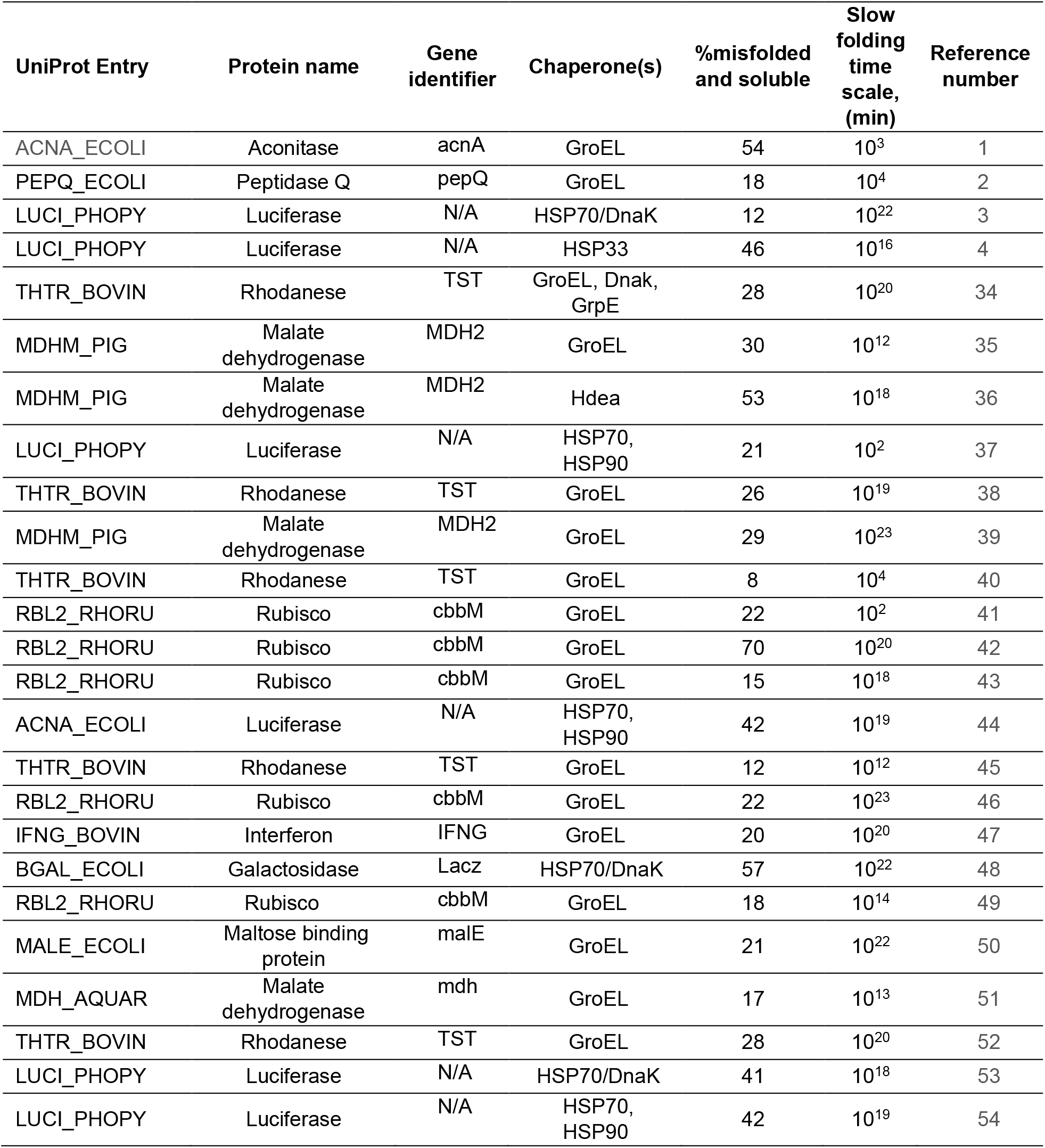
Meta-analysis of proteins that remain soluble and misfolded in the presence of chaperones.

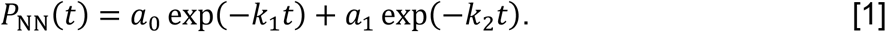

In Eq. 1, *a*_0_ + *a*_1_ ≡ 1, *t* is time, and *k*_1_ and *k*_2_ are refolding rate constants. A similar procedure was previously used to extract characteristic slow-folding timescales for protein folding via an obligate misfolded/intermediate state^7^ (Supplementary Reference Material 1). Figure S3 displays the original experimental data, *P*_NN_(*t*) values, and fit results, while Table S1 summarizes all fit parameters.

### Selection of chaperones and the client proteins

Monomers composing the molecular chaperone GroEL consist of three domain termed the apical, equatorial, and interconnecting domains. Client proteins bind to a specific region within the apical domain. All structures of GroEL used in this study are based on PDB structure 1KP8, which has been used widely in GroEL studies^8-10^. In line with previous studies of GroEL-protein interactions, we consider only the apical domain’s binding to client proteins^10^. We modelled the interactions of six client proteins with GroEL: (1) Transcription factor 1 (PDB ID: 1K7J), (2) Purine nucleoside phosphorylase (PDB ID: 1A69), (3) S-Adenosylmethionine synthetase (PDB ID: 1P7L), (4) Enolase (PDB ID: 2FYM), (5) Isochorismate synthase (PDB ID: 3HWO), and (6) Galactitol-1-phosphate dehydrogenase (PDB ID: 4A2C). We selected these proteins because they are confirmed GroEL clients^11-13^ and each was previously observed to populate long-lived misfolded states in coarse-grain simulations^7^. Unfolded, misfolded, and folded structures were selected at random from ensembles of structures generated in these previous coarse-grain simulations. We simulated interactions between GroEL and the folded, near-native misfolded, and unfolded conformations of each of these six client proteins.

In addition to simulations with GroEL, we also examined the interaction of HtpG with purine nucleoside phosphorylase^14^. A coarse-grained model of full-length HtpG was constructed based on PDB ID: 2IOQ, which was used in several earlier HtpG simulation studies^15-17^. The centre of mass of this full-length structure was positioned at the origin of the CHARMM internal coordinate system. Simulations of HtpG were otherwise conducted in the same fashion as those described for GroEL and its client proteins.

### Construction of coarse-grained protein representations

We use a Cα coarse-grained representation for GroEL, HtpG, and their client proteins^18,19^. The potential energy of a conformation within this model is calculated according to the expression

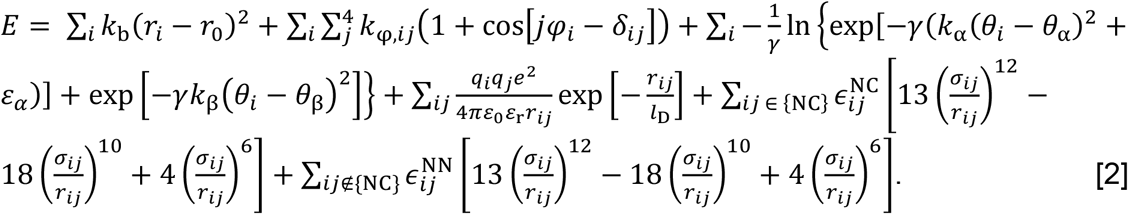

In Eq. 2 the summations represent, from left to right, contributions from virtual Cα−Cα bonds, torsion angles, bond angles, electrostatic interactions, Lennard-Jones-like native interactions, and repulsive non-native interactions to the total potential energy (*E*) of a given coarse-grain model configuration. The bond, dihedral, and angle terms have been reported elsewhere^20,21^. Electrostatic interactions are considered using Debye−Hückel theory with a Debye length, *l*_*D*_, of 10 Å and a dielectric constant of 78.5. Interaction sites representing the positively charged amino acids lysine and arginine are assigned *q* = +*e*, sites representing glutamic acid and aspartic acid are assigned *q* = −*e*, and all other interaction sites are taken to have a charge of zero^18^. We compute the contribution from native contacts to *E* using the 12−10−6 interaction potential of Karanicolas and Brooks^20^. The value of 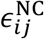, the depth of the energy minimum for any particular native contact, is calculated as 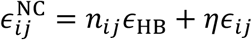 represents the energy contribution from hydrogen bonds, while *ϵ*_*ij*_ represents the energy contribution from the van der Waals contacts between a pair of residues *i* and *j* found to be in contact within the protein all-atom reference structure. *n*_*ij*_ indicates the number of hydrogen bonds formed between a pair of residues *i* and *j*. The value of *ϵ*_*ij*_ is initially set using the Betancourt−Thirumalai pairwise potential^22^ and multiplied by a constant *η* to construct a reasonably stable coarse-grain model as described below. The collision diameters, *σ*_*ij*_, between all the Cα interactions sites involved in native contacts are set equal to the distance between the Cα atoms of the corresponding amino acid residues in the crystal structure divided by 2^1/6^. van der Waals interaction energies between pairs of residues that do not share a native contact are instead computed in the final summation. For all the non-native interactions, 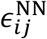 is set to be 0.000132 kcal/mol and *σ*_*ij*_ is computed as reported previously^20^.

### Selection of *η* for chaperone and client protein coarse-grain models

Values of *η* were initially set based on a previously published training set^23^ and sets of ten 1-µs Langevin dynamics simulations used to determine their suitability. All simulations were run in CHARMM^24^ version c35b5 at 310 K with a frictional coefficient of 0.050 ps^-1^, a 15-fs integration time step, and the SHAKE^25^ algorithm used to constrain all bond lengths. Coordinates were saved every 5,000 integration time steps (every 75 ps). A particular *η* value is considered suitable if the coarse-grain model has a fraction of native contacts, *q*, greater than 0.69 for at least 98% of simulation frames. Based on this procedure, we selected *η* = 1.359 for native contacts within each client protein (*i*.*e*., their intra-domain contacts), *η* = 1.800 for intra- and inter-domain contacts within and between GroEL monomers, and *η* = 1.400 for interactions between client proteins and GroEL. The *η* for HtpG-client protein interactions is also set to 1.400, and the *η* for intra-HtpG interactions is set to 1.400. Full details of these parameters are included in Tables S2, S3, and S4.

### Simulation of GroEL and HtpG interactions with client proteins

Simulations were initialized with the center of mass of the GroEL coarse-grain model at the CHARMM internal coordinate system origin. The client protein of interest was then placed in a random orientation such that the distance between its center of mass and the center of mass of the apical GroEL domain was 50 Å, with no contacts initially formed between them. Spherical harmonic restraints with force constant 0.1 kcal/(mol × Å^2^) were placed on all GroEL interaction sites to maintain its conformation and position at the origin throughout the simulation. Root Mean Square Deviation (RMSD) restraints with a force constant of 0.2 kcal/(mol× Å^2^) were used to maintain the client proteins in their initial conformations. This system is then placed in a sphere of radius 70 Å. This sphere radius of 70 Å was found to be the minimum sphere radius that can easily accommodate each of the client proteins we consider. We find qualitatively similar results using a sphere radius of 85 Å (Figure 4A, data for 0.65 M). For each unfolded, folded, and near-native misfolded client protein conformation, we ran simulations with ten different initial client protein conformations generated by randomly rotating the starting client protein conformation. Each system was then equilibrating for 20 ns before running 800 ns of production simulation for each GroEL-client protein system. All simulations were performed using CHARMM with a Langevin thermostat as described in the coarse-grain model parameterization Methods section. Control simulations were also run in which interactions between GroEL and client proteins are deactivated (Figure S3). We define the client protein and GroEL to be bound to one another in a particular simulation frame if ≥10 residues form native contacts. Our results are not sensitive to the value of this binding threshold, with qualitatively consistent results found when thresholds of ≥5 or ≥1 residue are used (Figure S1). The binding probabilities between GroEL and each client protein (Figures 3 and S2) were computed as the number of frames in which the client protein is bound divided by the total number of simulation frames across all ten trajectories.

Simulation of interactions between HtpG and its client protein purine nucleoside phosphorylase were carried out in an analogous fashion, with HtpG’s centre of mass placed at the origin of a 70-Å sphere with the client protein initially placed in a random orientation 50-Å away. All restraints, force constants, and other simulation parameters were otherwise the same as for GroEL-client protein simulations.

### All-atom simulations of GroEL and client proteins

We randomly chose one of the client proteins, Isochorismate synthase, used in our coarse-grained simulations and simulated its interactions with GroEL at all-atom resolution. We chose ten representative structures each from the ensembles of unfolded, folded, and near-native misfolded Isochorismate synthase/GroEL systems and back-mapped these 30 coarse-grained structures to all-atom resolution using a previously reported procedure^19^. Next, each of these all-atom composite structures of the GroEL heptamer and client protein were solvated in a box of SPC/E water^26^ with dimensions 14 × 14 × 14 nm^3^ and then neutralized by the addition of 58 sodium atoms. This neutralized system was then energy minimization with the steepest descent algorithm. A spherical harmonic restraint with force constant 1000 kJ/(mol × nm^2^) was applied to the GroEL heptamer structure. All-atom simulations were carried out with GROMACS 4.6.5^27^ using the AMBER03 force field^28^. Long-range electrostatic interactions were calculated with the Particle Mesh Ewald method^29^. Lennard-Jones interactions were calculated with a distance cut-off of 1.2 nm, and the temperature and pressure were maintained throughout the simulations at 310 K and 1 atm with a Nose-Hoover thermostat^30,31^ and Parrinello-Rahman barostat^32^, respectively. All bonds were constrained using the LINCS algorithm^33^ and an integration time step of 5 fs used. We performed 500 ps of equilibration followed by a 1-ns production simulation with each of the 30 all-atom conformations.

### Calculation of odds ratios of binding probabilities with and without attractive interactions between client proteins and chaperones

Odds ratios of the binding probabilities between chaperones and unfolded (U) or folded (F) conformations of client proteins with attractive van der Waals interactions on or off were calculated as

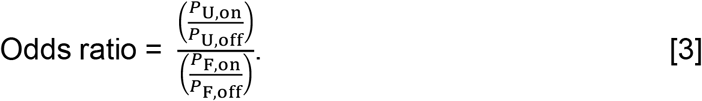

In Eq. 3 the terms *P*_U,on_ and *P*_U,off_ are the probabilities of protein/chaperone binding with interactions turned on or off, respectively, for unfolded client protein conformations. The terms *P*_F,on_ and *P*_F,off_ are the analogous values computed from simulations initialized with the client protein in the folded state. Odds ratios for interactions between misfolded or folded client protein conformations with chaperone interactions turned either on or off were computed using the equation

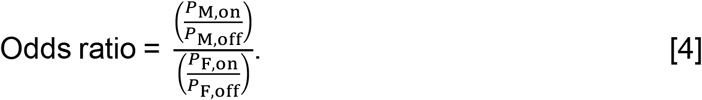

In Eq. 4, *P*_M,on_ and *P*_M,off_ are the binding probabilities of a misfolded client protein to chaperone with attractive van der Waal interactions turned on or off, respectively.

### Identification of entangled protein conformations

The six proteins whose interactions with GroEL/HtpG we model here were previously identified to populate entangled conformations^7^ when they misfold. These entanglements are local non-covalent lasso-type entanglements that are not present in the native state that are associated with long-lived misfolded states within the *E. coli* proteome. We calculated the entanglement (*G*) of the native and near-native like misfolded states (Table S5) based on previously described protocol^7^. The Perl code used to compute *G* is available on GitHub at https://github.com/obrien-lab/topology_analysis. The value of *G* is computed as

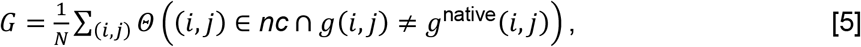

where (*i, j*) is one of the native contacts in the native crystal structure; *nc* is the set of native contacts formed in the current structure; *g*(*i, j*) and *g*^native^(*i, j*) are, respectively, the total linking number of the native contact (*i, j*) in the current and native structures; *N* is the total number of native contacts within the native structure; and the selection function *Θ* equals 1 when the condition is true and 0 when it is false. The larger *G* is the larger the number of residues that have changed their entanglement status relative to the native state. That is, *G* reports on the presence of non-native entanglements in structures.

## RESULTS

### Soluble misfolded proteins bypass the *E. coli* chaperone machinery *in vitro*

We carried out a meta-analysis of the experimental literature reporting time-courses of protein refolding and acquisition of function (Figure 1a). We focus on *in vitro* studies because they are capable of controlling for a number of factors that are currently not possible to control for *in vivo*. A typical experiment involves splitting a purified protein sample into two test tubes, applying a denaturant (such as urea) to one sample, then performing a dilution jump experiment to initiate protein refolding and measuring the time course of the fraction of functional protein. For such a study to make it into our analysis we require: (i) that the signal be normalized by the activity of the non-denatured protein sample; (ii) that centrifugation or ultracentrifugation be performed at each time point to remove insoluble aggregates before analysis; and (iii) the fraction of folded-functional protein be measured in the presence of one or more chaperones. Twenty-five papers spanning three decades meet these criteria^1-4,34-54^ (see Table 1 and Supplementary File 1). Five different chaperones are represented in these studies – GroEL, HSP70, HSP90, HSP33 and Hdea – and ten different client proteins – Malate dehydrogenase, Rhodanese, Luciferase, Rubisco, Aconitase, Peptidase Q, Maltose binding protein, Interferon gamma, Dihydropicolinate synthase and Galactosidase. Twenty-one of these studies measured protein folding in the presence of one chaperone, three studies used two different chaperones, and one study used a mixture of three different chaperones. The duration of the time-courses monitoring refolding in these studies ranged from 5 minutes to 600 minutes, with an average of 150 minutes and a median of 140 minutes. These details are summarized for each study in Table 1.

In each of these studies there is always a fraction of soluble protein that does not attain a folded, functional state by the last time point. The percentage of molecules that did not fold range from a low of 8% to a high of 50%. Since protein structure equals function, these percentages reflect the fraction of protein molecules that are soluble, non-functional and likely misfolded in solution. Thus, there is always a subpopulation of soluble protein that misfolds and whose folding is not catalysed by the presence of these chaperones. One example is shown in Figure 2, where the unfolded client protein Rhodanese is incubated with GroEL/GroES, DnaJ, and DnaK. In this example, even 150 minutes after refolding was initiated with a dilution jump, 35% of soluble Rhodanese remains misfolded.

**Figure 2.**
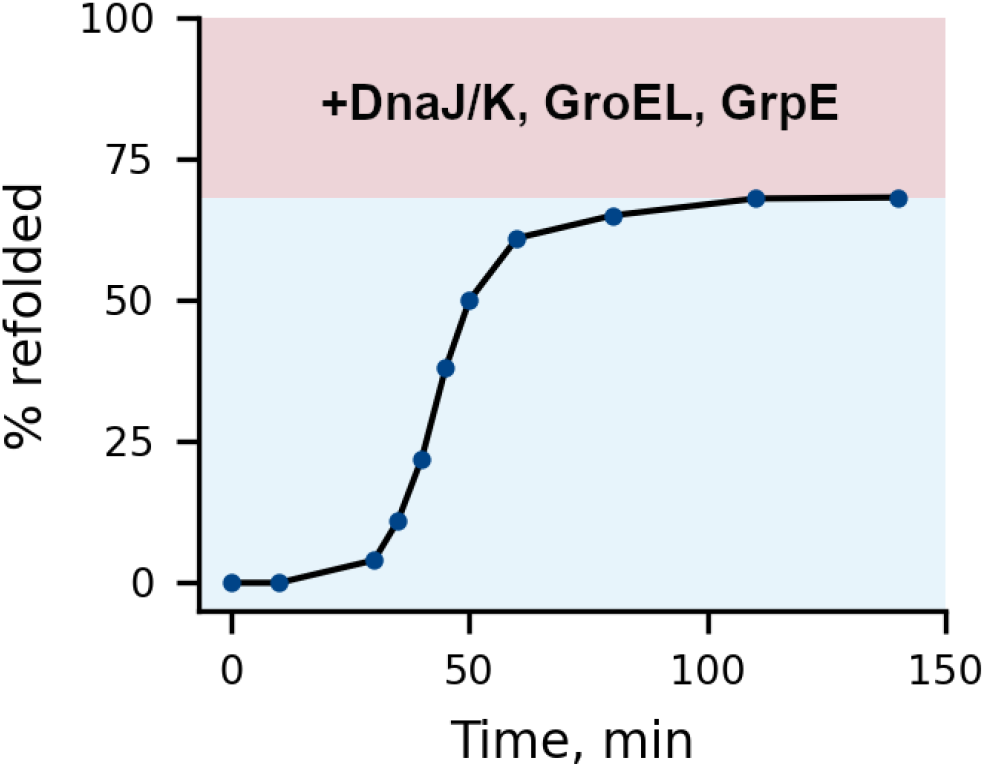
Time course of reactivation/refolding of soluble rhodanese in a mixture of the chaperones DnaJ/K, GroEL and GrpE. Rhodanese was initially unfolded using guanidine hydrochloride and refolding then monitored after a dilution jump and the addition of chaperones. Note that >25% of Rhodanase (red shaded region) is unable to reach its fully folded conformation even in the presence of these chaperones during the time course of the experiment. A kinetic fit (Eq. 1) indicates this subpopulation will take 10^20^ min (∼360,000,000 years) to fold. Experimental data were extracted from Ref. 34 Figure. 4a using PlotDigitizer (see Supplementary File 1).

### Refolding of soluble, misfolded states typically takes months or longer

The time courses reported in these studies allow us to estimate how long it takes for the subpopulation of soluble, misfolded states to fold and function. Applying a double exponential fit (see Methods, Figure S3 and Table S1) to the time courses, and interpreting the slower characteristic time scale as the folding time of the soluble misfolded fraction, we find that these states take between 0.08 and 10^18^ days to fold, with a mean of 10^13^ days and a median of 10^9^ days. Thus, for most proteins, their soluble misfolded states convert to the native state extremely slowly in the presence of chaperones.

### Misfolded states have similar binding affinities to GroEL and HtpG as the native state ensemble

We next asked how is it possible that long-lived misfolded proteins are able to bypass the post-translational cellular chaperone machinery? To address this question we used coarse-grained Langevin dynamics to calculate the binding affinity between the chaperone GroEL – one of the best studied chaperones in *E. coli* – and three distinct conformational states of client proteins: folded, unfolded, and misfolded (see Methods). Six client proteins were selected based on their known interactions with GroEL^10-12^ and previous simulations that indicate these protein can populate long-lived misfolded states^7^ (Figure 1b). In addition to computing the binding affinities of each of these proteins to GroEL, we also consider the interactions of Purine nucleoside phosphorylase with the chaperone HtpG (Figure 1c). Structural properties of each of the protein conformational states used in these simulations are reported in Table 2.

**Table 2.** Structural comparison of unfolded, folded, and near-native misfolded states

| Client protein | Conformational State | Fraction of native contacts, $Q$ | Hydrophobic solvent accessible surface area, $\text{\AA}^2$ |
| --- | --- | --- | --- |
| Isochorismate synthase | Unfolded | 0.59 | 87.4 |
|  | Misfolded | 0.85 | 25.7 |
|  | Folded | 0.92 | 20.2 |
| Enolase | Unfolded | 0.70 | 81.1 |
|  | Misfolded | 0.87 | 20.9 |
|  | Folded | 0.92 | 16.8 |
| Galactitol -1-phosphate dehydrogenase | Unfolded | 0.51 | 77.5 |
|  | Misfolded | 0.86 | 24.8 |
|  | Folded | 0.94 | 20.1 |
| Transcription factor-1 | Unfolded | 0.19 | 98.1 |
|  | Misfolded | 0.85 | 29.4 |
|  | Folded | 0.93 | 25.3 |
| S-Adenosylmethionine synthetase | Unfolded | 0.44 | 69.3 |
|  | Misfolded | 0.84 | 30.5 |
|  | Folded | 0.94 | 27.9 |
| Purine nucleoside phosphorylase | Unfolded | 0.38 | 90.4 |
|  | Misfolded | 0.88 | 30.3 |
|  | Folded | 0.93 | 26.6 |

We find, as expected, that the unfolded ensembles of all six client proteins are more likely to bind to GroEL than their native state ensembles (Figure 3). Purine nucleoside phosphorylase’s unfolded ensemble also binds with a higher probability than its folded ensemble to HtpG. Surprisingly, however, the misfolded states of all six proteins interact with GroEL with a similar probability as the native state, and the same trend is seen for Purine nucleoside phosphorylase with HtpG (Figure 3).

**Figure 3.**
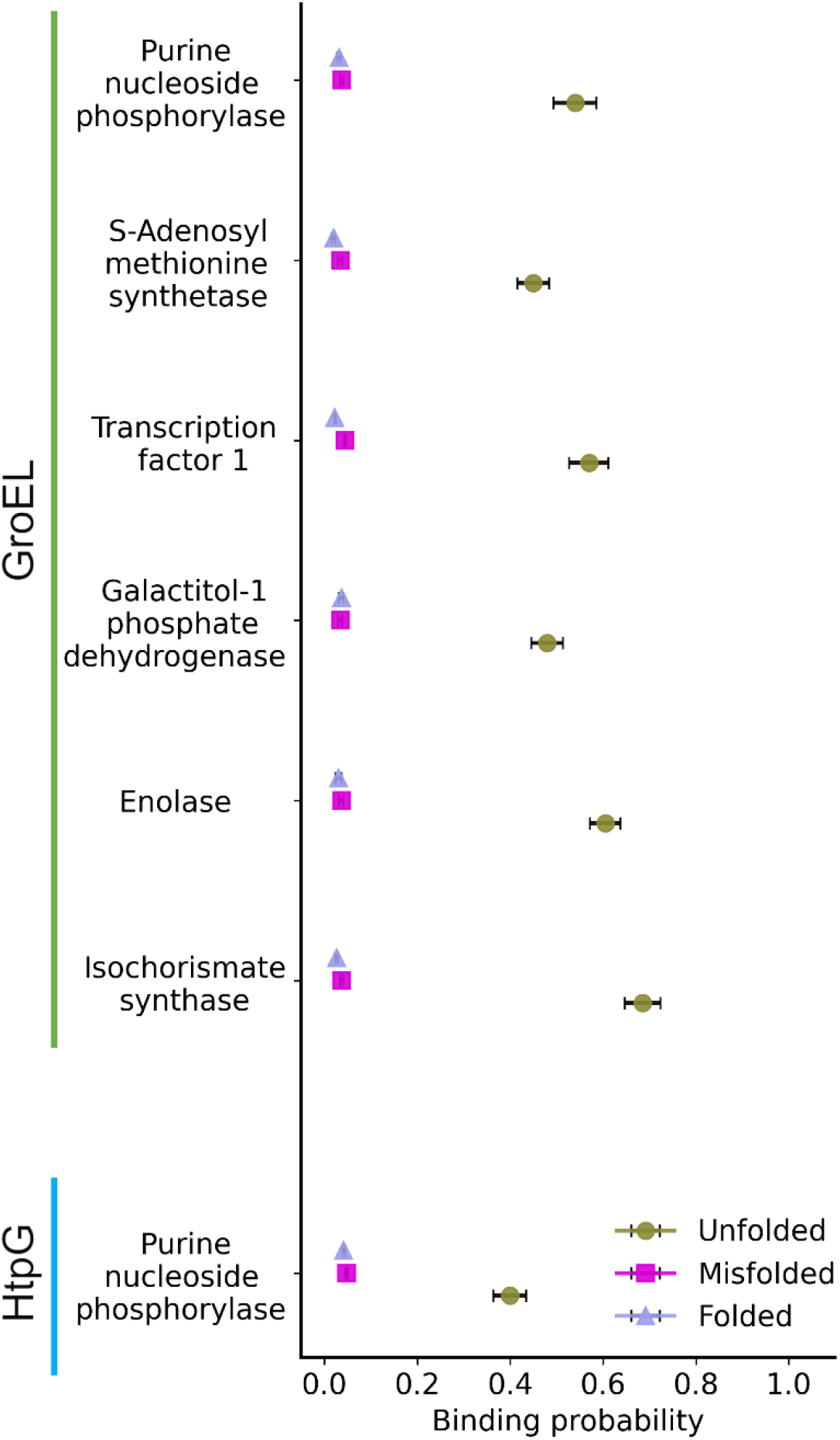
Binding probabilities of the unfolded, misfolded, and folded states of client proteins to chaperones at 310 K and a 1.15 mM concentration of protein. The binding probabilities between the chaperone GroEL with the unfolded (light brown), misfolded (magenta), or folded (light blue) conformations of Isochorismate synthase, Enolase, Galactitol-1-phosphate dehydrogenase, Transcription factor 1, S-Adenosylmethionine synthetase, and Purine nucleoside phosphorylase are displayed to the right of the green line. The binding probability of the chaperone HtpG with Purine nucleoside phosphorylase is indicated by the light blue line. The binding probability of each of the unfolded, misfolded, and folded states for each client protein and GroEL or HtpG were averaged over ten independent trajectories. Error bars are 95% confidence intervals generated by bootstrapping 10^6^ times.

Two different hypotheses can explain these observations. First, the finite size of the simulation environment, coupled with the larger size of the unfolded state, may lead the unfolded state to more frequently contact GroEL than the more compact native and misfolded conformations. Alternatively, differences in attractive interactions between these states and GroEL/HtpG may drive the observed behavior. To control for size effects and rule out the first hypothesis, we reran all the simulations allowing only excluded volume interactions between the client protein and GroEL (Figure S2), and calculated two sets of odds ratios. First, we calculated the odds that the unfolded state interacts with GroEL versus the folded state when attractive interactions are present versus the same odds computed from simulations in which attractive interactions are deactivated (Eqs. 3-4, Table 3). For example, the unfolded Purine nucleoside phosphorylase is 4.3 times more likely to interact with GroEL compared to the folded state when attractive interactions between GroEL and Purine nucleoside phosphorylase are present (Table 3). Thus, it is the interactions between the client protein and GroEL, not the differences in the sizes of the protein conformational states, that primarily drives the observed differences in binding probability observed in Figure 3 between the folded and unfolded states. Second, we calculated the odds ratio of misfolded states binding GroEL with and without interactions present compared to the folded state (Table 3). We find that most of these ratios are statistically no different than 1, meaning that neither size differences nor interaction differences contribute to differences in GroEL binding probabilities between native and misfolded states. This result indicates that the native and misfolded states interact with GroEL to a similar extent.

**Table 3.** Odd’s ratios between probabilities of binding in the unfolded, misfolded, and folded states computed using Eqs. 3 & 4; *p*-values were calculated using the student t-test with *α* = 0.05.

| Client protein | Odds Ratio (U/F) | $p$-value | Odds Ratio (M/F) | $p$-value |
| --- | --- | --- | --- | --- |
| Isochorismate synthase | 2.8 | $9 \times 10^{-5}$ | 1.2 | 0.2 |
| Enolase | 2.4 | $1 \times 10^{-4}$ | 1.1 | 0.1 |
| Galactitol-1-phosphate-dehydrogenase | 3.4 | $2 \times 10^{-3}$ | 0.9 | 0.6 |
| Protein Transcription factor 1 | 2.7 | $1 \times 10^{-5}$ | 1.2 | 0.006 |
| S-adenosylmethionine synthetase | 3.2 | $2 \times 10^{-5}$ | 1.1 | 0.04 |
| Purine nucleoside phosphorylase | 4.3 | $9 \times 10^{-6}$ | 1.0 | 0.5 |

We conclude that long-lived misfolded states can bypass GroEL and HtpG because they exhibit no excess interaction with these chaperones beyond that of the native state ensembles’ interaction propensity.

### Conclusions are robust to changes in concentration and model resolution

We next examined if these conclusions are sensitive to changes in concentration or model resolution. To test this first issue we simulated Isochorismate synthase in the presence of GroEL at a lower concentration of 0.65 mM (earlier simulations were performed at 1.15 mM). We find that the qualitative differences in binding affinities between different conformational states persist at this lower concentration (Figure 4a). The unfolded state has a higher binding probability, and the folded and misfolded states have similar affinities. Thus, our conclusions are independent of protein concentration.

**Figure 4.**
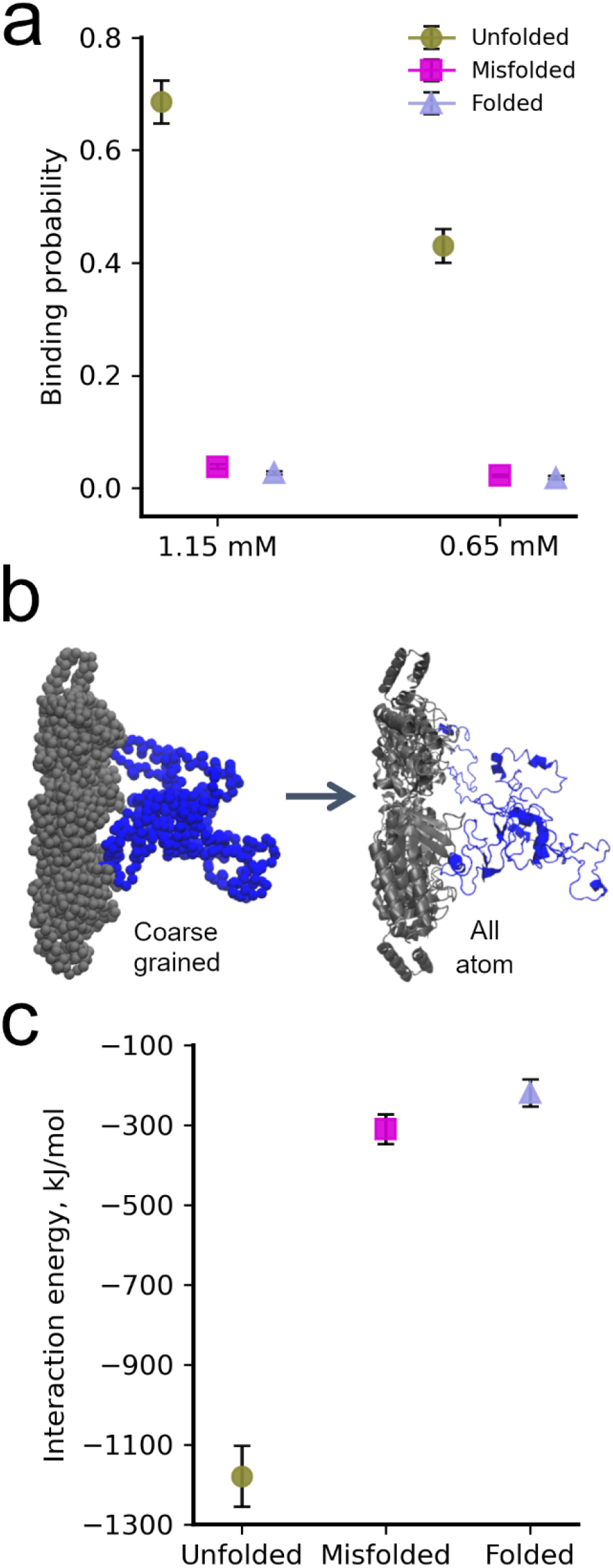
Effect of concentration and model resolution on binding probabilities. (a) Binding probability of the unfolded, misfolded, and folded states of Isochorismate synthase to GroEL at client protein concentrations of 1.15 mM and 0.65 mM. (b) Snapshot of Isochorismate synthase bound to GroEL in their coarse-grained (before backmapping) and all-atom (after backmapping) representations. (c) The all-atom interaction energy between GroEL and unfolded, misfolded, and folded Isochorismate synthase is displayed. Each interaction energy is the average from ten independent simulations. Error bars are 95% CIs from bootstrapping 10^6^ times.

To test if our results are dependent on model resolution we backmapped each of the ten coarse-grain folded, unfolded, and misfolded conformations of Isochorismate synthase bound to GroEL to an all-atom representation (Figure 4b) and ran 1-ns all-atom simulations in explicit water for each of these 30 systems (see Methods). We then calculated the average interaction energy between Isochorismate synthase and GroEL during the simulations. We find that the interaction energy of the unfolded, misfolded, and folded states are, respectively, -1,190 kJ/mol (95% CI[-1,266:-1,114]), -300 kJ/mol (95% CI [-337:-263]), and -220 kJ/mol(95% CI [-253:-188]) calculated via 10^6^ bootstrapping iterations (Figure 4c). Thus, regardless of model resolution, the folded and misfolded states have highly similar interaction energies with GroEL.

### Misfolded states bypass chaperones because they are similar to the native state

To understand the structural origins of our binding results we characterized the size, interface, and how native-like each conformational ensemble was by calculating, respectively, the ensemble-averaged radius-of-gyration, solvent accessible surface area, and fraction of native contacts. We observe (Table 2) that the unfolded ensemble is consistently larger and has more exposed surface area than the native state for all client proteins, explaining why it binds more strongly to GroEL and, in the case of Purine nucleoside phosphorylase, to HtpG. In contrast, the misfolded states are much more similar to the native state than they are to the unfolded state. Averaging across all client proteins, the misfolded state is just 8% larger than the native state (characterized by the percent difference in *R*_g_), has 90% of the native contacts formed, and has a surface area that is only 9% larger than the native state on average. Thus, the misfolded states have structural properties that are similar to the native state, explaining why they interact with these chaperones to a similar degree as the native state.

The reason why these particular misfolded states are kinetically long-lived was previously explained^7^. These misfolded states involve non-native changes in non-covalent lasso entanglements. A non-covalent lasso entanglement involves two structural components: a geometrically closed protein backbone loop, and a N- or C-terminal segment that threads through that loop. The loop is closed by a non-covalent native contact. One third of structures in the protein databank have one-or-more non-covalent lasso entanglements, while two thirds do not^55^. A non-native change of entanglement, characterized by our metric *G* (see Eq. 5 and Methods), means that a protein that forms one of these self-entanglements in the native state does not form it in the misfolded state, while a protein that does not form one of these self-entanglements in the native state does form it in the misfolded state. Each of the six proteins we simulated misfold into states that exhibit a non-native gain of entanglement relative to the native state. When such non-native changes of entanglement occur in near-native misfolded states, it is an energetically costly and slow process to reach the native state because the protein must unfold to allow the correct entanglement state to be achieved. An illustration of a non-native gain of entanglement (which is present in its misfolded conformation that we simulated) is illustrated for protein Enolase in Figure 5, where the arrow points to the crossing point of the threading segment through the loop in Figure 5b.

**Figure 5.**
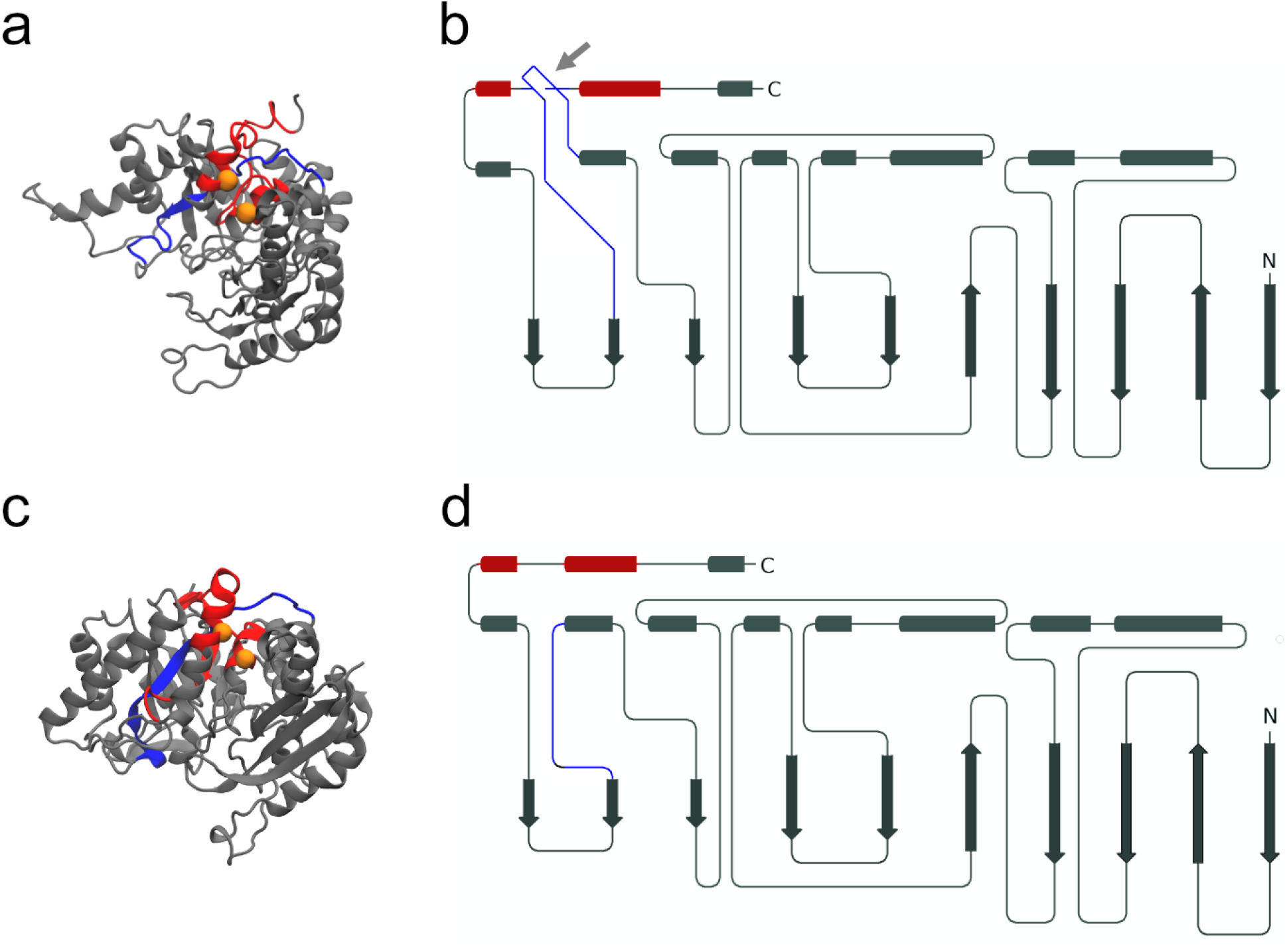
Structural representations of near-native entangled and native Enolase. (a) Cartoon model of the near-native entangled state of Enolase. The closed loop and threading segment that form the entanglement are colored red and blue, respectively. The pair of residues that form the native contact that closes the loop are highlighted by gold spheres at the location of their Cα atoms. (b) Schematic representation of the gain in entanglement for Enolase shown in (a). The threading of the C-terminus through a loop is indicated by the grey arrow. (c) Cartoon model of the native state of Enolase, which contains zero entanglements. (d) Schematic representation of the native state of Enolase with no entanglements.

## DISCUSSION

In this study we have used a combination of published experimental data and multi-scale simulations to answer a number of basic molecular biology and biochemistry questions concerning nascent protein structure and function *in vivo*. The observations that synonymous mutations can have long term effects on protein structure and function *in vivo* strongly imply that soluble, misfolded subpopulations persist in cells and that chaperones do not catalyse their folding on biologically relevant time scales.

This motivated us to re-analyze the last several decades of relevant literature to examine if there was quantitative *in vitro* data to test this inference. We indeed find that in every single *in vitro* experiment in which there are rigorous controls and normalization there is always a subpopulation of soluble, misfolded, less-functional protein that do not fold in the presence of chaperones. These subpopulations can be as high as 50% of the total protein molecules in solution. Applying a kinetic model to the experimental time courses, we find these particular soluble misfolded states can take months or years to fold in the presence of chaperones. Thus, the *in vivo* and *in vitro* data demonstrate the same phenomenon: some soluble, misfolded proteins can bypass the chaperone machinery.

Does this mean that all misfolded proteins bypass chaperones? At equilibrium proteins adopt an ensemble of distinct structures, existing on a continuum from more to less ordered and hence, for globular proteins, span from exposing less to more hydrophobic surface area. Thus, some protein conformations will be more or less likely to interact with chaperones, and hence different misfolded conformations will have different affinities for chaperones. Indeed, in our simulation results we observe that when the protein is less ordered and more unfolded the binding affinity for the chaperones increase. Thus, not all misfolded states will bypass chaperones.

A key aspect of this study is that the simulations utilized six proteins that have been previously shown^7^ to populate long-lived misfolded states, and compared their chaperone binding affinity to that of the unfolded and folded states. The fact that these are long-lived misfolded states is essential for two reasons. First, if the misfolded states rapidly folded they would not need chaperones to acquire their function. Second, it is these kinetically trapped misfolded states that are biologically relevant as they can have long-term impacts on subcellular processes and phenotype through a loss of function. Through these comparisons we were able to demonstrate – using both coarse-grained and all-atom protein representations – that the misfolded and native states have similar affinities for chaperones, indicating that chaperones do not treat these particular long-lived misfolded states much differently than they do the native state. The structural and energetic origin of this lack of differentiation comes from the high structural and surface similarity of the misfolded and native states.

It was previously shown^7^ for these six misfolded states that kinetic trapping and native state similarity are intimately connected. These six misfolded states are long lived for two reasons. They form a non-covalent lasso entanglement – meaning part of their protein backbone created what can be geometrically defined as a closed loop, and the N- or C-terminal segment threads through this loop – but also they contain significant native structure. This combination means that to disentangle and reach the native state large portions of the misfolded protein must unfolded, which is a very slow process^56^. Hence, the large amount of native structure around an entanglement leads to long lived states. By choosing to study misfolded states that were kinetically long lived we concomitantly selected for misfolded states that were native like.

Interestingly, from two simulation studies^7,57^ (Supplementary Reference Material 1 and Supplementary Reference Material 2) it was predicted that many soluble misfolded states could take anywhere from days to years to fold. Our analysis of the published experimental data yields a similar range. Thus, the previous simulation predictions are qualitatively accurate, and are further evidence that these simulation techniques can complement experimental efforts in the area of protein structure and function *in vivo*.

Our identification of near-native self-entanglements as a relevant mechanism is not mutually exclusive with other soluble misfolding and misfunctioning mechanisms that can occur *in vivo*. These other mechanisms can include non-native dimer swapped structures^58-60^, aberrant protein isoforms from mRNA splicing^61^, post-translational modifications^62-63^, and chemical processes that age proteins such as oxidation^64-65^. Future experiments that seek to detect these non-native changes of entanglement must rule out these alternative explanations.

One critique of our meta-analysis is that we only analyzed *in vitro* data, and the lack of an *in vivo* environment, which includes vectorial synthesis by the ribosome and the presence of many different chaperones, artificially increased the subpopulations of soluble misfolded protein. While it is possible that the absolute amount of soluble, misfolded protein may decrease in the cellular context they are unlikely to be entirely eliminated, and such a result would not change our molecular explanation for this phenomenon. It has been observed that even when a protein is synthesized by the ribosome in the presence of a chaperone it still populates states that remain soluble and non-functional – thus, vectorial synthesis does not eliminate these subpopulations^66^. Furthermore, the aforementioned studies of the influence of synonymous codons on protein function in cells are consistent with these subpopulations existing *in vivo*. Thus, the *in vitro* results point towards the ability of many proteins to exhibit subpopulations of soluble misfolded states that bypass chaperones and are long-lived kinetic traps. More broadly, these issues emphasize that the experimental community needs to carry out quantitative measurements of folding and function *in vivo* that are as accurate and precise as the measurements *in vitro*.

These and other recent findings^67^ are providing a new perspective on protein structure and function *in vivo* in which proteins can populate a fourth state that is soluble, misfolded, less functional, not rapidly degraded, not likely to aggregate, nor acted upon excessively by chaperones. The population of this fourth state can be influenced by both translation-elongation kinetics, as demonstrated by synonymous mutation studies, or through rounds of chemical denaturation and renaturation, as seen in our meta-analysis. It is natural to hypothesize other perturbations could influence their populations as well, such as changes in temperature^68^. Experimental efforts to structurally characterize these self-entangled states is likely to be a fruitful area of future research, as the implications of these states for protein structure, function and homeostasis are broad and fundamental.

## Supporting information

Supplementary Reference Material 1

Supplementary Reference Material 2

## ACKNOWLEDGEMENTS

E.P.O. acknowledges support from the National Science Foundation (MCB-1553291) as well as from the National Institutes of Health (R35-GM124818). Portions of numerical computations and data analysis in this work have been carried out on the CyberLAMP cluster, which is supported by NSF-MRI-1626251 and operated by the Institute for Computational and Data Sciences at The Pennsylvania State University.

## Supporting Information

**Figure S1.**
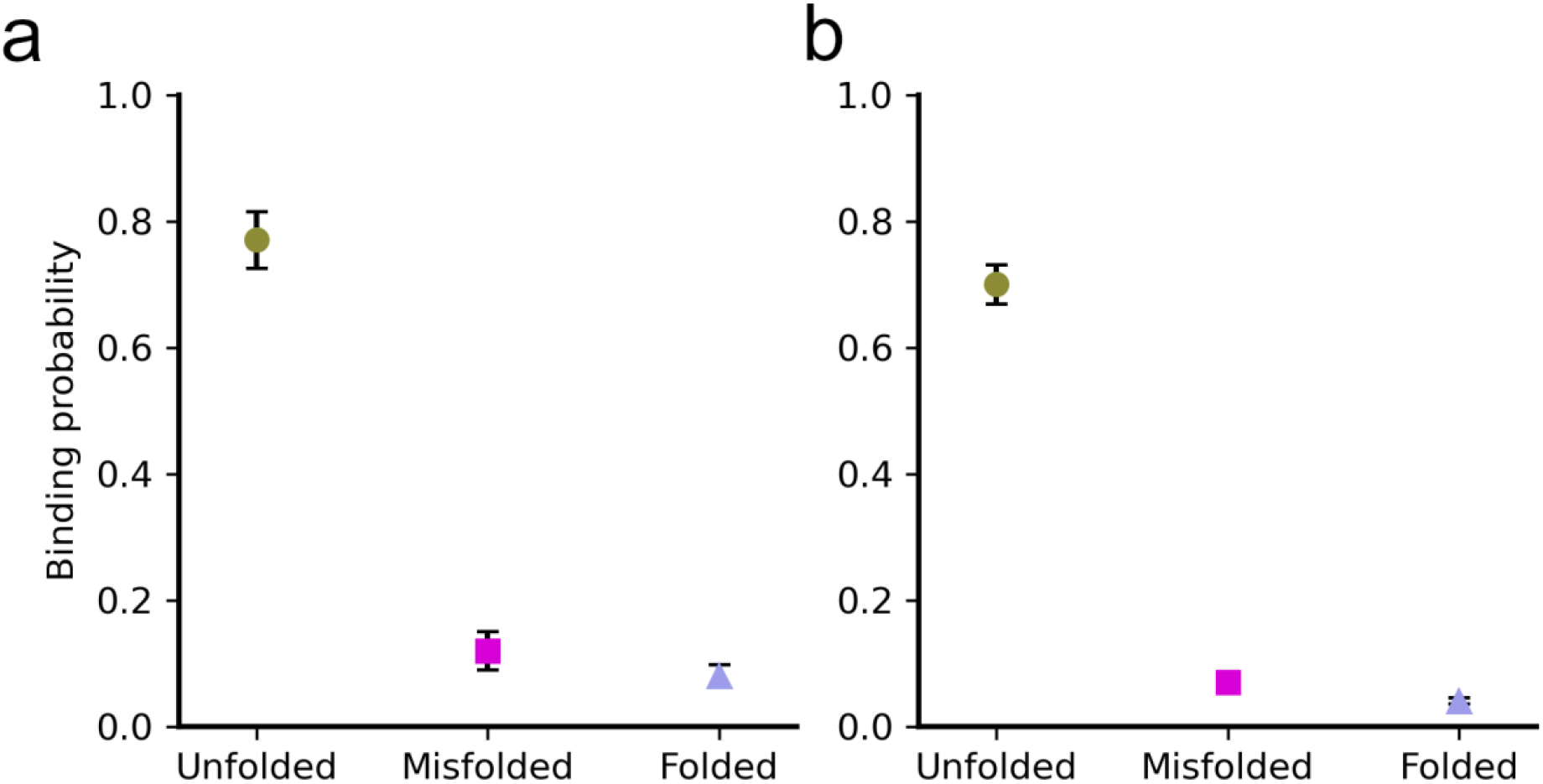
Influence of different contact thresholds on GroEL/Isochorismate synthase binding probabilities. (a) Results when a threshold of ≥1 residues is used. (b) Results when a threshold of ≥5 residues is used. In both cases the unfolded state shows much higher binding probabilities in comparison to the misfolded and folded states. Error bars are 95% CIs computed by bootstrapping 10^6^ times.

**Figure S2.**
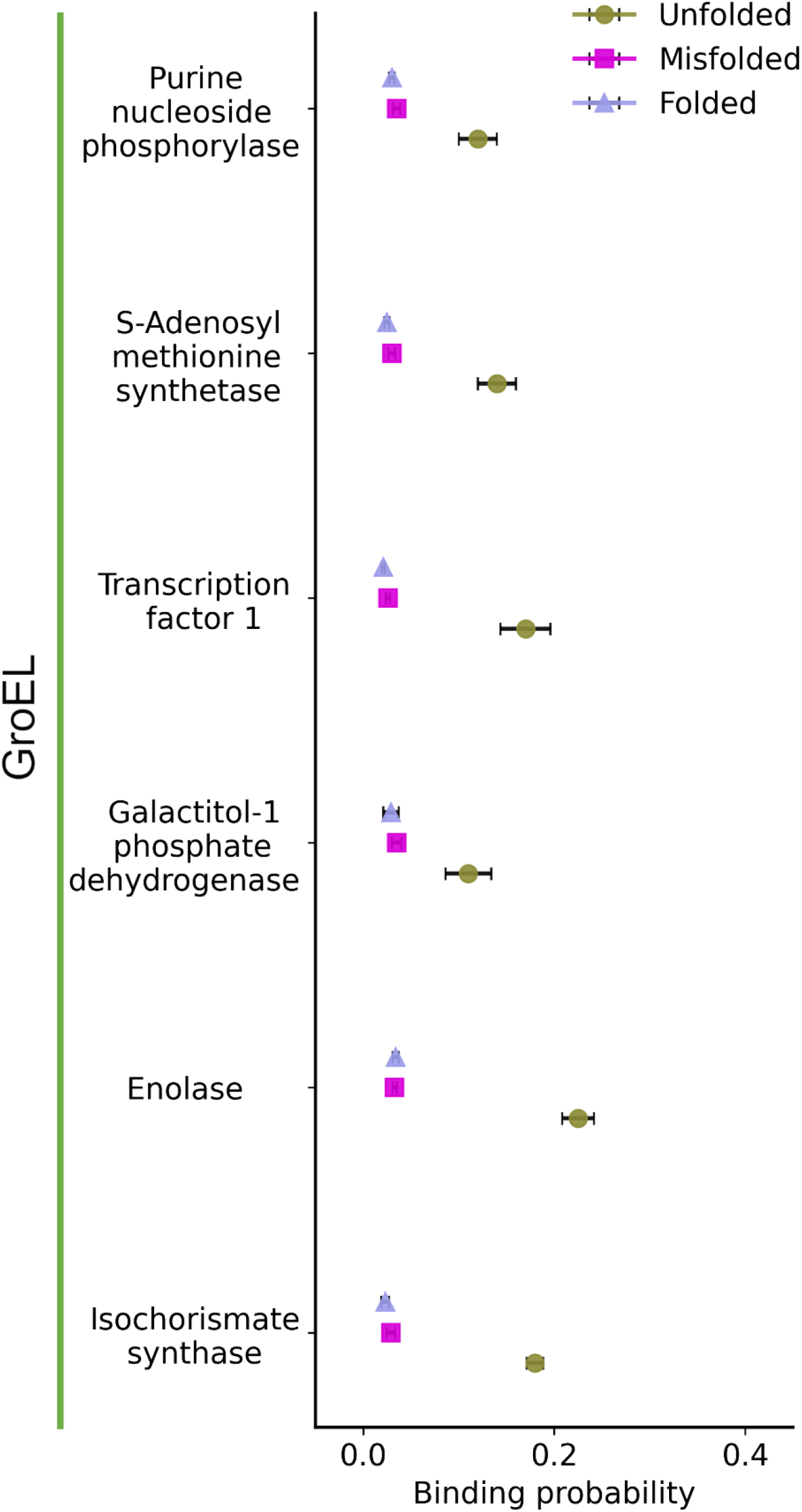
Binding probabilities of unfolded, misfolded, and folded states of client proteins for GroEL in the absence of attractive interactions. In these simulations, interactions between client proteins and GroEL are dependent totally on steric effects. The binding probability of each of the unfolded, misfolded and folded states for each client protein and GroEL was averaged over ten independent simulations. Error bars are 95% CIs computed by bootstrapping 10^6^ times.

**Figure S3.**
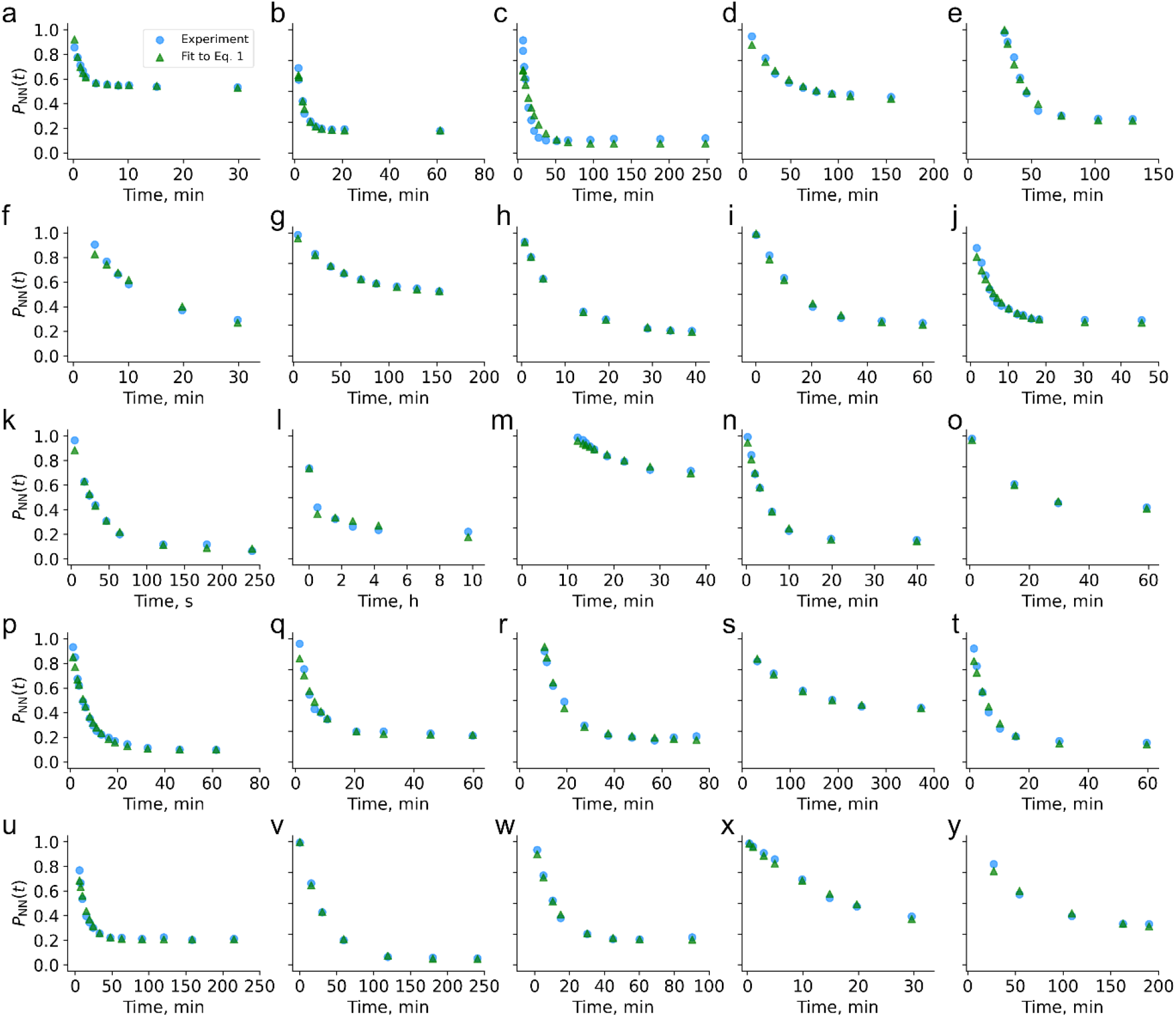
Curve-fitting results for 25 experimental refolding studies. Values of *P*_NN_(*t*) computed as described in the Methods from the original experimental data were fit to Eq. 1 using SciPy in Python3. The resulting fit parameters and Pearson *R*^2^ values are listed in Table S1 while the original figures from which data were extracted with PlotDigitizer are provided in a Supplementary Excel file.

**Table S1.** Results of curve fitting to Eq. 1 for 25 experimental refolding time courses

| Label of plots in Figure S3 | Ref. | $a_0$ | $k_1$ (time<sup>-1</sup>)<sup>†</sup> | $a_1$ | $k_2$ (time<sup>-1</sup>)<sup>†</sup> | Pearson $R^2$ |
| --- | --- | --- | --- | --- | --- | --- |
| a | 1 | 0.43 | 0.99 | 0.57 | $2 \times 10^{-3}$ | 0.98 |
| b | 2 | 0.81 | 0.37 | 0.19 | $1 \times 10^{-4}$ | 0.98 |
| c | 3 | 0.92 | 0.06 | 0.08 | $9 \times 10^{-22}$ | 0.91 |
| d | 4 | 0.57 | 0.02 | 0.43 | $1 \times 10^{-16}$ | 0.98 |
| e | 34 | 0.90 | 0.05 | 0.10 | $1 \times 10^{-20}$ | 0.97 |
| f | 35 | 0.49 | 0.01 | 0.51 | $1 \times 10^{-12}$ | 0.97 |
| g | 36 | 0.80 | 0.01 | 0.20 | $4 \times 10^{-18}$ | 1.00 |
| h | 37 | 0.75 | 0.15 | 0.25 | $1 \times 10^{-2}$ | 1.00 |
| i | 38 | 0.72 | 0.06 | 0.28 | $1 \times 10^{-19}$ | 0.99 |
| j | 39 | 0.01 | 0.06 | 0.11 | $5 \times 10^{-23}$ | 0.98 |
| k | 40 | 0.36 | 0.03 | 0.93 | $7 \times 10^{-4}$ | 0.99 |
| l | 41 | 0.79 | 24.38 | 0.21 | $8 \times 10^{-2}$ | 0.95 |
| m | 42 | 0.62 | 0.06 | 0.38 | $3 \times 10^{-20}$ | 0.98 |
| n | 43 | 0.88 | 0.20 | 0.12 | $4 \times 10^{-18}$ | 1.00 |
| o | 44 | 0.77 | 0.07 | 0.23 | $9 \times 10^{-19}$ | 1.00 |
| p | 45 | 0.76 | 0.14 | 0.24 | $1 \times 10^{-12}$ | 0.99 |
| q | 46 | 0.95 | 0.16 | 0.05 | $2 \times 10^{-23}$ | 0.97 |
| r | 47 | 0.70 | 0.05 | 0.30 | $9 \times 10^{-20}$ | 0.97 |
| s | 48 | 0.80 | 0.01 | 0.20 | $2 \times 10^{-22}$ | 0.99 |
| t | 49 | 0.57 | 0.16 | 0.43 | $3 \times 10^{-14}$ | 0.98 |
| u | 50 | 0.59 | 0.08 | 0.41 | $7 \times 10^{-22}$ | 0.98 |
| v | 51 | 0.90 | 0.02 | 0.10 | $1 \times 10^{-13}$ | 1.00 |
| w | 52 | 0.85 | 0.09 | 0.50 | $3 \times 10^{-20}$ | 1.00 |
| x | 53 | 0.79 | 0.04 | 0.21 | $3 \times 10^{-18}$ | 0.99 |
| y | 54 | 0.73 | 0.01 | 0.27 | $2 \times 10^{-19}$ | 0.98 |

<sup>†</sup>Units are min<sup>-1</sup> for all rates except for sub-panels k and l in Figure. S3, which have units of s<sup>-1</sup> and h<sup>-1</sup>, respectively.

**Table S2.** Identities of the client proteins and their corresponding *η* values

| **Client protein** | **PDB ID** | **Optimum $\eta$ used**<br>**for coarse-**<br>**grained modelling**<br>**of client proteins** |
| --- | --- | --- |
| Isochorismate synthase | 3HWO | 1.359 |
| Enolase | 2FYM | 1.359 |
| Galactitol-1-phosphate-dehydrogenase | 4A2C | 1.359 |
| Transcription factor 1 | 1K7J | 1.359 |
| S-Adenosylmethionine synthetase | 1P7L | 1.359 |
| Purine nucleoside phosphorylase | 1A69 | 1.359 |

**Table S3.** Identities of the chaperones and their corresponding *η* values

| Chaperone | PDB ID | $\eta$ used for coarse-grained modelling |
| --- | --- | --- |
| GroEL | 1KP8 | 1.8 |
| HtpG | 2IOQ | 1.4 |

**Table S4.** *η* values for interactions between client proteins and chaperones

| Chaperone-Client protein Complex | $\eta$ used for chaperone-client protein binding simulations |
| --- | --- |
| GroEL-Isochorismate synthase | 1.4 |
| GroEL-Enolase | 1.4 |
| GroEL-Galactitol-1-phosphate dehydrogenase | 1.4 |
| GroEL-Transcription factor 1 | 1.4 |
| GroEL-S-Adenosine methionine synthetase | 1.4 |
| GroEL-Purine nucleoside phosphorylase | 1.4 |
| HtpG-Purine nucleoside phosphorylase | 1.4 |

**Table S5.** Entanglement (*G*, see Eq. 5) calculated for the of native and near-native like misfolded states of client proteins

| Client protein name | Entanglement in the native state? | $G$ | Entanglement present in near-native misfolded state? | $G$ |
| --- | --- | --- | --- | --- |
| Isochorismate synthase | No | 0.00 | Yes | 0.05 |
| Enolase | No | 0.00 | Yes | 0.09 |
| Galactitol-1-phosphate-dehydrogenase | No | 0.00 | Yes | 0.01 |
| Transcription factor 1 | No | 0.00 | Yes | 0.10 |
| S-adenosylmethionine synthetase | No | 0.00 | Yes | 0.08 |
| Purine nucleoside phosphorylase | No | 0.00 | Yes | 0.09 |

## Notes

### Competing Interest Statement

The authors have declared no competing interest.

