## Supplementary Reference Material 1 for "Subpopulations of soluble, misfolded proteins commonly bypass chaperones: How it happens at the molecular level"

### Universal protein misfolding intermediates can bypass the proteostasis network and remain soluble and non-functional

Daniel Nissley[1,ǂ], Yang Jiang[1,ǂ], Fabio Trovato[1], Ian Sitarik[1], Karthik Narayan[1], Edward P. O'Brien[1,2,3*]

[1] Department of Chemistry, Pennsylvania State University, University Park, PA 16802, USA
[2] Bioinformatics and Genomics Graduate Program, The Huck Institutes of the Life Sciences, Pennsylvania State University, University Park, PA 16802, USA
[3] Institute for Computational and Data Sciences, Pennsylvania State University, University Park, PA 16802, USA
ǂ These authors contributed equally to this research project

#### ABSTRACT

Misfolded protein conformations with decreased functionality can bypass the proteostasis machinery and remain soluble *in vivo*. This is an unexpected phenomenon as several cellular quality control mechanisms have evolved to rid cells of misfolded proteins. Three questions, then, are: how is it structurally possible for long-lived, soluble, misfolded proteins to bypass the proteostasis machinery and processes? How widespread are these soluble, misfolded states across the proteome? And how long do they persist for? Here, we address these questions using coarse-grain molecular dynamics simulations of the synthesis, termination, and post-translational dynamics of a representative set of cytosolic *E. coli* proteins. We find that half of all proteins exhibit subpopulations of misfolded conformations that are likely to bypass molecular chaperones, avoid aggregation, and not be degraded. These misfolded states can persist for months or longer for some proteins. Structurally characterizing these misfolded states, we observe they have a large amount of native structure, but also contain localized misfolded regions from non-native changes in entanglement, in which a protein segment threads through a loop formed by another portion of the protein that is not found in the native state. The surface properties of these misfolded states are native like, allowing them to bypass the proteostasis machinery and processes to remain soluble, while their entanglements make these states long-lived kinetic traps, as disentanglement requires unfolding of already folded portions of the protein. In terms of function, one-third of proteins have subpopulations that misfold into less-functional states that have structurally perturbed functional sites yet remain soluble. These results explain how proteins misfold into soluble, non-functional conformations that bypass cellular quality controls, and indicate that, unexpectedly, this is a wide-spread cellular phenomenon that can lead to reduced protein function across the cytosolic proteome. Such entanglements are observed in many native structures, suggesting the non-native entanglements we observe are plausible. More broadly, these near-native entangled structures suggest a hypothesis for how synonymous mutations can modulate downstream protein structure and function, with these mutations partitioning nascent proteins between these kinetically trapped states.

#### INTRODUCTION

How soluble, misfolded protein populations with reduced functionality[1–3] bypass cellular quality control mechanisms[4] for long time periods is poorly understood. Further, how common this phenomenon is across organismal proteomes has not been assessed. These are important gaps in our knowledge to fill as the answers will offer a more complete picture of protein structure[5] and function in cells, may lead to refinement of the protein homeostasis model[6] of proteome maintenance, are likely to be relevant to how synonymous mutations have long-term impacts on protein structure and function[7], and could reveal long-term misfolding on a scale greater than previously thought.

Protein homeostasis ("proteostasis") refers to the maintenance of proteins at their correct concentrations and in their correct conformational states through the action of a large ensemble of chaperones, proteolytic enzymes, and disaggregation agents[6,8]. This model posits that under normal (*i.e.*, not stressed) cellular growth conditions globular proteins *in vivo* typically attain one of three states: folded/functional, misfolded/aggregated, or degraded. Various molecules work together to maintain proteostasis by catalyzing the interconversion of proteins between these states[9–11]. For example, some chaperones in *E. coli*, such as GroEL/GroES[9] and DnaK[10], catalyze the folding of misfolded or unfolded proteins. Others, such as the set of enzymes associated with the ubiquitin-proteasome system in eukaryotes, covalently tag misfolded proteins for degradation[12]. Yet others, such as *E. coli*'s ClpX, have the potential to break apart aggregates, allowing newly released monomeric proteins to either fold or be degraded[13]. Many caveats and nuances exist in the proteostasis model. For example, some insoluble aggregates, such as carboxysomes, are biologically beneficial by spatially concentrating protein function[14]. Some non-native protein oligomers are soluble[15], and recently discovered biomolecular condensates[16] represent a form of phase separation in which proteins within a condensate are soluble but preferentially interact with each other over other cellular components. Additionally, soluble proteins are not always functional because some require co- or post-translational modifications[17,18].

The timescales involved with many proteostasis processes are often quite short. Co-translationally acting chaperones bind ribosome nascent chain complexes on time scales of 10's of ms[19], the ubiquitin-degradation machinery tags up to 30% of eukaryotic nascent chains for immediate degradation after synthesis[20,21], and post-translationally acting chaperones can fold misfolded proteins in just a few seconds[22]. Indeed, the FoldEco kinetic model of *E. coli* proteostasis indicates that conversions between various states within the network occur with rate constants typically on the order of seconds[23]. Thus, according to the proteostasis model, misfolded proteins will be either converted in a matter of seconds to minutes into their folded state, be degraded, or form aggregates provided cells are not stressed and the proteostasis machinery is not overwhelmed by a high concentration of unfolded proteins[24].

Synonymous mutations change the triplet of nucleotides between degenerate mRNA codons encoding the same amino acid, leading to an altered mRNA sequence that encodes the same protein primary structure. Despite leaving the amino acid sequence unchanged, such mutations can alter the translation-elongation rate of ribosomes and have been found to alter the structure and function of proteins for long timescales[7]. For example, translation of a synonymous variant of the *frq* gene in the fungus *Neurospora* resulted in the synthesis of FRQ protein with altered conformations that bound 50% less to a partner protein, resulting in a significantly altered circadian rhythm that persisted for multiple days[2]. Many other proteins have been reported to exhibit altered structure or function upon the introduction of synonymous mutations[25–27]. The fact that these changes occur in the soluble fraction of the proteome indicates that alterations to translation speed do not cause higher-order oligomer formation. Most importantly, such observations are inconsistent with the proteostasis model, which predicts that any proteins with misfolded and non-functional structures should refold, aggregate, or be degraded on faster timescales.

One hypothesis that would resolve this discrepancy is that proteins can populate a fourth state. In this state, proteins are kinetically trapped over long timescales in misfolded conformations with reduced functionality, but they do not have a propensity to aggregate or interact with proteostasis machinery in excess of that of folded proteins. If correct, this hypothesis raises a number of questions, including: (*i*) what type of structures are adopted in this fourth state? (*ii*) how do those conformations simultaneously avoid folding, aggregation, and degradation in excess of that observed for the native ensemble? (*iii*) how long do they persist? And (*iv*) what fraction of the proteome exhibits this behavior?

To test this hypothesis and address these questions we utilize coarse-grained simulations in which we simulate protein synthesis, co-translational and post-translational folding, and estimate the fraction of molecules that fold, misfold, interact with chaperones, aggregate, are degraded, or attain a functional conformation. Simulating a representative subset of the cytosolic *E. coli* proteome in this way, we discover that a large proportion of newly synthesized proteins can adopt misfolded conformations that are near-native in structure and thus likely to interact with co- and post-translational chaperones in a manner similar to that of the native state. These misfolded conformations expose a similar amount of aggregation-prone surface area as the native ensemble, and therefore do not have an increased propensity to aggregate. For some proteins, misfolding is localized near their functional sites, indicating that they will exhibit decreased biological function. We estimate that many of these near-native misfolded states are kinetically trapped, exhibiting lifetimes on the order of days to months. We discover the common structural feature of these proteome-wide, soluble misfolded states.

#### RESULTS

**A coarse-grain model reproduces experimentally observed misfolding of Firefly Luciferase**. The model we use for protein synthesis, folding, and function has been shown to accurately predict experimentally measured changes in enzyme specific activities[28], indicating it reasonably describes protein structure-function relationships. As an additional test, here we examine if the model is able to identify if a protein will have misfolded subpopulations. Firefly Luciferase, a 550-residue protein with four domains, folds co-translationally[29]. Specific activity experiments[25] have found that some soluble, nascent Luciferase molecules misfold when translation speed is increased. To test our model we simulated Luciferase's synthesis, ejection from the ribosome exit tunnel, and post-translational dynamics using a coarse-grain representation of the protein and ribosome (Figure 1a, b, and c, Table S1, Methods, and Eq. 1 and 2). Fifty statistically independent trajectories were run. To characterize Luciferase's native conformational ensemble, we also simulated ten trajectories initiated from Luciferase's crystal structure, which we refer to as native-state simulations. To assess whether or not a given Luciferase trajectory is misfolded we utilize the time-dependent mode of the fraction of native contacts ($Q_{\mathrm{mode}}$) and the probability that a non-native entanglement ($P(G_k)$) has formed. We consider a trajectory misfolded if it has (*i*) a mean $Q_{\text{mode}}$, over the final 100 ns of the post-translational phase of the simulation, that is less than the average from the native-state simulations, or (*ii*) a mean $P(G_k)$ for the different possible changes in non-covalent lasso threading denoted $k = \{0,1,2,3,4\}$ (see Methods) of 0.1 or greater over the final 100 ns of the trajectory, or (*iii*) both *(i)* and *(ii)* occur (see Methods). Conditions (*i*) and (*ii*) correspond to perturbations of structure relative to the native state as defined by fraction of native contacts and entanglement, respectively. Based on this definition, 46% (95% Confidence Interval [32%, 60%], calculated from bootstrapping $10^6$ times) of nascent Luciferase molecules misfold (Figure 1d and e, Methods). We observe that when Luciferase misfolds, it misfolds 100% of the time in the second domain (based on $\langle Q_{\text{mode}} \rangle$), which is composed of residues 13-52 and 212-355, and 35% of the time at the interface between domains 1 and 2. These misfolded Luciferase structures are near-native, with a 5.5% decrease in the overall fraction of native contacts ($\langle Q_{\text{overall}} \rangle = 0.86$, computed over the final 100 ns of misfolded trajectories) compared to the native ensemble for the same domain ($\langle Q_{\text{overall}} \rangle = 0.91$). Thus, a large proportion of nascent Luciferase misfolds into near-native conformations that typically involve misfolding of the second domain or its interface.

To assess whether these misfolded states are likely to display reduced specific activity *in vivo* we need to determine whether they evade chaperones to remain misfolded, do not aggregate or become degraded such that they remain soluble (as specific activity measurements are carried out on the soluble fraction), and finally if the structure of the residues involved in Luciferase function is perturbed. In *E. coli*, the chaperone trigger factor (TF) binds nascent proteins co-translationally, while GroEL/GroES and DnaK interact with proteins post-translationally. Interactions with TF[30] or GroEL/GroES[9] are thought to occur by the non-specific recognition of exposed hydrophobic patches on client proteins co- and post-translationally, respectively. DnaK, however, interacts with specific binding sites within a protein's sequence[31].

To estimate whether misfolded Luciferase is likely to interact with TF we computed the average relative difference between the hydrophobic solvent accessible surface area (SASA) of each misfolded trajectory to the folded population (denoted $\langle \zeta_{\text{hydrophobic}}^{\text{co-t}} \rangle$, see Methods and Eq. 9) during synthesis. The value of $\langle \zeta_{\text{hydrophobic}}^{\text{co-t}} \rangle$ is ≤10% for 16 of 23 misfolded trajectories, meaning that they display less than a 10% increase in hydrophobic SASA during synthesis relative to the folded population of trajectories. This indicates that a majority of misfolded Luciferase molecules will not interact with TF much more than a properly folded Luciferase molecule. Though TF accelerates protein folding under force[32], under normal conditions it is also thought to act as a holdase[30]. Thus, we conclude that our co-translational Luciferase misfolded states can misfold into conformations that do not interact with TF in a manner that accelerates folding, allowing these misfolded states to persist post-translationally.

Next, to determine whether misfolded conformations of Luciferase are likely to interact with GroEL/GroES post-translationally, we computed the average relative difference between the hydrophobic SASA of each misfolded conformation in the final 100 ns and its value in the native-state simulations ($\langle \zeta_{\text{hydrophobic}} \rangle$, see Methods and Eq. 10). The value of $\langle \zeta_{\text{hydrophobic}} \rangle$ for Luciferase is ≤10% for 9 of 23 misfolded trajectories, indicating that these misfolded states expose only a small excess of hydrophobic SASA relative to the native ensemble, and are therefore not likely to be engaged by GroEL/GroES.

Finally, to estimate whether misfolded Luciferase structures are more likely to interact with DnaK than the native state, we computed $\langle \zeta_{\mathrm{DnaK}} \rangle$ (Eq. 11), the average relative difference in SASA of residues predicted to be DnaK binding sites by the Limbo algorithm[31] in the final 100 ns for all misfolded trajectories. We find that all 23 misfolded trajectories have $\langle \zeta_{\mathrm{DnaK}} \rangle \leq 10\%$, indicating that DnaK is unlikely to preferentially bind to these misfolded states any more than it is to the native state. Thus, some of Luciferase's misfolded states are unlikely to interact with TF, GroEL/GroES, or DnaK, and thus bypass the *E. coli* chaperone network.

The next key question is whether or not these misfolded Luciferase structures, having bypassed chaperone quality controls, are likely to remain soluble or to aggregate or be degraded. To estimate whether misfolded Luciferase will aggregate we used the AMYLPRED2[33] webserver to identify residues in the Luciferase amino acid sequence that lead to aggregation when exposed to solvent. We then computed $\langle \zeta_{\mathrm{agg}} \rangle$ (Eq. 12), the average relative difference in SASA between these aggregation-prone residues in the final 100 ns of each misfolded trajectory in comparison to the same residues in the native-state simulations. We find that 12 of 23 misfolded trajectories have $\langle \zeta_{\mathrm{agg}} \rangle \leq 10\%$, indicating that these misfolded conformations display only a minor increase in aggregation propensity and are likely to remain soluble.

Finally, we considered the likelihood that misfolded Luciferase will be targeted for degradation. Degradation in *E. coli* is carried out by various proteolytic enzymes, including ClpX, that recognize and degrade misfolded or aggregated proteins. It is well established that perturbed protein structures lead to degradation[34]; we therefore use $\langle \zeta_{\mathrm{hydrophobic}} \rangle$ to quantify how similar misfolded Luciferase conformations are to the native state. For 9 of 23 misfolded Luciferase trajectories $\langle \zeta_{\mathrm{hydrophobic}} \rangle$ is $\leq 10\%$, indicating they are unlikely to be degraded more quickly than native Luciferase.

Having determined that some misfolded conformations of Luciferase can evade chaperones, aggregation, and degradation, the final question is whether their function is decreased relative to native Luciferase. To answer this question, we identified the residues that take part in Luciferase's bioluminescence, defined as those resides within 4.5 Å of the 5′-O-[N-dehydroluciferyl)-sulfamoyl]-adenosine ligand in PDB structure 4G36, in addition to all residues identified in the UniProt database[35] to have a role in its catalytic mechanism. To quantify the difference in structure of residues involved in Luciferase's catalytic mechanism, we compute the average relative difference between the structures sampled in the final 100 ns of each misfolded trajectory and native Luciferase in terms of the structural overlap function ($\langle \chi_{\mathrm{func}} \rangle$) over residues implicated in its function (see Eq. 13-15 and Figure 1b, c, f, g, and h). Positive values of $\chi_{\mathrm{func}}$ indicate perturbed structure relative to the native state simulations. We find that 14 of 23 misfolded Luciferase trajectories have $\langle \chi_{\mathrm{func}} \rangle \geq 10\%$, indicating that they have significantly perturbed structure at functionally important sites relative to the native state (Figure 1f). For example, Figures 1g and 1h show the binding pocket at the final frames of folded and misfolded, soluble, but non-functional Luciferase trajectories. The binding pocket structure is perturbed such that it impinges on the substrate location, suggesting this misfolded conformation will have decreased function. Since structure equals function, this result indicates that the efficiency of the enzymatic reaction carried out by misfolded Luciferase will be less efficient than in its native fold.

Cross-referencing the lists of misfolded trajectories that are likely to avoid chaperones, aggregation, degradation, and exhibit reduced function, we find that one trajectory displays all of these characteristics and likely remains soluble but less functional than native Luciferase. Our simulation results are thus qualitatively consistent with the experimental observation that some of nascent Luciferase molecules misfold into a soluble conformation and that these misfolded conformations display reduced function. Our coarse-grain simulation protocol for nascent protein synthesis, ejection, and post-translational dynamics is thus able to recapitulate nascent protein misfolding.

**Simulating a representative subset of the *E. coli* cytosolic proteome**. It is not computationally feasible to simulate all 2,600 cytosolic *E. coli* proteins. Therefore, to investigate the extent of nascent protein misfolding within the *E. coli* proteome we constructed models for a representative subset of 122 proteins. This set of proteins has the same distributions of protein length and structural class, and a similar ratio of multi- to single-domain proteins as the entire *E. coli* proteome[36] (Table S2). The details of the parameterization of these models are described in Ref. 36. Each protein was synthesized on the same coarse-grain ribosome representation as Luciferase and their post-translational dynamics simulated for 30 CPU days per trajectory. Larger proteins take longer to simulate. Therefore, this fixed post-translational simulation run time resulted in trajectories of different durations due to different protein sizes. The simulation time in the post-translational phase therefore ranged in duration between 2.7 and 154.1 $\mu$s per trajectory. Because our coarse-grain model exhibits an approximately four-million fold acceleration of folding dynamics[37], due to decreased solvent viscosity[38] and a smoother free-energy landscape[39], these post-translational simulation times correspond to experimental times of 11.0 to 611 seconds, respectively. As with Luciferase, ten trajectories were also initiated from the crystal structure of each protein and simulated for 30 CPU days to serve as reference simulations representing the native-state structural ensemble.

**Two thirds of nascent *E. coli* proteins populate misfolded states**. A fundamental question our simulation data set can address is how common nascent protein misfolding is across *E. coli*'s cytosolic proteome. As with Luciferase, we use the fraction of native contacts and entanglement as measures of misfolding. We find that 66% of proteins (80 out of 122) remain misfolded in at least one trajectory, 40% of proteins are misfolded in at least 20% of trajectories (49 out of 122), and 7% are misfolded in 100% of trajectories (9 out of 122). The proteins in these various categories are summarized in Table S4; Figure 2a displays a histogram of the probability of misfolding over the 122 different E. coli proteins simulated. In total, 27% of the simulation trajectories (1,631 out of 6,100) of the *E. coli* cytosolic proteome remain in misfolded conformations after 30 CPU days of post-translational dynamics.

**Many misfolded states are similar to the native state.** Misfolded conformations that are very different from the native state will likely interact with the proteostasis machinery. To characterize the closeness of misfolded states to the native state across our set of misfolded conformations we calculated the absolute percent change in the mean overall fraction of native contacts $\langle Q_{\mathrm{overall}} \rangle$ (in this case, computed over all residues in secondary structures within each protein, rather than for individual domains or interfaces) between each protein's native state simulations and the mean Q in the final 10 ns of each misfolded trajectory (Figure 2b). We observe that 76% of misfolded trajectories (1,242 out of 1,631) have ≤20% change in mean Q in comparison to the native state, while 58% of trajectories (939 out of 1,631) misfold and have a ≤10% change in Q. 9% of trajectories (144 out of 1,631) have a ≤1%

change in Q. These calculations indicate that a large proportion of trajectories that misfold populate states that are native-like. Therefore, many *E. coli* proteins are kinetically trapped in near-native conformations that are structurally similar to the native state.

**Misfolded states can persist for days or longer after release from the ribosome.** Misfolded conformations that persist for just a few seconds before properly folding are unlikely to have downstream consequences in a cell. To estimate the range of folding times for misfolded conformations we computed the survival probability of the unfolded state, $S_{\mathrm{U}}(t)$, for each protein domain and interface and extracted their characteristic folding timescales using a three-state folding model, which reports folding timescales for the fast- and slow-folding phases (see Methods). For a protein to be folded all its component domains and interfaces must be folded. Furthermore, since folding pathways that pass through misfolded states take longer to reach the native state the slow-folding phase reflects the time scale of these pathways. Therefore, for a given protein, we interpret the longest, slow-folding phase time as the time scale of the misfolded state reaching the native state. In total, we are able to reliably determine folding times for 73 out of 122 proteins, with fit equations for other domains having small Pearson $R^2$ values indicative of low-quality estimates. These extrapolated folding times for the slow phase were then mapped onto experimental times using the acceleration factor associated with the coarse-grained model[37] (see Methods). The 25$^{\text{th}}$, 50$^{\text{th}}$, 75$^{\text{th}}$, and 95$^{\text{th}}$ percentile mean folding times for the slow phase are 1.41 s, 50.9 s, 1.19 x $10^7$ d, and 3.83 x $10^{16}$ d, respectively, and the full range of times extend from 0.04 s to 1.08 x $10^{22}$ d (Figure 2c, Table S5). While values at very long times have larger uncertainties, as small differences in the fit parameters will lead to large variation in the extrapolated folding times, these results clearly indicate that many of these misfolded states can persist for many days or months after synthesis.

**Half of the proteome misfolds and bypasses the chaperone machinery.** Misfolded proteins are engaged by various chaperones both co- and post-translationally that help direct their correct folding. So, we next determined how many of the trajectories that exhibit misfolding in our simulations are likely to be evade chaperone-dependent quality control mechanisms. As was done for Luciferase, we considered the interactions of each of our 1,631 misfolded trajectories with TF, GroEL/GroES, and DnaK based on the relative difference between the SASA of specific subsets of residues in the misfolded ensemble versus the native state ensemble (see Methods, Eqs. 9-11). We find that 1,053 misfolded trajectories, representing 70 unique proteins, are not likely to interact with TF, as they display $\langle \zeta^{\text{co-t}}_{\text{hydrophobic}} \rangle \leq 10\%$ or are too short to engage with it co-translationally (see Methods, Table S6, and Figure S1). A total of 1,411 of misfolded trajectories representing 80 unique proteins are either not known GroEL/GroES[40–42] clients or have $\langle \zeta_{\text{hydrophobic}} \rangle \leq 10\%$, and are therefore not likely to interact excessively with GroEL/GroES (Table S6, Figure 2d). Finally, we find that 1,115 misfolded trajectories representing 74 unique proteins are either not confirmed DnaK clients or have $\langle \zeta_{\text{DnaK}} \rangle \leq 10\%$ and are therefore unlikely to interact with DnaK excessively (Figure S2). A total of 731 trajectories representing 64 different proteins are misfolded and unlikely to interact with TF, GroEL/GroES, or DnaK (Table S6). These results indicate that 52% of proteins (64 out of 122 unique proteins) exhibit misfolded subpopulations that can bypass chaperones.

**Half of the proteome misfolds and remains soluble.** We next assessed how many of the 1,631 trajectories in which the protein misfolds represent conformational states that are likely to remain soluble. For each protein we computed $\langle \zeta_{\text{agg}} \rangle$, the average relative difference in SASA of residues predicted to be aggregation prone computed over the final 100 ns for each misfolded trajectory, to quantify the difference in aggregation propensity for the misfolded population relative to the native state simulations (see Methods, Eq. 12). Of the 1,631 misfolded trajectories, 814 have with $\langle \zeta_{\text{agg}} \rangle \leq 10\%$, indicating they are not likely to aggregate in excess of what is observed for the native state (Table S7, Figure S3). We conclude that these trajectories, representing 56% of the proteome dataset (68 out of 122), are thus unlikely to aggregate.

**Half of the proteome misfolds and does not exhibit excess degradation.** Next, we examined how many misfolded proteins are likely to avoid rapid degradation. We did this by computing $\langle \zeta_{\text{hydrophobic}} \rangle$, which characterizes the percent difference between the total hydrophobic SASA of misfolded trajectories in comparison to the set of native-state simulations (Eq. 10). The values of $\langle \zeta_{\text{hydrophobic}} \rangle$ for 896 misfolded trajectories are ≤10%, indicating they are unlikely to be targeted for degradation. These 896 misfolded trajectories predicted to bypass degradation represent 57% (70 out of 122) unique proteins (Table S7, Figure 2d). Thus, a majority of proteins can populate, to varying degrees, misfolded states that do not exhibit degradation in excess of those proteins that are natively folded.

**Half of the proteome misfold into conformations that bypass all aspects of the proteostasis machinery in *E. coli*.** Misfolded conformations that do not engage chaperones, do not aggregate, and are not degraded in excess of the native state will remain soluble within the cell for a similar time scale as the native state. We cross-referenced our lists of misfolded trajectories that fall into each of these categories, finding that 47% of proteins (57 out of 122) and 8% of all trajectories simulated (517 out of 6,100) misfold into such soluble conformations (Table S8). Thus, a near majority of proteins have subpopulations of misfolded states that will bypass all aspects of protein homeostasis, and stay misfolded for biologically long time periods.

**Half of the proteome misfolds and will exhibit altered function.** Next, we examined what percentage of the proteome misfolds and is likely to exhibit altered function. To answer this question, we constructed a database identifying residues that take part in the function of each of the 122 proteins in our data set based on information available in PDB and UniProt database entries. These functional residues were identified based on whether they were in contact with substrates (such as other biomolecules, small-molecule compounds, or ions) in their PDB structures, as well as based on UniProt's identification of functional residues (see Methods). We then computed the mean relative difference in the structural overlap function of these functional residues in the final 100 ns of misfolded trajectories relative to the native state reference simulations ($\langle \chi_{\text{func}} \rangle$, see Methods). We find that 62% of misfolded trajectories (1,019 out of 1,631) have $\langle \chi_{\text{func},} \rangle \geq 10\%$, indicating that structure at their functional loci is significantly perturbed and consequently their function is also likely to be perturbed. These trajectories represent misfolded conformations of 69 unique proteins, indicating that 57% of the proteome can populate misfolded conformations likely to exhibit reduced function (Table S8, Figure 2e).

**One-third of proteins exhibit soluble, misfolded, native-like states with reduced functionality.** We next determined which of our 122 proteins misfold, evade chaperones, aggregation, degradation, and display reduced function. We find that 31% of proteins (38 out of 122) and 3% of all trajectories (186 out of 6,100) can bypass proteostasis machinery and display decreased function (Table S8, Figure 2f). The extrapolated folding times of these 38 soluble but non-functional proteins range from 2.13 s to 1.07 x $10^{22}$ days with a median predicted folding time of 1.05 x $10^{12}$ days, indicating that their function is likely to be perturbed for long timescales.

**Intra-molecular entanglement drives long-lived, soluble misfolded conformations.** To determine what characteristics, if any, the misfolded conformations of different proteins have in common, we used the Gauss linking number calculated from linking between a closed loop formed by a native contact between residues *i* and *j* and the pseudo-closed loops formed by the flanking termini, $g(i,j)$[43]. This quantity provides a useful measure of whether subsections of the protein chain are entangled with each other[44]. Misfolding, or changes in the linkage between the two closed loops, can then be identified by changes in the Gauss linking number of specific native contacts between a reference structure and a target structure (Figure 3a, b).

To determine whether or not misfolded states tend to be entangled, we generated a 2x2 contingency table (Figure 3c) tabulating the co-occurrence of misfolding based on $\langle Q_{\mathrm{mode}} \rangle$ and the presence of entanglement. We find an odds ratio of 48.1 ($p < 1 \times 10^{-10}$, Fisher's Exact test), indicating that entanglement and misfolding almost always co-occur, with few instances of misfolding that do not include entanglement. Thus, misfolding is primarily driven by entanglement of segments of the nascent protein with each other.

We hypothesized that due to the large energetic barrier needed to disentangle entanglements, the most long-lived misfolded states in the *E. coli* proteome would tend to be entangled. To test this hypothesis we generated a second 2x2 contingency table and counted how frequently slow- and fast-folding proteins tend to be entangled (Figure 3d). Proteins with an extrapolated folding time for the slow phase greater than the median were considered to be slow-folding; a protein's misfolded states are considered entangled if ≥50% of its misfolded trajectories display entanglement. We find an odds ratio of 15.1 ($p = 5.0 \times 10^{-7}$, Fisher's Exact test) indicating that the presence of entangled misfolded structures are 15 times more likely to be associated with slow folding. Thus, entanglement is the primary cause of long-lived misfolded states.

Taken together, these results demonstrate that misfolding is primarily driven by the formation of entanglements, and that these entangled misfolded states lead to long-lived kinetic traps.

**An in-depth case study.** To illustrate our key findings it is useful to consider one protein in-depth. We focus on glycerol-3-phosphate dehydrogenase. This protein consists of two domains composed of residues 1-387 and 388-501, respectively (Figure 4a). As part of its biological function, glycerol-3-phosophate dehydrogenase must chemically modify the flavin adenine dinucleotide ligand (Figure 4a, dark blue). In our post-translational simulations 74% (=37/50) of trajectories remain misfold in its first domain, yet only exhibit a 4.7% decrease in the fraction of native contacts relative the native-state simulations (Figures 4b, c). Thus, these misfolded states resemble the native ensemble. This protein also folds extremely slowly, with Domain 1, Domain 2, and the interface between Domain 1 and 2 estimated to require, respectively, on the order of $10^{16}$, $10^{15}$, and $10^{21}$ seconds to fold (Figure 4d). These misfolded states are, however, largely expected to evade chaperones, aggregation, and degradation to remain soluble based on the similarity of their surface properties to that of the native ensemble (Figure 4e). Twenty-six misfolded trajectories (52%) also exhibit notably reduced structure at functional sites, including around the flavin adenine dinucleotide substrate, despite being well folded overall (Figure 4f, g, and h). In 92% (=34/37) of these misfolded trajectories a non-native entanglement is present. These results exemplify how entangled misfolded states can perturb portions of a protein critical for function yet can remain soluble for long timescales (Figure 4d and h).

#### DISCUSSION

Soluble, long-lived, non-functional protein misfolded states are observed to occur based on changes in protein function arising from alteration of translation-elongation kinetics. This is the first study to estimate the extent of this phenomenon across the nascent proteome of an organism and examine the structural and kinetic properties of these kinetically trapped states. We have found that a majority of cytosolic *E. coli* proteins exhibit subpopulations of misfolded, kinetically trapped states, and that many of these misfolded states are similar enough to the native state to evade the various quality control mechanisms of the proteostasis machinery in *E. coli*. We estimate that one-third of cytosolic *E. coli* proteins have subpopulations that misfold into near-native conformations that have reduced function and bypass the proteostasis network to remain soluble and non-functional anywhere for days or longer.

To appreciate these results, it is useful to understand the types of misfolding that can and cannot occur in our simulation model. The coarse-grain forcefield is parameterized for each protein based on its crystal structure, with this native-state conformation encoded as the potential energy minimum in the form of a Gō-based energy function (Eq. 1). This means that the native state is the global free energy minimum at our simulation temperatures; any other state is metastable. Another consequence of this type of model is that misfolding involving non-native tertiary structure formation is not possible. Thus, the misfolded states our model can populate are topologically frustrated states that are kinetic traps[45]. A kinetic trap is a local minimum separated from other conformations in the ensemble by energy barriers much larger than thermal energy, making the attainment of the native state a slow process for some protein subpopulations. In our model, intra-molecular entanglements can occur. These entanglements consist of two parts: a contiguous segment of the protein that forms a 'closed' loop, where the loop closure is geometrically defined as a backbone segment that has a native contact between two residues at its ends, and another segment of the protein that threads through this loop (Fig. 3).

It was previously hypothesized[46] that like protein topological knots[47] (which persist when pulling on both termini), this type of entanglement, which we refer to as a non-covalent lasso entanglement[48], would generate topological frustration and be a kinetic trap. Simulations of proteins with topological knots in their native state[45] observed that the wrong knot could form and that many of these states were long-lived kinetic traps as they required 'backtracking'[49] (*i.e.*, unfolding) to fix the knot. While only 3 of the 122 proteins in our study contain topological knots in the native state, the non-covalent lasso entanglement intermediates we observe are non-native pseudoknots[47] (which unravel when pulling on both termini), and require either reptation of the threaded protein segment out of the closed loop (Fig. 3) or unfolding of the loop surrounding the threaded segment to disentangle. Thus, the results of this study bridge the rich field of polymer topology with biologically important consequences for in vivo protein structure and function.

A potential criticism of this work is that the non-native entanglements we observe may just be an artifact of our coarse-grained modeling of proteins. Several lines of evidence indicate this criticism is unfounded. A sufficiently long linear polymer performing a random walk will always sample knotted structures. Thus, it is a fundamental polymer property that knots and entanglements have the potential to form[50]. In a recent study, four entangled structures produced from our coarse-grained model were back-mapped and simulated using classical, all-atom molecular dynamics[51]. The entanglements, and native-like structure of these states persisted for the entire 1 μs simulation time. In another study, one-third of the protein crystal structures in the CATH database were found to contain in the native state the same types of entanglements we observe as intermediates[52]. Taken together, these results indicate that the entanglements we observe are realistic non-native intermediates that have the potential to be populated by many proteins.

The majority of proteins in this study misfold in some capacity. And 38 unique proteins, or about one-third of the proteins we simulated, have one or more trajectories that remain soluble and non-functional due to misfolding. Projecting this proportion across the entire set of 2,600 proteins that make up the cytosolic *E. coli* proteome, we estimate that approximately 806 proteins may exhibit misfolding into soluble states. Given that a reduction in the function of a protein has the potential to influence multiple cellular processes, this result suggests that these misfolded states could exert wide-spread influences on cell behavior and phenotype.

One of the most fundamental timescales of a protein is its half-life, which gives a measure of the lifetime of a typical copy of a protein between its synthesis and degradation. If misfolded states persist on the same timescale as the protein half-life then protein function will be perturbed for most of that protein's existence. Unfortunately, we are unaware of any proteome-wide studies of protein half-lives in *E. coli*. However, based on studies of protein lifetimes in budding yeast[53] and human cells[54], which found median half-lives of 43 min (range 2 min to 81 days) and 36 h (range: 8 h to 153 days), respectively, we can estimate that typical half-lives in *E. coli* range from minutes to hours. Many of our extrapolated folding times are on the same order of magnitude as these values or greater, indicating that misfolded but non-functional states can persist for the entire lifetime of a protein. This is consistent with the experimental observation that misfolding can influence folding and function for extended periods[1,2,55].

If the half-life is a fundamental time scale of a protein, then the cell-division time is a fundamental time scale of a bacterium. During cell division, binary fission of the mother cell results in the formation of two genetically identical daughter cells that each contain roughly half of the copies of each protein from the mother cell. Cell-division timescales for bacteria are typically expressed in terms of the doubling time, which is the time required for the number of cells to double. In *E. coli*, doubling times during exponential growth phase range from 10s of minutes to hours depending on the growth medium[56]. A total of 31 of our 122 proteins have extrapolated folding times for the slow phase longer than 40 min. And, of the 38 proteins that misfold into soluble but non-functional states, 14 have times longer than 40 min. Since these folding times are on a similar time scale as the doubling time, soluble misfolded conformations will be split between the daughter cells. This suggests that the memory of those events can be encoded in these kinetically trapped states and transferred to the daughter cells. It will be an interesting area of future study to determine whether inheritance of soluble, non-functional proteins can act as a mechanism for epigenetic inheritance and influence daughter cell behavior.

A key question answered by our results is what allows these misfolded states to remain misfolded in non-functional states for such long timescales. Entanglements allow these misfolded states to persist for long time scales, and their largely native topologies mean they are not excessively acted upon by the proteostasis machinery. In many instances, large-scale unfolding would need to take place in order for the entangled protein to disentangle[45] to a state from which the native fold is more readily accessible.

In summary, we have found that the majority of *E. coli* proteins misfold, and that some proteins misfold into states that likely bypass cellular proteostasis machinery to remain soluble but non-functional. We find that these misfolded conformations are able to remain soluble because they are, overall, very similar to the native state, but with certain entanglements that lead to perturbed structure and function. Given that self-entanglement is a fundamental polymer property, the entanglements we have observed may represent a universal type of misfolding that has the potential to impact a range of protein functions. Specifically, our results suggest the hypothesis that entangled states may be the source of reduced dimerization[2], enzymatic function[57], and small-molecular transport[27] upon changes in translation kinetics induced by synonymous mutations. Future theoretical and experimental efforts should focus on the structural characterization of non-native entangled states and their influence on protein function.

#### METHODS

**Selection of proteins and parameterization of their coarse-grain models.** A data set of 50 multi- and 72 single-domain proteins was selected at random from a previously developed database of *E. coli* proteins with solved X-ray diffraction or NMR structures[36,58]. This data set contains proteins with realistic distributions of protein size and structural class (see Figure S1 and Table S1, respectively, of Ref. 36). Small sections of missing residues (<10) were rebuilt and minimized in CHARMM, while large missing sections for some multi-domain proteins were rebuilt based on homologous protein structures (see Table S4 of Ref. 36). Each of the rebuilt all-atom models was then converted to a $C_\alpha$ coarse-grain representation. The potential energy forcefield of this coarse-grain model is given by the equation

$$E = \sum_i k_\mathrm{b}(r_i - r_0)^2 + \sum_i \sum_j^4 k_{\varphi,ij}\left(1 + \cos\left[j\varphi_i - \delta_{ij}\right]\right) + \sum_i -\frac{1}{\gamma}\ln\left\{\exp\left[-\gamma\left(k_\alpha(\theta_i - \theta_\alpha)^2 + \varepsilon_\alpha\right)\right] + \exp\left[-\gamma k_\beta\left(\theta_i - \theta_\beta\right)^2\right]\right\} + \sum_{ij} \frac{q_i q_j e^2}{4\pi\varepsilon_0\varepsilon_\mathrm{r} r_{ij}} \exp\left[-\frac{r_{ij}}{l_\mathrm{D}}\right] + \sum_{ij\,\in\,\{\mathrm{NC}\}} \epsilon_{ij}^\mathrm{NC}\left[13\left(\frac{\sigma_{ij}}{r_{ij}}\right)^{12} - 18\left(\frac{\sigma_{ij}}{r_{ij}}\right)^{10} + 4\left(\frac{\sigma_{ij}}{r_{ij}}\right)^{6}\right] + \sum_{ij\notin\{\mathrm{NC}\}} \epsilon_{ij}^\mathrm{NN}\left[13\left(\frac{\sigma_{ij}}{r_{ij}}\right)^{12} - 18\left(\frac{\sigma_{ij}}{r_{ij}}\right)^{10} + 4\left(\frac{\sigma_{ij}}{r_{ij}}\right)^{6}\right]. \quad [1]$$

These forcefield terms represent, from left to right, the contributions from $C_\alpha - C_\alpha$ bonds, torsion angles, bond angles, electrostatic interactions, Lennard-Jones-like native interactions, and repulsive non-native interactions to the total potential energy. Full details of the model parameters can be found in Ref. 36. The value of $\epsilon_{ij}^\mathrm{NC}$, which determines the global energy minimum for a native contact, is calculated as

$$\epsilon_{ij}^\mathrm{NC} = n_{ij}\epsilon_\mathrm{HB} + \eta\epsilon_{ij}. \quad [2]$$

In Eq. 2 $\epsilon_{\mathrm{HB}}$ and $\epsilon_{ij}$ represent energetic contributions from the hydrogen bonds and van der Waals interactions between residues $i$ and $j$ found within the all-atom structure of the protein, respectively. $n_{ij}$ is the integer number of hydrogen bonds between residues $i$ and $j$ and $\epsilon_{\mathrm{HB}} = 0.75\ \mathrm{kcal/mol}$. The value of $\epsilon_{ij}$ is initially set based on the Betancourt-Thirumalai potential[59] and the value of $\eta$ for each individual domain and interface set based on a previously published training set[37]. The values of $\eta$ used for all production simulations are listed in Ref. 36 Tables S2 and S3 alongside all protein names and the chain identifiers used during model building. For simplicity, all proteins are referred to using the PDB ID of the entry from which they were primarily derived. The parameters for the coarse-grain model of firefly luciferase (PDB ID: 4G36) have not previously been reported and are therefore provided in Table S1.

**Simulations of nascent protein synthesis, ejection, and post-translational dynamics.** All simulations were performed using CHARMM and the coarse-grain forcefield described in Eq. 1 with an integration timestep of 0.015 ps and a Langevin integrator with friction coefficient of 0.050 $ps^{-1}$ at a temperature of 310 K. The synthesis and ejection of each protein was simulated using a previously published protocol and a coarse-grain cutout of the ribosome exit tunnel and surface (for complete simulation details see Ref. 36). In this model, ribosomal RNA is represented by one interaction site each for each ribose sugar, phosphate group, and pyrimidine base and two interaction sites for each purine bases. Ribosomal proteins are represented at the $C_\alpha$ level. Post-translational dynamics simulations were initiated from the final protein structure obtained after ejection with the ribosome deleted. Fifty statistically independent trajectories were run for each of the 122 proteins in the *E. coli* cytosolic proteome data set and for Luciferase. Post-translational dynamics was run for 30 CPU days for each trajectory. Ten trajectories were also initiated from the native-state coordinates for all proteins and run for 30 CPU days each to provide a realistic reference ensemble for each protein's folded state.

**Identification of misfolded trajectories.** Two order parameters, $Q_{\mathrm{mode}}$ and $P(G_k \mid PDB,\ traj)$, were used to determine whether or a not a given trajectory folds. Detailed definitions of these order parameters are given in the following two Methods sections. A given trajectory is considered to be misfolded if either its $Q_{\mathrm{mode}}$ or $f_c(G_k \mid PDB, traj)$ values (or both, as described below) indicate that the trajectory is significantly different from the native state reference simulations.

**Calculation of $Q_{\mathrm{mode}}$ and its use as an order parameter for protein folding.** The fraction of native contacts, $Q$, was calculated for each domain and interface of all 122 proteins during their synthesis, ejection, and post-translational dynamics. Only contacts between pairs of residues both within secondary structural elements as identified by STRIDE[60] based on the final rebuilt all-atom structures were considered. To determine when a given domain or interface within a protein folded, the mode of the $Q$ values over a 15-ns sliding window ($Q_{\mathrm{mode}}$) was compared to the representative value of the native state computed as the average $Q_{\mathrm{mode}}$ over all windows of the ten native-state simulations denoted ($\langle Q_{\mathrm{mode}}^{\mathrm{NS}} \rangle$). A given trajectory is defined as misfolded if its average $Q_{\mathrm{mode}}$ over the final 100 ns of the post-translational dynamics portion of the simulation, denoted $\langle Q_{\mathrm{mode}}^{\mathrm{NS}} \rangle$, is less than $\langle Q_{\mathrm{mode}}^{\mathrm{NS}} \rangle - 3\sigma$, where $\sigma$ is the standard deviation of $\langle Q_{\mathrm{mode}}^{\mathrm{NS}} \rangle$.

**Generation of entanglement metric distributions and use as an order parameter for protein folding**. To detect non-covalent lasso entanglements we use linking numbers[61]. A link is defined as the entanglement of two closed curves; here, we use (1) the closed curve composed of the backbone trace connecting residues $i$ and $j$ that form a native contact and (2) the open curves formed by the terminal tails. The native contact between $i$ and $j$ in (1) is considered to close this loop, even though there is no covalent bond between these two residues. Outside this loop is an N-terminal segment, composed of residues 1 through $i-1$, and a C-terminal segment, composed of residues $j+1$ through $N$, whose entanglement through the closed loop we characterize with partial linking numbers denoted $g_{\mathrm{N}}$ and $g_{\mathrm{C}}$[43]. For a given structure of an *N*-length protein, with a native contact present at residues $(i,j)$, the coordinates $\boldsymbol{R}_l$ and the gradient $\mathrm{d}\boldsymbol{R}_l$ of the point $l$ on the curves were calculated as

$$\begin{cases} \boldsymbol{R}_l = \dfrac{1}{2}(\boldsymbol{r}_l + \boldsymbol{r}_{l+1}) \\ \mathrm{d}\boldsymbol{R}_l = \boldsymbol{r}_{l+1} - \boldsymbol{r}_l \end{cases}, \tag{3}$$

where $\boldsymbol{r}_l$ is the coordinates of the C$_\alpha$ atom in residue $l$. The linking numbers $g_{\mathrm{N}}(i,j)$ and $g_{\mathrm{C}}(i,j)$ were calculated as

$$\begin{cases} g_{\mathrm{N}}(i,j) = \dfrac{1}{4\pi}\displaystyle\sum_{m=6}^{i-5}\sum_{n=i}^{j-1}\dfrac{\boldsymbol{R}_m - \boldsymbol{R}_n}{|\boldsymbol{R}_m - \boldsymbol{R}_n|^3}\cdot(\mathrm{d}\boldsymbol{R}_m \times \mathrm{d}\boldsymbol{R}_n) \\ g_{\mathrm{C}}(i,j) = \dfrac{1}{4\pi}\displaystyle\sum_{m=i}^{j-1}\sum_{n=j+4}^{N-6}\dfrac{\boldsymbol{R}_m - \boldsymbol{R}_n}{|\boldsymbol{R}_m - \boldsymbol{R}_n|^3}\cdot(\mathrm{d}\boldsymbol{R}_m \times \mathrm{d}\boldsymbol{R}_n) \end{cases}, \tag{4}$$

where we excluded the first 5 residues on the N-terminal curve, last 5 residues on the C-terminal curve and 4 residues before and after the native contact for the purpose of eliminating the error introduced by both the high flexibility and contiguity of the termini and trivial entanglements in local structure. The above summations yield two non-integer values, and the total linking number for a native contact $(i,j)$ was therefore estimated as

$$g(i,j) = \mathrm{round}\big(g_{\mathrm{N}}(i,j)\big) + \mathrm{round}\big(g_{\mathrm{C}}(i,j)\big), \tag{5}$$

Comparing the absolute value of the total linking number for a native contact $(i,j)$ to that of a reference state allows us to ascertain a gain or loss of linking between the backbone trace loop and the terminal open curves as well as any switches in chirality. Therefore, there are 6 change in linking cases we should consider (Table S3) when using this approach to quantify entanglement.

To examine the distribution of change in linking (entanglement) detected for a given protein model and statistically independent post-translational trajectory we can generate a discrete probability distribution of the 6 cases in Table S3 as

$$P(G_k \mid PDB, traj) = \frac{N_k}{N_E} \tag{6}$$

Where $G_k$ for $k \in \{0,1,2,3,4,5\}$ is the change in entanglement case of interest from Table S3, $N_E$ is the total number of changes in entanglement instances detected in the trajectory, and $N_k$ is the total number of changes in entanglement in the trajectory of type $k$. As the change in entanglement is held relative to the static crystal structure, it is necessary to correct the post-translational distribution to remove transient changes in entanglement present in the reference state dynamics. This was done by subtraction of the reference distribution from the post-translational distribution considering nontrivial changes in linkage:

$$f_c(G_k \mid PDB, traj) = \left| P_{pt}(G_k \mid PDB, traj) - P_{ref}(G_k \mid PDB) \right|, \quad k \in \{0,1,2,3,4\}. \quad [7]$$

For a given case of entanglement change $k$ the magnitude of $f_c(G_k \mid PDB, traj)$ increases as the probability of that mode of entanglement change deviates from the reference simulations. We define a given trajectory as misfolded if the average of any of its corrected entanglement values from Eq. 7 over the final 100 ns of the trajectory are ≥0.1.

This trajectory-level analysis is useful for classifying statistically independent sample sets by the level and types of changes in entanglement they exhibit, but a time series metric which conveys the same information was desired to allow for folding time extrapolations. *G* is a time dependent order parameter that reflects the extent of the topological entanglement changes in a given structure compared to the native structure and is calculated as

$$G(t) = \frac{1}{N} \sum_{(i,j)} \Theta\left((i,j) \in nc \cap g(i,j,t) \neq g^{\text{native}}(i,j)\right), \quad [8]$$

where (*i*, *j*) is one of the native contacts in the native crystal structure; *nc* is the set of native contacts formed in the current structure at time $t$; $g(i,j,t)$ and $g^{\text{native}}(i,j)$ are, respectively, the total entanglement number of the native contact (*i*, *j*) at time $t$, and native structures estimated using Eq. 5; *N* is the total number of native contacts within the native structure and the selection function $\Theta$ equals 1 when the condition is true and equals 0 when it is false. The larger *G* is, the greater the number of native contact residues that have changed their entanglement status relative to the native state. The utility of entanglement for detecting structural perturbations not apparent by fraction of native contacts or root mean square deviation is visually described in Figure S4.

**Calculation and extrapolation of folding times.** Folding times were determined for each domain and interface from their post-translational $Q_{\text{mode}}$ and $G$ time series. The folding time for a domain or interface is taken as the first $t$ at which $Q_{\text{mode}}$ is greater than or equal to $\langle Q_{\text{mode}}^{\text{NS}} \rangle - 3\sigma$ and with $G \leq G_{\text{xs}}$. The survival probability of the unfolded state was then computed based on these folding times and fit to the double-exponential function $S_U(t) = f_1 \exp(k_1 * t) + f_2 \exp(k_2 * t)$ with $f_1 + f_2 \equiv 1$. This double-exponential fit equation represents a kinetic scheme in which the unfolded state first passes to an obligate intermediate before proceeding irreversibly to the folded state. Folding times for each kinetic phase were computed as $\tau_{\text{F},1} = \frac{1}{k_1}$ and $\tau_{\text{F},2} = \frac{1}{k_2}$, with the overall folding time of the domain or interface taken as the longer of the two folding times. The folding time reported for each protein is the longest folding time from any of its constituent domains or interfaces. The assumption of double-exponential folding kinetics is not a good assumption for all 122 proteins simulated. We therefore only consider domains and interfaces whose $S_U(t)$ time series are fit with a Pearson $R^2 > 0.90$. We are also unable to compute folding times for the nine proteins for which no trajectories folded. In total, we found reliable folding times for 73 of our 122 proteins.

**Identifying misfolded proteins unlikely to interact with trigger factor.** To determine whether or not a given protein in our data set is likely to interact with trigger factor (TF) during synthesis we computed the relative difference in the hydrophobic SASA between misfolded trajectories and folded trajectories on the ribosome using the equation

$$\zeta_{\text{hydrophobic}}^{\text{co-t}}(t,l) = \left(\frac{A_{\text{hydrophobic}}(t,l)}{\langle A_{\text{hydrophobic}}(t,l)\rangle_{\text{F}}} - 1\right) * 100\%. \quad [9]$$

In Eq. 9, $A_{\text{hydrophobic}}(t,l)$ is the total hydrophobic SASA of residues in the nascent protein at time $t$ and nascent chain length $l$ exposed outside of the ribosome exit tunnel (defined as having an x-coordinate ≥100 Å in the internal CHARMM coordinate system; see Ref. 36 for details). The term $\langle A_{\text{hydrophobic}}(t,l)\rangle_{\text{F}}$ is the mean total hydrophobic SASA of residues at time $t$ and length $l$ outside of the exit tunnel calculated over all frames of synthesis trajectories identified to be folded by $Q_{\text{mode}}$ and $G$ analysis. Note well, the reference states for Equations 10-12 are the native-state reference simulations initiated in bulk solution, but due to the co-translational nature of TF interactions we use the folded subpopulation of synthesis trajectories as the reference state in Eq. 9. As TF is thought to only interact with nascent proteins of 100 residues or longer[62], we compute Eq. 9 for $l = \{100, 101, \ldots, N\}$ for each protein and trajectory, where $N$ is the total number of residues in the full-length protein. Proteins shorter than 100 amino acids are considered not to interact with TF. To quantify the overall propensity of a misfolded trajectory of a given protein to interact with TF we compute $\langle \zeta_{\text{hydrophobic}}^{\text{co-t}}\rangle$ as the mean of $\zeta_{\text{hydrophobic}}^{\text{co-t}}(t,l)$ for all $t$ and allowed values of $l$. Based on examination of $\zeta_{\text{hydrophobic}}^{\text{co-t}}(t,l)$ time series, we determined that a value of $\langle \zeta_{\text{hydrophobic}}^{\text{co-t}}\rangle \leq 10\%$ corresponds to proteins that are unlikely to engage TF significantly more than folded conformations on the ribosome. For the purposes of this calculation and all others in this work that consider sets of hydrophobic residues, coarse-grain interactions sites representing {Ile, Val, Leu, Phe, Cys, Met, Ala, Gly, Trp} are considered to be hydrophobic.

**Identifying misfolded proteins unlikely to interact with GroEL/GroES.** Our data set of 122 *E. coli* proteins was first cross-referenced with the list of 276 confirmed GroEL/GroES substrates[40–42]. The 58 proteins that do not appear in this list of confirmed clients are considered to not be GroEL/GroES client proteins. GroEL/GroES is thought to identify and bind regions of exposed hydrophobic surface area on nascent proteins. To determine whether the misfolded conformations of proteins that can interact with GroEL/GroES are likely to do so, we compared the total hydrophobic SASA of misfolded conformations with the native-state ensemble using the equation

$$\zeta_{\text{hydrophobic}}(t) = \left(\frac{A_{\text{hydrophobic}}(t)}{\langle A_{\text{hydrophobic}}(t)\rangle_{\text{NS}}} - 1\right) * 100\%. \quad [10]$$

This equation provides the quantity $\zeta_{\text{hydrophobic}}(t)$, which measures the relative difference between the SASA of hydrophobic residues within a misfolded conformation at time $t$ ($A_{\text{hydrophobic}}(t)$) in comparison to to the mean hydrophobic SASA calculated over all frames of the native-state reference simulations ($\langle A_{\text{hydrophobic}}(t)\rangle_{\text{NS}}$). As GroEL/GroES interacts with proteins post-translationally, Eq. 10 was applied to the post-translational simulation data for all misfolded conformations of a given protein and the average over the final 100 ns computed for each trajectory ($\langle \zeta_{\text{hydrophobic}}\rangle$). The full set of proteins with misfolded conformations that we predict will not interact with GroEL/GroES is taken as the union of the sets of all trajectories for proteins not in the list of experimentally confirmed clients and the list of trajectories with $\langle \zeta_{\text{hydrophobic}}\rangle \leq 10\%$.

**Identifying misfolded proteins unlikely to interact with DnaK.** We first cross-referenced our list of 122 simulated proteins with a list of DnaK client proteins. The 21 proteins that do not appear on this list are considered to not interact with DnaK. To determine whether misfolded conformations of proteins that can interact with DnaK are likely to do so, we predicted DnaK binding sites using the Limbo webserver[31] and then compared the total SASA of residues in predicted binding sites between misfolded conformations and the native state using the equation

$$\zeta_{\mathrm{DnaK}}(t) = \left(\frac{A_{\mathrm{DnaK}}(t)}{\langle A_{\mathrm{DnaK}}(t)\rangle_{\mathrm{NS}}} - 1\right) * 100\%. \quad [11]$$

In this equation $A_{\mathrm{DnaK}}(t)$ is the total SASA of residues in DnaK binding sites as a function of time $t$ and $\langle A_{\mathrm{DnaK}}(t)\rangle_{\mathrm{NS}}$ is the mean total SASA of residues in DnaK binding sites averaged over all frames of the native-state reference simulations. Equation 11 was applied to the post-translational simulation data of all misfolded conformations for each misfolded trajectory and the average over the final 100 ns of each trajectory computed ($\langle\zeta_{\mathrm{DnaK}}\rangle$). The full set of trajectories with misfolded conformations that we predict will not interact with DnaK is taken as the union of the sets of trajectories for proteins not in the list of experimentally confirmed clients and the list of trajectories with $\langle\zeta_{\mathrm{DnaK}}\rangle \leq 10\%$. We note that three proteins, PDB IDs 2JRX, 2V81, and 2KFW, are predicted by Limbo to have no DnaK binding sites. 2JRX and 2KFW appear on the list of confirmed DnaK clients, and are therefore considered to be DnaK binders. 2V81, however, is not a confirmed DnaK client and is thus counted as not likely to interact with DnaK.

**Identifying misfolded proteins unlikely to aggregate.** We used the AMYLPRED2 webserver[33] to predict the sets of aggregation-prone residues within the primary sequences of our 122 proteins. Whether or not a given trajectory for a protein is likely to aggregate was determined by comparing the SASA of aggregation-prone regions within the misfolded trajectory to the mean SASA of aggregation-protein regions in the native-state reference ensemble using the equation

$$\zeta_{\mathrm{agg}}(t) = \left(\frac{A_{\mathrm{agg}}(t)}{\langle A_{\mathrm{agg}}(t)\rangle_{\mathrm{NS}}} - 1\right) * 100\%. \quad [12]$$

In this equation, $A_{\mathrm{agg}}(t)$ is the total SASA of residues in aggregation-prone regions as a function of time, $t$, and $\langle A_{\mathrm{agg}}(t)\rangle_{\mathrm{NS}}$ is the mean total SASA of residues in aggregation-prone regions averaged over all frames of the native-state reference simulations. Equation 12 was applied to the post-translational simulation data for all misfolded trajectories of a given protein and the average over the final 100 ns for each trajectory ($\langle\zeta_{\mathrm{agg}}\rangle$). Trajectories with $\langle\zeta_{\mathrm{agg}}\rangle \leq 10\%$ are considered to be unlikely to aggregate.

**Identifying misfolded proteins unlikely to be degraded.** Whether or not a protein's misfolded conformations are likely to be targeted for degradation was determined on the basis of $\zeta_{\mathrm{hydrophobic}}$ (Eq. 10). Proteins with $\langle\zeta_{\mathrm{hydrophobic}}\rangle \leq 10\%$ are considered to be unlikely to be degraded.

**Creating a database of functional residues for *E. coli* proteins.** Information from UniProt and RCSB was unified and parsed to create a database of residues implicated in function for each of our 122 proteins. Residues involved in interactions with small molecules were identified as those residues with heavy atoms within 4.5 Å of any heteroatom (identified by the HETATM keyword in PDB records) other than water and non-native amino acids such as selenidomethionine (*i.e.*, MSE residues). Many proteins must form multimeric complexes in order to exercise their function. To consider these interactions, we also identified residues with heavy atoms within 4.5 Å of heavy atoms in a different chain ID within the same PDB structure.

Our 122 coarse-grain models were built from single PDB structures or, in the case of some multi-domain protein models, the merging of multiple structures. These structures used for model building often lack ligands or protein binding partners that may be required for function due to differences in crystallographic conditions and/or the intention of the original crystallographers. To provide a broader view of functional residues we therefore also considered all PDB structures identified by UniProt to represent the same gene product. Functional residues were identified in these alternative structures as described above for the initial structures. PDB entries representing the same protein often have different residue numbering schemes and a small number of mutations. We therefore used amino acid alignments in BLAST to determine the mapping from alternative numbering schemes to the numbering scheme within the structure used for model building. Only those domains with at least 97% sequence identity were considered for this analysis to allow for small mutational changes while excluding significantly different proteins. A summary of each of the terms in the database and their meanings is provided in Table S9.

**Determining which misfolded conformations are likely non-functional.** The relative difference in function between misfolded trajectories and native-state reference trajectories was determined by calculating the relative difference of the structural overlap of residues identified to be involved in protein function using the equation

$$\chi_{\mathrm{func}} = \left(1 - \frac{\chi(t)}{\langle \chi(t) \rangle_{\mathrm{NS}}}\right) * 100\%. \qquad [13]$$

In this equation, $\chi(t)$ is the structural overlap between residues implicated in function at time $t$ in a misfolded trajectory with the native-state reference structure and $\langle \chi(t) \rangle_{\mathrm{NS}}$ is the mean of this same value computed over all simulation frames of the native-state reference simulations. The value of $\chi(t)$ is calculated as

$$\chi(t) = \frac{1}{N} \sum_{i \in \{\mathrm{Func}\}} \sum_{j \geq i+2;\, j \in \{\mathrm{Func}\}} \Theta\left[\epsilon - \left|r_{ij}(t) - r_{ij}^{0}\right|\right] \qquad [14]$$

and gives the fraction of pairwise distances that are at native-like values at time $t$. The set $\{\mathrm{Func}\}$ contains all residues implicated in protein function. The indices $i$ and $j$ correspond to residues in $\{\mathrm{Func}\}$ for the protein being analyzed. $N$ is the total number of pairwise contacts between residues $i$ and $j$ both in $\{\mathrm{Func}\}$ that also satisfy the condition $j \geq i + 2$. The parameters $r_{ij}(t)$ and $r_{ij}^{0}$ are the distances between residues $i$ and $j$ at time $t$ and between $i$ and $j$ in the native state reference structure, respectively. $\Theta(x)$ is the step function given by

$$\Theta(x) = \begin{cases} 1, & x \geq 0 \\ 0, & x < 0 \end{cases}. \qquad [15]$$

The value of $\epsilon$ is taken as $0.2 \cdot \mathrm{r}_{\mathrm{c}_\alpha}$ where $\mathrm{r}_{\mathrm{c}_\alpha} = 3.81$ Å is the virtual bond length between coarse-grain interaction sites in the coarse-grain simulation model. A particular pair of residues $i$ and $j$ contribute 1 to $\chi(t)$ if $\left|r_{ij}(t) - r_{ij}^{0}\right|$ is less than $\epsilon$, such that $\epsilon - \left|r_{ij}(t) - r_{ij}^{0}\right| \geq 0$, and in all other situations contribute 0 to $\chi(t)$. Similar forms of this equation have been used previously to observe structural transitions in simulations of proteins[63]. The value of $\chi_{\mathrm{func}}$ was averaged over the final 100 ns for given protein trajectory to give $\langle \chi_{\mathrm{func}} \rangle$, the mean relative difference in structure of functional residues for the misfolded conformation in comparison to the native state. Trajectories with $\langle \chi_{\mathrm{func}} \rangle \geq 10\%$ are considered to be less functional than the native state.

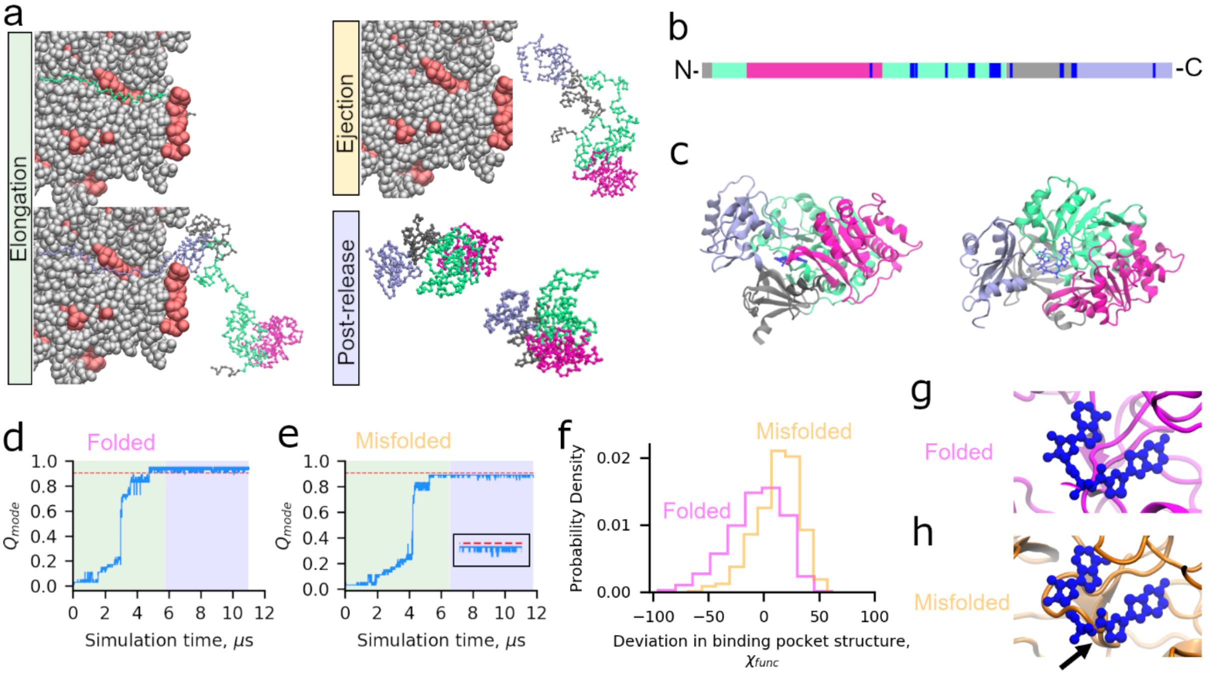


**Figure 1**. **Luciferase exhibits subpopulations that misfold into soluble but less-functional conformations**. (a) Simulations of translation elongation and ejection of nascent Luciferase were performed with a coarse-grain ribosome representation (ribosomal proteins and RNA are displayed in red and grey respectively). Domains 1, 2, 3, and 4 of Luciferase are displayed in silver, light green, magenta, and light purple, respectively. After ejection, the ribosome is removed and post-release dynamics is simulated for 30 CPU days per trajectory. (b) Primary structure diagram of Luciferase colored as described in (a); positions involved in the catalytic function of Luciferase as described in Methods are colored blue. (c) Cartoon diagram of Luciferase native state colored as described in (a) with the 5′-O-[N-dehydroluciferyl)-sulfamoyl]-adenosine ligand colored dark blue. (d) $Q_{\mathrm{mode}}$ (see Methods) versus time for Domain 2 of a trajectory of Luciferase that folded correctly. Portions of the plot colored green, yellow, and blue correspond to the synthesis, ejection, and post-translation phases of the simulation. Note that the relatively short duration of ejection for this protein renders that section of the plot invisible at this resolution. The red line corresponds to $\langle Q_{\mathrm{mode}}^{\mathrm{NS}} \rangle$ minus three standard deviations and represents the threshold for defining this domain as folded (see Methods). (e) Same as (d) but for a trajectory that misfolds. Inset shows the final microsecond of the $Q_{\mathrm{mode}}$ time series. (f) Distributions of $\chi_{\mathrm{func}}$ (Eq. 13) over the final 100 ns of the folded (magenta) and misfolded (orange) trajectories displayed in panels (d) and (e). The misfolded distribution shows greater structural distortion (i.e., values of $\chi_{\mathrm{func}} > 0$) of the binding pocket. (g) Backmapped all-atom structure from the final frame of the folded simulation shown in (d) aligned based on the residues implicated in function to the native state. (h) Same as (g) except for the final structure from the misfolded trajectory in (e), showing a strand misfolding in the ligand binding pocket (indicated by black arrow). Steric conflict between where the substrate binds and the surrounding binding pocket of the misfolded structures indicates this misfolded state will have reduced function.

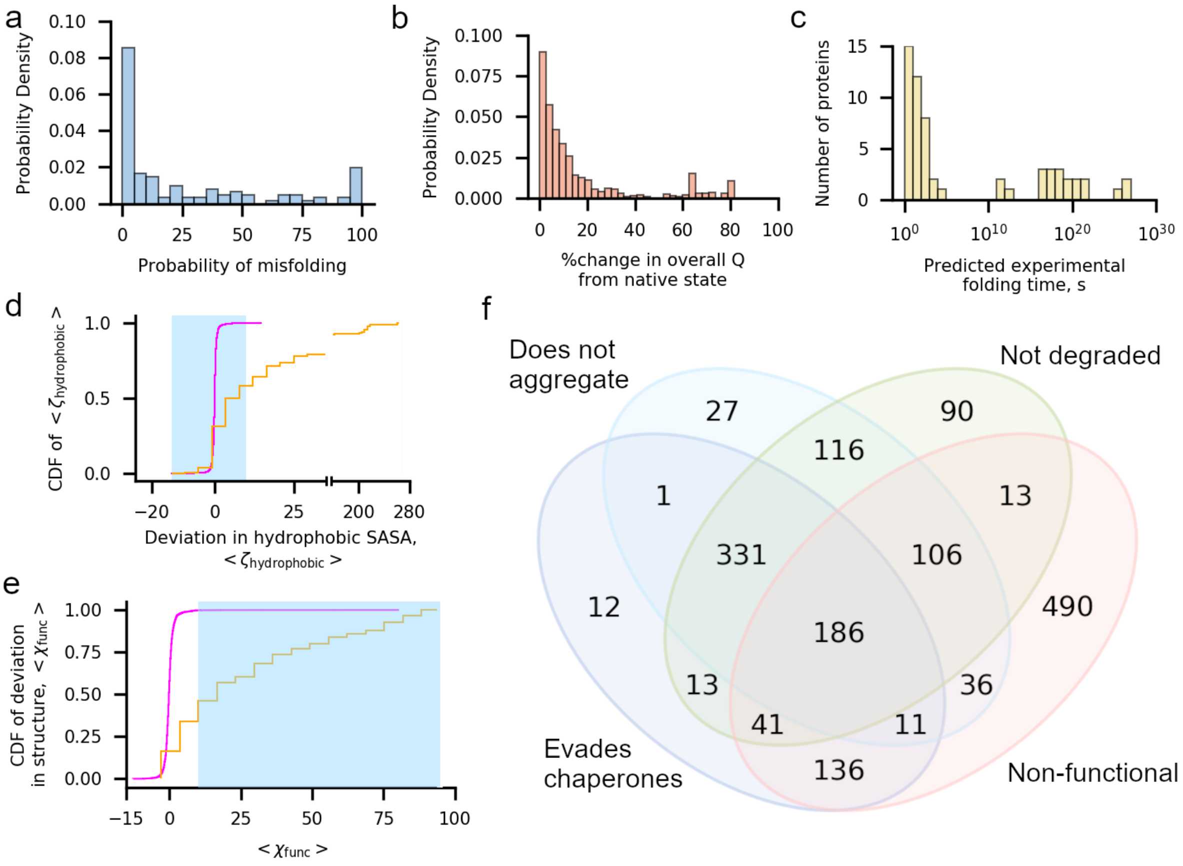


**Figure 2. One in three proteins exhibit subpopulations that misfold into soluble but less-functional conformations that evade proteostasis machinery.** (a) Histogram of the probability of misfolding, computed as the number of misfolded trajectories divided by 50, for each of the 122 proteins in the cytosolic *E. coli* proteome set. (b) Histogram of the percent change, computed as $\frac{|\langle Q_{\text{overall}}\rangle - \langle Q_{\text{overall}}^{\text{NS}}\rangle|}{\langle Q_{\text{overall}}^{\text{NS}}\rangle} * 100\%$, in fraction of native contacts within the final 100 ns of each of the 1,631 misfolded trajectories ($\langle Q_{\text{overall}}\rangle$) in the *E. coli* proteome data set relative to the average value from each protein's native state simulations ($\langle Q_{\text{overall}}^{\text{NS}}\rangle$). The majority of misfolded proteins are within 10% of the native value. (c) Histogram of extrapolated folding times for the slow-folding kinetic phase from survival probability curves for the 73 proteins in the cytosolic *E. coli* dataset with a reliable estimate (see Methods). (d) Cumulative distribution function (CDF) of $\langle \zeta_{\text{hydrophobic}} \rangle$ computed over the values of $\langle \zeta_{\text{hydrophobic}} \rangle$ (Eq. 10) for 1,631 misfolded (orange) and 4,469 folded (magenta) trajectories. The blue shaded region indicates the set of $\langle \zeta_{\text{hydrophobic}} \rangle$ values considered to have no significant increase in hydrophobic solvent-accessible surface area relative to the native-state ensemble. (e) Same as (d) but CDFs are computed over values of $\langle \chi_{\text{func}} \rangle$ for trajectories in the misfolded and folded populations. (f) Venn diagram indicating the number of the 1,631 misfolded trajectories that evade chaperones (TF, DnaK, and GroEL/GroES), do not aggregate, are not degraded, and are non-functional. The 186 trajectories at the center of this diagram are misfolded states that are expected to evade the proteostasis machinery, remain soluble but non-functional.

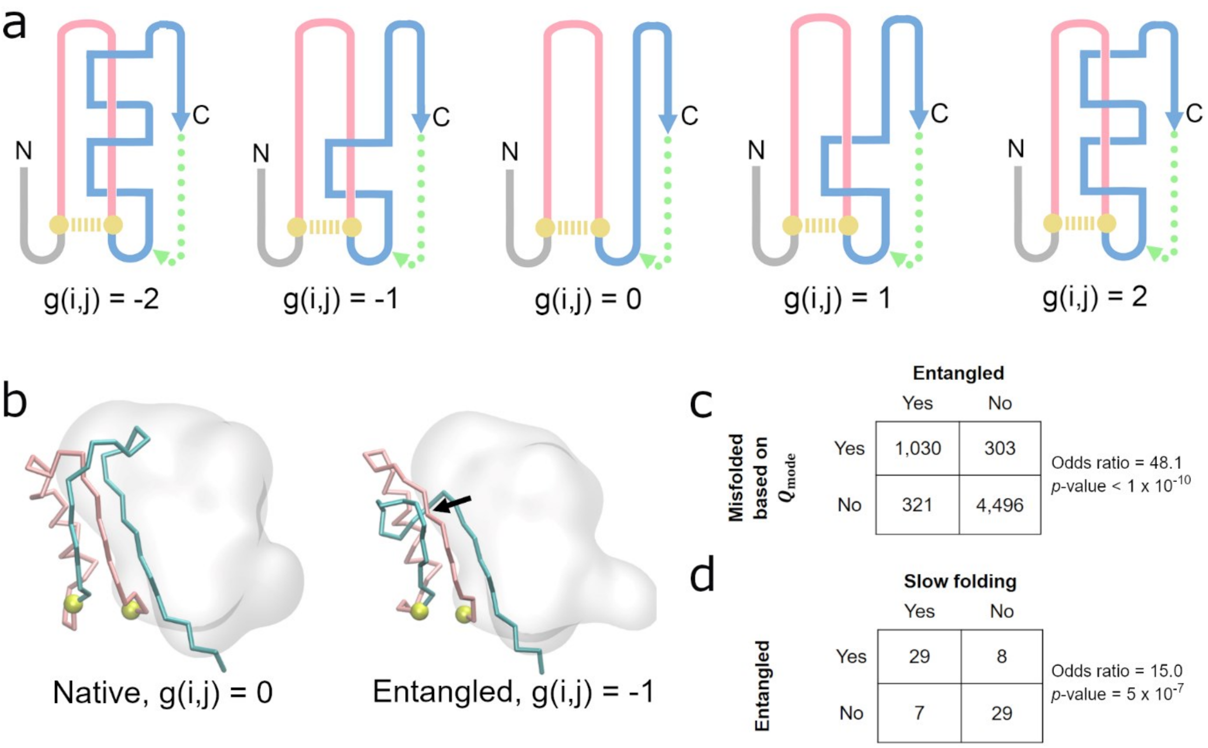


**Figure 3**. **Detecting non-native entanglements in a monomeric protein structure.** (a) Schematic of how self-entanglements can be detected by examining the change in the Gauss linking number g(*i,j*) (Eq. 5) between a closed loop (pink) formed by the backbone segment between residues *i* and *j* that form a native contact (gold dashed line) and another pseudo-closed loop formed by the C-terminal backbone segment (blue) and a pseudo-vector (dashed green line) connecting the C-terminal residue and the start of the C-terminal segment, which begins at residue *j*+1. Threading of the N-terminal segment (composed of residues 1 through *i*-1) is determined in a similar manner. Examples of different Gauss linking numbers and their corresponding structures are shown in this hypothetical illustration. The magnitude of g(*i,j*) is proportional to the number of threading events of the blue segment through the pink loop, while its sign is a function of the relative positioning of primary structure vectors at crossing points between the pink and blue segments. The structure with g(*i,j*) =0 exhibits no entanglement. (b) An example of a gain in entanglement of the protein YJGH (PDB: 1PF5), where the C-termini (cyan) threads a loop (pink) formed by the native contact between residues D72 & Y104 (gold). Black arrow indicates the location of the crossing point of the two entangled loops. (c) Contingency table indicating the number of trajectories that are misfolded/folded across our 122 proteins based on $Q_{\mathrm{mode}}$ analysis and entangled/not entangled. Indicated $p$-values and odds ratios were computed in SciPy using the Fisher Exact Test. (d) Same as (c) except contingency table displays the number of trajectories that are entangled/not entangled and predicted to be slow folding/fast folding. For the purposes of this analysis, a protein is considered slow- or fast-folding if its computed folding time is above or below the median folding time from the set of 73 proteins with reliable estimates, respectively. A protein is considered entangled if ≥50% of its misfolded trajectories are entangled.

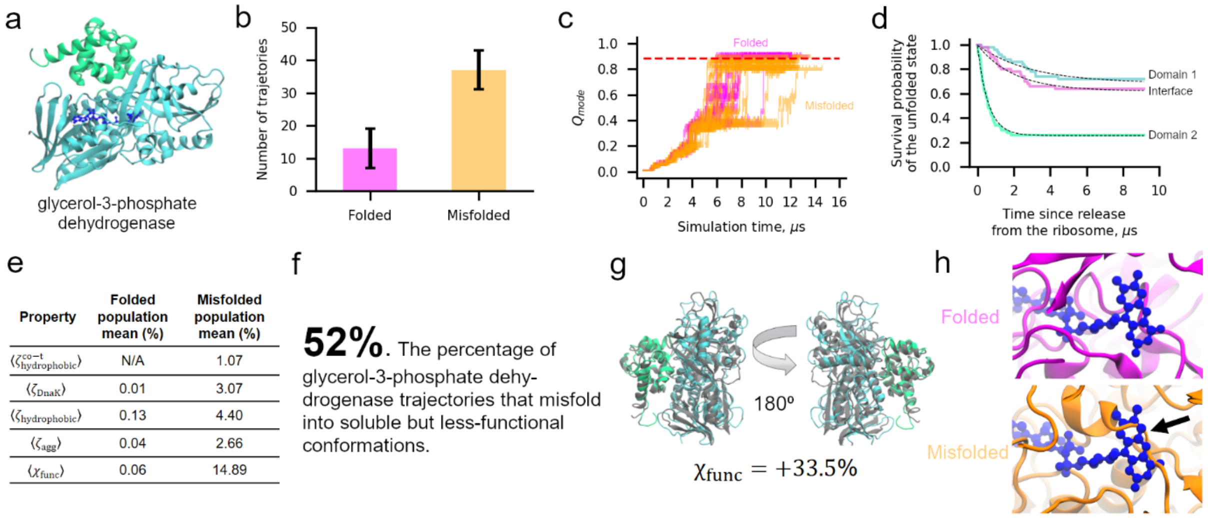


**Figure 4. The two-domain glycerol-3-phosphate dehydrogenase protein displays widespread misfolding into soluble but less-functional conformations.** (a) Ribbon structure of glycerol-3-phosphate dehydrogenase (PDB ID: 2QCU). Domains 1 and 2 are composed by residues 1-387 and 388-501 and are colored cyan and green, respectively. The flavin adenine dinucleotide ligand is shown in a dark blue representation. (b) Number of folded and misfolded trajectories for this protein in our simulations. Error bars are 95% confidence intervals from 1 x $10^6$ bootstrapped samples. (c) $Q_{\mathrm{mode}}$ versus time for the subpopulations of folded (magenta) and misfolded (orange) trajectories. Each line represents one independent trajectory. (d) Survival probability of the unfolded state versus time computed for Domain 1 (cyan), Domain 2 (green), and the Domain 1|2 interface as described in Methods. Dotted black lines are double-exponential fits used to extract rate constants. (e) Summary of key parameters for the folded and misfolded populations. (f) Of the 37/50 trajectories that are identified to be misfolded, 26 are predicted to remain misfolded but soluble in less-functional conformations. (g) Representative misfolded structure colored as in (a) aligned to the native-state reference structure. Despite a high $\chi_{\mathrm{func}}$ values indicative of a less-functional conformation, the protein is largely native. (h) Representative folded and misfolded structures back-mapped to all-atom resolution and then aligned to the native state based on the flavin adenine dinucleotide binding pocket residues. Steric conflict (indicated by black arrow) can be seen between the substrate binding location and the misfolded binding pocket, indicating reduced function of this conformation.

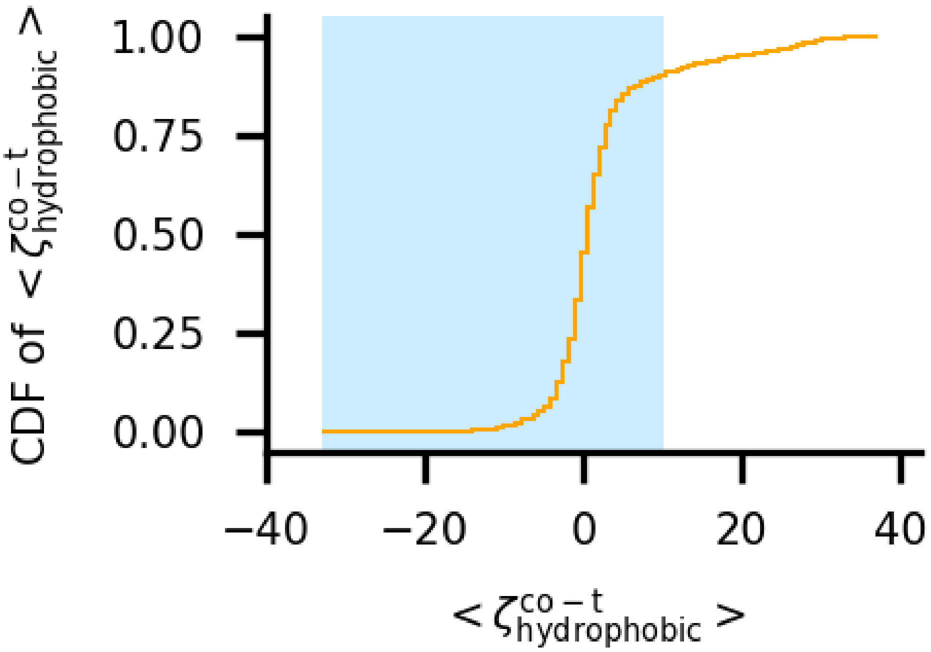


**Figure S1**. Cumulative distribution function of $\langle\zeta^{co-t}_{hydrophobic}\rangle$ over the subset of 1,051 misfolded trajectories for which it was computed (see Methods). The blue shaded region indicates the subset of values taken to indicate no significant increase in trigger factor interactions relative to the folded population.

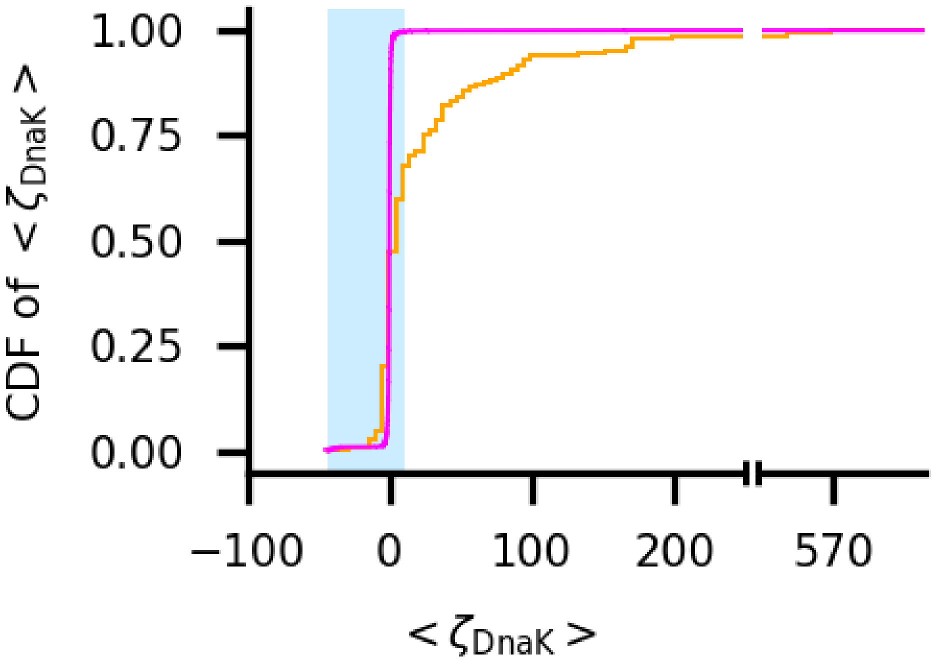


**Figure S2**. Cumulative distribution function of $\langle \zeta_{\mathrm{DnaK}} \rangle$ over the subset of 1,631 misfolded (orange) and 4,469 folded (magenta) trajectories. The blue shaded region indicates the subset of values taken to indicate no significant increase in DnaK interactions relative to the native-state population.

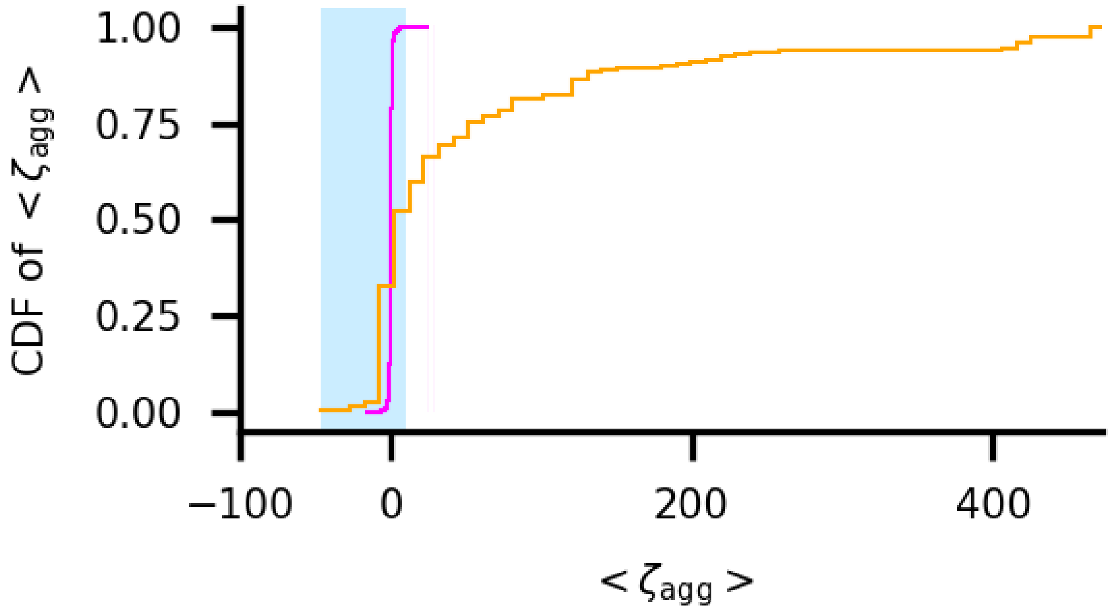


**Figure S3**. Cumulative distribution function of $\langle \zeta_{\text{Agg}} \rangle$ over the subset of 1,631 misfolded (orange) and 4,469 folded (magenta) trajectories. The blue shaded region indicates the subset of values taken to indicate no significant increase in aggregation propensity relative to the native state simulations.

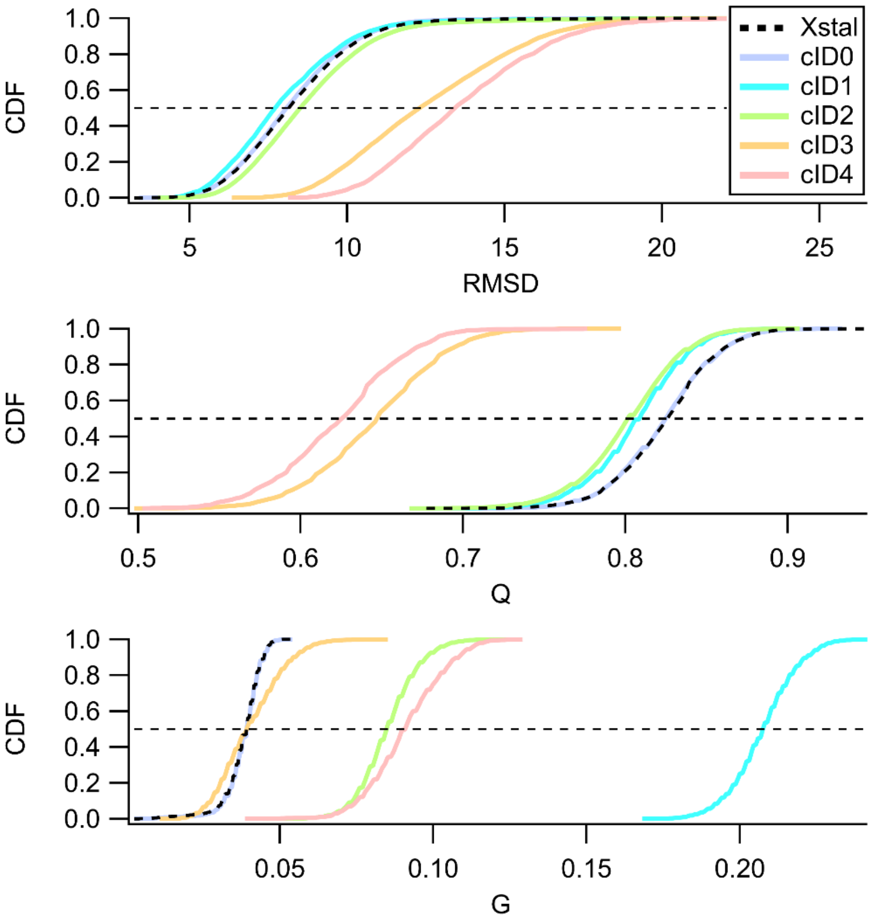


**Figure S4**. The utility of the change in entanglement metric (G) for detecting structural perturbations not apparent by fraction of native contacts (Q) or root mean square deviation (RMSD) is exemplified by examining clusters of the trajectories for aminoimidazole ribonucleotide synthetase (PDB: 1CLI) relative to the cluster of trajectories started from the reference state. The 50 trajectories obtained from synthesis simulations can be clustered into 5 separate clusters characterized by the $\langle Q_{\text{mode}} \rangle$ and the corrected discrete distribution of a give type of change in entanglement (Eq. 7). cID0 contains 8/50 trajectories all of which terminate to the native state with no changes in entanglement. cID1 contains 3/50 trajectories that are near native like (i.e. $\langle Q_{\text{mode}} \rangle \approx \langle Q_{\text{mode}}^{NS} \rangle$) that show appreciable gain and loss of entanglements as well as pure switch in chirality. cID2 contains 32/50 trajectories that show appreciable gain and loss of entanglements with less near native like conformations (i.e. $\langle Q_{\text{mode}} \rangle \geq \langle Q_{\text{mode}}^{NS} \rangle - \sigma \langle Q_{\text{mode}}^{NS} \rangle$). cID3 contains 6/50 trajectories that show non-native like conformations (i.e. $\langle Q_{\text{mode}} \rangle < \langle Q_{\text{mode}}^{NS} \rangle - \sigma \langle Q_{\text{mode}}^{NS} \rangle$) with appreciable gain and loss of entanglements. cID4 contains 1/50 trajectories that show non-native like conformations with appreciable gain and loss of entanglements and pure change in chirality. For the clusters which contain near native like conformations and changes in entanglement of any kind (cID1 light blue and cID2) Q and RMSD fail to distinguish different clusters while G does not. For those clusters that contain non-native like conformations and changes in entanglement of any kind (cID3 & cID4) Q and RMSD are much more reliable.

**Table S1.** Coarse-grain model parameters for PDB ID 4G36

| PDB ID | Protein name | Domain or interface | Structural class | $\eta$ used in model building |
|---|---|---|---|---|
| 4G36 | Firefly Luciferase | Domain 1:1-12; 356-437 | β | 1.442 |
| | | Domain 2:13-52; 212-355 | α/β | 1.114 |
| | | Domain 3: 53-211 | α/β | 1.114 |
| | | Domain 4: 438-550 | α/β | 1.916 |
| | | Interface 1\|2 | - | 1.235 |
| | | Interface 1\|3 | - | 1.235 |
| | | Interface 1\|4 | - | 1.507* |
| | | Interface 2\|3 | - | 1.235 |
| | | Interface 2\|4 | - | 1.235 |
| | | Interface 3\|4 | - | 1.235 |

*indicates interface is unstable at all tested values of $\eta$ and so median interface value from set of stable interfaces is used for all simulations

**Table S2** *E. coli* **proteome dataset information**

| Index | PDB ID | Chain ID | UniProt Entry name | Protein name | Gene names |
|---|---|---|---|---|---|
| 1 | 1A69 | A | DEOD_ECOLI | Purine nucleoside phosphorylase DeoD-type | deoD, pup, b4384, JW4347 |
| 2 | 1A6J | B | PTSN_ECOLI | Nitrogen regulatory protein | ptsN, rpoP, yhbI, b3204, JW3171 |
| 3 | 1A82 | A | BIOD1_ECOLI | ATP-dependent dethiobiotin synthetase BioD 1 | bioD1, b0778, JW0761 |
| 4 | 1AG9 | A | FLAV_ECOLI | Flavodoxin 1 | fldA, b0684, JW0671 |
| 5 | 1AH9 | Model 1 | IF1_ECOLI | Translation initiation factor IF-1 | infA, b0884, JW0867 |
| 6 | 1AKE | A | KAD_ECOLI | Adenylate kinase | Adk, dnaW, plsA, b0474, JW0463 |
| 7 | 1B9L | A | FOLX_ECOLI | Dihydroneopterin triphosphate 2'-epimerase | folX, b2303, JW2300 |
| 8 | 1CLI | A | PUR5_ECOLI | Phosphoribosylformylglycinamidine cyclo-ligase | purM, purG, b2499, JW2484 |
| 9 | 1D2F | A | MALY_ECOLI | Protein MalY | malY, b1622, JW1614 |
| 10 | 1DCJ | Model 1 | TUSA_ECOLI | Sulfur carrier protein TusA | tusA, sirA, yhhP, b3470, JW3435 |
| 11 | 1DFU | P | RL25_ECOLI | 50S ribosomal protein L25 | rplY, b2185, JW2173 |
| 12 | 1DUV | G | OTC1_ECOLI | Ornithine carbamoyltransferase subunit I | argI, b4254, JW4211 |
| 13 | 1DXE | A | GARL_ECOLI | 5-keto-4-deoxy-D-glucarate aldolase | garL, yhaF, b3126, JW3095 |
| 14 | 1EF9 | A | SCPB_ECOLI | Methylmalonyl-CoA decarboxylase | scpB, mmcD, ygfG, b2919, JW2886 |
| 15 | 1EIX | C | PYRF_ECOLI | Orotidine 5'-phosphate decarboxylase | pyrF, b1281, JW1273 |
| 16 | 1EM8 | A | HOLC_ECOLI | DNA polymerase III subunit chi | holC, b4259, JW4216 |
| 17 | 1EUM | A | FTNA_ECOLI | Bacterial non-heme ferritin | ftnA, ftn, gen-165, rsgA, b1905, JW1893 |
| 18 | 1FJJ | A | YBHB_ECOLI | UPF0098 protein YbhB | ybhB, b0773, JW0756 |
| 19 | 1FM0 | D | MOAD_ECOLI | Molybdopterin synthase sulfur carrier subunit | moaD, chlA4, chlM, b0784, JW0767 |
| 20 | 1FTS | A | FTSY_ECOLI | Signal recognition particle receptor FtsY | ftsY, b3464, JW3429 |
| 21 | 1FUI | A | FUCI_ECOLI | L-fucose isomerase, Fuclase | fucI, b2802, JW2773 |
| 22 | 1GER | B | GSHR_ECOLI | Glutathione reductase | gor, b3500, JW3467 |
| 23 | 1GLF | O | GLPK_ECOLI | Glycerol kinase | glpK, b3926, JW3897 |
| 24 | 1GQE | A | RF2_ECOLI | Peptide chain release factor RF2 | prfB, supK, b2891, JW5847 |
| 25 | 1GQT | B | RBSK_ECOLI | Ribokinase | rbsK, b3752, JW3731 |
| 26 | 1GT7 | A | RHAD_ECOLI | Rhamnulose-1-phosphate aldolase | rhaD, rhuA, b3902, JW3873 |
| 27 | 1GYT | L | AMPA_ECOLI | Cytosol aminopeptidase | pepA, carP, xerB, b4260, JW4217 |
| 28 | 1GZ0 | C | RLMB_ECOLI | 23S rRNA (guanosine-2'-O-)-methyltransferase RlmB | rlmB, yjfH, b4180, JW4138 |
| 29 | 1H16 | A | PFLB_ECOLI | Formate acetyltransferase 1 | pflB, pfl, b0903, JW0886 |
| 30 | 1H75 | A | NRDH_ECOLI | Glutaredoxin-like protein NrdH | nrdH, ygaN, b2673, JW2648 |

**Table S2** cont.

| Index | PDB ID | Chain ID | UniProt Entry name | Protein name | Gene names |
|---|---|---|---|---|---|
| 31 | 1I6O | B | CAN_ECOLI | Carbonic anhydrase 2 | can, cynT2, yadF, b0126, JW0122 |
| 32 | 1JNS | Model 1 | PPIC_ECOLI | Peptidyl-prolyl cis-trans isomerase C | ppiC, parVA, b3775, JW3748 |
| 33 | 1JW2 | A | HHA_ECOLI | Hemolysin expression-modulating protein Hha | hha, b0460, JW0449 |
| 34 | 1JX7 | A | YCHN_ECOLI | Protein YchN | ychN, b1219, JW1210 |
| 35 | 1K7J | A | YCIO_ECOLI | Uncharacterized protein YciO | yciO, b1267, JW5196 |
| 36 | 1KO5 | A | GNTK_ECOLI | Thermoresistant gluconokinase | gntK, b3437, JW3400 |
| 37 | 1KSF | X | CLPA_ECOLI | ATP-dependent Clp protease ATP-binding subunit ClpA | clpA, lopD, b0882, JW0866 |
| 38 | 1L6W | A | FSAA_ECOLI | Fructose-6-phosphate aldolase 1 | fsaA, fsa, mipB, ybiZ, b0825, JW5109 |
| 39 | 1M3U | A | PANB_ECOLI | 3-methyl-2-oxobutanoate hydroxymethyltransferase | panB, b0134, JW0130 |
| 40 | 1MZG | B | SUFE_ECOLI | Cysteine desulfuration protein SufE | sufE, ynhA, b1679, JW1669 |
| 41 | 1NAQ | A | CUTA_ECOLI | Divalent-cation tolerance protein CutA | cutA, cutA1, cycY, b4137, JW4097 |
| 42 | 1NG9 | A | MUTS_ECOLI | DNA mismatch repair protein MutS | mutS, fdv, b2733, JW2703 |
| 43 | 1ORO | A | PYRE_ECOLI | Orotate phosphoribosyltransferase | pyrE, b3642, JW3617 |
| 44 | 1P7L | A | METK_ECOLI | S-adenosylmethionine synthase | metK, metX, b2942, JW2909 |
| 45 | 1P91 | A | RLMA_ECOLI | 23S rRNA (guanine(745)-N(1))-methyltransferase | rlmA, rrmA, yebH, b1822, JW1811 |
| 46 | 1PF5 | A | YJGH_ECOLI | RutC family protein YjgH | yjgH, b4248, JW4206 |
| 47 | 1PMO | B | DCEB_ECOLI | Glutamate decarboxylase beta | gadB, b1493, JW1488 |
| 48 | 1PSU | B | PAAI_ECOLI | Acyl-coenzyme A thioesterase PaaI | paaI, ydbV, b1396, JW1391 |
| 49 | 1Q5X | A | RRAA_ECOLI | Regulator of ribonuclease activity A | rraA, menG, yiiV, b3929, JW3900 |
| 50 | 1QF6 | A | SYT_ECOLI | Threonine-tRNA ligase | thrS, b1719, JW1709 |
| 51 | 1QTW | A | END4_ECOLI | Endonuclease 4 | Nfo, b2159, JW2146 |
| 52 | 1RQJ | A | ISPA_ECOLI | Farnesyl diphosphate synthase | ispA, b0421, JW0411 |
| 53 | 1SG5 | Model 1 | ROF_ECOLI | Protein rof | Rof, yaeO, b0189, JW0184 |
| 54 | 1SV6 | A | MHPD_ECOLI | 2-keto-4-pentenoate hydratase | mhpD, b0350, JW0341 |
| 55 | 1SVT | J | CH60_ECOLI | 60 kDa chaperonin | groL, groEL, mopA, b4143, JW4103 |
| 56 | 1T4B | A | DHAS_ECOLI | Aspartate-semialdehyde dehydrogenase | asd, hom, b3433, JW3396 |
| 57 | 1T8K | A | ACP_ECOLI | Acyl carrier protein | acpP, b1094, JW1080 |
| 58 | 1U0B | B | SYC_ECOLI | Cysteine--tRNA ligase | cysS, b0526, JW0515 |
| 59 | 1U60 | A | GLSA1_ECOLI | Glutaminase 1 | glsA1, ybaS, b0485, JW0474 |
| 60 | 1UUF | A | YAHK_ECOLI | Aldehyde reductase YahK | yahK, b0325, JW0317 |
| 61 | 1W78 | A | FOLC_ECOLI | Dihydrofolate synthase/folylpolyglutamate synthase | folC, dedC, b2315, JW2312 |

**Table S2** cont.

| Index | PDB ID | Chain ID | UniProt Entry name | Protein name | Gene names |
|---|---|---|---|---|---|
| 62 | 1W8G | A | PLPHP_ECOLI | Pyridoxal phosphate homeostasis protein | yggS, b2951, JW2918 |
| 63 | 1WOC | C | PRIB_ECOLI | Primosomal replication protein N | priB, b4201, JW4159 |
| 64 | 1XN7 | Model 1 | FEOC_ECOLI | Probable [Fe-S]-dependent transcriptional repressor FeoC | feoC, yhgG, b3410, JW3373 |
| 65 | 1XRU | A | KDUI_ECOLI | 4-deoxy-L-threo-5-hexosulose-uronate ketol-isomerase | kduI, yqeE, b2843, JW2811 |
| 66 | 1XVI | A | MPGP_ECOLI | Mannosyl-3-phosphoglycerate phosphatase | yedP, b1955, JW1938 |
| 67 | 1YQQ | A | XAPA_ECOLI | Purine nucleoside phosphorylase 2 | xapA, pndA, b2407, JW2398 |
| 68 | 1ZYL | A | SRKA_ECOLI | Stress response kinase A | srkA, rdoA, yihE, b3859, JW3831 |
| 69 | 1ZZM | A | YJJV_ECOLI | Uncharacterized metal-dependent hydrolase YjjV | yjjV, b4378, JW4341 |
| 70 | 2A6Q | E | YOEB_ECOLI | Toxin YoeB | yoeB, b4539, JW5331 |
| 71 | 2AXD | Model 1 | HOLE_ECOLI | DNA polymerase III subunit theta | holE, b1842, JW1831 |
| 72 | 2D1P | B | TUSC_ECOLI | Protein TusC | tusC, yheM, b3344, JW3306 |
| 73 | 2FEK | Model 1 | WZB_ECOLI | Low molecular weight protein-tyrosine-phosphatase Wzb | wzb, b2061, JW2046 |
| 74 | 2FYM | A | ENO_ECOLI | Enolase | eno, b2779, JW2750 |
| 75 | 2GQR | A | PUR7_ECOLI | Phosphoribosylaminoimidazole-succinocarboxamide synthase | purC, b2476, JW2461 |
| 76 | 2H1F | A | A0A0H2VC26_ECOL6 | Lipopolysaccharide heptosyltransferase-1 | rfaC, c4447 |
| 77 | 2HD3 | K | EUTN_ECOLI | Ethanolamine catabolic microcompartment shell protein EutN | eutN, cchB, yffY, b2456, JW2440 |
| 78 | 2HG2 | A | ALDA_ECOLI | Lactaldehyde dehydrogenase | aldA, ald, b1415, JW1412 |
| 79 | 2HGK | Model 1 | YQCC_ECOLI | Uncharacterized protein YqcC | yqcC, b2792, JW2763 |
| 80 | 2HNA | A | MIOC_ECOLI | Protein MioC | mioC, yieB, b3742, JW3720 |
| 81 | 2HNH | A | DPO3A_ECOLI | DNA polymerase III subunit alpha | dnaE, polC, b0184, JW0179 |
| 82 | 2HO9 | Model 1 | CHEW_ECOLI | Chemotaxis protein CheW | cheW, b1887, JW1876 |
| 83 | 2ID0 | A | RNB_ECOLI | Exoribonuclease 2 | rnb, b1286, JW1279 |
| 84 | 2JEE | C | ZAPB_ECOLI | Cell division protein ZapB | zapB, yiiU, b3928, JW3899 |
| 85 | 2JO6 | Model 1 | NIRD_ECOLI | Nitrite reductase (NADH) small subunit | nirD, b3366, JW3329 |
| 86 | 2JRX | Model 1 | YEJL_ECOLI | UPF0352 protein YejL | yejL, b2187, JW2175 |
| 87 | 2KC5 | Model 1 | HYBE_ECOLI | Hydrogenase-2 operon protein HybE | hybE, b2992, JW2960 |
| 88 | 2KFW | Model 1 | SLYD_ECOLI | FKBP-type peptidyl-prolyl cis-trans isomerase SlyD, PPIase | slyD, b3349, JW3311 |
| 89 | 2KX9 | A | PT1_ECOLI | Phosphoenolpyruvate-protein phosphotransferase | ptsI, b2416, JW2409 |
| 90 | 2O1C | A | NUDB_ECOLI | Dihydroneopterin triphosphate diphosphatase | nudB, ntpA, b1865, JW1854 |
| 91 | 2OQ3 | Model 1 | PTMA_ECOLI | Mannitol-specific cryptic phosphotransferase enzyme IIA component | cmtB, b2934, JW2901 |
| 92 | 2PTH | A | PTH_ECOLI | Peptidyl-tRNA hydrolase | pth, b1204, JW1195 |
| 93 | 2PTQ | A | PUR8_ECOLI | Adenylosuccinate lyase | purB, b1131, JW1117 |
| 94 | 2QCU | B | GLPD_ECOLI | Aerobic glycerol-3-phosphate dehydrogenase | glpD, glyD, b3426, JW3389 |
| 95 | 2QVR | A | F16PA_ECOLI | Fructose-1,6-bisphosphatase class 1 | fbp, fdp, b4232, JW4191 |

**Table S2** cont.

| Index | PDB ID | Chain ID | UniProt Entry name | Protein name | Gene names |
|---|---|---|---|---|---|
| 96 | 2R5N | A | TKT1_ECOLI | Transketolase 1 | tktA, tkt, b2935, JW5478 |
| 97 | 2UYJ | A | TDCF_ECOLI | Putative reactive intermediate deaminase TdcF | tdcF, yhaR, b3113, JW5521 |
| 98 | 2V81 | A | DGOA_ECOLI | 2-dehydro-3-deoxy-6-phosphogalactonate aldolase | dgoA, yidU, b4477, JW5628 |
| 99 | 2WIU | A | HIPA_ECOLI | Serine/threonine-protein kinase toxin HipA, Ser/Thr-protein kinase HipA | hipA, b1507, JW1500 |
| 100 | 2WW4 | A | ISPE_ECOLI | 4-diphosphocytidyl-2-C-methyl-D-erythritol kinase, CMK | ispE, ipk, ychB, b1208, JW1199 |
| 101 | 2YVA | A | DIAA_ECOLI | DnaA initiator-associating protein DiaA | diaA, yraO, b3149, JW3118 |
| 102 | 3ASV | B | YDFG_ECOLI | NADP-dependent 3-hydroxy acid dehydrogenase YdfG | ydfG, b1539, JW1532 |
| 103 | 3BMB | B | RNK_ECOLI | Regulator of nucleoside diphosphate kinase | rnk, b0610, JW0602 |
| 104 | 3BRQ | B | ASCG_ECOLI | HTH-type transcriptional regulator AscG | ascG, b2714, JW5434 |
| 105 | 3GN5 | B | MQSA_ECOLI | Antitoxin MqsA | mqsA, ygiT, b3021, JW2989 |
| 106 | 3HWO | A | ENTC_ECOLI | Isochorismate synthase EntC | entC, b0593, JW0585 |
| 107 | 3IV5 | B | FIS_ECOLI | DNA-binding protein Fis | fis, b3261, JW3229 |
| 108 | 3M7M | X | HSLO_ECOLI | 33 kDa chaperonin | hslO, yrfI, b3401, JW5692 |
| 109 | 3N1S | J | HINT_ECOLI | Purine nucleoside phosphoramidase | hinT, ycfF, b1103, JW1089 |
| 110 | 3NXC | A | SLMA_ECOLI | Nucleoid occlusion factor SlmA | slmA, ttk, yicB, b3641, JW5641 |
| 111 | 3OFO | D | RS4_ECOLI | 30S ribosomal protein S4 | rpsD, ramA, b3296, JW3258 |
| 112 | 3PCO | D | SYFB_ECOLI | Phenylalanine--tRNA ligase beta subunit | pheT, b1713, JW1703 |
| 113 | 3QOU | A | CNOX_ECOLI | Chaperedoxin | cnoX, ybbN, b0492, JW5067 |
| 114 | 4A2C | A | GATD_ECOLI | Galactitol 1-phosphate 5-dehydrogenase | gatD, b2091, JW2075 |
| 115 | 4DCM | A | RLMG_ECOLI | Ribosomal RNA large subunit methyltransferase G | rlmG, ygjO, b3084, JW5513 |
| 116 | 4DZD | A | CAS6_ECOLI | CRISPR system Cascade subunit CasE | casE, cas6e, ygcH, b2756, JW2726 |
| 117 | 4E8B | A | RSME_ECOLI | Ribosomal RNA small subunit methyltransferase E | rsmE, yggJ, b2946, JW2913 |
| 118 | 4FZW | A | PAAF_ECOLI | 2,3-dehydroadipyl-CoA hydratase | paaF, ydbR, b1393, JW1388 |
| 119 | 4HR7 | A | ACCC_ECOLI | Biotin carboxylase | accC, fabG, b3256, JW3224 |
| 120 | 4IM7 | A | A0A0H2V7F2_ECOL6 | Hypothetical oxidoreductase ydfI | c1968 |
| 121 | 4IWX | A | RIMK_ECOLI | Ribosomal protein S6--L-glutamate ligase | rimK, b0852, JW0836 |
| 122 | 4KN7 | A | RPOB_ECOLI | DNA-directed RNA polymerase subunit beta, RNAP subunit beta | rpoB, groN, nitB, rif, ron, stl |

**Table S3**. Definitions of entanglement types

| Type | Change in entanglement | Change in chirality | Conditional* |
|---|---|---|---|
| G0 | Gain | No | $\lvert g^{\text{current}}(i,j)\rvert > \lvert g^{\text{native}}(i,j)\rvert \text{and } g^{\text{current}}(i,j) \times g^{\text{native}}(i,j) \geq 0$ |
| G1 | Gain | Yes | $\lvert g^{\text{current}}(i,j)\rvert > \lvert g^{\text{native}}(i,j)\rvert \text{and } g^{\text{current}}(i,j) \times g^{\text{native}}(i,j) < 0$ |
| G2 | Lose | No | $\lvert g^{\text{current}}(i,j)\rvert < \lvert g^{\text{native}}(i,j)\rvert \text{and } g^{\text{current}}(i,j) \times g^{\text{native}}(i,j) \geq 0$ |
| G3 | Lose | Yes | $\lvert g^{\text{current}}(i,j)\rvert < \lvert g^{\text{native}}(i,j)\rvert \text{and } g^{\text{current}}(i,j) \times g^{\text{native}}(i,j) < 0$ |
| G4 | None | Yes | $\lvert g^{\text{current}}(i,j)\rvert = \lvert g^{\text{native}}(i,j)\rvert \text{and } g^{\text{current}}(i,j) \times g^{\text{native}}(i,j) < 0$ |
| G5 | None | None | $\lvert g^{\text{current}}(i,j)\rvert = \lvert g^{\text{native}}(i,j)\rvert \text{and } g^{\text{current}}(i,j) \times g^{\text{native}}(i,j) \geq 0$ |

* conditional used to determine if a native contact between residues $i$ and $j$ has a change in a given type of entanglement in the current structure relative to the native state structure.

**Table S4.** Proteins categorized by percent of trajectories misfolded based

| Category | Number of proteins | PDB IDs |
|---|---|---|
| At least 2% misfolded | 80 | 1A69, 1A6J, 1A82, 1AH9, 1DFU, 1DXE, 1FJJ, 1GQT, 1GT7, 1H16, 1I6O, 1K7J, 1L6W, 1M3U, 1NAQ, 1ORO, 1P91, 1PF5, 1PMO, 1Q5X, 1SV6, 1U60, 1W8G, 1WOC, 1YQQ, 1ZYL, 2HNA, 2JO6, 2O1C, 2PTH, 2V81, 2YVA, 3BMB, 3HWO, 4A2C, 1CLI, 1D2F, 1DUV, 1EF9, 1FTS, 1FUI, 1GER, 1GLF, 1GYT, 1GZ0, 1KSF, 1NG9, 1P7L, 1QF6, 1SVT, 1T4B, 1U0B, 1UUF, 1W78, 1XRU, 1XVI, 2FYM, 2H1F, 2HG2, 2HNH, 2ID0, 2KFW, 2KX9, 2PTQ, 2QCU, 2R5N, 2WIU, 2WW4, 3BRQ, 3GN5, 3M7M, 3PCO, 4DCM, 4DZD, 4E8B, 4FZW, 4HR7, 4IM7, 4IWX, 4KN7 |
| At least 20% misfolded | 49 | 1A69, 1DFU, 1DXE, 1GQT, 1H16, 1K7J, 1P91, 1SV6, 1U60, 1YQQ, 1ZYL, 2HNA, 2JO6, 2V81, 3HWO, 4A2C, 1CLI, 1DUV, 1FTS, 1FUI, 1GLF, 1GYT, 1GZ0, 1KSF, 1NG9, 1P7L, 1QF6, 1SVT, 1T4B, 1U0B, 1W78, 1XRU, 1XVI, 2FYM, 2HG2, 2HNH, 2ID0, 2KFW, 2KX9, 2QCU, 2R5N, 2WIU, 3GN5, 3PCO, 4E8B, 4HR7, 4IM7, 4IWX, 4KN7 |
| At least 50% misfolded | 27 | 1GQT, 1H16, 1K7J, 1P91, 1U60, 1YQQ, 2JO6, 2V81, 1CLI, 1FUI, 1GYT, 1GZ0, 1KSF, 1P7L, 1T4B, 1U0B, 1XVI, 2HNH, 2ID0, 2KX9, 2QCU, 2R5N, 3PCO, 4E8B, 4HR7, 4IM7, 4KN7 |
| 100% misfolded | 9 | 1GQT, 1H16, 1FUI, 1GYT, 1GZ0, 1XVI, 4E8B, 4HR7, 4KN7 |

**Table S5**. Extrapolated folding times for the 73 proteins with a reliable estimate

| PDB ID | Simulated $\tau_F$, ns | Estimated experimental $\tau_F$, s | Estimated experimental $\tau_F$, min | Estimated experimental $\tau_F$, h | Estimated experimental $\tau_F$, d |
|---|---|---|---|---|---|
| 1SG5 | 1.08E+01 | 4.29E-02 | 7.15E-04 | 1.19E-05 | 4.97E-07 |
| 2AXD | 1.08E+01 | 4.29E-02 | 7.15E-04 | 1.19E-05 | 4.97E-07 |
| 3IV5 | 1.15E+01 | 4.55E-02 | 7.58E-04 | 1.26E-05 | 5.27E-07 |
| 2JRX | 1.57E+01 | 6.22E-02 | 1.04E-03 | 1.73E-05 | 7.20E-07 |
| 2KC5 | 1.68E+01 | 6.67E-02 | 1.11E-03 | 1.85E-05 | 7.72E-07 |
| 2HO9 | 2.10E+01 | 8.34E-02 | 1.39E-03 | 2.32E-05 | 9.65E-07 |
| 2JEE | 2.28E+01 | 9.05E-02 | 1.51E-03 | 2.52E-05 | 1.05E-06 |
| 3QOU | 3.77E+01 | 1.50E-01 | 2.49E-03 | 4.16E-05 | 1.73E-06 |
| 1H75 | 4.25E+01 | 1.69E-01 | 2.81E-03 | 4.68E-05 | 1.95E-06 |
| 1T8K | 4.30E+01 | 1.70E-01 | 2.84E-03 | 4.73E-05 | 1.97E-06 |
| 2FEK | 4.30E+01 | 1.70E-01 | 2.84E-03 | 4.74E-05 | 1.97E-06 |
| 1MZG | 5.86E+01 | 2.32E-01 | 3.87E-03 | 6.45E-05 | 2.69E-06 |
| 3NXC | 9.97E+01 | 3.96E-01 | 6.60E-03 | 1.10E-04 | 4.58E-06 |
| 1FM0 | 1.07E+02 | 4.26E-01 | 7.10E-03 | 1.18E-04 | 4.93E-06 |
| 2HGK | 2.84E+02 | 1.13E+00 | 1.88E-02 | 3.13E-04 | 1.30E-05 |
| 4DCM | 2.92E+02 | 1.16E+00 | 1.93E-02 | 3.21E-04 | 1.34E-05 |
| 1GQE | 3.42E+02 | 1.36E+00 | 2.26E-02 | 3.76E-04 | 1.57E-05 |
| 1ZZM | 3.51E+02 | 1.39E+00 | 2.32E-02 | 3.87E-04 | 1.61E-05 |
| 2KX9 | 3.55E+02 | 1.41E+00 | 2.35E-02 | 3.92E-04 | 1.63E-05 |
| 2OQ3 | 3.63E+02 | 1.44E+00 | 2.40E-02 | 4.00E-04 | 1.67E-05 |
| 1EIX | 4.34E+02 | 1.72E+00 | 2.87E-02 | 4.78E-04 | 1.99E-05 |
| 1KO5 | 4.39E+02 | 1.74E+00 | 2.90E-02 | 4.84E-04 | 2.02E-05 |
| 1GT7 | 5.36E+02 | 2.13E+00 | 3.54E-02 | 5.91E-04 | 2.46E-05 |
| 3N1S | 5.83E+02 | 2.31E+00 | 3.85E-02 | 6.42E-04 | 2.68E-05 |
| 2QVR | 6.26E+02 | 2.48E+00 | 4.14E-02 | 6.90E-04 | 2.87E-05 |
| 1AG9 | 8.60E+02 | 3.41E+00 | 5.68E-02 | 9.47E-04 | 3.95E-05 |
| 1EUM | 9.16E+02 | 3.64E+00 | 6.06E-02 | 1.01E-03 | 4.21E-05 |
| 3OFO | 9.94E+02 | 3.94E+00 | 6.57E-02 | 1.10E-03 | 4.56E-05 |
| 2KFW | 1.81E+03 | 7.20E+00 | 1.20E-01 | 2.00E-03 | 8.33E-05 |
| 3BMB | 3.36E+03 | 1.33E+01 | 2.22E-01 | 3.70E-03 | 1.54E-04 |
| 2PTH | 3.39E+03 | 1.35E+01 | 2.24E-01 | 3.74E-03 | 1.56E-04 |
| 2GQR | 3.82E+03 | 1.51E+01 | 2.52E-01 | 4.21E-03 | 1.75E-04 |
| 1QTW | 4.13E+03 | 1.64E+01 | 2.73E-01 | 4.55E-03 | 1.90E-04 |
| 1JX7 | 4.43E+03 | 1.76E+01 | 2.93E-01 | 4.88E-03 | 2.03E-04 |
| 2YVA | 5.42E+03 | 2.15E+01 | 3.59E-01 | 5.98E-03 | 2.49E-04 |
| 1PSU | 1.05E+04 | 4.16E+01 | 6.93E-01 | 1.15E-02 | 4.81E-04 |
| 1GLF | 1.28E+04 | 5.09E+01 | 8.49E-01 | 1.41E-02 | 5.89E-04 |
| 1A6J | 1.54E+04 | 6.12E+01 | 1.02E+00 | 1.70E-02 | 7.09E-04 |
| 1DXE | 1.81E+04 | 7.18E+01 | 1.20E+00 | 2.00E-02 | 8.31E-04 |
| 2HD3 | 1.93E+04 | 7.65E+01 | 1.28E+00 | 2.13E-02 | 8.86E-04 |
| 4IM7 | 2.46E+04 | 9.77E+01 | 1.63E+00 | 2.71E-02 | 1.13E-03 |
| 1XRU | 3.38E+04 | 1.34E+02 | 2.23E+00 | 3.72E-02 | 1.55E-03 |

**Table S5** cont.

| PDB ID | Simulated $\tau_\mathrm{F}$, ns | Estimated experimental $\tau_\mathrm{F}$, s | Estimated experimental $\tau_\mathrm{F}$, min | Estimated experimental $\tau_\mathrm{F}$, h | Estimated experimental $\tau_\mathrm{F}$, d |
|---|---|---|---|---|---|
| 1I6O | 4.03E+04 | 1.60E+02 | 2.66E+00 | 4.44E-02 | 1.85E-03 |
| 1EF9 | 4.66E+04 | 1.85E+02 | 3.08E+00 | 5.13E-02 | 2.14E-03 |
| 1ORO | 4.82E+04 | 1.91E+02 | 3.18E+00 | 5.31E-02 | 2.21E-03 |
| 1SV6 | 4.94E+04 | 1.96E+02 | 3.26E+00 | 5.44E-02 | 2.27E-03 |
| 1AH9 | 6.57E+04 | 2.61E+02 | 4.34E+00 | 7.24E-02 | 3.02E-03 |
| 3GN5 | 7.30E+04 | 2.90E+02 | 4.83E+00 | 8.04E-02 | 3.35E-03 |
| 1ZYL | 8.02E+04 | 3.18E+02 | 5.30E+00 | 8.84E-02 | 3.68E-03 |
| 2PTQ | 5.28E+05 | 2.09E+03 | 3.49E+01 | 5.82E-01 | 2.42E-02 |
| 2V81 | 5.58E+05 | 2.22E+03 | 3.69E+01 | 6.15E-01 | 2.56E-02 |
| 1FTS | 4.60E+06 | 1.83E+04 | 3.04E+02 | 5.07E+00 | 2.11E-01 |
| 3M7M | 7.33E+13 | 2.91E+11 | 4.85E+09 | 8.08E+07 | 3.36E+06 |
| 4IWX | 7.61E+13 | 3.02E+11 | 5.03E+09 | 8.39E+07 | 3.50E+06 |
| 1KSF | 2.59E+14 | 1.03E+12 | 1.71E+10 | 2.85E+08 | 1.19E+07 |
| 3BRQ | 3.83E+18 | 1.52E+16 | 2.53E+14 | 4.22E+12 | 1.76E+11 |
| 2FYM | 1.28E+19 | 5.06E+16 | 8.44E+14 | 1.41E+13 | 5.86E+11 |
| 1CLI | 2.35E+19 | 9.31E+16 | 1.55E+15 | 2.59E+13 | 1.08E+12 |
| 2O1C | 3.30E+19 | 1.31E+17 | 2.18E+15 | 3.64E+13 | 1.52E+12 |
| 1NG9 | 1.35E+20 | 5.37E+17 | 8.94E+15 | 1.49E+14 | 6.21E+12 |
| 1U60 | 1.65E+20 | 6.53E+17 | 1.09E+16 | 1.81E+14 | 7.56E+12 |
| 1K7J | 6.72E+20 | 2.67E+18 | 4.45E+16 | 7.41E+14 | 3.09E+13 |
| 1W78 | 7.66E+20 | 3.04E+18 | 5.07E+16 | 8.45E+14 | 3.52E+13 |
| 2HNH | 7.85E+20 | 3.11E+18 | 5.19E+16 | 8.65E+14 | 3.60E+13 |
| 1U0B | 2.82E+21 | 1.12E+19 | 1.87E+17 | 3.11E+15 | 1.30E+14 |
| 1UUF | 7.80E+21 | 3.09E+19 | 5.15E+17 | 8.59E+15 | 3.58E+14 |
| 2ID0 | 4.21E+22 | 1.67E+20 | 2.78E+18 | 4.64E+16 | 1.93E+15 |
| 1QF6 | 7.43E+22 | 2.95E+20 | 4.91E+18 | 8.19E+16 | 3.41E+15 |
| 2HG2 | 2.83E+23 | 1.12E+21 | 1.87E+19 | 3.12E+17 | 1.30E+16 |
| 2QCU | 1.66E+24 | 6.60E+21 | 1.10E+20 | 1.83E+18 | 7.64E+16 |
| 2WIU | 4.80E+27 | 1.91E+25 | 3.18E+23 | 5.29E+21 | 2.20E+20 |
| 1GER | 3.73E+28 | 1.48E+26 | 2.47E+24 | 4.11E+22 | 1.71E+21 |
| 1YQQ | 2.34E+29 | 9.30E+26 | 1.55E+25 | 2.58E+23 | 1.08E+22 |

**Table S6.** Proteins with at least one misfolded trajectory expected to bypass chaperones

| Category | Number of unique proteins | PDB IDs |
|---|---|---|
| Not expected to interact with TF | 70 | 2ID0, 4IWX, 1A69, 2WIU, 1U60, 4DZD, 2HNH, 3HWO, 1KSF, 1ORO, 1ZYL, 3BRQ, 1XRU, 1EF9, 2PTQ, 2FYM, 1NG9, 1DXE, 2JO6, 4A2C, 1PMO, 2KX9, 2R5N, 1K7J, 1GT7, 2WW4, 3M7M, 3PCO, 1GLF, 1GER, 2H1F, 1YQQ, 1DFU, 1SVT, 2V81, 1NAQ, 1U0B, 1PF5, 1FJJ, 2YVA, 1DUV, 1UUF, 3BMB, 1A6J, 1SV6, 1W78, 4IM7, 2KFW, 1Q5X, 1P7L, 1QF6, 1T4B, 4FZW, 1AH9, 1FTS, 2QCU, 1CLI, 2PTH, 2O1C, 2HNA, 1W8G, 3GN5, 1WOC, 4DCM, 1A82, 2HG2, 1L6W, 1D2F, 1I6O, 1M3U |
| Not expected to interact with GroEL/GroES | 80 | 2ID0, 1XVI, 4IWX, 1A69, 1FUI, 2WIU, 1U60, 4DZD, 2HNH, 3HWO, 1KSF, 1ORO, 1P91, 1ZYL, 3BRQ, 1XRU, 1EF9, 2PTQ, 2FYM, 1NG9, 1DXE, 4HR7, 2JO6, 2HG2, 4A2C, 1PMO, 2KX9, 2R5N, 1K7J, 1GT7, 1GQT, 2WW4, 3M7M, 3PCO, 1GLF, 1GER, 2H1F, 1YQQ, 1DFU, 1SVT, 2V81, 1NAQ, 1U0B, 1PF5, 1FJJ, 2YVA, 1DUV, 1UUF, 3BMB, 1A6J, 1H16, 1SV6, 1W78, 4IM7, 2KFW, 1Q5X, 1P7L, 1QF6, 2PTH, 3GN5, 4FZW, 1AH9, 1FTS, 2QCU, 1GZ0, 1CLI, 2O1C, 2HNA, 4E8B, 1W8G, 1WOC, 4DCM, 1A82, 1L6W, 1T4B, 1D2F, 1I6O, 4KN7, 1GYT, 1M3U |
| Not expected to interact with DnaK | 74 | 2ID0, 1XVI, 1A69, 1FUI, 2WIU, 1U60, 4DZD, 2HNH, 3HWO, 1KSF, 1ORO, 1P91, 1ZYL, 3BRQ, 1XRU, 1EF9, 2PTQ, 2FYM, 4HR7, 1NG9, 1DXE, 2JO6, 2HG2, 4A2C, 1PMO, 2KX9, 2R5N, 1K7J, 1GT7, 2WW4, 3M7M, 3PCO, 2H1F, 1YQQ, 1DFU, 1SVT, 2V81, 1NAQ, 1U0B, 1PF5, 1FJJ, 1DUV, 1UUF, 3BMB, 1H16, 1SV6, 1W78, 4IM7, 1Q5X, 1P7L, 1QF6, 2PTH, 3GN5, 4FZW, 1AH9, 2QCU, 1GZ0, 2O1C, 1A6J, 4E8B, 1W8G, 1FTS, 1GYT, 1WOC, 4DCM, 1A82, 2HNA, 1L6W, 1T4B, 1D2F, 1GER, 4KN7, 1CLI, 1M3U |
| Not expected to interact with TF, GroEL/GroES, or DnaK | 64 | 2ID0, 1A69, 2WIU, 1U60, 4DZD, 2HNH, 3HWO, 1KSF, 1ORO, 1ZYL, 3BRQ, 1XRU, 1EF9, 2PTQ, 2FYM, 1NG9, 1DXE, 2JO6, 4A2C, 1PMO, 2KX9, 2R5N, 1K7J, 1GT7, 2WW4, 3M7M, 3PCO, 2H1F, 1YQQ, 1DFU, 1SVT, 2V81, 1NAQ, 1U0B, 1PF5, 1FJJ, 1DUV, 1UUF, 3BMB, 1SV6, 1W78, 4IM7, 1Q5X, 1P7L, 1QF6, 4FZW, 1AH9, 2QCU, 2PTH, 2O1C, 1A6J, 1W8G, 1FTS, 3GN5, 1WOC, 4DCM, 1A82, 2HNA, 1L6W, 1T4B, 1D2F, 1GER, 1CLI, 1M3U |

**Table S7**. Proteins with at least one misfolded trajectory expected not to aggregate or be degraded

| Category | Number of unique proteins | PDB IDs |
|---|---|---|
| Not expected to aggregate | 68 | 2ID0, 1XVI, 1A69, 1FUI, 2WIU, 1U60, 4DZD, 2HNH, 3HWO, 1KSF, 1ORO, 1P91, 1ZYL, 3BRQ, 1XRU, 2PTQ, 2FYM, 4HR7, 1NG9, 2JO6, 2HG2, 4A2C, 1PMO, 2KX9, 2R5N, 1K7J, 1GT7, 2WW4, 3M7M, 3PCO, 1DFU, 1SVT, 1U0B, 1PF5, 1FJJ, 1DUV, 1SV6, 1W78, 4IM7, 1YQQ, 2KFW, 1Q5X, 1P7L, 1QF6, 2PTH, 3GN5, 4FZW, 1AH9, 1FTS, 2QCU, 1GZ0, 1CLI, 2O1C, 2HNA, 1A6J, 1W8G, 1WOC, 4DCM, 1A82, 1L6W, 1T4B, 1D2F, 1NAQ, 1I6O, 1GER, 4KN7, 1GYT, 1M3U |
| Not expected to be degraded | 70 | 2ID0, 1XVI, 1A69, 1FUI, 2WIU, 1U60, 4DZD, 2HNH, 3HWO, 1KSF, 1ORO, 1P91, 1ZYL, 3BRQ, 1XRU, 2PTQ, 2FYM, 1NG9, 4HR7, 2JO6, 2HG2, 4A2C, 1PMO, 2KX9, 2R5N, 1K7J, 1GT7, 2WW4, 3M7M, 3PCO, 1GER, 1YQQ, 1DFU, 1SVT, 1U0B, 1PF5, 1FJJ, 1DUV, 1SV6, 1W78, 4IM7, 2KFW, 1Q5X, 1NAQ, 1P7L, 1QF6, 2PTH, 3GN5, 4FZW, 1AH9, 1FTS, 2QCU, 1GZ0, 1CLI, 2O1C, 1A6J, 4E8B, 1W8G, 1WOC, 4DCM, 1A82, 2HNA, 1L6W, 1T4B, 1UUF, 1D2F, 1I6O, 4KN7, 1GYT, 1M3U |

**Table S8**. Proteins with misfolded conformations expected to bypass proteostasis machinery and remain soluble but less functional

| Category | Number of unique proteins | PDB IDs |
|---|---|---|
| Expected to bypass chaperones, not aggregate, and not be degraded | 57 | 2ID0, 1A69, 2WIU, 1U60, 4DZD, 2HNH, 3HWO, 1KSF, 1ORO, 1ZYL, 3BRQ, 1XRU, 2PTQ, 2FYM, 1NG9, 2JO6, 4A2C, 1PMO, 2KX9, 2R5N, 1K7J, 1GT7, 2WW4, 3M7M, 3PCO, 1DFU, 1SVT, 1U0B, 1PF5, 1FJJ, 1DUV, 1SV6, 1W78, 4IM7, 1YQQ, 1Q5X, 1P7L, 1QF6, 4FZW, 1AH9, 2QCU, 2PTH, 2O1C, 1A6J, 1W8G, 1FTS, 3GN5, 1WOC, 4DCM, 1A82, 2HNA, 1L6W, 1T4B, 1D2F, 1GER, 1CLI, 1M3U |
| Expected to exhibit reduced function | 69 | 1XVI, 4IWX, 1A69, 1FUI, 2WIU, 1U60, 4DZD, 2HNH, 3HWO, 2ID0, 1ZYL, 1XRU, 1P91, 1EF9, 2FYM, 4HR7, 1NG9, 1DXE, 2HG2, 1PMO, 2R5N, 1K7J, 1GQT, 3PCO, 1GLF, 2H1F, 1YQQ, 1DFU, 1SVT, 2V81, 1NAQ, 1U0B, 2YVA, 4KN7, 1DUV, 1UUF, 3BMB, 1A6J, 1H16, 1SV6, 1W78, 4IM7, 1Q5X, 2PTH, 1T4B, 4FZW, 1FTS, 2QCU, 1P7L, 1GZ0, 1KSF, 1CLI, 1GYT, 1WOC, 2PTQ, 1A82, 2WW4, 4A2C, 3GN5, 1L6W, 1GT7, 1QF6, 1D2F, 1I6O, 1GER, 2KX9, 2O1C, 1PF5, 3M7M |
| Expected to bypass proteostasis machinery and to exhibit reduced function | 38 | 1A69, 2WIU, 1U60, 4DZD, 3HWO, 1ZYL, 2FYM, 1NG9, 1PMO, 2R5N, 1K7J, 1DFU, 1SVT, 1U0B, 1SV6, 1W78, 4IM7, 1YQQ, 1Q5X, 4FZW, 2QCU, 1KSF, 2PTH, 1FTS, 2HNH, 1WOC, 2PTQ, 1A82, 3GN5, 3PCO, 1L6W, 1T4B, 1A6J, 1GT7, 1D2F, 1GER, 1DUV, 2O1C |

**Table S9**. Functional residues database terms and descriptions

| Term in database | Description |
|---|---|
| PDB used in model | Uniprot: PDB ID used in the simulation |
| Gene name | Uniprot: Name of gene |
| Protein Uniprot ID | Uniprot: ID associated with gene in uniprot |
| Ordered Locus | Uniprot: Ordered locus associated with gene |
| Function | Uniprot: Function of the gene product |
| Catalytic activity | Uniprot: Catalytic activity of protein |
| Subunit | Uniprot: Information about n-mers structure |
| GO functions | Uniprot: Gene ontology functions |
| Binding site | Uniprot: Information about mapped and unmapped binding site |
| Active site | Uniprot: Information about mapped and unmapped active site |
| Metal binding site | Uniprot: Information about mapped and unmapped metal binding site |
| Site | Uniprot: Information about mapped and unmapped site |
| Absorption | Uniprot: Information about ligands absorbed |
| Calcium binding | Uniprot: Information about calcium binding |
| DNA binding | Uniprot: Information about DNA binding |
| EC number | Uniprot: Enzyme Commission number |
| Metal binding | Uniprot: Information about mapped and unmapped metal binding |
| Nucleotide binding | Uniprot: Information about mapped and unmapped about nucleotide binding |
| Pathway | Uniprot: Information about mechanism pathway |
| pH dependence | Uniprot: Information about the protein function's pH dependence |
| Temperature dependence | Uniprot: Information about the protein function's temperature dependence |
| Cofactor | Uniprot: Information about the protein's cofactors |

**Table S9** continued

| Term in database | Description |
|---|---|
| Kinetics | Uniprot: Information about the kinetics of the protein |
| mRNA sequence | mRNA sequence of the gene product |
| AA sequence | Amino acid sequence of the gene product |
| CIF sequence of # - X | PDB: Amino acid sequence of the chain X in CIF # |
| Unmapped active residues from CIF # | PDB: Chain ID, Residue # in CIF file, Residue Name, Ligand in contact with |
| Mapped active residues from CIF # | PDB: Chain ID, Mapped residue #, Residue Name, Ligand in contact with |
| Ligand abbreviation from CIF # | PDB: 3 letter abbreviation, ligand name |
| Unmapped interfacial residues from CIF # | PDB: Chain 1, residue name, unmapped residue # in CIF file, Chain 2, residue name, unmapped residue # in CIF file |
| Mapped interfacial residues from CIF # | PDB: Chain 1, residue name, mapped residue, Chain 2, residue name, unmapped residue # in CIF file |
| Small molecule active residues (mapped) | PDB: List of residues in contact with small molecules and ligands |
| Interfacial active residues (mapped) | PDB: List of interfacial residues in the crystal structure |
| Combined active residues (mapped) | Uniprot and PDB: List of functional residues from Uniprot, residues interacting with small molecules and interfacial residues |
