## Supplementary Reference Material 2 for "Subpopulations of soluble, misfolded proteins commonly bypass chaperones: How it happens at the molecular level"

### How synonymous mutations alter enzyme structure and function over long time scales

Yang Jiang[1], Syam Sundar Neti[1], Priya Pradhan[1], Squire J. Booker[1, 2, 3] and Edward P. O'Brien[1, 4, 5, *]

[1] Department of Chemistry, Penn State University, University Park, Pennsylvania, United States

[2] Department of Biochemistry and Molecular Biology, Pennsylvania State University, University Park, Pennsylvania, United States

[3] Howard Hughes Medical Institute, Pennsylvania State University, University Park, Pennsylvania, United States

[4] Bioinformatics and Genomics Graduate Program, The Huck Institutes of the Life Sciences, Penn State University, University Park, Pennsylvania, United States

[5] Institute for Computational and Data Sciences, Penn State University, University Park, Pennsylvania, United States

#### Abstract

The specific activity of enzymes can be altered over long time scales in cells by synonymous mutations, which change an mRNA molecule's sequence but not the encoded protein's primary structure. How this happens at the molecular level is unknown. Here, we resolve this issue by applying multiscale modeling to three *E. coli* enzymes - type III chloramphenicol acetyltransferase, D-alanine–D-alanine ligase B, and dihydrofolate reductase. This modeling involves coarse-grained simulations of protein synthesis and post-translational behavior, all-atom simulations as a test of robustness, and QM/MM calculations to characterize function. We first demonstrate that our model accurately predicts experimentally measured changes in specific activity due to synonymous mutations. Then, we show that changes in codon translation rates induced by synonymous mutations cause shifts in co-translational and post-translational folding pathways that kinetically partition molecules into subpopulations that very slowly interconvert to the native, functional state. These long-lived states exhibit reduced catalytic activity, as demonstrated by their increased activation energies for the reactions they carry out. Structurally, these states resemble the native state, with localized misfolding near the active sites of the enzymes. The localized misfolding involves noncovalent lasso entanglements - a topology in which the protein backbone forms a loop closed by noncovalent native contacts which is then threaded by another portion of the protein. Such entanglements are often kinetic traps, as they can require a large proportion of the protein to unfold, which is energetically unfavorable, before they disentangle and attain the native state. The near-native-like structures of these misfolded states allow them to bypass the proteostasis machinery and remain soluble, as they exhibit similar hydrophobic surface areas as the native state. These entangled structures persist in all-atom simulations as well, indicating that these conclusions are independent of model resolution. Thus, synonymous mutations cause shifts in the co- and post-translational structural ensemble of proteins, whose altered subpopulations lead to long-term changes in the specific activities of some enzymes. The formation of entangled subpopulations is therefore a mechanism through which changes in translation elongation rate alter ensemble-averaged specific activities, which can ultimately affect the efficiency of biochemical pathways and phenotypic traits.

#### Introduction

In both *in vitro* and in vivo experiments, the specific activity of protein enzymes – that is, the catalytic turnover per unit time per unit mass of soluble protein – can change depending on the codons used to encode the protein[1, 2, 3, 4, 5, 6]. The specific activity of the *E. coli* enzyme type III chloramphenicol acetyltransferase (CAT-III), for example, decreases by approximately 20% when fast-translating synonymous mutations are introduced into its transcript[1, 6]. Synonymous mutations change the sequence of nucleotides composing an mRNA molecule, which in turn changes the speed at which translation elongation occurs[7] but not the protein primary structure. Specific activity measurements control for differences in protein expression and the formation of insoluble aggregates through centrifugation or gel separation[1, 2, 3, 8]. For enzymes that do not require post-translational modifications, these observed changes in specific activity indicate that inside cells, newly expressed proteins can populate long-lived conformational states that are not native, have reduced functionality, somehow bypass the chaperone and degradation machinery, and do not aggregate. Furthermore, these observations indicate that the distribution of these kinetically trapped states is sensitive to changes in translation elongation speed.

This fourth state of soluble proteins is distinct from the three typical states[9] of a protein being either folded and functional, misfolded and aggregated, or degraded. The structural, kinetic, and energetic properties of this fourth state of proteins is a fundamental, unanswered question in biochemistry and molecular biology. Soluble, nonfunctional states do not just occur for enzymes, other protein functions can also be altered. The hormone-transporting protein transthyretin, for example, can have 20% of its soluble fraction in a nonfunctional state[10]. There are hints in the literature of some of the structural properties of this fourth state. NMR-derived structures of β-gamma-crystallin produced from two synonymous mRNA variants found that it populated two conformations differing in only the formation of a disulfide bond[11]. Disulfide bonds represent an energetically strong constraint on folding topologies and are uncommon, only 5% and 20%, respectively, of *E. coli* and human cytoplasmic proteins contain disulfide bonds[12]. Furthermore, many enzymes that exhibit specific activity changes due to synonymous mutations do not contain disulfide bonds[1, 2, 3, 5]. Thus, the nature of these structurally altered subpopulations has not yet been resolved, although experimental data rule out misfolding involving quaternary structures. For example, gel separation and chromatography experiments rule out the formation of off-pathway dimers and higher-order oligomers for many enzymes that normally function in a monomeric state[1, 2, 4, 8].

Here, we use a novel multiscale approach across the dimensions of time, space, and energy to simulate the synthesis, post-translational behavior, and function of enzymes under different translation rate schedules that arise from synonymous codons. We first establish that our modeling approach can correctly predict the experimentally measured changes in specific activity for CAT-III and D-alanine–D-alanine ligase B (DDLB). We then apply the model to enzymes whose activity is sensitive to synonymous mutations and enzymes whose activity is not. Dissecting the structures and kinetics of co- and post-translational folding that occur in our simulations, we show that synonymous mutations shift the folding pathways and populations of near-native non-covalent lasso entanglements, each of which has its own intrinsic activity ($k_{cat}$) and leads to long-term changes in the enzymatic specific activity.

#### Methods

**Coarse-grained (CG) simulation model.** To model transitions between the native state and the unfolded state, we utilized a Gō-based CG model[13, 14, 15, 16, 17] for all the proteins studied. Briefly, this CG model represents each residue as a single interaction site centered on the Cα position and uses a structure-based potential energy function (see Supplementary equation (1)). The force field parameters were optimized by training the parameters to reproduce the folding stability free energies of 18 small single-domain proteins[18], followed by assignment of the minimum values that can reproduce the structural stability for a given protein. To map the simulation timescale to an experimental timescale, we use the scaling factor $\alpha$, which is the ratio of bulk experimental folding times to simulated folding times (see Supplementary Table 4). The high-resolution crystal structure PDB 4v9d[19] was used to coarse-grain the *E. coli* ribosome, with the A- and P-site tRNA molecules modeled based on the PDB structure 5jte.[20] The entire 50S subunit, including the A- and P-site tRNAs, was coarse-grained using the three/four-point model of ribosomal RNA[13] and the Cα model for ribosomal protein[13] and then truncated to include only those CG interaction sites within 30 Å of the centerline of the exit tunnel, within 20 Å of the peptidyl transferase center (PTC, identified as A2602 in the 23S rRNA of the *E. coli* ribosome), and the interaction sites at the ribosome surface near the exit tunnel opening. This resulted in 4,577 interaction sites for the cropped CG *E. coli* ribosome.

Protein synthesis on the ribosome was modeled using a continuous synthesis protocol that describes A-site tRNA binding, peptidyl transfer, tRNA translocation, and ribosome trafficking[21] at each nascent chain length using codon translation rates obtained from the model of Fluitt & Viljoen[22] (see Supplementary equations (11) and (13)). Post-translational dynamics were modeled by simulating the nascent protein in the absence of the ribosome. Simulations for both co- and post-translational folding were performed via Langevin dynamics with a collision frequency of 0.05 ps$^{-1}$ and a time step of 15 fs using OpenMM[23]. Details of the model setup can be found in Supplementary Methods Sections 1 to 8.

**Virtual screening for enzymes that exhibit kinetic traps.** To identify candidate enzymes that may have kinetic traps, we created a dataset of well-characterized monomeric *E. coli* enzymes by searching the relevant databases EzCatDB[24, 25], UniProt[26] and RCSB PDB[27]. The wild-type enzymes were then parameterized, and their synthesis and post-translational dynamics were simulated with a 14-day wall time. The candidates were selected by using the scoring function

$$\text{Score} = \frac{1}{2} \times \left[ \left( 1 - e^{-\left\langle \tau_{\text{F}}^{\text{post}} \right\rangle} \right) + \Theta_{\text{binding}} \right], \tag{1}$$

where $\left\langle \tau_{\text{F}}^{\text{post}} \right\rangle$ is the mean post-translational folding time calculated by using the double-pathway kinetics scheme of Supplementary equation (6) with no delay time (*i.e.*, $t_1 = t_2 = 0$) and $\Theta_{\text{binding}}$ is an indicator of whether (equals 1) or not (equals 0) misfolding occurs at or near the substrate binding site. A higher score indicates a higher possibility for the enzyme to have long-lived kinetic traps that have misfolding in the substrate binding pocket during its post-translational folding dynamics. Further details of the screening can be found in Supplementary Methods Section 9.

**Characterizing post-translational misfolded structures.** The misfolded structures of the post-translational folding process were characterized by using the metrics $G$ and $Q_{\text{act}}$. $G$ is an order parameter that measures the extent to which there is a change of entanglement in a given structure compared to the native structure and is calculated as

$$G = \frac{1}{N} \sum_{(i,j)} \Theta\left( (i,j) \in nc \cap g(i,j) \neq g^{\text{native}}(i,j) \right), \tag{2}$$

where $(i, j)$ is one of the native contacts in the native crystal structure; $nc$ is the set of native contacts formed in the current structure; $g(i,j)$ and $g^{\text{native}}(i,j)$ are, respectively, the total linking number of the native contact $(i, j)$ in the current and native structures estimated using

Supplementary equation (16) (see Supplementary Methods Section 11 for details); $N$ is the total number of native contacts within the native structure; and the selection function $\Theta$ equals 1 when the condition is true and 0 when it is false. The larger $G$ is the larger the number of residues that have changed their entanglement status relative to the native state. That is $G$ reports on the presence of non-native entanglements in structures.

$Q_{act}$ is the fraction of native contacts that have formed in the enzyme substrate binding pocket. Residues composing the substrate binding pocket were identified as those residues within 8 Å of any atoms of the relevant ligands present in the crystal structure. The $Q_{act}$ values were calculated for all the native contacts between one atom within the substrate binding pocket and any other atom, as shown below:

$$Q_{\text{act}} = \frac{\sum_{i\in I}\sum_{j\in J}\Theta(i,j|\text{Current})}{\sum_{i\in I}\sum_{j\in J}\Theta(i,j|\text{Native})}, \tag{3}$$

where $i$ and $j$ are the residue indices and satisfy $j > i + 3$; $I$ is the intersection set of residues within secondary structure elements (α-helical or β-strands) and the substrate binding pocket; $J$ is the set of residues within secondary structure elements; $\Theta(i,j|\text{Current})$ and $\Theta(i,j|\text{Native})$ are step functions that equal 1 when residue $i$ and $j$ have native contact and 0 when $i$ and $j$ do not have native contact in the current structure and native structure, respectively. Native contacts are considered formed when the distance between the Cα atoms of residues $i$ and $j$ does not exceed 1.2 times their native distance and the native distance does not exceed 8 Å. In the case of CAT-III, residue set $I$ also includes native contacts in the trimer interface region (residues 25 to 33 and 150 to 157) to monitor the folding of the trimer interface as well. Note that the native contacts used in estimating $Q_{act}$ are restricted to those within secondary structure elements, while the entire set of native contacts is used to estimate $G$. Details of the assessment procedure can be found in Supplementary Methods Section 12.

**Specific activity estimation.** For each metastable state $i$, identified using the Markov state modeling procedure reported in Supplementary Methods Section 12, we randomly select five conformations from all the microstates based on the probability distribution of the microstates within the metastable state and back-map them to all-atom structures (see Supplementary Methods Section 13). We then used QM/MM simulations (see Supplementary Methods Section 14) to calculate the transition state barrier for each of the five conformations, and from these, we determined the median activation barrier height $\Delta G_i^{\ddagger}$. Only the metastable states that form a near-native active site ($Q_{act}$ ≥ 0.6), as well as the native state, were taken to estimate $\Delta G_i^{\ddagger}$, while the others were considered having an infinite barrier height (zero reaction rate). Assuming the rate constants of each metastable state have similar pre-exponential factors that can be treated as a constant, the specific activity for a protein can be estimated by using the probability distribution of metastable states (state probability $p_i$) and the activation free energy barrier height ($\Delta G_i^{\ddagger}$) of each state as

$$\text{Specific activity} = \frac{A}{m}\sum_{i=1}^{N} p_i \cdot e^{-\frac{1}{RT}\Delta G_i^{\ddagger}}, \tag{4}$$

where $m$ is the molecular weight of the enzyme; $A$ is the pre-exponential factor allowing us to convert $\Delta G_i^{\ddagger}$ to the reaction rate constant; $N$ is the total number of metastable states; $R$ is the gas constant; and $T$ is the temperature. In many codon usage studies[1, 3, 8], the relative specific activity is often used to compare the enzymatic activities of a protein produced by synonymous mRNA variants. In this study, the maximum specific activity among the fast and slow synonymous mRNA variants of an enzyme was used for normalization:

$$\begin{cases} \mathrm{SA}^*_{\mathrm{fast}} = \dfrac{\mathrm{SA}_{\mathrm{fast}}}{\max\{\mathrm{SA}_{\mathrm{fast}}, \mathrm{SA}_{\mathrm{slow}}\}} \\ \mathrm{SA}^*_{\mathrm{slow}} = \dfrac{\mathrm{SA}_{\mathrm{slow}}}{\max\{\mathrm{SA}_{\mathrm{fast}}, \mathrm{SA}_{\mathrm{slow}}\}} \end{cases}, \tag{5}$$

where $\mathrm{SA}^*$ is the relative specific activity. Note that the pre-exponential factor $A$ and the molecular weight $m$ cancel out when specific activity is normalized; therefore, they do not need to be estimated. The details of estimating the state probability $p_i$ (accounting for the soluble fraction only) and the activation free energy barrier height ($\Delta G_i^{\ddagger}$) are presented in Supplementary Methods Section 14. The materials and experimental methods for measuring the specific activity of DDLB variants are presented in Supplementary Methods Section 18.

#### Results

**Recapitulating experimental trends of CAT-III.** We first applied our modeling protocol to the *E. coli* enzyme CAT-III to test if the method accurately predicts the influence of synonymous mutations on specific activity. CAT-III has been studied experimentally and shown to have a reduced specific activity when faster-translating codons are introduced through synonymous mutations[1]. We created both fast- and slow-translating synonymous mRNA sequences (denoted, respectively, CAT-III$_{\mathrm{fast}}$ and CAT-III$_{\mathrm{slow}}$) by replacing each wild-type codon with its fastest or slowest synonymous variant (see Fig. 1b and Supplementary Methods Section 19). The resulting slow-mRNA variant takes two times longer to translate than the fast variant (see Fig. 1f). We simulated the synthesis and post-translational behavior of CAT-III resulting from the fast- and slow-mRNA variants and calculated their respective specific activities (equation (4)) at the end of the post-translational simulations. In our model, CAT-III$_{\mathrm{fast}}$ exhibited 81.4% (95% confidence interval (CI): [70.0%, 94.8%] from bootstrapping, $p$-value=0.0035 from random permutation test, $10^6$ permutations) of the specific activity of CAT-III$_{\mathrm{slow}}$ (Fig. 1g). This comparison, known as the 'relative specific activity' (equation (5)), is common in biochemical studies[1, 3]. A 20% decrease in activity has been observed experimentally[1]. Thus, our modeling approach qualitatively recapitulates experimentally observed changes in enzyme activity due to synonymous mutations.

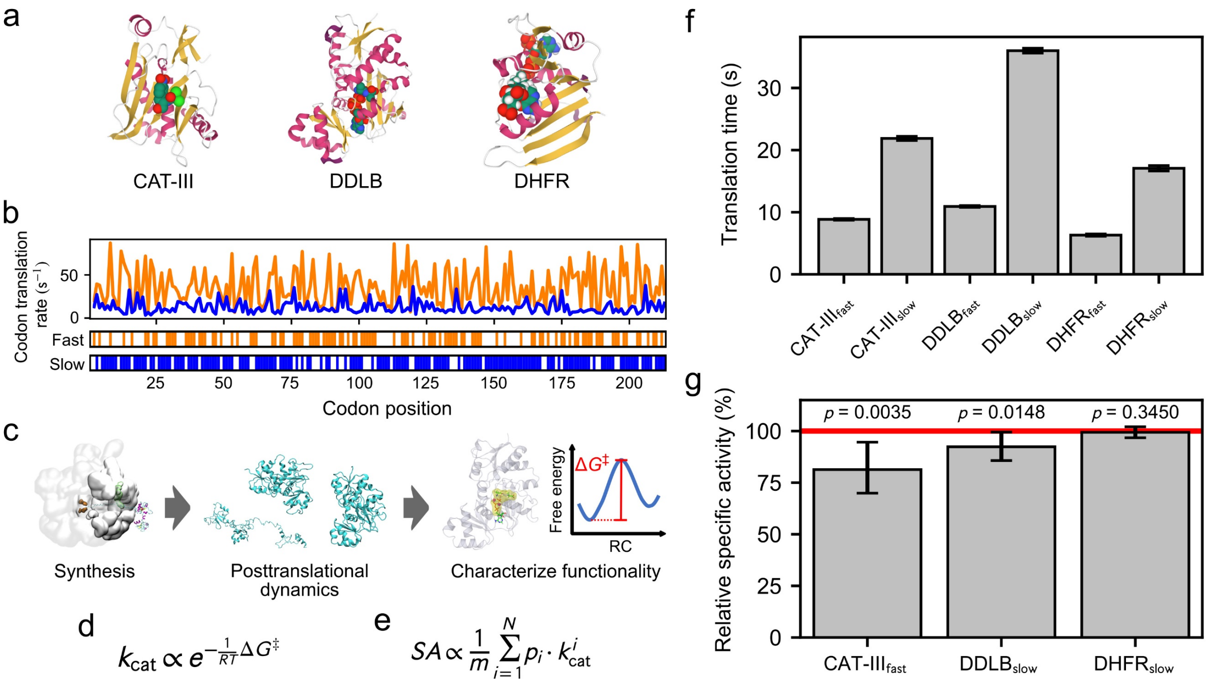

**Figure 1**. **A multiscale approach for understanding the influence of synonymous codons on the structure and function of enzymes.** a, Crystal structures of the three enzymes investigated in this study, with secondary structure elements highlighted and substrates presented; b, codon translation rate of CAT-III fast and slow synonymous mRNA variants, with the mutation sites presented on the bottom bars; c scheme diagram of the multiscale approach, including the CG protein synthesis and post-translational dynamics, and the all-atom characterization of enzymatic functionality; d, enzymatic reaction rate $k_{cat}$ estimated from the activation free energy $\Delta G^{\ddagger}$; e, enzyme specific activity (SA) estimated as the ensemble average of reaction rates $k_{cat}$ (see equation (4)); f, comparison of the translation time for the fast and slow synonymous mRNA variants; g, relative SAs for CAT-III, DDLB and DHFR. The SA values are normalized to the higher SA value found in fast- and slow-mRNA variants. Error bars in panels f and g are 95% CIs about the mean, which were estimated using bootstrapping. p-values characterize the difference in SAs between the proteins produced from the fast and slow variants. p-values were calculated using the permutation test. The SAs of CAT-III and DDLB are sensitive to translation speed changes, and that of DHFR is not.

**Other enzymes that are sensitive to changes in translation speed.** We next applied our model to identify other *E. coli* enzymes whose specific activities are sensitive to translation speed changes. To do this, we developed a virtual screening strategy that identifies misfolding-prone proteins exhibiting long-lived, post-translational kinetic traps (see Methods), as we hypothesized that the activity of these enzymes is more likely to be sensitive to translation speed changes. Due to the large number of *E. coli* enzymes that have an unknown catalytic mechanism or uncharacterized protein-substrate complex structures, we focused on only 14 well-characterized monomeric enzymes (Table 1). We synthesized each protein on the ribosome using the translation rate arising from their wild-type mRNA sequences, followed by release from the ribosome and a post-translational simulation phase. To identify good candidates, we scored (equation (1)) each of the 14 enzymes based on whether they misfolded near the substrate binding site and whether the misfolding events were long-lived. Porphobilinogen deaminase never exhibited any folding events during the simulations and therefore was excluded from further study. The protein with the highest score, D-alanine–D-alanine ligase B (DDLB), was selected as a candidate whose enzymatic activity we hypothesized is likely to be sensitive to translation speed changes, whereas the protein with the lowest score, dihydrofolate reductase (DHFR), was identified as an enzyme whose activity is likely to be insensitive to synonymous mutations.

**Table 1.** The 14 *E. coli* enzymes studied using our virtual screening strategy. $\langle\tau_F^{\text{post}}\rangle$, calculated post-translational mean folding time. $\Theta_{\text{binding}}$ indicates whether misfolding is ('1') or is not ('0') located at or near the substrate binding site. 'Score' is the scoring function calculated from equation (1).

| Protein Name | Length (residues) | PDB ID | $\langle\tau_F^{\text{post}}\rangle$ (s) | $\Theta_{\text{binding}}$ | Score |
|---|---|---|---|---|---|
| D-alanine─D-alanine ligase B (DDLB) | 306 | 4C5C | $9.8 \times 10^9$ | 1 | 1.00 |
| Quinone oxidoreductase 2 | 303 | 2ZCV | 3.9 | 1 | 0.99 |
| Peptide deformylase | 169 | 3K6L | 3.4 | 1 | 0.98 |
| Phosphoenolpyruvate carboxykinase | 540 | 1AQ2 | $5.2 \times 10^7$ | 0 | 0.50 |
| Adenylate kinase | 214 | 1AKE | 9.5 | 0 | 0.50 |
| Dephospho-CoA kinase | 206 | 1VHL | 4.8 | 0 | 0.50 |
| Protein-L-isoaspartate O-methyltransferase | 208 | 3LBF | 3.4 | 0 | 0.48 |
| Methionyl-tRNA formyltransferase | 315 | 2FMT | 1.0 | 0 | 0.32 |
| Phosphoribosylglycinamide formyltransferase | 212 | 1C2T | 0.65 | 0 | 0.24 |
| Flavodoxin/ferredoxin--NADP reductase | 248 | 1FDR | 0.09 | 0 | 0.05 |
| DNA-3-methyladenine glycosylase 2 | 282 | 3CW7 | 0.07 | 0 | 0.03 |
| 2-Dehydropantoate 2-reductase | 303 | 2OFP | 0.05 | 0 | 0.02 |
| Dihydrofolate reductase (DHFR) | 159 | 4KJK | 0.03 | 0 | 0.01 |
| Porphobilinogen deaminase | 313 | 1PDA | - | - | - |

Next, we simulated the synthesis and post-translational dynamics of DDLB and DHFR from their fast- and slow-translating mRNA variants (sequences are presented in Supplementary Methods Section 19) using the same simulation protocol as applied to CAT-III. The slow variants of DDLB and DHFR translate, respectively, three and two times slower than their fast variants (Fig. 1f). We found that the specific activity of $\text{DDLB}_{\text{slow}}$ was 92.4% (95% CI: [85.7%, 99.3%]) that of $\text{DDLB}_{\text{fast}}$ ($p$-value = 0.0148) 60 s after synthesis was completed. $\text{DHFR}_{\text{slow}}$ exhibited a specific activity 99.4% (95% CI: [96.7%, 102.1%]) that of $\text{DHFR}_{\text{fast}}$, which were not significantly different ($p$-value = 0.3450). Therefore, our model describes enzymes whose specific activity is either sensitive (DDLB) or insensitive (DHFR) to changes in translation speed over long-time scales.

**Accurate prediction of changes in specific activity.** To experimentally test if $\text{DDLB}_{\text{slow}}$ has a reduced enzymatic activity compared to $\text{DDLB}_{\text{fast}}$ we recombinantly expressed the fast and slow DDLB variants in *E. coli* using the same mRNA sequences as in the simulations, purified the enzyme, and then assayed the reaction kinetics (see Supplementary Methods Section 18). We observe that the fast variant had a higher level of protein expression 5 hours after induction than the slow variant, consistent with the fast mRNA variant translating faster (Supplementary Fig. 12). The reaction rate constant $k_{\text{cat}}$ was measured across five biological replicates of the fast and slow variants (see Supplementary table 11). We find the specific activity of $\text{DDLB}_{\text{slow}}$ is, on average, 88% ($\langle k_{\text{cat}}^{\text{slow}}/k_{\text{cat}}^{\text{fast}}\rangle$, 95% CI: [81.3%, 94.8%]) that of $\text{DDLB}_{\text{fast}}$ ($p$-value = 0.0186, one-tailed t-test for $\langle k_{\text{cat}}^{\text{slow}}/k_{\text{cat}}^{\text{fast}}\rangle < 1$). Thus, our modeling approach successfully predicts changes in activity for DDLB and CAT-III, indicating it is accurate.

**Near-native, lasso-like entangled structures are populated by the three enzymes.** Next, in our simulations we identified the structures, catalytic properties, and folding pathways that cause these activity changes. We first examined the structural distributions of CAT-III, DDLB and DHFR in the post-translational simulations by using a clustering algorithm that uses both structural information and temporal interconversion rates between metastable states. Specifically, after numerous tests of different metrics, we structurally clustered on the basis of the fraction of native contacts formed in substrate binding regions (denoted as $Q_{act}$, equation (3)) and the fraction of native contacts that exhibit non-native topological entanglements (denoted as $G$, equation (2), see Methods for description). Microstates were identified by clustering the post-translational trajectories of both synonymous variants together for each protein using the *k*-means algorithm[28], and then the microstates were coarse-grained to a smaller number of metastable states using the PCCA+ algorithm[29] (see Fig. 2a and b for CAT-III, Fig. 3a and b for DDLB and Fig. 4a and b for DHFR).

Across the metastable states, we observed diverse misfolded structures for CAT-III (see Supplementary Fig. 5) and DDLB (see Supplementary Fig. 6). Most of the misfolded structures exhibited topological entanglements, and many of these entangled structures were near native ($Q_{act} \geq 0.6$ and $G \geq 0.02$; interactive visualization of some entangled structures is provided on the supplementary website https://yuj179.github.io/topo_entanglements/). Misfolded, metastable states P9, P10, P11, P12 and P13 of CAT-III exhibited $Q_{act}$ and $G$ values of (0.63, 0.21), (0.64, 0.05), (0.79, 0.02), (0.87, 0.13), and (0.85, 0.09), respectively, while the native state (P14) had values of (0.87, 0.00) (Fig. 2h and Supplementary Fig. 5). The DDLB misfolded states P4, P5, P6, P8, P9, P10 and P11 exhibited values of (0.62, 0.13), (0.66, 0.06), (0.62, 0.09), (0.85, 0.20), (0.88, 0.08), (0.85, 0.04) and (0.96, 0.14), respectively, while the native state (P12) had values of (0.93, 0.01) (Fig. 3h and Supplementary Fig. 6). In contrast, DHFR exhibited fewer misfolded states and only one entangled, misfolded state, with $Q_{act}$ and $G$ values of (0.68, 0.05) compared to the values (0.90, 0.00) for its native state (Fig. 4h and Supplementary Fig. 7). All of the topological entanglements that we observe had a noncovalent lasso topology[30, 31, 32, 33, 34], where native contacts within a certain folded region established a closed loop along the protein backbone and another segment in the same chain threaded through this loop to become entangled (Fig. 5). None of them are topologically knotted, as identified by a knot detection algorithm[35], meaning that pulling on their termini will result in a fully extended conformation. Thus, all three enzymes sampled near-native, entangled but unknotted structures during folding.

**Some entangled structures are long-lived kinetic traps.** To disentangle a misfolded structure, it is often necessary to unfold some portion of the properly folded segments to attain the native state. This can be energetically costly. Therefore, we hypothesized that these near-native, entangled structures are long-lived kinetic traps. To test this hypothesis, we calculated the post-translational probability of being in each metastable state as a function of time using a master equation model (Supplementary Methods Section 14.1, Supplementary equation (17)) and report the results in panels c of Figs. 2, 3 and 4. For CAT-III and DDLB, entangled states (P10, P12, and P13) and (P5, P9, and P10), respectively, persisted with appreciable populations (>0.1%) until 60 s after nascent protein release from the ribosome. For DHFR, the single entangled state (P3), populated only 0.2% of the time, disentangled by 60 s. As a further test, we examined whether any post-translational trajectory of CAT-III or DDLB that reaches an entangled state ever converts to an unentangled state. For CAT-III and DDLB, we find that 83.2% and 96.6% of trajectories sample an entangled state, and of these, 87.8% and 34.1% do not convert to a state that is not entangled by the end of the simulations. Thus, we conclude that many of these entangled structures represent long-lived kinetic traps that convert to the native state over very long time scales.

**Entangled structures have altered catalytic properties.** The noncovalent lasso entanglement intermediates we observe are a form of misfolding, as they represent ordered structures that are not native. Therefore, we hypothesized that some of these entangled structures have catalytic properties different than those of the native ensemble. To test this hypothesis, we calculated the transition state barrier height of the enzyme reaction of each metastable state using a back-mapping procedure to an all-atom representation and subsequent QM/MM umbrella sampling simulations of the catalytic reaction (see the potential of mean force plots in Supplementary Fig. 8-10). Thus, for each metastable state, we obtained the activation free energy barrier $\Delta G^{\ddagger}$ going from reactants to products. For all three proteins, the native state had the lowest activation energy (panel f in Figs. 2, 3 and 4), and the other metastable states had higher barriers. Thus, these misfolded and entangled intermediates contribute to changes in specific activity. Coupled with the observation that entangled structures tend to be long-lived kinetic traps, we also conclude that these specific metastable states lead to reduced specific activity over long time scales.

**Distinctions between shallow and deep entanglements explain why CAT-III and DDLB cannot quickly fold but DHFR can.** Not all entangled structures are long-lived kinetic traps; otherwise, the entangled structure of DHFR (state P3 in Fig. 4h and Supplementary Fig. 7) would never disentangle. Sliding a small number of residues out of the closed loop (Fig. 5, also see Supplementary Methods Section 11) tends to be easier than sliding a large number of residues. In addition, unfolding a small number of residues during disentanglement also tends to be easier than unfolding a large number of residues. Therefore, we hypothesized that the minimum number of residues involved in the threading segment, whose reptation can cause disentanglement, and the minimum number of residues needed to unfold during the disentanglement process should correlate with the ability of entangled metastable states to interconvert to unentangled states. To test this hypothesis, we analyzed the representative structures from each metastable state. The entanglement of DHFR involved only, on average, 5 residues in the threading segment, and sliding these segments through the loop should not cause any portion of the protein to unfold. For CAT-III and DDLB, entangled states that never disentangle in the simulations involved, on average, 34 and 8 residues, respectively, in the threading segment and had 27 and 53 residues that need to unfold during disentanglement (see https://yuj179.github.io/topo_entanglements/). These results are consistent with our hypothesis and suggest that because DHFR's entanglement is 'shallow' (*i.e.*, involves a few residues and does not need to unfold to disentangle), thermal fluctuations can easily disentangle this structure, while thermal energy is not sufficient to quickly disentangle the CAT-III and DDLB 'deep' entanglements.

**Native-like entangled states are much less likely than other misfolded states to interact with chaperones, aggregate or be degraded and hence remain soluble.** The proteostasis machinery in cells has the potential to catalyze the folding or degradation of these entangled structures. The entangled structures could also potentially aggregate and be removed from the pool of soluble enzymes. We accounted for the effects of these processes by estimating the aggregation, degradation and Hsp70 binding propensities of the different metastable states (see Supplementary Methods Section 14.2, Supplementary equation (21)). We observed, as expected, that the less structured metastable states have a higher propensity to aggregate, be degraded or interact with chaperones. For example, relative to the native state, states P1 through P8 of CAT-III (see structures in Supplementary Fig. 5) are much more likely to experience one of these processes, as indicated by the larger magnitude of the bar plots in Fig. 2d. However, states P9 to P13 exhibit a similar propensity as the native state (P14) to aggregate, be degraded, or become chaperone substrates. Similar results are observed for DDLB (Fig. 3d) and DHFR (Fig. 4d). Thus, our model indicates that some of these entangled structures do not interact with the proteostasis machinery any more than the native state does. Therefore, the altered specific activity we observe is likely to persist inside cells over long-time scales.

We estimated the percent of proteins in each metastable state that are likely to remain soluble by accounting for the aggregation, degradation and Hsp70 binding propensities within each metastable state (see Supplementary Methods Section 14.2, Supplementary equation (19)). We find that for both CAT-III and DDLB, many of the near-native entangled structures remained soluble. At least 94% of the proteins in entangled states P10, P12 and P13 for CAT-III are estimated to remain soluble (Fig. 2e), while at least 89% of the proteins in entangled states P5, P9 and P10 of DDLB are expected to remain soluble (Fig. 3e). Thus, many of the long-lived, near-native entangled states are likely to remain soluble and free from aggregation, degradation, and catalyzed folding by chaperones.

The reason for this is that these near-native structures sequester the residues and sequence motifs that promote these processes to an extent similar to that in the native state. For example, state P12 of CAT-III exposes a similar amount of hydrophobic surface area as the native state ensemble (36.2 $nm^2$ versus 33.9 $nm^2$). The exposed surface areas of residues that promote aggregation and interactions with the chaperone Hsp70 are also similar between state P12 and the native state, respectively, 20.4 $nm^2$ versus 17.8 $nm^2$ and 39.4 $nm^2$ versus 37.3 $nm^2$.

**Entanglements persist in all-atom models.** To test that the entangled structures we observe are also kinetic traps in all-atom models, we back-mapped two entangled conformations (states P12 and P13) of CAT-III and two from DDLB (states P5 and P10) to an all-atom representation (see Supplementary Methods Section 13). We then simulated three independent trajectories of each conformation in fully solvated, unrestrained molecular dynamics at 310 K (Supplementary Methods Section 14.4). We find that all entanglements persist in each 1 μs trajectory (Supplementary Fig. 11). These results indicate that the entangled structures we have identified are likely to be long-lived kinetic traps regardless of the model resolution.

**Synonymous mutations alter the post-translational populations of entangled states.** Next, we examined how translation speed changes affect the post-translational populations of entangled structures. Using the master equation model, we observe that for CAT-III, 60 s after translation termination, 74.1% (95% CI: [71.3%, 76.9%]) of the structures are entangled when synthesis was fast, while 71.2% (95% CI: [68.1%, 74.0%]) are entangled when synthesis was slow. Likewise, the entangled population of DDLB accounted for 34.6% (95% CI: [31.64%, 37.6%]) and 29.4% (95% CI: [26.6%, 32.3%]) of the total under slow and fast translation, respectively. In contrast, the population of entangled structures for DHFR remained zero regardless of translation speed 60 s post termination. Thus, synonymous mutations can alter the population distributions of entangled states over long-time scales for deep entanglements.

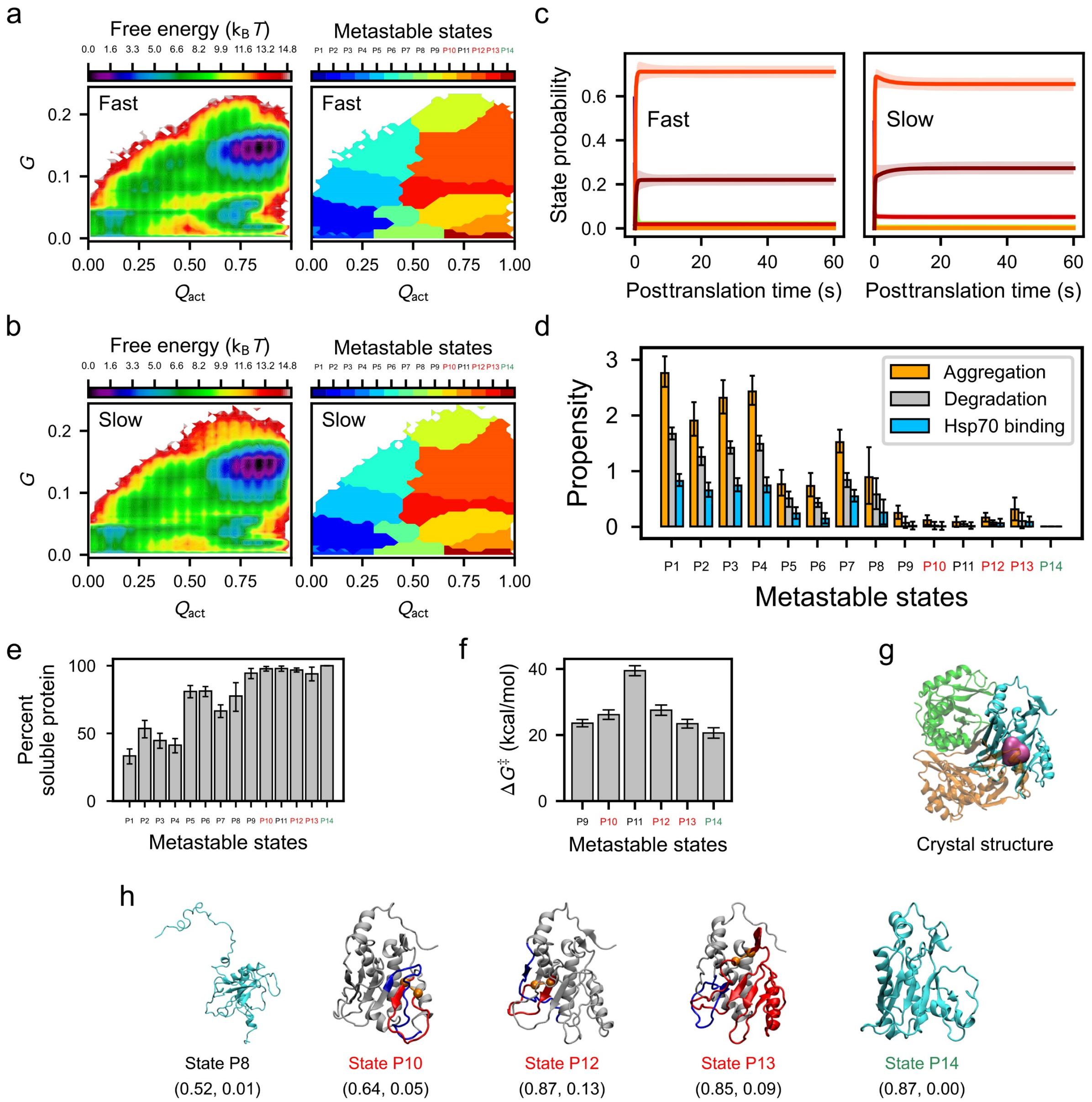

**Figure 2. Fast translation partitions more CAT-III into post-translational kinetically trapped entangled states.** a and b, Free energy surfaces for the post-translational folding of fast-translating (a) and slow-translating (b) mRNA variants as a function of the order parameters $Q_{act}$ and $G$ (left) and the regions corresponding to different metastable states (right). c, Time courses of gross state probabilities (soluble + insoluble; same colors as the metastable states in panels a and b) with error bars shown as transparent stripes for fast (left) and slow (right) variants. d, The aggregation, degradation and Hsp70 binding propensities of each metastable state calculated using Supplementary equation (21). e, The percent soluble protein of each metastable state calculated by Supplementary equation (19). f, The median transition state barrier heights ($\Delta G^{\ddagger}$) for the near-native metastable states calculated from the QM/MM simulations. g, The trimer crystal structure (3cla) with substrate shown in magenta and three monomers shown in green, orange, and cyan. h, From left to right, the representative structures of the intermediate state P8; near-native kinetically trapped states P10, P12 and P13 (the closed loop and threading segment of their entangled regions are colored in red and blue, respectively); and the native state P14. The average $Q_{act}$ and $G$ coordinates for each state are reported below the structure in the format ($Q_{act}$ value, $G$ value). For panels a through h, kinetically trapped entangled states are labeled in red; the native state (*i.e.*, state P14) is labeled in green; and others are labeled in black. All error bars represent 95% CIs calculated by

bootstrapping. The structures of P10, P12 and P13 can be explored interactively at https://yuj179.github.io/topo_entanglements/.

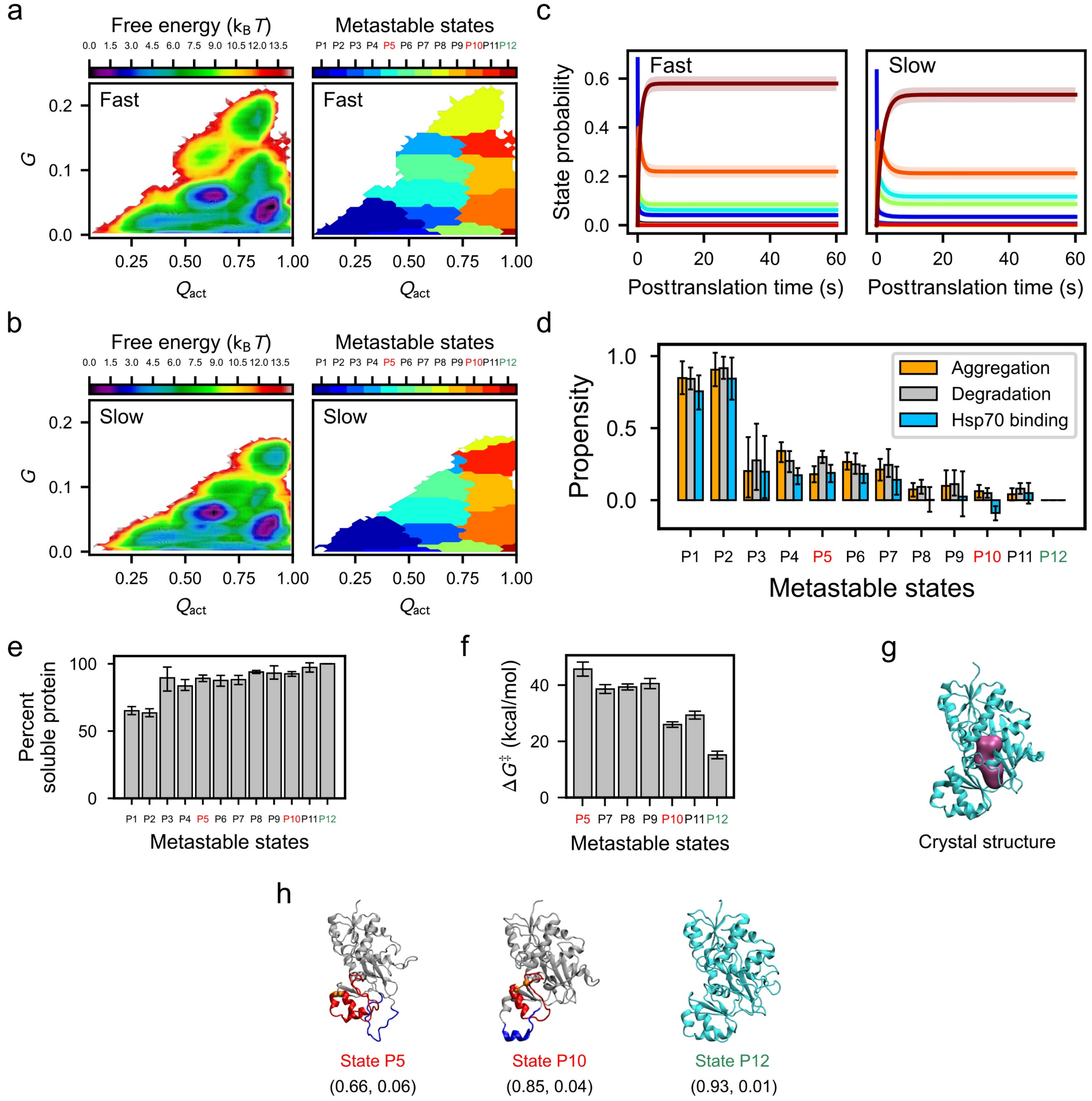

**Figure 3. Slow translation partitions more DDLB into post-translational kinetically trapped entangled states.** a and b, Free energy surfaces for the post-translational folding of fast-translating (a) and slow-translating (b) mRNA variants as a function of the order parameters $Q_{act}$ and $G$ (left) and the regions corresponding to metastable states (right). c, Time courses of gross state probabilities (soluble + insoluble; same colors as the metastable states in panels a and b) with error bars shown as transparent stripes for fast (left) and slow (right) variants. d, The aggregation, degradation and Hsp70 binding propensities of each metastable state calculated using Supplementary equation (21). e, The percent soluble protein of each metastable state calculated by Supplementary equation (19). f, The median transition state barrier heights ($\Delta G^{\ddagger}$) for the near-native metastable states calculated from the QM/MM simulations. State P4 and P6 are excluded from the $\Delta G^{\ddagger}$ estimation due to the low probability of being populated in the simulations (< 0.05%). g, Crystal structure (4c5c) with substrate shown in magenta. h, from left to right, the representative structures of the near-native kinetically trapped states P5 and P10 (the closed loop and threading segment of their entangled regions are colored in red and blue, respectively) and the native state P12. The average

$Q_{act}$ and $G$ coordinates for each state are reported below the structure in the format ($Q_{act}$ value, $G$ value). For panels a through h, kinetically trapped entangled states are labeled in red; the native state (*i.e.*, state P12) is labeled in green; and others are labeled in black. All error bars represent 95% CIs calculated by bootstrapping. The structures of P5 and P10 can be explored interactively at https://yuj179.github.io/topo_entanglements/.

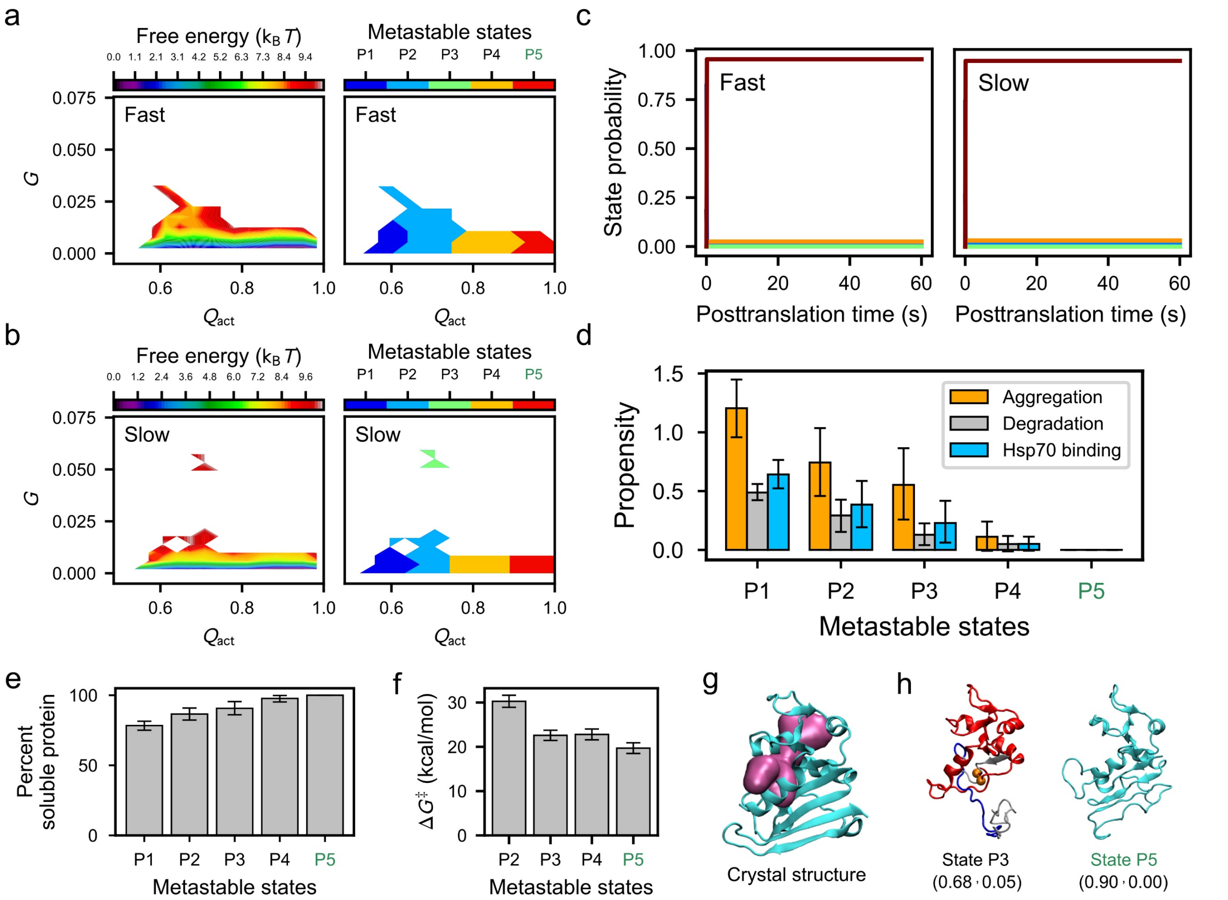

**Figure 4. No kinetically trapped states arise in synonymous variants of DHFR.** a and b, Free energy surfaces for the post-translational folding of fast (a) and slow (b) mRNA variants as a function of the order parameters $Q_{act}$ and $G$ (left) and the regions corresponding to metastable states (right). c, Time courses of gross state probabilities (soluble + insoluble; same colors as the metastable states in panels a and b) with error bars shown as transparent stripes for fast (left) and slow (right) variants. d, The aggregation, degradation and Hsp70 binding propensities of each metastable state calculated using Supplementary equation (21). e, The percent soluble protein of each metastable state calculated by Supplementary equation (19). f, The median transition state barrier heights ($\Delta G^{\ddagger}$) for the near-native metastable states calculated from the QM/MM simulations. g, Crystal structure (4kjk) with substrate shown in magenta. h, From left to right, the representative structures of transient misfolded state P3 (the closed loop and threading segment of the shallow entangled regions are colored in red and blue, respectively) and native state P5. The average $Q_{act}$ and $G$ coordinates for each state are reported below the structure in the format ($Q_{act}$ value, $G$ value). For panels a through h, kinetically trapped entangled states are labeled in red, the native state (*i.e.*, state P5) is labeled in green, and others are labeled in black. All error bars represent 95% CIs calculated by bootstrapping. The structure of P3 can be explored interactively at https://yuj179.github.io/topo_entanglements/.

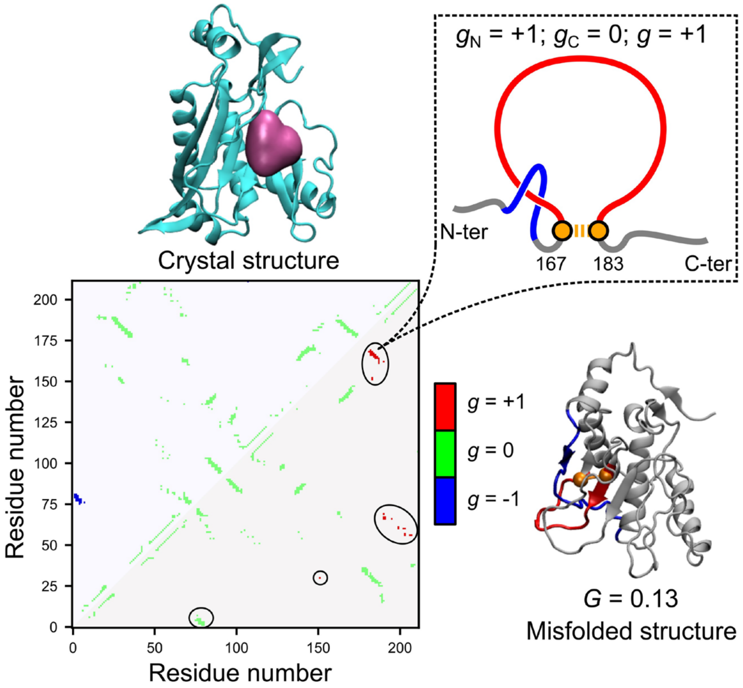

**Figure 5**. **Illustration of the *G* metric and the noncovalent lasso topology.** A contact map of the native contacts is presented on the bottom left, where the upper and lower triangular regions represent the native contacts within the native crystal structure and the misfolded structure, respectively. The native contacts are colored based on the linking number *g* calculated by Supplementary equation (16). A topology diagram of the noncovalent lasso entanglement formed by the native contact of residues 167 and 183, as an instance, is presented at the top right, where the closed loop is shown in red and the threading segment is shown in blue. As described by equation (2), the *G* metric of this misfolded structure, *G* = 0.13, was estimated by counting the number of native contacts within the misfolded structure, whose *g* value is different from that of the same native contact within the crystal structure (*i.e.*, the number of native contacts covered by the black circles on the lower triangular region of the contact map), and normalizing them to the total number of native contacts in the crystal structure.

**Synonymous mutations cause a divergence in the distribution of co-translational structures that persist post-translationally.** The post-translational changes in the distribution of entangled structures must have originated co-translationally. We hypothesized that the changes in translation speed alter the co-translational folding pathways of the protein. To test this hypothesis, we first assessed how different were the nascent chain structural distributions under fast and slow translation. To do this, we applied the normalized Jensen-Shannon divergence metric (Supplementary equation (23)) to the population of microstates identified as part of our Markov state modeling. A value of 0 means there is no difference between the distributions, while a value of 1 means the distributions are completely different. Calculating this metric at each nascent chain length during synthesis, we found that for all three proteins, the structural divergence induced by synonymous mutations was small at short nascent chain lengths and tended to increase as the nascent chain length increased (Fig. 6b, e and h). The maximum divergence occurred at or near the longest nascent chain length before the nascent chain was released from the ribosome. Specifically, for DHFR, the structural ensembles arising from fast and slow synthesis started to diverge at approximately 110 residues in length, while for CAT-III and DDLB, the two distributions start to diverge at 190 and 210 residues, respectively. Thus, the fast- and slow-translation rates altered the co-translational distributions of conformations for all three enzymes.

Applying this divergence metric to the time-dependent post-translational structural distributions, we observed that for DHFR, the initially different structural ensembles arising from fast and slow synthesis rapidly converged to nearly the same distribution post-translationally (*i.e.*, have divergence values very close to zero in Fig. 6h). For CAT-III and DDLB, the two distributions do not reconverge 60 s after synthesis. Thus, the changes in conformations caused by synonymous mutations are quickly 'forgotten' by DHFR due to its rapid equilibration, while CAT-III and DDLB retain a 'memory' of those co-translational differences due to kinetic trapping.

**Changes in co-translational folding pathways give rise to altered post-translational populations.** Next, we characterized the co- and post-translational folding pathways arising from changes in the translation rate. To do this, we calculated the populations and pathway probabilities of transitions between metastable states occurring co- and post-translationally under the different translation schedules (see Supplementary Methods Section 16). To display similarities and differences in folding pathways, we used arrows with different colors. Black arrows between metastable states indicate that a transition was observed between those states in the top 80% of populated pathways, blue arrows indicate that the transition was seen only in the slow-translation schedule, and orange arrows indicate that the transition was seen only in the fast translation schedule.

We identified nine co- and five post-translational metastable states (representative structures shown in Fig. 6g) for DHFR. The co- and post-translational folding pathways were very similar, as most of the transitions were seen in both translation schedules (black arrows in Fig. 6g). There were some differences though. Transitions from state C7 to C9 and from C9 to P1 were observed for only fast and slow schedules, respectively. The pathway probabilities, however, demonstrated that even though the initial post-translational structural distribution differed, nearly all states converted to the native state by the end of the post-translational simulations (Fig. 6i). For example, the pathway probabilities of C9→P5 are 100% for both fast and slow translation. The pathway probability of C8→P5 is 100% for fast translation and 99.8% for slow translation. An exception is the transition probability of C6→P5, where state C6 was not sampled by the slow variant, and no such transition was observed in our simulation. However, due to the small population of C6 (2% in Fig. 6i) sampled by the fast variant, this difference was negligible. Thus, the co-translational folding of DHFR was slightly affected by differences in translation speed; however, these differences quickly disappeared due to rapid folding from all post-translational metastable states.

In contrast, CAT-III exhibited a co- and post-translational folding reaction network (Fig. 6a) and pathway probabilities that were sensitive to translation speed changes and whose resulting differences persisted post-translationally. Nine co-translational and fourteen post-translational metastable states were identified for CAT-III (Fig. 6a). Six of these post-translational metastable states (P6, P9, P10, P11, P12 and P13) exhibited entangled structures, while no co-translational states exhibited entanglement. Thus, entanglement was a post-translational process for CAT-III (see Supplementary videos I and II visualizing, respectively, the process of entanglement versus correct folding). Very different co-translational folding pathways were observed in the top 80% of pathways starting at a nascent chain length of approximately 195 residues, where the divergence metric exhibited a large increase (Fig. 6b), and transitions into and out of co-translational metastable state C9 started to differ between the fast and slow translation schedules (as indicated by the blue and orange arrows in Fig. 6a). For example, only in the slow schedule can C9 transition to P13, while only in the fast schedule can C5 transition to P1. These differences in allowed transitions persisted post-translationally as well, with a number of states, such as P12 and P13, being effective sinks during the post-translational simulations, *i.e.*, allowing no direct or indirect transitions to the native state (P14) once they were reached. These sinks are entangled structures, which is consistent with our earlier observation that these entangled states tend to be kinetic traps (indicated by the red borders around metastable states P12 and P13 in Fig. 6a).

Thus, the translation speed causes differences in the CAT-III co-translational folding pathways once at least 195 residues have been synthesized, and those differences lead to changes in the populations of post-translationally entangled states, thereby altering the transition state barrier for the reaction this enzyme carries out (Fig. 2f) and affecting the specific activity of CAT-III over long time scales (Fig. 1g).

Finer-grained consideration of the folding pathways (estimated by using the master equation model) shows that post-translationally, CAT-III$_{fast}$ partitioned 5.7% (95% CI [1.7%, 9.9%]) more protein (gross amount, soluble + insoluble, same hereafter) into the near-native kinetic trap P12, while CAT-III$_{slow}$ partitioned 5.2% (95% CI [1.4%, 9.2%]) more protein into native state P14. The transitions towards the native state P14 were enhanced for CAT-III$_{slow}$, whereas the transitions towards the kinetic trap P12 were enhanced for CAT-III$_{fast}$. This is also indicated by the pathway probabilities shown in Fig. 6c. Synonymous mutations also led to smaller changes in other states, including the kinetic trap P13 and intermediate state P8.

Similar to CAT-III, DDLB also exhibits co- and post-translational folding pathways that were sensitive to changes in translation speed across its eight co- and twelve post-translational metastable states. However, unlike CAT-III, DDLB co-translationally formed entangled structures (states C8 and C9) that persisted to form entangled post-translational states (states P1, P3 and P5). In CAT-III, the divergence in co-translational folding pathways between fast and slow translation schedules monotonically increased with increasing chain length (Fig. 6b). However, the co-translational folding pathways of DDLB diverged starting at nascent chain length 110 but reconverged around 190 residues, diverging again starting at 210 residues (Fig. 6e). The first divergence involved a transition to state C4 with the fast schedule that did not occur with the slow schedule (orange arrow in Fig. 6d). This state ultimately interconverted to state C3, which is an obligatory intermediate in both schedules. The second divergence started at state C7, in which 6% more nascent chains went to entangled state C8 with the slow schedule than with the fast schedule, and the rest went to state C10. Parallel co- and post-translational folding pathways then arose. In one pathway, transitions were observed from C10→C11→P2, while the other transitions from C8→C9→P1 involved entangled structures in all states (see Fig. 6d, also see Supplementary videos III and IV illustrating the processes of entanglement and folding). The increased probability of this entangled pathway during slow translation decreased the probability that DDLB would post-translationally reach the native state P12 (3.1% versus 9.3% in slow versus fast, Fig. 6f), while DDLB$_{slow}$ partitioned 5.5% (95% CI [3.1%, 7.8%]) more protein (gross amount, soluble + insoluble) into the near-native kinetic trap P5 after post-translation (as estimated by using the master equation model). Thus, we again see that entangled states act as sinks with altered transition state barriers for the enzyme's reaction (Fig. 3f). This population shifts among these states and the folding pathways they take part in leads to long-lived changes in specific activity.

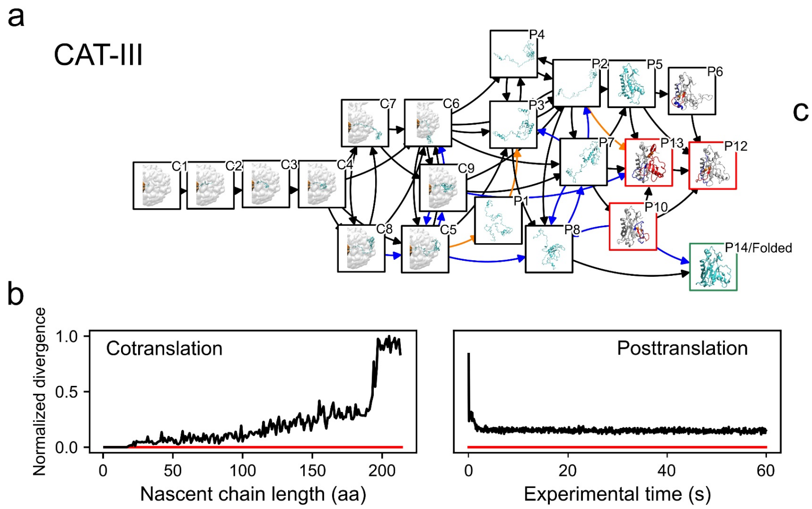

c

| Pathway probability | | |
|---|---|---|
| | Fast | Slow |
| * → C9 | 7.0% | 24.0% |
| C9 → P14/Folded | 14.3% | 12.9% |
| * → C6 | 63.0% | 53.0% |
| C6 → P14/Folded | 20.5% | 30.0% |
| * → C5 | 30.0% | 23.0% |
| C5 → P14/Folded | 28.3% | 35.7% |

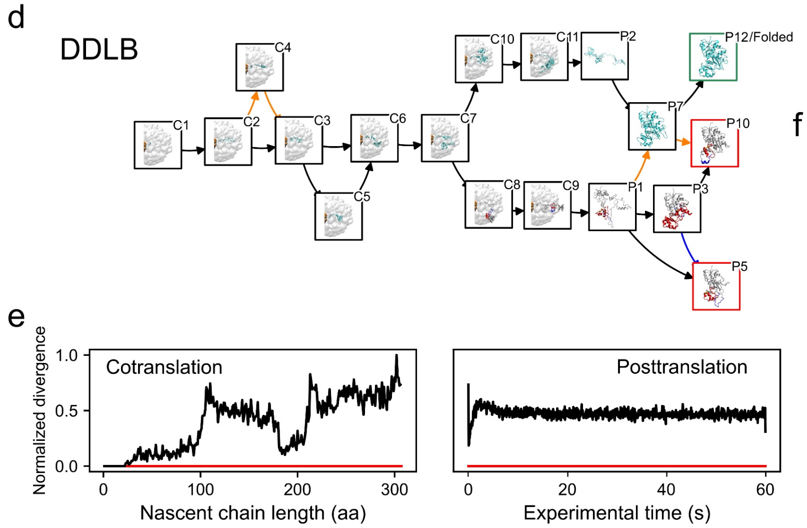

f

| Pathway probability | | |
|---|---|---|
| | Fast | Slow |
| * → C11 | 70.0% | 64.0% |
| C11 → P12/Folded | 79.6% | 81.1% |
| * → C9 | 30.0% | 36.0% |
| C9 → P12/Folded | 9.3% | 3.1% |

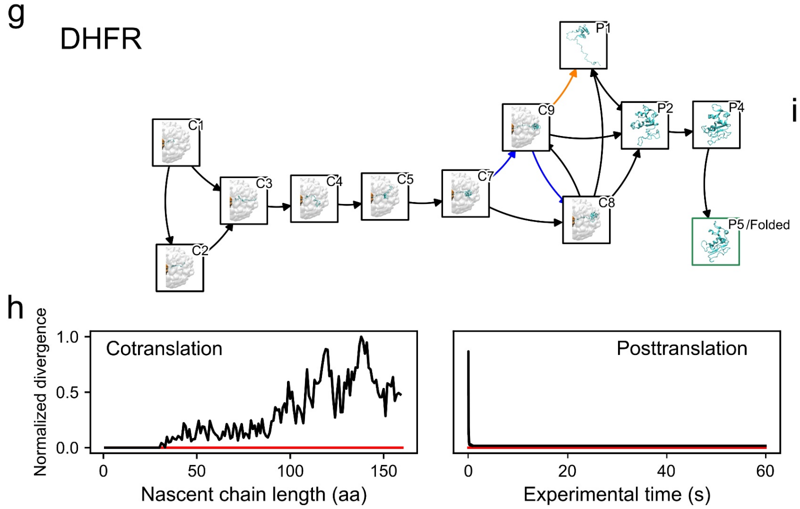

i

| Pathway probability | | |
|---|---|---|
| | Fast | Slow |
| * → C9 | 10.0% | 14.0% |
| C9 → P5/Folded | 100.0% | 100.0% |
| * → C8 | 88.0% | 86.0% |
| C8 → P5/Folded | 100.0% | 99.8% |
| * → C6 | 2.0% | 0.0% |
| C6 → P5/Folded | 100.0% | NaN |

**Figure 6. Co- and post-translational folding pathways of CAT-III, DDLB and DHFR.** a, d, and g, The most probable folding pathways are presented as a directed graph where the nodes represent metastable states, with representative structures shown in the boxes, and the edges represent the transitions in the 80% most likely pathways. The nodes corresponding to the native state and the kinetically trapped states

are framed in green and red, respectively. The transitions (edges) that are observed in only the fast variants or the slow variants are marked in orange and blue, respectively. b, e and h, Normalized Jensen-Shannon divergence (Supplementary equation (23)) of the co- and post-translational microstate distributions comparing fast and slow variants, with a red line indicating zero divergence. c, f and i, The co-translational pathway probabilities (* → end state C*) and the post-translational pathway probabilities from the co-translational end states to the native state (C* → native state) are summarized in the tables. The pathway probabilities whose changes were greater than 5% are highlighted in yellow. The post-translational pathway probabilities are normalized by the corresponding co-translational pathway probabilities that involve the particular co-translational end state.

#### Discussion

How synonymous mutations alter the specific activity of enzymes has been an unanswered question. A detailed understanding of the molecular mechanisms giving rise to this phenomenon would provide biochemists, molecular biologists, evolutionary biologists, and biomedical researchers a framework to interpret experiments on the influence of codon usage on protein structure, function, cellular phenotype, disease, and the selection pressures shaping mRNA sequence evolution. In this study, we developed a novel multiscale model that recapitulates the experimental observations for the specific activity of CAT-III variants[1] and correctly predicts the changes in specific activity of DDLB variants, giving us confidence that the model is realistic. This provided us the opportunity to study the structures, pathways, and kinetics that give rise to this phenomenon.

Our key findings are that (1) changes in elongation kinetics induced by synonymous mutations can alter co-translational nascent chain structural ensembles and folding pathways; (2) for some enzymes, such as CAT-III and DDLB, these changes in the structural ensemble can persist long after the nascent chain has been released from the ribosome; (3) this persistence arises from conformational states that are long-lived kinetic traps; (4) these kinetic traps arise at the molecular level from deep entanglements that slowly disentangle because they require the unfolding of already folded protein segments; (5) these entanglements are non-covalent lasso topologies in which a closed loop is formed by a backbone segment connecting two residues that form a non-covalent native contact, and another segment threads through this loop; (6) many of these entangled structures have decreased catalytic efficiencies due to structural perturbation of their active sites; (7) some entangled structures are very similar to the native state – exposing similar extents of hydrophobic surface area and chaperone binding motifs; and (8) because of their near-native conformations, these entangled structures can bypass the chaperone and degradation machinery of the cell and do not exhibit an increased propensity to aggregate. This situation results in a soluble fraction of enzymes with decreased specific activities that can persist for long time periods in cells.

Two concepts central to our explanation – intra-molecular entanglement and subpopulations of kinetically trapped states – are not without precedent. The material properties of entangled synthetic polymers have long been studied and modeled[36, 37] in the field of polymer physics. And there has been a large amount of research focused on knotted proteins that often contain disulfide bonds or topologies in which when the protein's ends are pulled in opposite directions the protein does not fully extend[31, 38, 39, 40]. These characteristics, however, are not present in our entanglements. The entanglements we observe form a closed loop due to a non-covalent native contact, and if both ends of the protein were pulled disentanglement would occur. Recently, this type of entanglement has been detected in almost one third of the protein structures deposited in the Protein Data Bank[32]. Thus, the potential for protein segments to form non-native, noncovalent lassos is plausible. Kinetically trapped states of proteins have been observed in single-molecule experiments probing the functioning of flavoenzymes[41, 42]. In one study, heterogeneous populations of the protein cholesterol oxidase were observed to stochastically switch very slowly between active and nonactive states[41]. However, in that study, the influence of protein synthesis on the distribution of conformational states was not probed. Thus, a novel aspect of our discovery is that it combines the phenomena of entanglement and kinetic trapping as essential to the mechanism by which synonymous mutations affect co- and post-translational protein structure and function.

We also observed in our simulations the counterintuitive phenomenon that the folding of some proteins is promoted when their rapid synthesis gives them less time to fold on the ribosome. This phenomenon was first predicted from chemical kinetic models[43] of co-translational folding. Specifically, we observed that synthesizing DDLB three times faster increased the final post-translational folding probability of DDLB by 4.6% (95% CI [0.4%, 8.9%]) and increased its activity. Consistent with the earlier prediction[43], this phenomenon arose because having less time to fold on the ribosome decreases DDLB partitioning into the misfolded, kinetically trapped state. This adds to the growing list of proteins for which slowing synthesis decreases folding efficiency, which results in altered function, and challenges the common rule of thumb that slow-translating codons promote folding.

An important aspect of our choice of protein systems is that we included a protein, DHFR, whose activity is not sensitive to synonymous mutations. This provided us with the opportunity to gain insights by comparing and contrasting its behavior with those of CAT-III and DDLB. A key insight from this comparison is that even though the DHFR co-translational structures and folding pathways were sensitive to synonymous mutations, these differences did not persist post-translationally because the only entangled state that DHFR populates and disentangled rapidly. This is consistent with previous experimental and theoretical studies that show that DHFR has fast folding kinetics and an absence of significant off-pathway intermediates[44] and kinetic traps[45]. We demonstrated that the two reasons for this are that DHFR only negligibly populates an entangled conformation and that the entanglement it forms is 'shallow' - its disentanglement requires only 5 residues to slide out of the closed loop and does not need to unfold any other portions of the protein to do so. Thus, this shallow entanglement can be easily disentangled without large structural rearrangements. In contrast, CAT-III and DDLB entanglements have many more residues involved, requiring large structural rearrangements to become disentangled, leading to more-persistent entangled structures. Thus, the presence of entanglement is not sufficient to guarantee a kinetic trap. What is important is the nature of the entanglement and the native structure surrounding it.

Like any simulation study, limitations of force fields, time scales, and sampling statistics are of concern. In the CG simulations, we used a structure-based force field to encode the native state as the global free energy minimum. One of the many benefits of this is that we automatically know that the entangled structures we observe are metastable states, as the native state is encoded to be the global free energy minimum below the protein's melting temperature. However, one potential drawback is that this force field does not allow ordered, nonnative secondary and tertiary structures to form (*e.g.*, the model does not allow β-sheets to switch to α-helical bundles). This means that additional structures beyond entanglement could also contribute to the influence of synonymous mutations on enzyme activity. At the monomeric protein level, however, such nonnative ordered structure seems unlikely. Except for fold-switching proteins, which are rare in nature (only 0.5 to 4% of proteins in the Protein Data Bank are estimated to switch folds[46]) and usually require quaternary interactions to induce structural changes[47], there have been no reported experimental data for alternative, ordered tertiary structures occurring on the folding pathways of monomeric proteins in bulk solution. In the QM/MM simulations, we used the 3rd-order density-functional tight-binding (DFTB) method[48] as a balance between computational speed and accuracy. Transition state barrier heights can change depending on the level of theory and basis set used. However, it is often the case that trends across related compounds or conformations are robust. Therefore, the differences in the barrier heights between metastable states are likely to be qualitatively correct, meaning our overall conclusions would remain unchanged even if a higher level of theory was used.

Another limitation of our simulations is that we only simulated one protein chain at a time. Thus, we can never observe homodimer-swapped structures that could also be long-lived kinetic traps and influence function[49]. However, as noted, in the experimental examples provided in the Introduction, such dimer-swapped structures were ruled out. To address sampling issues, we applied rigorous statistical analyses of all our reported quantities, including confidence intervals and p-values. All our conclusions were drawn from statistically significant signals, indicating that sampling was not an issue. For these reasons, our results are robust and statistically meaningful, giving us confidence that they represent realistic scenarios of what is happening at the molecular level.

The predictions that monomeric enzymes can become intramolecularly entangled and populate long-lived states can be tested experimentally. Ensemble-level experiments, such as NMR and X-ray crystallography, often need appreciable populations of a state to detect that state, which might make entanglement detection difficult. Single-molecule techniques, such as FRET, can detect heterogeneous populations but contain little structural information since the signal arises from only the two points where dyes are attached to a protein. Therefore, cryo-EM, with its ability to build structural clusters of heterogeneous populations, and chemical-exchange mass spectrometry, which can detect time-dependent distributions of chemical modifications on proteins, seem the most promising techniques to experimentally detect these entanglements.

This study explains how synonymous mutations can alter enzyme activity in cells. Synonymous mutations alter translation elongation speeds and change the population of nascent chain conformations in entangled states that are near native but have lower catalytic efficiencies than that of the native state. Hence, the specific activity, a quantity averaged over the populations of proteins in different conformational states, can increase or decrease due to synonymous mutations. The experimental search for these entangled structures and their roles in influencing protein structure, function, and phenotypes in cells is likely to be a fruitful area of research in the future.

**Acknowledgements**

E.P.O. acknowledges support from the National Science Foundation (MCB-1553291) as well as the National Institutes of Health (R35-GM124818). S.J.B. acknowledges support from the NIH (GM-122595), the Eberly Family Distinguished Chair in Science, and the Howard Hughes Medical Institute. Computations in this work have been carried out on the Extreme Science and Engineering Discovery Environment (XSEDE) supercomputer, which is supported by MCB-160069, and the Pennsylvania State University's Institute for Computational and Data Sciences' Roar supercomputer.

**Author contributions**

E.P.O. designed the research; Y.J. developed the computational methods with contributions from E.P.O.; Y.J. wrote the computer code and carried out the simulations and computations; S.S.N. and S.J.B. designed the experimental validation for DDLB variants; S.S.N. and P.P. performed the experiments; All of the authors analyzed the data and wrote the manuscript.

**Code availability**

All computer code developed in this work is available in the GitLab repositories https://git.psu.edu/yuj179/cg_simtk_protain_folding and https://git.psu.edu/yuj179/activation-energy-estimation-workflow, under the MIT license.

#### Graphical Abstract

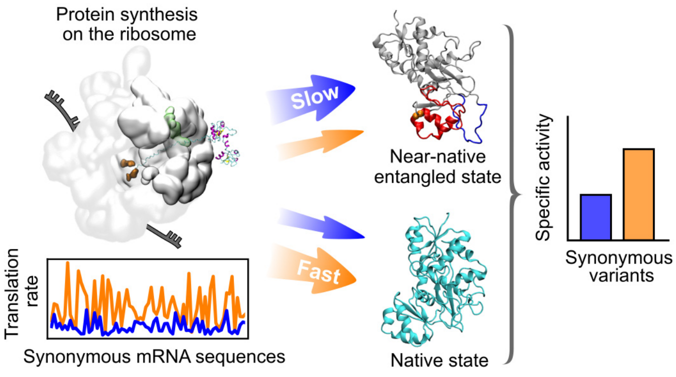

Supplementary information for

### How synonymous mutations alter enzyme structure and function over long time scales

Yang Jiang[1], Syam Sundar Neti[1], Priya Pradhan[1], Squire J. Booker[1, 2, 3] and Edward P. O'Brien[1, 4, 5, *]

[1] Department of Chemistry, Penn State University, University Park, Pennsylvania, United States

[2] Department of Biochemistry and Molecular Biology, Pennsylvania State University, University Park, Pennsylvania, United States

[3] Howard Hughes Medical Institute, Pennsylvania State University, University Park, Pennsylvania, United States

[4] Bioinformatics and Genomics Graduate Program, The Huck Institutes of the Life Sciences, Penn State University, University Park, Pennsylvania, United States

[5] Institute for Computational and Data Sciences, Penn State University, University Park, Pennsylvania, United States

#### Supplementary Methods

##### 1. Coarse-grained (CG) protein model

We utilized a Gō-based force field[1, 2, 3, 4, 5] for all the proteins investigated in this study. Briefly, this CG model represents each residue as a single interaction site centered on the Cα atom position. The potential energy function is:

$$\begin{aligned} E_{\text{tot}} &= E_{\text{bond}} + E_{\text{angle}} + E_{\text{dihedral}} + E_{\text{elec}} + E_{\text{vdW}} \\ &= \sum_i K_{\text{b}}(b_i - b_0)^2 + \sum_i -\frac{1}{\gamma}\ln\left\{e^{-\gamma[K_\alpha(\theta_i-\theta_\alpha)^2+\varepsilon_\alpha]} + e^{-\gamma K_\beta(\theta_i-\theta_\beta)^2}\right\} \\ &\quad + \sum_i \sum_{j=1}^{4} K_{\text{D}_j}\left[1+\cos\left(j\varphi_i - \delta_j\right)\right] + \sum_{i,j} \frac{q_i q_j}{4\pi\varepsilon_0\varepsilon_\gamma r_{ij}} \cdot e^{-\frac{r_{ij}}{l_{\text{D}}}} \\ &\quad + \sum_{i,j} \varepsilon_{ij}\left[13\left(\frac{R_{ij}}{r_{ij}}\right)^{12} - 18\left(\frac{R_{ij}}{r_{ij}}\right)^{10} + 4\left(\frac{R_{ij}}{r_{ij}}\right)^{6}\right], \end{aligned} \quad (1)$$

where $K_\text{b}$ is the bond force constant and equals 50 kcal/mol/Å$^2$; $b_i$ is the $i^{\text{th}}$ pseudo bond length between two adjacent interaction sites; $b_0$ is the equilibrium pseudo bond length and is set to 3.81 Å, which is the average distance between two adjacent Cα atoms in the protein sequence; $\gamma$, $K_\alpha$, $\theta_\alpha$, $\varepsilon_\alpha$, $K_\beta$ and $\theta_\beta$ are all constants of the double-well angle potential[6], which was designed to describe bond angles associated with both α-helix and β-sheet conformations; $\theta_i$ is the $i^{\text{th}}$ angle of two adjacent pseudo bonds; $K_{\text{D}_j}$ and $\delta_j$ are the dihedral force constant and the phase at periodicity *j*, respectively; $\varphi_i$ is the $i^{\text{th}}$ pseudo dihedral angle; $q_i$ is the net charge of the $i^{\text{th}}$ interaction site, which equals the net charge of the corresponding amino acid residue; $\varepsilon_0$ and $\varepsilon_\gamma$ are the dielectric constants of vacuum and water, respectively; $l_\text{D}$ is the Debye length and is set[1] to 10 Å; $\varepsilon_{ij}$ and $R_{ij}$ are the well depth and the vdW radius in the LJ 12-10-6 potential[7], which takes into account of desolvation barriers between interaction sites *i* and *j* and $r_{ij}$ is their distance. The 1-4 nonbonding interactions were included in our model without scaling any parameters. The nonbonding interactions were smoothly switched to zero starting at a distance of 18 Å and ending at 20 Å. Constraints on the bond length were applied in all the simulations.

Parameters $\varepsilon_{ij}$ and $R_{ij}$ were set according to the interaction types. For the interaction sites that have native contacts, $\varepsilon_{ij} = \varepsilon_{ij}^{\text{HB}} + n_{\text{scal}} \cdot \varepsilon_{ij}^{\text{SC-SC}} + \varepsilon_{ij}^{\text{BB-SC}}$, where the hydrogen bond (HB) contact well depth $\varepsilon_{ij}^{\text{HB}}$ was set as 0.75 kcal/mol for a single HB contact and 1.5 kcal/mol for multiple HB contacts; the sidechain-sidechain (SC-SC) interaction well depth $\varepsilon_{ij}^{\text{SC-SC}}$ was set according to the Betancourt−Thirumalai statistical potential[8] and then scaled by a multiplicative factor $n_{\text{scal}}$ to achieve a realistic native-state stability for a particular protein; the backbone-sidechain (BB-SC) interaction well depth $\varepsilon_{ij}^{\text{BB-SC}}$ was set as 0.37 kcal/mol. All the native contacts were identified for the residue pairs that are separated by no less than 2 residues within the native structure. The native HB contacts were identified by STRIDE[9]. The native SC-SC contacts and BB-SC contacts were identified as those residue pairs that have any heavy atom within 4.5 Å in the crystal structure. The value of $R_{ij}$ for a native contact was set as the native distance $d_{ij}$ between interaction sites *i* and *j*. For the non-native contacts, to make the interaction mostly repulsive, we used $\varepsilon_{ij} = \sqrt{\varepsilon_i \cdot \varepsilon_j}$ and $R_{ij} = R_i + R_j$, where $\varepsilon_i$ was set as 0.000132 kcal/mol and $R_i$ was set as the non-native collision diameter $\sigma_i$ of interaction sites *i* multiplied by $2^{1/6}$ and divided by 2. The collision diameters were identified based on the protein's native structure according to Karanicolas−Brooks method[7]. All the CG force field parameters are summarized in Supplementary Table 1. All the CG simulations in this study were performed in OpenMM[10].

#### 2. Defining native-state stabilities by parameterizing $n_{\text{scal}}$

Setting a realistic free energy of stability is important for accurately modeling protein folding in the CG model. However, due to the time-consuming parameterization process, especially for a large multi-domain protein, it is impractical to parameterize all the proteins that we are interested in. We therefore utilized a training set that contains 18 small single-domain proteins to optimize each $n_{\text{scal}}$ to reproduce the corresponding experimental protein stability[5, 11, 12, 13, 14] ($\Delta G_{\text{UN}}^{\text{exp}}$). The training-set proteins can be classified into 3 structural classes: mainly α-helix (marked as α class, see Supplementary Table 2a), mainly β-sheet (marked as β class, see Supplementary Table 2b) and α/β fold (marked as α/β class, see Supplementary Table 2c). We used the resulting $n_{\text{scal}}$ data of those structural classes to further parameterize the proteins that we are interested in.

Each training-set protein was coarse-grained using the interaction sites at every Cα position within its crystal structure. The CG structures were then parameterized using 4 different $n_{\text{scal}}$ values, resulting 4 CG models for each training-set protein. We utilized parallel temperature replica exchange molecular dynamic (pt-REMD)[15] simulations to efficiently sample the conformational space of each CG model. 100,000 replica exchange attempts were performed with the sampling time 75 ps between two attempts, resulting the total sampling time as 7.5 μs, for each CG model. The temperature windows for each pt-REMD simulations were chosen within the range of 280 K to 400 K and progressively optimized so that the successful exchange ratios are no less than 0.6 by using the minimum number of windows. The heat capacity $C_{\mathrm{v}}$ and the probability of protein folding $P_{\mathrm{N}}$ were then estimated using the Weighted Histogram Analysis Method[16] (WHAM). The melting temperature $T_{\mathrm{m}}$ was identified as the temperature where the $C_{\mathrm{v}}$ of the particular protein reaches its maxima. We used the fraction of native contact *Q* within secondary structure elements as the order parameter (calculated using the main text equation (3), where *I* and *J* are both the set of residues within secondary structure elements) and the threshold $Q_{\mathrm{eq}}$ used to identify the folded and unfolded states was chosen as the Q value where the cumulative probability reaches 0.5 at $T_{\mathrm{m}}$. The folded states were then identified as the structure whose $Q > Q_{\mathrm{eq}}$. The protein stability $\Delta G_{\mathrm{UN}}^{\mathrm{sim}}$ at the experimental temperature $T^{\mathrm{exp}}$ was then estimated by

$$\Delta G_{\mathrm{UN}}^{\mathrm{sim}}(T^{\mathrm{exp}}) = -k_{\mathrm{B}}T^{\mathrm{exp}} \cdot \ln\left[\frac{P_{\mathrm{N}}(T^{\mathrm{exp}})}{1 - P_{\mathrm{N}}(T^{\mathrm{exp}})}\right], \tag{2}$$

where $k_{\mathrm{B}}$ is Boltzmann's constant.

There are several proteins in α and α/β classes whose conformational space cannot be sufficiently sampled within the 100,000 exchange attempts, i.e., replicas cannot cross the free energy barrier to visit both of the folded and unfolded states frequently. To improve the reliability of the estimated protein stability, we performed an extra pt-REMD simulation at each $n_{\mathrm{scal}}$ value starting from the unfolded states that were obtained from the MD simulations at 800 K. The final $\Delta G_{\mathrm{UN}}^{\mathrm{sim}}$ at a particular $n_{\mathrm{scal}}$ value for those insufficiently sampled proteins (shown in Supplementary Table 3) were calculated as the average of the protein stability values estimated using the pt-REMD simulations starting from their native states and unfolded states.

The optimized $n_{\mathrm{scal}}$ for each training-set protein (shown in Supplementary Table 2) was determined through extrapolation as

$$n_{\mathrm{scal}}^{*} = \frac{\Delta G_{\mathrm{UN}}^{\mathrm{exp}} - b}{m}, \tag{3}$$

where *m* and *b* are the slope and intercept obtained by the linear regression of $\Delta G_{\mathrm{UN}}^{\mathrm{sim}}$ as a function of $n_{\mathrm{scal}}$, respectively.

##### 3. Stepwise $n_{\mathrm{scal}}$ optimization strategy for arbitrary proteins

To further parameterize the proteins that are more complex than the training set proteins, we used a stepwise optimization strategy. Five levels of $n_{\mathrm{scal}}$ for each structural class were predefined as shown in Supplementary Table 5. At the first level, $n_{\mathrm{scal}}$ was chosen as the mean value of the structural class $\langle n^*_{\text{scal}}\rangle_{\text{class}}$, as well as the overall mean value of the 18 training-set proteins for the domain interface. At the second level, $n_{\text{scal}}$ was chosen as $\langle n^*_{\text{scal}}\rangle_{\text{class}}$ increased by $\langle \Delta\Delta G/\Delta G^{\text{class}}_{\text{UN}}\rangle_{-} \times 100\% = 23\%$, where $\Delta\Delta G = \Delta G^{\text{exp}}_{\text{UN}} - \Delta\Delta G^{\text{class}}_{\text{UN}}$, $\Delta\Delta G^{\text{class}}_{\text{UN}} = m \cdot \langle n_{\text{scal}}\rangle_{\text{class}} + b$ and $\langle \dots \rangle_{-}$ means the average is only calculated for the proteins that are destabilized using $\langle n^*_{\text{scal}}\rangle_{\text{class}}$ within the training set. At the third level, $n_{\text{scal}}$ was chosen as $\langle n^*_{\text{scal}}\rangle_{\text{class}}$ increased by $\max_{-}\{\Delta\Delta G/\Delta G^{\text{class}}_{\text{UN}}\} \times 100\% = 46\%$, where $\max_{-}\{\dots\}$ means the maximum value only accounts for the proteins that are destabilized using $\langle n^*_{\text{scal}}\rangle_{\text{class}}$ within the training set. At the fourth level, $n_{\text{scal}}$ was chosen as $\langle n_{\text{scal}}\rangle_{\text{class}}$ increased by $\min\{\Delta\Delta G\}/\langle \Delta G^{\text{class}}_{\text{UN}}\rangle \times 100\% = 70\%$, which is the largest destabilization relative to the overall average $\Delta G^{\text{class}}_{\text{UN}}$. At the last level, $n_{\text{scal}}$ was chosen as the overall maximum value of all levels.

An arbitrary protein was parameterized using the following strategy: The protein was coarsen-grained using our CG force field (Supplementary equation (1)). The native vdW interactions were divided into groups based on the domains defined by CATH[17]; $n_{\text{scal}}$ was assigned as the first-level value according to the domain's structural class (or interface). We then performed 10 parallel 1-μs MD simulations at 310 K for this CG model and monitor the *Q* value of each domain and interface. The domain or interface was regarded stabilized when all the 10 trajectories have the *Q* values greater than the threshold $\langle Q_{\text{kin}}\rangle$ for no less than 98% of the simulation time. We then increased the $n_{\text{scal}}$ values of the destabilized domains and interfaces to the next level and keep the other's $n_{\text{scal}}$ values invariant and generate the next CG model. We repeated the above procedure until all the domains and interfaces were stabilized. If a domain or interface cannot be stabilized even using the highest level of $n_{\text{scal}}$, the median $n_{\text{scal}}$ value (i.e., level 3) of the corresponding structural class was used to generate the final CG model regardless of the stability.

#### 4. Estimation of *in silico* protein folding times

The *in silico* protein folding time $\langle \tau^{\text{sim}}_{\text{F}}\rangle$ was estimated as the mean first passage time (MFPT) of protein folding for each training-set protein by using temperature quenching simulations. We performed MD simulations for all the CG protein models built using corresponding $n^*_{\text{scal}}$ at 800 K to force them unfolded, followed by running MD simulations at 310 K to monitor protein folding. The trajectory where the *Q* value is higher than a threshold $Q_{\text{kin}}$ for at least 150 ps was identified as a folded trajectory and then terminated. $Q_{\text{kin}}$ is defined as $Q_{\text{kin}} = \frac{1}{2}(Q_{\text{eq}} + Q_{310})$, where $Q_{310}$ is the most probable *Q* value at 310 K for each training-set protein estimated by using WHAM in the pt-REMD trial whose $n^*_{\text{scal}}$ is closest to the optimized value $n^*_{\text{scal}}$.

500 trajectories were run for each training-set protein and the proportion of unfolded trajectories ($S_{\mathrm{U}}$) was fit to an exponential decay as a function of simulation time. We first considered the simplest folding kinetics with one pathway $\mathrm{U} \xrightarrow{k_{\mathrm{F}}} \mathrm{F}$. In this case, $S_{\mathrm{U}}$ follows a single-exponential equation

$$S_{\mathrm{U}}(t) = \begin{cases} 1, & 0 \le t < t_0 \\ e^{-k_{\mathrm{F}}(t-t_0)}, & t \ge t_0 \end{cases}, \tag{4}$$

where $t_0$ is the delay time of protein folding. The single-exponential fitting works well except for protein ABP1, SH3, and Tenascin, whose fitting quality $R^2$ are both lower than 0.99. We further considered the protein folding kinetics that has two parallel pathways from U to F with folding rates $k_1$ and $k_2$ for each. In this case, $S_{\mathrm{U}}$ follows the double-exponential equation

$$S_{\mathrm{U}}(t) = \begin{cases} 1 & 0 \le t < t_1 \\ f_1 e^{-k_1(t-t_1)} + f_2, & t_1 \le t < t_2 \\ f_1 e^{-k_1(t-t_1)} + f_2 e^{-k_2(t-t_2)}, & t \ge t_2 \end{cases}, \tag{5}$$

where $f_1$ and $f_2$ are the proportions of the folding events along the first and second pathway, respectively, and thus satisfy $0 < f_1, f_2 < 1$ and $f_1 + f_2 \equiv 1$; $t_1$ and $t_2$ are the corresponding delay times for the first and second folding pathway. Using the double-exponential equation, the fitting quality $R^2$ of protein ABP1, SH3, and Tenascin was above 0.99.

The *in silico* protein folding time $\langle \tau_{\mathrm{F}}^{\mathrm{sim}} \rangle$ was then estimated by

$$\begin{aligned} \langle \tau_{\mathrm{F}}^{\mathrm{sim}} \rangle &= \int_0^{\infty} \frac{d[1 - S_{\mathrm{U}}(t)]}{dt} \cdot t \, \mathrm{d}t \\ &= \begin{cases} t_0 + \dfrac{1}{k_{\mathrm{F}}}, & \text{for single-pathway kinetics;} \\ f_1\left(t_1 + \dfrac{1}{k_1}\right) + f_2\left(t_2 + \dfrac{1}{k_2}\right), & \text{for double-pathway kinetics;} \end{cases} \end{aligned} . \tag{6}$$

The scaling factor $\alpha$ of the *in silico* protein folding time $\langle \tau_{\mathrm{F}}^{\mathrm{sim}} \rangle$ with respect to the experimental protein folding time $\langle \tau_{\mathrm{F}}^{\mathrm{exp}} \rangle$ was calculated as $\alpha = \overline{\langle \tau_{\mathrm{F}}^{\mathrm{exp}} \rangle / \langle \tau_{\mathrm{F}}^{\mathrm{sim}} \rangle}$. All the relevant quantities for calculating $\langle \tau_{\mathrm{F}}^{\mathrm{sim}} \rangle$ and $\alpha$ are listed in Supplementary Table 4.

##### 5. CG model of ribosome

The high-resolution crystal structure PDB 4v9d[18] was used to build a coarse-grain model of the *E. coli* ribosome. To model the tRNA molecules present in the A-site and P-site, we inserted the A- and P-site tRNAs obtained from PDB 5jte[19] into the A-site and P-site of PDB 4v9d by aligning the 50S subunit of PDB 4v9d to the 50S subunit of PDB 5jte. The entire 50S subunit, including

23S rRNA, 5S rRNA, ribosomal proteins, A-site tRNA and P-site tRNA, was coarsen-grained using the three/four-point model of RNA[1] and the Cα model of protein[1]. Briefly, nucleotides containing pyrimidines and purines were represented as 3 and 4 interaction sites, respectively, with one interaction site (P) located at the phosphate position and having a q = -1*e* charge, another (R) at the centroid of the ribose ring, and one (named as PU1 and PU2, or PY) at the centroid of each conjugated ring in the base; and amino acid residues were represented as one interaction site at the Cα position with corresponding net charge. To accelerate the simulations we truncated the large subunit of the CG ribosome model by retaining those interaction sites within 30 Å of the center line of the exit tunnel, the interaction sites within 20 Å of the peptidyl transferase center (PTC, selected as A2602 in the 23S rRNA of the *E. coli* ribosome) and the interaction sites at the ribosome surface near the tunnel exit, resulting in 4,577 interaction sites in the cropped CG *E. coli* ribosome. The location and shape of the exit tunnel were detected by MOLE*online*[20, 21].

##### 6. CG force field for ribosome-nascent-chain complex (RNC)

The CG ribosome model was held fixed during the simulations except the ribosomal protein L24, which has a long loop that overhangs the exit tunnel opening. We therefore used the CG protein forcefield shown in Supplementary equation (1) to parameterize the *E. coli* ribosomal protein L24 and made residues 42-59 flexible. The fixed part of the CG ribosome model has no intra-ribosome interactions but has interactions with the nascent chain and the flexible part of the ribosome. The potential energy of the CG RNC model is thus described as

$$\begin{aligned} E_{\text{tot}} &= E_{\text{tot}}^{\text{flex-flex}} + E_{\text{tot}}^{\text{flex-fixed}} + E_{\text{tot}}^{\text{R-NC}} + E_{\text{tot}}^{\text{NC-NC}} \\ &= E_{\text{bonding}}^{\text{R-NC}} + \sum_{i\in\{\text{flex} \cup \text{flex|fixed} \cup \text{NC}\}} K_{\text{b}}(b_i - b_0)^2 \\ &\quad + \sum_{i\in\{\text{flex} \cup \text{flex|fixed} \cup \text{NC}\}} -\frac{1}{\gamma}\ln\left\{e^{-\gamma\left[K_\alpha(\theta_i-\theta_\alpha)^2+\varepsilon_\alpha\right]} + e^{-\gamma K_\beta(\theta_i-\theta_\beta)^2}\right\} \\ &\quad + \sum_{i\in\{\text{flex} \cup \text{flex|fixed} \cup \text{NC}\}} \sum_{j=1}^{4} K_{\text{D}_j}\left[1+\cos(j\varphi_i - \delta_j)\right] \\ &\quad + \sum_{\substack{i\in\{\text{R} \cup \text{NC}\},\\ j\in\{\text{NC}\}, i\neq j}} \left\{\frac{q_i q_j}{4\pi\varepsilon_0\varepsilon_\gamma r_{ij}} \cdot e^{-\frac{r_{ij}}{l_\text{D}}} + \varepsilon_{ij}\left[13\left(\frac{R_{ij}}{r_{ij}}\right)^{12} - 18\left(\frac{R_{ij}}{r_{ij}}\right)^{10} + 4\left(\frac{R_{ij}}{r_{ij}}\right)^{6}\right]\right\}, \quad (7) \end{aligned}$$

where $E_{\text{tot}}^{\text{flex-flex}}$ represents the intra-ribosome interactions of the flexible part (ribosomal protein L24 of the *E. coli* ribosome) using Supplementary equation (1); $E_{\text{tot}}^{\text{NC-NC}}$ represents the intra-NC interactions using Supplementary equation (1); $E_{\text{tot}}^{\text{flex-fixed}}$ represents the intra-ribosome interactions between the flexible part and the fixed part; and $E_{\text{R-NC}}$ represents the interactions between NC and the ribosome. Parameters of both $E_{\text{tot}}^{\text{flex-flex}}$ and $E_{\text{tot}}^{\text{NC-NC}}$were obtained from the corresponding native structures using the method described in Supplementary Methods Sections 1 to 3. All the nonbonded interactions in $E_{\text{tot}}^{\text{flex-fixed}}$ and $E_{\text{tot}}^{\text{R-NC}}$ were treated as non-native contacts and calculated in the same manner described in Supplementary Methods Section 1 using the parameters shown in Supplementary Table 6. The parameter $R_i$ of the ribosomal protein interaction sites were derived from the collision diameters $\sigma_i$ used in our previous work[1], while $R_i$ of the RNA interaction sites were calculated from the collision diameters as $\sigma_i = 2 \cdot \langle \bar{d}_{ij}(\text{R-NC}) - \frac{1}{2}\sigma_j \rangle$. The term $\bar{d}_{ij}(\text{R-NC})$ is the average minimum distance between the interaction site type *i* observed in the ribosome and interaction site type *j* observed in NC, whose collision diameter is $\sigma_j$, and was measured by using 6 high-resolution cryo-EM structures of RNCs (PDB 3jbu [22], 4uy8 [23], 5jte [19], 5ju8 [19], 5nwy [24] and 6i0y [25]). $E_{\text{tot}}^{\text{flex-fixed}}$ has extra bonding interactions (including bond, angle and dihedral terms, same as those in Supplementary equation (1)) for the linking bonds at flexible-fixed interface (flex|fixed), i.e., the bond between residue 41 and 42 and the bond between residue 59 and 60 of the ribosomal protein L24 of the *E. coli* ribosome. $E_{\text{tot}}^{\text{R-NC}}$ also has bonded interactions ($E_{\text{bonding}}^{\text{R-NC}}$) for the linking bonds at the NC-tRNA interface, where the force field varies at different steps within the continuous synthesis process, and is described in Supplementary Methods Section 7.

##### 7. Protein synthesis simulation protocol

The on-ribosome protein synthesis process was simplified as containing three sub-steps at each time the nascent chain elongates, as shown in Supplementary Fig. 1: (1) the cognate aminoacyl tRNA (aa-tRNA) binds at the A-site; (2) the *l*-length nascent-chain-tRNA complex ($NC_l$-tRNA) at the P-site and the aa-tRNA at the A-site undergo the peptidyl transfer reaction catalyzed by the PTC and then forms $NC_{l+1}$-tRNA at the A-site; (3) the P-site tRNA is translocated to the E-site and the A-site tRNA is translocated to the P-site, resulting in an empty A-site for the next round of aa-tRNA binding. We denote the processes from step (1) to step (2), step (2) to step (3) and step (3) to next round step (1) as PT, TL and AB, respectively.

The aa-tRNA molecule was modeled by adding an extra harmonic bond between the amino acid interaction site (CA) and the last nucleotide ribose interaction site (tRNA:76@R, where ':'

refers to residues and '@' refers to atoms; similarly hereafter), which introduces extra angle and dihedral energy terms into the aa-tRNA bonding potential energy $E_{\text{aa-tRNA}}^{\text{bonding}}$, resulting in

$$\begin{aligned} E_{\text{aa-tRNA}}^{\text{bonding}} &= E_{\text{bond}} + E_{\text{angle}} + E_{\text{dihedral}} \\ &= K_{\text{b}}(b - b_0)^2 + \sum_i K_{\theta}(\theta_i - \theta_0)^2 + K_{\varphi}(\varphi - \varphi_0)^2, \end{aligned} \tag{8}$$

where $K_{\text{b}}$ is the bond force constant and set as 200 kcal/mol/Å$^2$; $b$ is the pseudo bond length between CA and tRNA:76@R; $b_0$ is the corresponding equilibrium pseudo bond length; $K_{\theta}$ is the angle force constant and set as 25 kcal/mol/rad$^2$; $\theta_i$ includes the angle of CA–tRNA:76@R–tRNA:76@PU2 and the angle of CA–tRNA:76@R–tRNA:76@P; $\theta_0$ is the corresponding equilibrium angle; $K_{\varphi}$ is the dihedral force constant and set as 25 kcal/mol/rad$^2$; $\varphi$ is the dihedral angle of plane CA–tRNA:76@R–tRNA:76@P and plane tRNA:76@R–tRNA:76@P–tRNA:76@PU2; $\varphi_0$ is the corresponding equilibrium dihedral angle.

The NC$_l$-tRNA molecule was modeled by adding an extra harmonic bond between the last amino acid interaction site (NC:*l*@CA) and tRNA:76@R, which introduces the same energy terms with $E_{\text{aa-tRNA}}^{\text{bonding}}$ when *l* = 1. When *l* > 1, this bond introduces an extra double-well angle energy term into the $E_{\text{NC}_l\text{-tRNA}}^{\text{bonding}}$, resulting

$$E_{\text{NC}_l\text{-tRNA}}^{\text{bonding}} = E_{\text{aa-tRNA}}^{\text{bonding}} - \frac{1}{\gamma}\ln\left\{e^{-\gamma[K_{\alpha}(\theta-\theta_{\alpha})^2+\varepsilon_{\alpha}]} + e^{-\gamma K_{\beta}(\theta-\theta_{\beta})^2}\right\}, \tag{9}$$

where $\theta$ is the angle of NC:(*l*-1)@CA–NC:*l*@CA–tRNA:76@R and the parameters were set as those listed in Supplementary Table 1. All the equilibrium values for both P-site and A-site NC-tRNA (or aa-tRNA) presented in Supplementary equation (8) were set as the average values observed in high-resolution experimental structures: 3jbu [22], 4uy8 [23], 4v5c [26], 5jte [19], 5ju8 [19], 5nwy [24], 6enj [27], 6enu [27] and 6i0y [25]; and are shown in Supplementary Table 7.

The bonding interaction energy term $E_{\text{bonding}}^{\text{R-NC}}$ in Supplementary equation (7) during synthesizing at NC length *l* therefore can be calculated as

$$E_{\text{bonding}}^{\text{R-NC}} = \begin{cases} E_{\text{NC}_{l-1}\text{-tRNA}}^{\text{bonding}}(\text{P-site}) + E_{\text{aa-tRNA}}^{\text{bonding}}(\text{A-site}), & \text{within the process PT} \\ E_{\text{NC}_l\text{-tRNA}}^{\text{bonding}}(\text{A-site}), & \text{within the process TL} \\ E_{\text{NC}_l\text{-tRNA}}^{\text{bonding}}(\text{P-site}), & \text{within the process AB} \end{cases}, \tag{10}$$

where (P-site) means to use the equilibrium bond lengths, angles and dihedrals obtained from P-site tRNA; and (A-site) means to use the equilibrium values obtained from A-site tRNA (see Supplementary Table 7). Note that $E^{\text{bonding}}_{\text{NC}_{l-1}\text{-tRNA}}(\text{P-site}) = 0$ when *l* = 1. There are no constraints added on the bond lengths in $E^{\text{R-NC}}_{\text{bonding}}$.

To setup a continuous synthesis simulation, we used the corresponding mRNA sequence of the protein to be synthesized, the truncated CG ribosome structure of the corresponding organism and the CG RNC force field with optimized $n_{\text{scal}}$ for this protein. The synthesis begins at the start codon and ends at the stop codon. At each codon position, we obtained the initial RNC structure from the final structure calculated at the previous codon position, followed by putting the new amino acid interaction site near the A-site tRNA:76@R with an initial orientation based on the experimental data listed in Supplementary Table 7. At each sub-step, an energy minimization was performed on the last 15 residues of the C-terminal of NC, as well as the new amino acid at step 1, followed by running a MD simulation for a time $\tau^{\text{sim}}$. At step 1, we connected the bond between the new amino acid and the A-site tRNA. At step 2, we connect the ($l$-1)$^{\text{th}}$ amino acid interaction site to the new amino acid at A-site tRNA and break the original bond between the ($l$-1)$^{\text{th}}$ amino acid interaction site and the P-site tRNA. At step 3, the *l*-length NC was covalently connected to the P-site tRNA and the empty A-site was modeled by excluding all the nonbonded interactions of A-site tRNA with other interaction sites. The simulation time $\tau^{\text{sim}}$ at each step of each codon position was randomly sampled from the exponential distribution with the expectation $\langle\tau^{\text{sim}}\rangle$. Details on derivation of $\langle\tau^{\text{sim}}\rangle$ can be found in the Supplementary Methods Section 8.

Once the nascent chain was fully synthesized, the bond between NC:*l*@CA and tRNA:76@R was removed, as well as the corresponding bonding interaction energy terms. The nascent chain was ejected from the exit tunnel and then dissociated from the ribosome surface. We identified the complete dissociation as when the minimum distance between the nascent chain and the ribosome surface is greater than 20 Å for at least 750 ps. The final nascent chain structure was then used to study the off-ribosome post-translational folding.

#### 8. Setting realistic *in silico* codon translation times

Even though the folding process is accelerated in the coarse-grained model, we maintain a realistic ratio of folding and translation times. To set a realistic *in silico* mean translation time $\langle\tau^{\text{sim}}_{\text{trans}}(S)\rangle_i$ for codon *i* from the corresponding real mean translation time $\langle\tau^{\text{real}}_{\text{trans}}(S)\rangle_i$ given the mRNA sequence *S* and the organism, we use the scaling factor *α* from *in silico* protein folding times (see Supplementary Methods Section 4 and Supplementary Table 4) as $\langle \tau_{\text{trans}}^{\text{sim}}(S) \rangle_i = \frac{1}{\alpha} \langle \tau_{\text{trans}}^{\text{real}}(S) \rangle_i$. This takes into account of the influence of ribosome traffic $\left\langle \tau_{\text{trans}}^{\text{ribo-traffic}}(S) \right\rangle_i$ on the intrinsic translation time $\langle \tau_{\text{trans}}^{\text{intrinsic}} \rangle_i$ and is expressed as

$$\begin{aligned} \langle \tau_{\text{trans}}^{\text{real}}(S) \rangle_i &= \langle \tau_{\text{trans}}^{\text{intrinsic}} \rangle_i + \langle \tau_{\text{trans}}^{\text{ribo-traffic}}(S) \rangle_i \\ &= \langle \tau_{\text{AB}}^{\text{intrinsic}} \rangle_i + \langle \tau_{\text{PT}}^{\text{intrinsic}} \rangle + \langle \tau_{\text{TL}}^{\text{intrinsic}} \rangle + \langle \tau_{\text{trans}}^{\text{ribo-traffic}}(S) \rangle_i \\ &= \langle \tau_{\text{AB}}^{\text{intrinsic}} \rangle_i + \langle \tau_{\text{PT}}^{\text{intrinsic}} \rangle + \langle \tau_{\text{TL}}^{\text{real}}(S) \rangle_i, \end{aligned} \tag{11}$$

where $\langle \tau_{\text{AB}}^{\text{intrinsic}} \rangle_i$ is the intrinsic mean dwell time of the A-site aa-tRNA binding at codon *i* taking into account the competition of the non- and near-cognate tRNAs binding; $\langle \tau_{\text{PT}}^{\text{intrinsic}} \rangle$ is the intrinsic mean dwell time of the peptidyl transfer reaction at the PTC after A-site aa-tRNA binding and is independent of the codon identity; $\langle \tau_{\text{TL}}^{\text{real}}(S) \rangle_i$ is the real mean dwell time of tRNA translocation after the peptidyl transfer reaction and can be calculated as the summation of the intrinsic mean dwell time of translocation $\langle \tau_{\text{TL}}^{\text{intrinsic}} \rangle$ and the mean delay time due to the ribosome traffic $\left\langle \tau_{\text{trans}}^{\text{ribo-traffic}}(S) \right\rangle_i$ at codon *i*. $\langle \tau_{\text{trans}}^{\text{real}}(S) \rangle_i$ was estimated by a kinetic model proposed in our previous study[28] using the particular mRNA sequence *S*, the translation initiation rate and the intrinsic codon translation time of the particular organism. The translation initiation rate was set as the median value observed in experiments and simulations, which is 0.083 $s^{-1}$ for *E. coli*[29].

The intrinsic codon translation times $\langle \tau_{\text{trans}}^{\text{intrinsic}} \rangle_i$ of *E. coli*, as well as the intrinsic dwell times of sub-steps ($\langle \tau_{\text{PT}}^{\text{intrinsic}} \rangle$ and $\langle \tau_{\text{TL}}^{\text{intrinsic}} \rangle$), were obtained from Fluitt & Viljoen's work[30] (denoted as $\langle \tau_{\text{Fluitt}}^{\text{intrinsic}} \rangle_i$), followed by rescaling the data so that the average codon translation time equals the experimental average $\overline{\tau_{\text{trans}}^{\text{exp}}}$:

$$\langle \tau_{\text{trans}}^{\text{intrinsic}} \rangle_i = \frac{\overline{\tau_{\text{trans}}^{\text{exp}}}}{\sum_{k=1}^{64} w_k \cdot \langle \tau_{\text{Fluitt}}^{\text{intrinsic}} \rangle_k} \cdot \langle \tau_{\text{Fluitt}}^{\text{intrinsic}} \rangle_i, \tag{12}$$

where $w_k$ is the normalized codon usage frequency[31] of the $k^{\text{th}}$ codon in *E. coli*. The experimental codon translation rate varies from 12 to 21 $s^{-1}$ in *E. coli* at 310 K[32], resulting in an average rate of 16.5 $s^{-1}$. Thus, $\overline{\tau_{\text{trans}}^{\text{exp}}}$ was set as 0.061 s. All of the values for $\langle \tau_{\text{trans}}^{\text{intrinsic}} \rangle_i$, $\langle \tau_{\text{PT}}^{\text{intrinsic}} \rangle$, $\langle \tau_{\text{TL}}^{\text{intrinsic}} \rangle$ and $w_k$ can be found in Supplementary Table 8.

From Supplementary equation (11), the *in silico* mean dwell times of the sub-steps during translating codon *i* were then calculated as

$$
\begin{cases}
\left\langle \tau_{\text{AB}}^{\text{sim}} \right\rangle_i = \dfrac{1}{\alpha} \left\langle \tau_{\text{AB}}^{\text{intrinsic}} \right\rangle_i \\
\left\langle \tau_{\text{PT}}^{\text{sim}} \right\rangle = \dfrac{1}{\alpha} \left\langle \tau_{\text{PT}}^{\text{intrinsic}} \right\rangle \\
\left\langle \tau_{\text{TL}}^{\text{sim}}(S) \right\rangle_i = \dfrac{1}{\alpha} \left\langle \tau_{\text{TL}}^{\text{real}}(S) \right\rangle_i = \dfrac{1}{\alpha} \left[ \left\langle \tau_{\text{TL}}^{\text{intrinsic}} \right\rangle + \left\langle \tau_{\text{trans}}^{\text{ribo-traffic}}(S) \right\rangle_i \right] \\
\qquad\qquad = \dfrac{1}{\alpha} \left[ \left\langle \tau_{\text{TL}}^{\text{intrinsic}} \right\rangle + \left\langle \tau_{\text{trans}}^{\text{real}}(S) \right\rangle_i - \left\langle \tau_{\text{trans}}^{\text{intrinsic}} \right\rangle_i \right]
\end{cases}, \tag{13}
$$

#### 9. Virtual screening for proteins that get kinetically trapped during folding

To identify candidates whose catalytic property might be affected by synonymous mutations, we identified an initial set of enzymes by searching the databases EzCatDB[33, 34], UniProt[35] and PDB[36]. This initial set was of enzymes that (1) are endogenetic enzymes from *E. coli*; (2) have a known catalytic mechanism; (3) can be active as a monomer; (4) are located in the cytoplasm and not transmembrane; (5) have resolved crystal or NMR structures with bound substrates; and (6) do not have missing domains in their resolved structures (having several missing loops or fragments is acceptable). The catalytic mechanism information was obtained from EzCatDB; the enzymology information was obtained from UniProt; and the structural information was obtained from PDB. We found 14 *E. coli* enzymes that met these criteria. We chose the resolved structure that has the highest resolution and has the most relevant substrate binding, for those enzymes that have more than one resolved structure. The structural domains of those initial enzymes were then identified by using the CATH database and parameterized using our CG protein model (see Supplementary Methods Section 1) via the stepwise parameterization strategy (see Supplementary Methods Section 3). All the structural information and the parameterization information for these enzymes can be found in Supplementary Table 9. The wild-type enzymes were then continuously synthesized on the corresponding ribosome, followed by post-translational folding simulations (both have 10 replicas). The post-translational folding simulations for all the initial enzymes were stopped at a 14-day wall time. The candidates were then selected by using the score function equation (1) in the main text. The time point when the protein reaches the folded state was used to estimate $\left\langle \tau_{\text{F}}^{\text{post}} \right\rangle$ in the score function and was identified as the time when $Q^{\text{mod}}$ is higher than $\langle Q_{\text{native}}^{\text{mod}} \rangle - 3\sigma$ for 7.5 ns in simulation timescale. $Q^{\text{mod}}$ is the mode of Q-values calculated by histogramming a moving window of 200 data points along with the post-translational time course using a bin width 0.02;

$\langle Q_{\text{native}}^{\text{mod}} \rangle$ is the average of mode-filtered *Q* values of the native state calculated by using the 10 trajectories obtained from the final iteration of the stepwise parameterization strategy; and $\sigma$ is the standard deviation.

**10. Parameterization of the CG models for CAT-III**

CAT-III monomer is a single domain protein and was parameterized using the same approach as the training-set proteins (Supplementary Methods Section 2). The folding stability, -10.1 kcal/mol, was estimated via PREFUR algorithm[37] at 298 K. The native structure was obtained from PDB ID 3cla[38] and chain A was selected to create the CG model. Three $n_{\text{scal}}$ values (1.0, 1.1 and 1.2) were utilized to search for the optimized $n_{\text{scal}}^{*}$ value. At each $n_{\text{scal}}$, two pt-REMD simulations were initialized from native state and unfolded state, respectively. The parameterization results for CAT-III are shown in Supplementary Table 10.

**11. Detecting noncovalent lasso entanglements in protein structures**

To detect noncovalent lasso entanglements we use linking numbers[39], which requires at least one closed loop as an argument. We define this loop as being composed of the backbone trace connecting residues *i* and *j*, which have formed a native contact in the given protein conformation. The native contact between *i* and *j* is considered to close this loop, even though there is no covalent bond between these two residues. Outside this loop is an N-terminal segment, composed of residues 1 through *i*-1, and C-terminal segment composed of residues *j*+1 through *N*. These two segments represent open curves, whose entanglement through the closed loop we characterize with linking numbers denoted $g_{\text{N}}$ and $g_{\text{C}}$. We calculate these numbers using the partial Gauss double integration method proposed by Baiesi and co-workers[40]. For a given structure of an *N*-length protein, with a native contact present at residues (*i*, *j*), the coordinates $\boldsymbol{R}_l$ and the gradient $\mathrm{d}\boldsymbol{R}_l$ of the point *l* on the curves were calculated as

$$\begin{cases} \boldsymbol{R}_l = \dfrac{1}{2}(\boldsymbol{r}_l + \boldsymbol{r}_{l+1}) \\ \mathrm{d}\boldsymbol{R}_l = \boldsymbol{r}_{l+1} - \boldsymbol{r}_l \end{cases}, \quad (14)$$

where $\boldsymbol{r}_l$ is the coordinates of the Cα atom in residue *l*. The linking numbers $g_{\text{N}}(i,j)$ and $g_{\text{C}}(i,j)$ were calculated as

$$\begin{cases} g_{\text{N}}(i,j) = \dfrac{1}{4\pi} \displaystyle\sum_{m=6}^{i-5} \sum_{n=i}^{j-1} \frac{\boldsymbol{R}_m - \boldsymbol{R}_n}{|\boldsymbol{R}_m - \boldsymbol{R}_n|^3} \cdot (\mathrm{d}\boldsymbol{R}_m \times \mathrm{d}\boldsymbol{R}_n) \\ g_{\text{C}}(i,j) = \dfrac{1}{4\pi} \displaystyle\sum_{m=i}^{j-1} \sum_{n=j+4}^{N-6} \frac{\boldsymbol{R}_m - \boldsymbol{R}_n}{|\boldsymbol{R}_m - \boldsymbol{R}_n|^3} \cdot (\mathrm{d}\boldsymbol{R}_m \times \mathrm{d}\boldsymbol{R}_n) \end{cases}, \quad (15)$$

where we excluded the first 5 residues on the N-terminal curve, last 5 residues on the C-terminal curve and 4 residues before and after the native contact for the purpose of eliminating the error induced by both the high flexibility and contiguity of those tails. The above integrations yield two non-integer values, therefore the total linking number for a native contact (*i*, *j*) was estimated as

$$g(i,j) = \text{round}\big(g_{\text{N}}(i,j)\big) + \text{round}\big(g_{\text{C}}(i,j)\big). \quad (16)$$

As an illustration and test case we analyzed the post-translational trajectories of fast CAT-III variant. We calculated the maximum $g(i,j)$ for every frame of the trajectories and found that the maximum $g(i,j)$ varies from -2 to 2. We randomly chose structures corresponding to each maximum $g(i,j)$ detected and compared the structure with the corresponding linking diagram shown in Supplementary Fig. 2. We clearly see entanglements in these structures. As shown in Supplementary Fig. 2 b-f, all the maximum $g(i,j)$ values estimated from Supplementary equation (16) are consistent with those calculated as the total number of all the +1 and -1 crossings in the link diagram (left side in each panel) divided by 2.

#### 12. Assessing the influence of synonymous mutations on co- and post-translational folding

The translation schedules for the fastest (denoted as fast variant) and the slowest synonymous mRNA variant (denoted as slow variant) were predicted according to Fluitt & Viljoen's codon translation time in Supplementary Table 8. We performed continuous synthesis simulations for the fast and slow variants using 50 trajectories for DHFR and 100 trajectories for CAT-III and DDLB, followed by post-translational folding simulations in the bulk environment for 60 seconds on the experimental timescale (13.85 μs of simulation time). The number of post-translational trajectories was increased to 10 times larger than the continuous synthesis simulations (500 trajectories for DHFR and 1,000 trajectories for CAT-III and DDLB) for each variant. To reduce the computational cost, the post-translational simulation of each trajectory was early terminated if the protein folded, defined as $Q^{\text{mod}}$ attaining a value higher than $\langle Q^{\text{mod}}_{\text{native}}\rangle - 3\sigma$ for 7.5 ns of simulation time; $\langle Q^{\text{mod}}_{\text{native}}\rangle$ is the average of mode-filtered $Q$ values of the native state calculated by using 10, 1-μs CG simulation trajectories of the native state; and $\sigma$ is the standard deviation.

The structural distribution from the post-translational simulations was assessed as the free energy along the order parameters $G$ and $Q_{\text{act}}$ as described in the main text (equations (2) and (3)). To further analyze the post-translational kinetics, 400 clusters (micro-states) were grouped from the combined set of post-translational trajectories of both synonymous variants using the *k*-means algorithm[41, 42]. A Markov state model (MSM) was built and the clusters were coarsen-grained into a small number of metastable states using the PCCA+ algorithm[43]. The number of metastable states was chosen based on the existence of a gap in the eigenvalue spectrum of the transition probability matrix[44]. Five representative structures of each metastable state were randomly sampled from all microstates according to the probability distribution of the microstates within the given metastable state. The time evolution of the probability distribution of metastable states (see Supplementary Methods Section 14.1), as well as the representative structures, were used to estimate the specific activity of a synonymous variant. All the clustering and MSM building were performed by using PyEmma package[45].

#### 13. Back-mapping the CG protein model to an all-atom (AA) model

To be able to study enzymatic catalysis at the AA resolution, a customized back-mapping procedure was utilized to rebuild AA models for those CG protein structures. The CG interaction sites that represent the side-chain center-of-mass were first built near the corresponding Cα interaction sites, whose orientation was further refined by using energy minimizations under the Cα-SCM force field[46] with all Cα positions restrained. The backbone atoms were then rebuilt by Prodart2[47] and the side-chain atoms were rebuilt by Pulchra[48]. The final AA structure was obtained after a further energy minimization in vacuum with all Cα positions restrained.

#### 14. Calculating the specific activity from the structural distribution of enzyme conformations

##### 14.1. Estimation of specific activity for an enzyme

The specific activity (SA) for a given protein variant is estimated by utilizing the distribution of metastable states (whose state probability is $p_i$ for metastable state *i*) and the activation free energy barrier height ($\Delta G_i^{\ddagger}$) of each state as described in the main text (equation (4)), as well as the relative specific activity (SA*, see main text (equation (5)). To reduce the error of estimated reaction rates, $\Delta G_i^{\ddagger}$ was obtained as the median value of the successfully estimated activation barrier heights of the representative structures for each metastable state. To account for only the soluble conformations, the state probability $p_i$ was estimated as $p_i = p_i^0(60) \cdot f_i^{\mathrm{sol}} / \left[\sum_i p_i^0(60) \cdot f_i^{\mathrm{sol}}\right]$, where $f_i^{\mathrm{sol}}$ is the percent soluble protein of state *i* (see details in Supplementary Methods Section 14.2) and $p_i^0(60)$ is the raw probability of state *i* at post-translational time 60 s that was obtained by solving the master equation

$$\frac{\mathrm{d}\boldsymbol{P}(t)}{\mathrm{d}t} = \boldsymbol{M} \cdot \boldsymbol{P}(t), \tag{17}$$

where $\boldsymbol{P}(t)$ is a column vector of the state probabilities $\boldsymbol{P}(t) = [p_1^0(t), p_2^0(t), \cdots, p_N^0(t)]^{\mathrm{T}}$ and $\boldsymbol{M}$ is the rate matrix constructed by the transition rates $\omega_{ij}$ (transition from state *i* to *j*)

$$\boldsymbol{M} = \begin{bmatrix} -\sum_{j\neq 1} \omega_{1j} & \omega_{21} & \cdots & \omega_{N1} \\ \omega_{12} & -\sum_{j\neq 2} \omega_{2j} & \cdots & \omega_{N2} \\ \vdots & \vdots & \ddots & \vdots \\ \omega_{1N} & \omega_{2N} & \cdots & -\sum_{j\neq N} \omega_{Nj} \end{bmatrix}. \tag{18}$$

The transition rates $\omega_{ij}$ was estimated using[49]

$$\omega_{ij} = -\frac{1}{\Delta t} \cdot \frac{C_{ij}}{\sum_{i\neq j} C_{ij}} \cdot \ln\left(\frac{C_{ij}}{\sum_j C_{ij}}\right), \tag{19}$$

where $C_{ij}$ is the total number of transitions from state *i* to *j* and $\Delta t$ is the lag time for observing transitions, which is 0.32 ms on the experimental timescale. $C_{ij}$ was calculated by using the discrete metastable state trajectories obtained in Supplementary Methods Section 12. For the discrete trajectories whose length is shorter than 60 s, we extended the length to 60 s by repeating the most frequently sampled metastable state that were obtained in the last 200 frames of the original trajectory. The master equation (Supplementary equation (17)), with the initial value $\boldsymbol{P}(0)$ obtained from the state distribution adopted at the end of synthesis, was solved numerically using the Explicit Runge-Kutta method of order 5(4)[50] from time *t* = 0 s to *t* = 60 s.

##### 14.2. The percent of soluble protein from each metastable state

The percent soluble protein $f_i^{\text{sol}}$ was estimated as

$$f_i^{\text{sol}} = \frac{\chi_{\text{disordered}}^{\text{insol}} - \chi_i^{\text{insol}}}{\chi_{\text{disordered}}^{\text{insol}} - \min\left(\chi_i^{\text{insol}}\right)} \times 100\%, \tag{19}$$

where $\chi_i^{\text{insol}}$, $\chi_{\text{disordered}}^{\text{insol}}$ and $\min\left(\chi_i^{\text{insol}}\right)$ are the insolubility propensities of state *i*, the fully disordered structure and the minimum propensity value throughout all states, respectively. The fully disordered protein has no secondary and tertiary structure elements and was built as a linear sequence of amino acid residues. The insolubility propensity $\chi^{\text{insol}}$ was estimated by taking into account of the aggregation propensity $\chi^{\text{agg}}$, degradation propensity $\chi^{\text{deg}}$ and subtracting the Hsp70 binding propensity $\chi^{\text{Hsp70}}$ that is considered to prevent the misfolding protein from aggregation:

$$\chi^{\text{insol}} = \chi^{\text{agg}} + \chi^{\text{deg}} - \chi^{\text{Hsp70}}. \tag{20}$$

For a given metastable state, the aggregation, degradation and Hsp70 binding propensities were evaluated as the relative change of the mean solvent accessible surface area (SASA) of the aggregation-prone, degradation-prone and Hsp70 binding regions of the representative protein structures (or the modeled fully disordered structure) against the mean SASA of the native state, respectively:

$$\chi = \frac{\langle \text{SASA} \rangle}{\langle \text{SASA} \rangle_{\text{native}}} - 1. \tag{21}$$

A negative $\chi$ value indicates a lower propensity than the native state. The aggregation-prone region was predicted by the AmylPred2 server[51]. The degradation-prone region was defined as the hydrophobic residues (Ile, Val, Leu, Phe, Cys, Met, Ala, Gly and Trp). The Hsp70 binding region was predicted by using ChaperISM[52]. Note that, for CAT-III, the trimerization interface area (residue 25 to 33 and 150 to 157) was subtracted from the above SASAs. The 95% CIs of the propensities and the percent soluble protein were estimated using the bootstrapping method with 10,000 iterations.

##### 14.3. Estimating the activation free energy barrier for a protein conformation

We developed the following workflow (illustrated in Supplementary Fig. 3) to estimate activation free energy barrier height for a given enzyme structure:

(1) Take the representative structure (CG protein structure) of a metastable state. Back-map the CG structure to its AA structure.

(2) Refine the rebuilt AA structure by running energy minimizations in vacuum for 1,000 steps with all Cα positions restrained using a force constant 30 kcal/mol/Å$^2$ under a dielectric constant 78.5 to relax the side chain orientations, followed by running energy minimizations in a water box for 2,000 steps with all Cα positions restrained using a force constant 100 kcal/mol/Å$^2$ to relax water molecules and ions. The final refined AA structure was obtained after running energy minimizations in water for 2,000 steps with all Cα positions restrained using a force constant 10 kcal/mol/Å$^2$.

(3) For an enzyme that can only carry out the reaction in its homomultimer form, an estimation of the multimer structure was performed by using SymmDock program[53]. For example, CAT-III needs to form a trimer to conduct the reaction. The monomer structure of CAT-III obtained from step (2) was first refined to form a native-like trimer interface by running targeted energy minimizations in vacuum for 2,000 steps, where an RMSD restraint on the trimer interface region towards the corresponding native structure (crystal structure) with a force constant 100 kcal/mol/Å$^2$ was applied. The refined monomer structure was then used to predict

the trimer structure using SymmDock with distance restraints between monomers at the interface region to force the trimer to be formed on the native interface. If the prediction succeeded, the best candidate was obtained and the trimer interface was refined by running targeted energy minimizations in vacuum for 2,000 steps, where an RMSD restraint on the trimer interface region towards the corresponding native structure with a force constant 10 kcal/mol/Å$^2$ were applied. The final refined trimer structure was obtained after running a series of MD simulations in a water box: First, run energy minimizations for 2,000 steps with all Cα positions restrained using a force constant 10 kcal/mol/Å$^2$; Second, heat the system to 310 K in NVT ensemble with all Cα positions restrained using a force constant 10 kcal/mol/Å$^2$; Third, equilibrate the system at 310 K in the NPT ensemble for 0.5 ns with all Cα positions restrained using a force constant 5 kcal/mol/Å$^2$; Finally, run energy minimizations again for 500 steps with all Cα positions restrained using a force constant 10 kcal/mol/Å$^2$.

(4) After the AA structure of the monomer or multimer was refined, the structure was then used to predict the ligand-binding poses using AutoDock vina[54]. The docking boxes were set as the minimum box that covers each ligand presented in the crystal structure. The best binding pose (highest score) of each ligand was used to build the protein-ligand complex structure. The complex structure was then refined by running the following MD simulations: First, solvate the complex in TIP3P water molecules and neutralization ions (Na+ or Cl-); Second, run energy minimizations for 2,000 steps with all Cα positions restrained using a force constant 100 kcal/mol/Å$^2$, followed by running energy minimizations for 2,000 steps with all Cα positions restrained using a force constant 5 kcal/mol/Å$^2$; Third, heat the system to 310 K in the NVT ensemble with all Cα positions restrained using a force constant 3 kcal/mol/Å$^2$ for 20 ps; Forth, equilibrate the system in the NPT ensemble with all Cα positions restrained using a force constant 1 kcal/mol/Å$^2$ for 400 ps. A few distance restraints between ligands and surrounding residues were added to force the orientation of ligands to be the most native-like. The distance restraints for refining the binding poses of the reactants were set as the following: If the reactant is co-crystalized in the crystal structure, some of the representative interactions were chosen and the distances were restrained to the exact value; and if the reactant is not presented in any crystal structure, the possible interactions were obtained from the literature or inferred theoretically and the distances were restrained less than an expected value. The distance restraint force constant was stepwise increased to 500 kcal/mol/Å$^2$ in the first 200 ps and kept 500 kcal/mol/Å$^2$ in the later 200 ps; Finally, the refined complex structure was obtained after running a classical energy minimization for 2,000 steps, followed by a QM/MM energy minimization for 5,000 steps with the distance restraints between ligands and catalytic residues.

The distance restraint force constant was stepwise increased to 10 kcal/mol/Å$^2$ in the first 2,500 steps and kept 10 kcal/mol/Å$^2$ in the later 2,500 steps.

The activation free energy profile was estimated by running QM/MM umbrella sampling simulations along the predefined reaction coordinates (RCs). In each umbrella window, the simulation was run in the NPT ensemble at 310 K for 20 ps after a 1,000-step energy minimization. The umbrella restraint force constant was set as 250 kcal/mol/Å$^2$. The potential of mean force (PMF) was unbiased and estimated by the WHAM equation[16] using the last 10 ps trajectories (see Supplementary Figures 8, 9, 10). The activation free energy barrier height was obtained as the difference between the maximum PMF value and the minima before the maximum. The 95% CIs were estimated by using the Monte Carlo bootstrap error analysis[55] with 100 trials.

##### 14.4. Classical Molecular Dynamics Simulations

For a vacuum system in the above workflow, the MD simulations were performed without a nonbonded cutoff and using the dielectric constant set as 78.5. For a solvated system, the solute was embedded in a periodic TIP3P water box. Several ions were added to neutralize the system. Particle Mesh Ewald (PME) method[56] was used to calculate the long-range electrostatic interactions with a 10 Å cutoff. The NPT ensemble simulations were performed at 310 K temperature and 1 bar pressure via Langevin dynamics (the collision frequency is 1.0 ps$^{-1}$), with a coupling constant of 0.2 ps for both parameters. The SHAKE algorithm[57] was applied to the bonds involving hydrogen, which ensures the integral timestep to be 2 fs. All the classical MD simulations in the workflow were performed by Amber17[58] with *ff14SB* protein force field[59]. The ligands were parameterized by the general force field *gaff*[60]. The atomic charges of the ligands were estimated as the RESP charges[61] derived from the QM optimization with Hartree–Fock (HF) method at 6-31G* level using Gaussian 09[62].

##### 14.5. QM/MM Simulations

For QM/MM umbrella sampling simulations, the QM region (including ligands and catalytic residues) was simulated using the third-order density functional tight binding (DFTB3) Hamiltonian[63] with 3ob-3-1 parameter set[64, 65, 66, 67]. The MM region was simulated using *ff14SB* protein force field[59]. The QM/MM interface was built by inserting explicit link atoms (hydrogen atoms). The interaction on QM/MM interface was estimated using the electrostatic embedding scheme. Particle Mesh Ewald (PME) method[56] was used to calculate the long-range electrostatic interactions with a 10-Å cutoff. The NPT ensemble simulations were performed at

310 K temperature and 1 bar pressure via the Langevin dynamics (the collision frequency is 1.0 $ps^{-1}$), with a coupling constant of 0.2 ps for both parameters. The SHAKE algorithm[57] was applied to the bonds involving hydrogen in the MM region and no constraints were applied in the QM region to enable the proton transfer. The MD integration timestep was thus set as 1 fs. All the QM/MM umbrella sampling simulations were performed by Amber17[58].

##### 14.6. Detailed setup for each system

***CAT-III.*** The chain A of the crystal structure 3CLA was used as the template for back-mapping. The trimer structure of 3CLA was used as the template for estimating the trimer of representative structures. The trimer interface region in the monomer was identified as residue 25 to 33 and 150 to 157 (the residue index starts from 1 here, whereas it starts from 6 in the PDB file). The distance restraints applied during the prediction of trimer structure are on the distance between residue 25 in two monomers, distance between residue 32 in two monomers, distance between residue 150 in two monomers, distance between residue 157 in two monomers, distance between residue 25 in one monomer and 157 in another, and distance between residue 32 in one monomer and 150 in another. The binding poses of the reactant chloramphenicol (CLM) were searched within the box centered at (-1.000, 14.600, 11.200) Å with the dimension 14 Å × 14 Å × 14 Å. The binding poses of the reactant acetyl coenzyme A (ACO) were searched within the box centered at (-2.062, 16.000, 30.000) Å with the dimension 15 Å × 15 Å × 30 Å. The following distance restraints were used to refine the binding poses with classical MD simulations (residue indices start from 1 and are accumulated throughout the trimer. The Amber-style masking syntax is used to denote the selected atoms, where ':' denotes residuals and '@' denotes atoms, and the unit of distance is in Å, similarly hereinafter): *d*(:402@NE2, :CLM@O1) = 2.830; *d*(:402@NE2, :CLM@N2) = 8.750; *d*(:402@NE2, :CLM@C6) = 5.450; *d*(:402@NE2, :CLM@C2) = 4.020; *d*(:402@NE2, :CLM@C4) = 7.270; *d*(:402@NE2, :CLM@O5) = 5.650; *d*(:402@CG, :CLM@O1) = 4.940; *d*(:402@CG, :CLM@N2) = 9.180; *d*(:402@CG, :CLM@C6) = 6.680; *d*(:402@CG, :CLM@C2) = 5.800; *d*(:402@CG, :CLM@C4) = 8.720; *d*(:402@CG, :CLM@O5) = 7.400; *d*(:402@ND1, :CLM@O1) = 4.890; *d*(:402@ND1, :CLM@N2) = 10.090; *d*(:402@ND1, :CLM@C6) = 7.290; *d*(:402@ND1, :CLM@C2) = 5.970; *d*(:402@ND1, :CLM@C4) = 8.790; *d*(:402@ND1, :CLM@O5) = 7.750; *d*(:CLM@O1, :ACO@C22) < 3.000; *d*(:402@NE2, :ACO@S1) < 3.000; *d*(:402@ND1, :406@OD1) = 3.380. The following distance restraints were used to refine the active site with QM/MM energy minimization: *d*(:402@NE2, :CLM@O1) < 2.830; *d*(:CLM@O1, :ACO@C22) < 3.000; *d*(:402@NE2, :ACO@S1) < 3.000;

*d*(:402@ND1, :406@OD1) < 3.000. The QM region (see Supplementary Fig. 4a) includes reactant CLM, a part of ACO with the acetyl group (from C4 to C22), the side chain of His402[38, 68] and the side chain of Asp406[68]. The reaction coordinate (RC) was defined as RC = $d$(:ACO@C22, :ACO@S1) − $d$(:CLM@O1, :ACO@C22) and discretized into 34 windows: RC = -2.6, -2.4, -2.2, -2.0, -1.8, -1.6, -1.4, -1.2, -1.0, -0.9, -0.8, -0.7, -0.6, -0.5, -0.4, -0.3, -0.2, -0.1, 0.0, 0.1, 0.2, 0.3, 0.4, 0.5, 0.6, 0.7, 0.8, 0.9, 1.0, 1.2, 1.4, 1.6, 1.8, 2.0. The metastable states P9, P10, P11, P12, P13 and P14 were taken to evaluate $\Delta G_i^{\ddagger}$.

***DDLB.*** The chain A of the crystal structure 4C5C was used as the template for back-mapping. Two $Mg^{2+}$ ions (denoted as MC) that facilitate the proper orientation of ATP were docked at position (5.523, 31.727, 65.029) Å and (7.170, 33.772, 62.509) Å, respectively. The $Mg^{2+}$ ions were parameterized by the dummy-atom model[69], which has an enhanced performance in modeling metalloenzymes. The binding poses of reactant ATP were searched within the box centered at (7.538, 30.141, 68.311) Å with the dimension 7 Å × 11 Å × 16 Å. The binding poses of reactant zwitterionic D-alanine (-$NH_3^+$ and -$COO^-$, DAL) were searched within the box centered at (4.876, 31.014, 58.470) Å with the dimension 4 Å × 4 Å × 4 Å. The binding poses of reactant anionic D-alanine (-$COO^-$, DAN) were searched within the box centered at (2.031, 29.261, 60.338) Å with the dimension 4 Å × 4 Å × 4 Å. The following distance restraints were used to refine the binding poses with classical MD simulations: *d*(:272@OD1, :307@MC) = 2.16; *d*(:270@OE1, :307@MC) = 2.20; *d*(:270@OE2, :307@MC) = 2.13; *d*(:270@OE2, :308@MC) = 2.13; *d*(:257@OD2, :308@MC) = 2.05; *d*(:307@MC, :308@MC) = 3.64; *d*(:210@CA, :ATP@O2) = 3.68; *d*(:187@OE1, :ATP@O2) = 2.66; *d*(:187@OE2, :ATP@O1) = 2.43; *d*(:181@C, :ATP@N3) = 4.03; *d*(:144@NZ, :ATP@N2) = 3.00; *d*(:180@OE1, :ATP@N2) = 3.37; *d*(:180@C, :ATP@N3) = 4.67; *d*(:215@NZ, :ATP@O11) = 2.77; *d*(:ATP@O11, :308@MC) = 2.19; *d*(:ATP@O10, :151@CA) = 3.26; *d*(:ATP@O9, :97@NZ) = 2.74; *d*(:ATP@O13, :308@MC) = 2.03; *d*(:ATP@O7, :308@MC) = 1.97; *d*(:ATP@O12, :307@MC) = 1.96; *d*(:ATP@O10, :307@MC) = 2.02; *d*(:255@CZ, :DAL@O1) = 3.73; *d*(:15@OE2, :DAL@N1) = 2.95; *d*(:281@OG, :DAN@O1) = 2.61; *d*(:DAL@C1, :DAN@N1) < 3.50; *d*(:DAL@O2, :ATP@P3) < 3.00; *d*(:DAL@N1, :ATP@O12) = 2.80; *d*(:DAL@C1, :307@MC) = 5.66. The following distance restraints were used to refine the active site with QM/MM energy minimization: *d*(:DAL@C1 :DAN@N1) < 3.00; *d*(:DAL@O2, :ATP@P3) < 3.50; *d*(:255@CZ, :DAL@O1) < 3.00; *d*(:281@OG, :DAN@O1) < 3.00; *d*(:15@OE2, :DAL@N1) < 3.00, as well as all the P-O bonds in the ATP molecule. The QM region (see Supplementary Fig. 4b) includes reactant DAL, DAN, a part of ATP with all the phosphate groups (from C1 to P3) and the side chain of Arg255 (play an important role in stabilizing transition states and intermediates[70]). The reaction coordinate (RC) was defined as RC $=$ $0.6d($:ATP@O11$,$ :ATP@P3$) - 0.6d($:ATP@P3$,$ :DAL@O2$) + 0.4d($:DAL@C1$,$ :DAL@O2$) -$ $0.4d($:DAN@N1$,$ :DAL@C1$)$ and discretized into 33 windows: RC = -1.7, -1.5, -1.4, -1.3, -1.2, -1.1, -1.0, -0.9, -0.8, -0.7, -0.6, -0.5, -0.4, -0.3, -0.2, -0.1, 0.0, 0.2, 0.4, 0.6, 0.8, 0.9, 1.0, 1.1, 1.2, 1.3, 1.4, 1.5, 1.6, 1.7, 1.8, 1.9, 2.1. To ensure the reaction occurs within the RCs, a few extra distance restraints were applied in the umbrella sampling simulations: *d*(:ATP@O11, :ATP@P3) < 4.00; *d*(:ATP@O6, :DAL@C1) > 2.00; *d*(:ATP@O12, :DAL@C1) > 2.00; *d*(:ATP@O13, :DAL@C1) > 2.00; *d*(:DAL@C1, :DAL@O2) < 3.00, as well as all the P-O bonds in the ATP molecule except for the P3-O11 bond. The metastable states P5, P7, P8, P9, P10, P11 and P12 were taken to evaluate $\Delta G_i^{\ddagger}$. Near-native states P4 and P6 were excluded due to the low probability of being populated in the simulations (< 0.05%).

***DHFR.*** The crystal structure 4KJK was used as the template for back-mapping. The reaction mechanism of DHFR includes a proton transfer step, followed by a hydride transfer[71]. The hydride transfer from NADPH to the protonated dihydrofolate (DHF-H+) is the rate-limiting step[71]. We therefore only studied the hydride transfer step. The binding poses of reactant NADPH (NPH) were searched within the box centered at (4.333, 6.861, -16.028) Å with the dimension 40 Å × 54 Å × 20 Å. The binding poses of reactant DHF-H+ (DFH) were searched within the box centered at (0.861, -3.250, -6.861) Å with the dimension 20 Å × 28 Å × 32 Å. The following distance restraints were used to refine the binding poses with classical MD simulations: *d*(:78@CA, :NPH@N5) = 4.93; *d*(:44@CZ, :NPH@P3) = 4.09; *d*(:98@CZ, :NPH@O13) = 3.18; *d*(:46@OG1, :NPH@O10) = 2.70; *d*(:49@OG, :NPH@O2) = 3.71; *d*(:7@CA, :NPH@N1) = 4.01; *d*(:27@CG, :DFH@N5) = 3.54; *d*(:6@CA, :DFH@N4) = 3.63; *d*(:28@CA, :DFH@C1) = 4.32; *d*(:57@CZ, :DFH@C5) = 3.80; *d*(:DFH@C14, :NPH@C3) = 3.19. The following distance restraints were used to refine the active site with QM/MM energy minimization: *d*(:27@CG, : DFH @N5) = 3.54; *d*(:DFH @C14, :NPH@C3) = 3.19. The QM region (see Supplementary Fig. 4c) includes a part of DHF-H+ with the protonated pterin group, a part of NADPH with the dihydronicotinamide group and the side chain of Asp27 (play an important role in facilitating both the proton transfer and hydride transfer steps[71]). The reaction coordinate (RC) was defined as RC $= d($:NPH@C3$,$ :NPH@H42$) - d($:NPH@H4$,$ :DFH@C14$)$ and discretized into 31 windows: RC = -2.0, -1.8, -1.6, -1.4, -1.2, -1.0, -0.9, -0.8, -0.7, -0.6, -0.5, -0.4, -0.3, -0.2, -0.1, 0.0, 0.1, 0.2, 0.3, 0.4, 0.5, 0.6, 0.7, 0.8, 0.9, 1.0, 1.2, 1.4, 1.6, 1.8, 2.0. The metastable states P2, P3, P4 and P5 were taken to evaluate $\Delta G_i^{\ddagger}$.

##### 15. Error estimation and statistical tests

The 95% CIs of the raw state probabilities $p_i^0(t)$ were estimated using the following bootstrapping method: (1) Randomly resample the set of count matrixes of post-translational trajectories 10,000 times with replacement; (2) Calculate the resampled total count matrix for each variant as the summation of the set of count matrixes of post-translational trajectories, resulting in 10,000 resampled total count matrices for each variant; (3) For each sample, estimate the transition rate matrix using Supplementary equation (18) and the time series of raw state probability $p_i^0(t)$ by numerically solving Supplementary equation (17), resulting 10,000 resampled $p_i^0(t)$ for each state; (4) At a given time $t$, estimate the lower bound and upper bound of 95% CI as the 2.5% and 97.5% percentiles of the distribution of the 10,000 resampled $p_i^0(t)$ for each variant, respectively.

The 95% CIs of the relative specific activities SA$^*$ were estimated using the following bootstrapping method: (1) Create a set of $\Delta G_i^\ddagger$ values ($\{\Delta G_i^\ddagger\}$) that reproduces the observed distribution probabilities $\{p_i\}$ (with solubility correction) for each variant with a sample size equal to the number of post-translational trajectories; (2) Randomly resample the set $\{\Delta G_i^\ddagger\}$ 10,000 times with replacement; (3) For each sample, estimate SA$^*$ using the main text equations (4) and (5), where the state probabilities $p_i$ were calculated as the fraction of state *i* in the resampled $\{\Delta G_i^\ddagger\}$ set, resulting in 10,000 resampled SA$^*$; (5) Estimate the lower bound and upper bound of 95% CI as the 2.5% and 97.5% percentiles of the distribution of the 10,000 resampled SA$^*$ for each variant, respectively.

To test whether or not the observed change of SA$^*$ at 60 s is statistically significant, a p-value was calculated for each candidate enzyme using the permutation test method. The null hypothesis is that $SA_{fast}$ and $SA_{slow}$ do not have significant difference; and the alternative hypothesis is that $SA_{fast}$ is lower than $SA_{slow}$ or $SA_{slow}$ is lower than $SA_{fast}$, depending on which variant has the lower observed SA. The permutation test was performed as follows: (1) Create a set of $\Delta G_i^\ddagger$ values ($\{\Delta G_i^\ddagger\}$) that reproduces the observed distribution probabilities $\{p_i\}$ (with solubility correction) at 60 s for each variant with the sample size equals to the number of post-translational trajectories; (2) Combine $\{\Delta G_i^\ddagger\}_{fast}$ and $\{\Delta G_i^\ddagger\}_{slow}$ into one set and resample the combined set for 1,000,000 times without replacement, i.e., permutate the indices, resulting 1,000,000 resampled combined sets; (3) For each resampled combined set, partition the first half as the resampled $\{\Delta G_i^\ddagger\}_{fast}$ and the second half as the resampled $\{\Delta G_i^\ddagger\}_{slow}$, resulting

1,000,000 resampled $\{\Delta G_i^{\ddagger}\}_{\text{fast}}$ and $\{\Delta G_i^{\ddagger}\}_{\text{slow}}$; (4) For each sample, estimate $SA_{\text{fast}}$ and $SA_{\text{slow}}$, respectively, using the main text equation (4), where the state probabilities $p_i$ were calculated as the fraction of state *i* in the resampled $\{\Delta G_i^{\ddagger}\}_{\text{fast}}$ and $\{\Delta G_i^{\ddagger}\}_{\text{slow}}$ sets. Calculate $SA_{\text{slow}} - SA_{\text{fast}}$ (in the case that the alternative hypothesis is $SA_{\text{fast}} < SA_{\text{slow}}$) or $SA_{\text{fast}} - SA_{\text{slow}}$ (in the case that the alternative hypothesis is $SA_{\text{fast}} > SA_{\text{slow}}$); (5) Estimate the p-value as the frequency of observing the sampled difference that is greater than or equals to the observed difference. If the p-value is less than the significant level 0.05 then we reject the null hypothesis.

##### 16. Folding pathway analysis for co- and post-translational simulations

To analyze the co-translational folding pathways, the synthesis simulations were divided into blocks of nascent chain lengths. Three blocks of nascent chain lengths were used for CAT-III (1-100, 101-203 and 204-213 residues), whereas four and three blocks were used for DDLB (1-90, 91-180,181-296 and 297-306) and DHFR (1-80, 81-149 and 150-159). At each block, 200 clusters were grouped via a *k*-means algorithm[41, 42] based on the order parameters *Q* and *G* that were calculated for all the trajectories of fast and slow variants; and at maximum 3 metastable states were identified via the PCCA+ algorithm[43]. The co-translational discrete trajectories were constructed based on the metastable states assigned in all the blocks. The combined discrete trajectories for co- and post-translational folding were then constructed by appending the post-translational discrete trajectories (obtained in Supplementary Methods Section 12) to the corresponding co-translational discrete trajectories. The folding pathways were identified as follows: (1) For each discrete trajectory, put the starting state of the first frame into the pathway; (2) Move forward along the trajectory and find the state that is different from the last state recoded in the pathway. If the state has not yet been recorded in the pathway, then put it into the pathway. Otherwise, cut the pathway at the first place where this state is recorded and then move forward; (3) Repeat step (2) until the end of the trajectory. This will yield a pathway that has no loop on the route and only records the on-pathway states for each discrete trajectory. The distribution of distinct pathways and the pathway probabilities from one state to another can be estimated from the pathways of all the discrete trajectories.

##### 17. Structural distribution divergence analysis

To quantify the divergence of the co- and post-translational structural distributions between the fast variant and slow variant, the Jensen-Shannon divergence (JSD) metric[72] was applied on the probability distributions ($P_i^{\text{fast}}$ and $P_i^{\text{slow}}$) of the microstates ($M$) assigned by the *k*-means clusters at a given nascent chain length *i* or post-translational time step *i*:

$$\mathrm{JSD}\left(P_i^{\mathrm{fast}} \| P_i^{\mathrm{slow}}\right) = \frac{1}{2}\sum_{x \in M}\left[P_i^{\mathrm{fast}}(x)\ln\frac{P_i^{\mathrm{fast}}(x)}{\frac{1}{2}\left(P_i^{\mathrm{fast}}(x)+P_i^{\mathrm{slow}}(x)\right)} + P_i^{\mathrm{slow}}(x)\ln\frac{P_i^{\mathrm{slow}}(x)}{\frac{1}{2}\left(P_i^{\mathrm{fast}}(x)+P_i^{\mathrm{slow}}(x)\right)}\right] \tag{22}$$

where $P_i^{\mathrm{fast}}$ and $P_i^{\mathrm{slow}}$ were obtained from the last frame of the co-translational simulation trajectories at nascent chain length *i* and from frame *i* of the post-translational simulation trajectories, respectively. For the purpose of better comparison, the co-translational JSD values were normalized by their maximum value, while the post-translational JSD values were normalized to achieve that the starting value is identical to the end value of the co-translational JSD. That is,

$$\begin{cases} \mathrm{JSD}_{\mathrm{norm}}^{\mathrm{co\text{-}trans}}(i) = \dfrac{\mathrm{JSD}^{\mathrm{co\text{-}trans}}(i)}{\max\limits_i\left\{\mathrm{JSD}^{\mathrm{co\text{-}trans}}(i)\right\}} \\ \mathrm{JSD}_{\mathrm{norm}}^{\mathrm{post\text{-}trans}}(i) = \dfrac{\mathrm{JSD}^{\mathrm{post\text{-}trans}}(i)}{\mathrm{JSD}^{\mathrm{post\text{-}trans}}(0)} \cdot \mathrm{JSD}_{\mathrm{norm}}^{\mathrm{co\text{-}trans}}(N) \end{cases}, \tag{23}$$

where *N* is the protein chain length.

#### 18. Experimental validation for the predicted specific activity of DDLB variants

**Materials.** All commercial materials were used as received unless otherwise noted. Tris(hydroxymethyl)aminomethane (Tris), sodium chloride, and glycerol was purchased from Fisher Scientific. Imidazole was purchased from J. T. Baker Chemical Co. Isopropyl β-D-1-thiogalactopyranoside (IPTG) was purchased from Gold Biotechnology. Ni-NTA resin was purchased from Qiagen. 2-Mercaptoethanol, D-alanine, and Pyruvate kinase were purchased from Sigma-Aldrich. Lactate dehydrogenase, and phosphoenol pyruvate (PEP) were purchased from Roche.

**General methods.** UV-visible spectra were recorded on a Cary 60 spectrometer from Varian (Agilent Technologies, Santa Clara, CA) using the WinUV software package to control the instrument.

**Cloning, overexpression and purification of slow and fast translational variants of DDLB.** DNA sequences encoding the slow and fast variants of DDLB were obtained from our simulations and cloned into pET-26b(+) vector (New England Biolabs) containing C-terminal hexahistidine ($His_6$) tag using *Nde*I and *Sac*I restriction sites. The resulting constructs, pET-26b(+)-slow and pET-26(+)-fast DDLB, were verified by DNA sequencing and used to transform *E. coli* BL21 (DE3). A single colony was used to inoculate 200 ml of a lysogeny broth (LB) starter culture containing 50 mg/L kanamycin, which was shaken at 37 °C and 180 rpm for 12 h.

15 mL of this starter culture was used to inoculate 3 L of LB medium containing 50 mg/L kanamycin, which was incubated at 37 °C and 180 rpm until an optical density at 600 nm ($OD_{600}$) of ~0.6 was reached. Protein expression was induced by adding 0.25 mM isopropylthio-β-D-galactoside (IPTG), and incubation was continued at 25 °C at 180 rpm for an additional 12-15 h. The cells were harvested by centrifugation (4 °C, 15000 × *g*, 15 min). For protein purification, the cells were resuspended in lysis buffer (20 mM Tris-HCl, 200 mM NaCl, 5 mM imidazole, 10 mM $MgCl_2$, 2 mM β-mercaptoethanol pH 7.5), and were then disrupted by sonication with an ultrasonic cell disruptor (Branson Sonifier II "Modell W- 250", Heinemann) before the lysates were clarified by centrifugation (4 °C, 45000 × *g*, 45 min). The C-terminally $His_6$-tagged slow and fast DDLB variants were purified via immobilized metal affinity chromatography at 4 °C using Ni-NTA resin. The Ni-NTA resin was pre-equilibrated with lysis buffer. After loading the supernatant onto the Ni-NTA column, the resin was washed with 200 mL wash buffer-A (20 mM Tris-HCl, 200 mM NaCl, 20 mM imidazole, 5 mM $MgCl_2$, 2 mM β-mercaptoethanol pH 7.5) followed by a 50 mL wash with wash buffer-B (20 mM Tris-HCl, 200 mM NaCl, 40 mM imidazole, 5 mM $MgCl_2$, 2 mM β-mercaptoethanol pH 7.5) and a 50 mL wash with wash buffer-C (20 mM Tris-HCl, 200 mM NaCl, 80 mM imidazole, 5 mM $MgCl_2$, 2 mM β-mercaptoethanol pH 7.5). The slow and fast variant proteins were then eluted from the column with 30 mL of elution buffer (20 mM Tris-HCl, 200 mM NaCl, 300 mM imidazole, 2 mM β-mercaptoethanol pH 7.5). The proteins were concentrated using 10000 kDa MWCO filters (Millipore), and imidazole was removed using a PD-10 column (GE Healthcare) pre-equilibrated in storage buffer (20 mM Tris-HCl, 200 mM NaCl, 10% glycerol, 2 mM β-mercaptoethanol pH 7.5) according to the manufacturer's protocol. The enzyme was divided into 50 µL aliquots before it was snap-frozen in liquid $N_2$ and stored at −80 °C until use.

**DDLB kinetic assays.** Enzyme activity was measured spectrophotometrically using a previously described procedure with a few amendments[73, 74, 75]. Reactions contained the following in a final volume of 120 µL: 100 mM Tris–HCl (pH 7.8), 20 mM ATP, 10 mM $MgCl_2$, 10 mM KCl, 0.15 mg/mL lactate dehydrogenase, 0.3 mg/mL pyruvate kinase, 0.3 mM NADH, 7.5 mM phosphoenol pyruvate (PEP), and ~13-14 nM of DDLB with varying concentrations (0 – 8 mM) of D-alanine. Reactions were initiated by addition of D- alanine, and the disappearance of NADH was monitored over 5 min at 25 °C by the loss in absorbance at 340 nm ($\varepsilon_{340}$=6440 $M^{-1}$ $cm^{-1}$). Kinetic assays were conducted in triplicate (technical replicates) and the data were fitted to the Michaelis-Menten equation to extract the values of rate constant ($k_{cat}$) and Michaelis constant ($K_M$).

**Relative specific activity and hypothesis test.** In total 5 biological replicates were prepared and assayed for both the fast and slow variants. For each biological replicate, the relative specific activity of the slow variant with regard to the fast variant was calculated as $k_{\mathrm{cat}}^{\mathrm{slow}}/k_{\mathrm{cat}}^{\mathrm{fast}}$. The hypothesis that the average relative specific activity ($\langle k_{\mathrm{cat}}^{\mathrm{slow}}/k_{\mathrm{cat}}^{\mathrm{fast}}\rangle$) is less than 1 was tested by using the one-tailed Student's t-test.

**Time-course analysis of overexpression.** A single colony was used to inoculate 200 ml of a lysogeny broth (LB) starter culture containing 50 mg/L kanamycin, which was shaken at 37 °C and 180 rpm for 12 h. 1 mL of this starter culture was used to inoculate 100 mL of LB medium containing 50 mg/L kanamycin, which was incubated at 37 °C and 180 rpm until an optical density at 600 nm ($OD_{600}$) of ~0.6 was reached. Protein expression was induced by adding 0.25 mM isopropylthio-β-D-galactoside (IPTG), and 5 mL aliquots were collected at time points 0.5h, 1h, 2h, 3h, 4h, 5h after IPTG addition and the cells were harvested by centrifugation (4 °C, 4000 × *g*, 15 min). As a control, 5 mL of cells were also collected before IPTG addition. The aliquots were diluted accordingly based on the $OD_{600}$ so that the number of cells is controlled to be same. Protein expression at all time points including control was analyzed by both SDS-PAGE and western blot as the DDLB variants are $His_6$-tagged and the antibodies binds the $His_6$-tag are commercially available and were used to check the expression of both slow and fast variants.

##### 19. mRNA sequences used in this study

All the mRNA sequences used in the simulations and experiments are presented below. All sequences begin with the start codon and end with the stop codon.

**CAT-III fast variant:**

AUGAAUUAUACUAAAUUUGAUGUUAAAAAUUGGGUUCGUCGUGAGCACUUUGAGUUUUAUCGUCACCGUC
UGCCGUGUGGUUUUUCUCUGACUUCUAAAAUUGAUAUUACUACUCUGAAAAAAUCUCUGGAUGAUUCUGC
UUAUAAAUUUUAUCCGGUUAUGAUUUAUCUGAUUGCUCAAGCUGUUAAUCAAUUUGAUGAGCUGCGUAUG
GCUAUUAAAGAUGAUGAGCUGAUUGUUUGGGAUUCUGUUGAUCCGCAAUUUACUGUUUUUCACCAAGAGA
CUGAGACUUUUUCUGCUCUGUCUUGUCCGUAUUCUUCUGAUAUUGAUCAAUUUAUGGUUAAUUAUCUGUC
UGUUAUGGAGCGUUAUAAAUCUGAUACUAAACUGUUUCCGCAAGGUGUUACUCCGGAGAAUCACCUGAAU
AUUUCUGCUCUGCCGUGGGUUAAUUUUGAUUCUUUUAAUCUGAAUGUUGCUAAUUUUACUGAUUAUUUUG
CUCCGAUUAUUACUAUGGCUAAAUAUCAACAAGAGGGUGAUCGUCUGCUGCUGCCGCUGUCUGUUCAAGU
UCACCACGCUGUUUGUGAUGGUUUUCACGUUGCUCGUUUUAUUAAUCGUCUGCAAGAGCUGUGUAAUUCU
AAACUGAAAUAA

**CAT-III slow variant:**

AUGAACUACACAAAGUUCGACGUCAAGAACUGGGUCAGGAGGGAACAUUUCGAAUUCUACAGGCAUAGGC
UACCAUGCGGAUUCUCCCUAACAUCCAAGAUCGACAUCACAACACUAAAGAAGUCCCUAGACGACUCCGC
CUACAAGUUCUACCCAGUCAUGAUCUACCUAAUCGCCCAGGCCGUCAACCAGUUCGACGAACUAAGGAUG
GCCAUCAAGGACGACGAACUAAUCGUCUGGGACUCCGUCGACCCACAGUUCACAGUCUUCCAUCAGGAAA
CAGAAACAUUCUCCGCCCUAUCCUGCCCAUACUCCUCCGACAUCGACCAGUUCAUGGUCAACUACCUAUC
CGUCAUGGAAAGGUACAAGUCCGACACAAAGCUAUUCCCACAGGGAGUCACACCAGAAAACCAUCUAAAC
AUCUCCGCCCUACCAUGGGUCAACUUCGACUCCUUCAACCUAAACGUCGCCAACUUCACAGACUACUUCG
CCCCAAUCAUCACAAUGGCCAAGUACCAGCAGGAAGGAGACAGGCUACUACUACCACUAUCCGUCCAGGU
CCAUCAUGCCGUCUGCGACGGAUUCCAUGUCGCCAGGUUCAUCAACAGGCUACAGGAACUAUGCAACUCC
AAGCUAAAGUAG

**DDLB fast variant:**

AUGACUGAUAAAAUUGCUGUUCUGCUGGGUGGUACUUCUGCUGAGCGUGAGGUUUCUCUGAAUUCUGGUG
CUGCUGUUCUGGCUGGUCUGCGUGAGGGUGGUAUUGAUGCUUAUCCGGUUGAUCCGAAAGAGGUUGAUGU
UACUCAACUGAAAUCUAUGGGUUUUCAAAAAGUUUUUAUUGCUCUGCACGGUCGUGGUGGUGAGGAUGGU
ACUCUGCAAGGUAUGCUGGAGCUGAUGGGUCUGCCGUAUACUGGUUCUGGUGUUAUGGCUUCUGCUCUGU
CUAUGGAUAAACUGCGUUCUAAACUGCUGUGGCAAGGUGCUGGUCUGCCGGUUGCUCCGUGGGUUGCUCU
GACUCGUGCUGAGUUUGAGAAAGGUCUGUCUGAUAAACAACUGGCUGAGAUUUCUGCUCUGGGUCUGCCG
GUUAUUGUUAAACCGUCUCGUGAGGGUUCUUCUGUUGGUAUGUCUAAAGUUGUUGCUGAGAAUGCUCUGC
AAGAUGCUCUGCGUCUGGCUUUUCAACACGAUGAGGAGGUUCUGAUUGAGAAAUGGCUGUCUGGUCCGGA
GUUUACUGUUGCUAUUCUGGGUGAGGAGAUUCUGCCGUCUAUUCGUAUUCAACCGUCUGGUACUUUUUAU
GAUUAUGAGGCUAAAUAUCUGUCUGAUGAGACUCAAUAUUUUUGUCCGGCUGGUCUGGAGGCUUCUCAAG
AGGCUAAUCUGCAAGCUCUGGUUCUGAAAGCUUGGACUACUCUGGGUUGUAAAGGUUGGGGUCGUAUUGA
UGUUAUGCUGGAUUCUGAUGGUCAAUUUUAUCUGCUGGAGGCUAAUACUUCUCCGGGUAUGACUUCUCAC
UCUCUGGUUCCGAUGGCUGCUCGUCAAGCUGGUAUGUCUUUUUCUCAACUGGUUGUUCGUAUUCUGGAGC
UGGCUGAUUAA

**DDLB slow variant:**

AUGACAGACAAGAUCGCCGUCCUACUAGGAGGAACAUCCGCCGAAAGGGAAGUCUCCCUAAACUCCGGAG
CCGCCGUCCUAGCCGGACUAAGGGAAGGAGGAAUCGACGCCUACCCAGUCGACCCAAAGGAAGUCGACGU
CACACAGCUAAAGUCCAUGGGAUUCCAGAAGGUCUUCAUCGCCCUACAUGGAAGGGGAGGAGAAGACGGA
ACACUACAGGGAAUGCUAGAACUAAUGGGACUACCAUACACAGGAUCCGGAGUCAUGGCCUCCGCCCUAU
CCAUGGACAAGCUAAGGUCCAAGCUACUAUGGCAGGGAGCCGGACUACCAGUCGCCCCAUGGGUCGCCCU
AACAAGGGCCGAAUUCGAAAAGGGACUAUCCGACAAGCAGCUAGCCGAAAUCUCCGCCCUAGGACUACCA
GUCAUCGUCAAGCCAUCCAGGGAAGGAUCCUCCGUCGGAAUGUCCAAGGUCGUCGCCGAAAACGCCCUAC

AGGACGCCCUAAGGCUAGCCUUCCAGCAUGACGAAGAAGUCCUAAUCGAAAAGUGGCUAUCCGGACCAGA
AUUCACAGUCGCCAUCCUAGGAGAAGAAAUCCUACCAUCCAUCAGGAUCCAGCCAUCCGGAACAUUCUAC
GACUACGAAGCCAAGUACCUAUCCGACGAAACACAGUACUUCUGCCCAGCCGGACUAGAAGCCUCCCAGG
AAGCCAACCUACAGGCCCUAGUCCUAAAGGCCUGGACAACACUAGGAUGCAAGGGAUGGGGAAGGAUCGA
CGUCAUGCUAGACUCCGACGGACAGUUCUACCUACUAGAAGCCAACACAUCCCCAGGAAUGACAUCCCAU
UCCCUAGUCCCAAUGGCCGCCAGGCAGGCCGGAAUGUCCUUCUCCCAGCUAGUCGUCAGGAUCCUAGAAC
UAGCCGACUAG

**DHFR fast variant:**

AUGAUUUCUCUGAUUGCUGCUCUGGCUGUUGAUCGUGUUAUUGGUAUGGAGAAUGCUAUGCCGUGGAAUC
UGCCGGCUGAUCUGGCUUGGUUUAAACGUAAUACUCUGAAUAAACCGGUUAUUAUGGGUCGUCACACUUG
GGAGUCUAUUGGUCGUCCGCUGCCGGGUCGUAAAAAUAUUAUUCUGUCUUCUCAACCGGGUACUGAUGAU
CGUGUUACUUGGGUUAAAUCUGUUGAUGAGGCUAUUGCUGCUUGUGGUGAUGUUCCGGAGAUUAUGGUUA
UUGGUGGUGGUCGUGUUUAUGAGCAAUUUCUGCCGAAAGCUCAAAAACUGUAUCUGACUCACAUUGAUGC
UGAGGUUGAGGGUGAUACUCACUUUCCGGAUUAUGAGCCGGAUGAUUGGGAGUCUGUUUUUUCUGAGUUU
CACGAUGCUGAUGCUCAAAAUUCUCACUCUUAUUGUUUUGAGAUUCUGGAGCGUCGUUAA

**DHFR slow variant:**

AUGAUCUCCCUAAUCGCCGCCCUAGCCGUCGACAGGGUCAUCGGAAUGGAAAACGCCAUGCCAUGGAACC
UACCAGCCGACCUAGCCUGGUUCAAGAGGAACACACUAAACAAGCCAGUCAUCAUGGGAAGGCAUACAUG
GGAAUCCAUCGGAAGGCCACUACCAGGAAGGAAGAACAUCAUCCUAUCCUCCCAGCCAGGAACAGACGAC
AGGGUCACAUGGGUCAAGUCCGUCGACGAAGCCAUCGCCGCCUGCGGAGACGUCCCAGAAAUCAUGGUCA
UCGGAGGAGGAAGGGUCUACGAACAGUUCCUACCAAAGGCCCAGAAGCUAUACCUAACACAUAUCGACGC
CGAAGUCGAAGGAGACACACAUUUCCCAGACUACGAACCAGACGACUGGGAAUCCGUCUUCUCCGAAUUC
CAUGACGCCGACGCCCAGAACUCCCAUUCCUACUGCUUCGAAAUCCUAGAAAGGAGGUAG

#### Supplementary Tables

Supplementary Table 1. Force field parameters of the CG protein model

| Energy term | | Parameters |
|---|---|---|
| $E_{\text{bond}}$ | $K_{\text{b}}$ (kcal/mol/Å$^2$) | 50 |
| | $b_0$ (Å) | 3.81 |
| $E_{\text{angle}}$ | $\gamma$ (mol/kcal) | 0.1 |
| | $K_{\alpha}$ (kcal/mol/rad$^2$) | 106.4 |
| | $\theta_{\alpha}$ (rad) | 1.60 |
| | $\varepsilon_{\alpha}$ (kcal/mol) | 4.3 |
| | $K_{\beta}$ (kcal/mol/rad$^2$) | 26.3 |
| | $\theta_{\beta}$ (rad) | 2.27 |
| $E_{\text{dihedral}}$ | $K_{\text{D}_1}$ (kcal/mol/rad$^2$) | 0.6309 |
| | $\delta_1$ (rad) | 5.3291 |
| | $K_{\text{D}_2}$ (kcal/mol/rad$^2$) | 1.0621 |
| | $\delta_2$ (rad) | 4.5574 |
| | $K_{\text{D}_3}$ (kcal/mol/rad$^2$) | 0.045943 |
| | $\delta_3$ (rad) | 4.8229 |
| | $K_{\text{D}_4}$ (kcal/mol/rad$^2$) | 0.076257 |
| | $\delta_4$ (rad) | 2.1 |
| $E_{\text{elec}}$ | $q_i$ ($e$) | Amino acid dependent |
| | $\varepsilon_{\gamma}$ | 78.5 |
| | $l_{\text{D}}$ (Å) | 10 |
| $E_{\text{vdW}}$ | Native contacts [a] | |
| | $\varepsilon_{ij}$ (kcal/mol) | $\varepsilon_{ij}^{\text{HB}} + n_{\text{scal}} \cdot \varepsilon_{ij}^{\text{SC-SC}} + \varepsilon_{ij}^{\text{BB-SC}}$ |
| | $R_{ij}$ (Å) | $d_{ij}$ |
| | Non-native contacts [b] | |
| | $\varepsilon_{ij}$ (kcal/mol) | 0.000132 |
| | $R_{ij}$ (Å) | $\frac{2^{1/6}}{2}(\sigma_i + \sigma_j)$ |

[a] Native contacts include three terms: hydrogen bond (HB), sidechain-sidechain (SC-SC) and backbone-sidechain (BB-SC). The potential scaling factor $n_{\text{scal}}$ need to be further parameterized for a particular protein. $d_{ij}$ is the native contact distance.

[b] Non-native contacts have a fixed potential well depth. $\sigma_i$ is the collision diameter of the CG interaction site *i*.

Supplementary Table 2a. The protein name, PDB ID, protein length, experimental temperature $T^{\mathrm{exp}}$, experimental protein stability $\Delta G_{\mathrm{UN}}^{\mathrm{exp}}$, $n_{\mathrm{scal}}$ values used in pt-REMD, melting temperature $T_{\mathrm{m}}$, threshold used to identify folding states $Q_{\mathrm{eq}}$, estimated protein stability $\Delta G_{\mathrm{UN}}^{\mathrm{sim}}(T^{\mathrm{exp}})$, linear regression slope *m*, intercept *b*, coefficient of determination $R^2$ and optimized $n_{\mathrm{scal}}^{*}$ of α-class training-set proteins

| Protein | PDB ID | Length | Experimental parameters | | pt-REMD input and output parameters | | | | Linear regression parameters | | | |
|---|---|---|---|---|---|---|---|---|---|---|---|---|
| | | | $T^{\mathrm{exp}}$ (K) | $\Delta G_{\mathrm{UN}}^{\mathrm{exp}}$ (kcal/mol) | $n_{\mathrm{scal}}$ | $T_{\mathrm{m}}$ (K) | $Q_{\mathrm{eq}}$ | $\Delta G_{\mathrm{UN}}^{\mathrm{sim}}(T^{\mathrm{exp}})$ (kcal/mol) | *m* | *b* | $R^2$ | $n_{\mathrm{scal}}^{*}$ |
| EC298 | 1RYK | 69 | 298 | -2.72 [b] | 1.1 | 315.0 | 0.68 | -2.807 | -19.848 | 18.839 | 0.9952 | 1.0862 |
| | | | | | 1.2 | 331.1 | 0.66 | -5.119 | | | | |
| | | | | | 1.3 | 346.9 | 0.65 | -7.248 | | | | |
| | | | | | 1.4 | 361.5 | 0.64 | -8.713 | | | | |
| λ-repressor$_{6-85}$ | 1LMB | 80 | 298 | -4.25 [b] | 1.2 | 314.5 | 0.53 | -3.941 | -20.555 | 20.797 | 0.9995 | 1.2185 |
| | | | | | 1.3 | 330.0 | 0.54 | -5.802 | | | | |
| | | | | | 1.4 | 345.0 | 0.51 | -8.010 | | | | |
| | | | | | 1.5 | 359.8 | 0.50 | -10.057 | | | | |
| bACBP [a] | 2ABD | 86 | 298 | -6.40 [b] | 1.2 | 313.2 | 0.69 | -3.711 | -17.553 | 17.325 | 0.9970 | 1.3516 |
| | | | | | 1.3 | 327.3 | 0.69 | -5.406 | | | | |
| | | | | | 1.4 | 340.3 | 0.65 | -7.505 | | | | |
| | | | | | 1.5 | 354.8 | 0.67 | -8.863 | | | | |
| IM7 | 1CEI | 85 | 298 | -2.88 [b] | 1.0 | 305.8 | 0.62 | -1.809 | -16.987 | 14.735 | 0.9745 | 1.0369 |
| | | | | | 1.1 | 325.3 | 0.60 | -4.677 | | | | |
| | | | | | 1.2 | 342.8 | 0.60 | -5.528 | | | | |
| | | | | | 1.3 | 361.4 | 0.58 | -7.188 | | | | |
| Im9 | 1IMQ | 86 | 298 | -6.81 [c] | 1.1 | 309.7 | 0.53 | -2.458 | -18.194 | 17.111 | 0.9806 | 1.3147 |
| | | | | | 1.2 | 326.4 | 0.51 | -5.300 | | | | |
| | | | | | 1.3 | 340.6 | 0.51 | -6.723 | | | | |
| | | | | | 1.4 | 357.2 | 0.50 | -8.048 | | | | |
| Cytochrome B562 [a] | 256B | 106 | 293 | -6.60 [c] | 1.1 | 310.6 | 0.58 | -5.008 | -27.403 | 25.313 | 0.9982 | 1.1646 |
| | | | | | 1.2 | 322.8 | 0.59 | -7.402 | | | | |
| | | | | | 1.3 | 335.5 | 0.58 | -10.116 | | | | |
| | | | | | 1.4 | 347.5 | 0.57 | -13.238 | | | | |
| Average | | | | | | | | | | | | 1.1954 |

[a] Insufficiently sampled proteins whose pt-REMD results are the averages of those obtained from pt-REMD simulations starting from their native states and unfolded states. See details in Supplementary Table 3.

[b] Obtained from De Sancho et al.[11];

[c] Obtained from Leininger et al.[5];

Supplementary Table 2b. The protein name, PDB ID, protein length, experimental temperature $T^{\mathrm{exp}}$, experimental protein stability $\Delta G_{\mathrm{UN}}^{\mathrm{exp}}$, $n_{\mathrm{scal}}$ values used in pt-REMD, melting temperature $T_{\mathrm{m}}$, threshold used to identify folding states $Q_{\mathrm{eq}}$, estimated protein stability $\Delta G_{\mathrm{UN}}^{\mathrm{sim}}(T^{\mathrm{exp}})$, linear regression slope $m$, intercept $b$, coefficient of determination $R^2$ and optimized $n_{\mathrm{scal}}^{*}$ of β-class training-set proteins

| Protein | PDB ID | Length | Experimental parameters | | pt-REMD input and output parameters | | | | Linear regression parameters | | | |
|---|---|---|---|---|---|---|---|---|---|---|---|---|
| | | | $T^{\mathrm{exp}}$ (K) | $\Delta G_{\mathrm{UN}}^{\mathrm{exp}}$ (kcal/mol) | $n_{\mathrm{scal}}$ | $T_{\mathrm{m}}$ (K) | $Q_{\mathrm{eq}}$ | $\Delta G_{\mathrm{UN}}^{\mathrm{sim}}(T^{\mathrm{exp}})$ (kcal/mol) | $m$ | $b$ | $R^2$ | $n_{\mathrm{scal}}^{*}$ |
| ABP1 SH3 | 1JO8 | 58 | 298 | -3.07 [a] | 1.4 | 320.2 | 0.77 | -2.042 | -3.907 | 3.381 | 0.9952 | 1.6514 |
| | | | | | 1.5 | 334.6 | 0.78 | -2.535 | | | | |
| | | | | | 1.6 | 348.3 | 0.78 | -2.895 | | | | |
| | | | | | 1.7 | 362.7 | 0.78 | -3.224 | | | | |
| Fyn SH3 | 1SHF | 59 | 293 | -6.68 [a] | 1.2 | 307.6 | 0.54 | -2.888 | -12.596 | 12.167 | 0.9987 | 1.4962 |
| | | | | | 1.3 | 320.4 | 0.55 | -4.327 | | | | |
| | | | | | 1.4 | 335.4 | 0.55 | -5.410 | | | | |
| | | | | | 1.5 | 351.0 | 0.52 | -6.726 | | | | |
| CspB Bc | 1C9O | 66 | 298 | -1.81 [b] | 1.3 | 304.8 | 0.51 | -0.949 | -16.429 | 20.274 | 0.9969 | 1.3442 |
| | | | | | 1.4 | 319.6 | 0.49 | -2.8442 | | | | |
| | | | | | 1.5 | 334.7 | 0.49 | -4.5401 | | | | |
| | | | | | 1.6 | 349.0 | 0.49 | -5.8598 | | | | |
| CspA | 1MJC | 69 | 298 | -3.00 [c] | 1.4 | 307.9 | 0.50 | -1.6971 | -16.644 | 21.483 | 0.9980 | 1.4709 |
| | | | | | 1.5 | 320.8 | 0.51 | -3.6163 | | | | |
| | | | | | 1.6 | 334.2 | 0.50 | -5.2519 | | | | |
| | | | | | 1.7 | 347.3 | 0.49 | -6.7000 | | | | |
| Tenascin | 1TEN | 89 | 298 | -6.71 [a] | 1.3 | 307.4 | 0.41 | -1.6397 | -29.161 | 36.208 | 0.9999 | 1.4718 |
| | | | | | 1.4 | 323.9 | 0.39 | -4.7050 | | | | |
| | | | | | 1.5 | 338.7 | 0.39 | -7.5436 | | | | |
| | | | | | 1.6 | 354.4 | 0.39 | -10.4139 | | | | |
| Twitchin | 1WIT | 93 | 293 | -5.00 [a] | 1.3 | 310.1 | 0.46 | -3.0763 | -18.441 | 20.899 | 0.9995 | 1.4044 |
| | | | | | 1.4 | 324.6 | 0.46 | -4.9700 | | | | |
| | | | | | 1.5 | 338.7 | 0.45 | -6.6515 | | | | |
| | | | | | 1.6 | 353.3 | 0.44 | -8.6626 | | | | |
| Average | | | | | | | | | | | | 1.4732 |

[a] Obtained from De Sancho et al.[11];

[b] Obtained from Leininger et al.[5];

[c] Obtained from Reid et al.[12];

Supplementary Table 2c. The protein name, PDB ID, protein length, experimental temperature $T^{\mathrm{exp}}$, experimental protein stability $\Delta G_{\mathrm{UN}}^{\mathrm{exp}}$, $n_{\mathrm{scal}}$ values used in pt-REMD, melting temperature $T_{\mathrm{m}}$, threshold used to identify folding states $Q_{\mathrm{eq}}$, estimated protein stability $\Delta G_{\mathrm{UN}}^{\mathrm{sim}}(T^{\mathrm{exp}})$, linear regression slope *m*, intercept *b*, coefficient of determination $R^2$ and optimized $n_{\mathrm{scal}}^{*}$ of α/β-class training-set proteins

| Protein | PDB ID | Length | Experimental parameters | | pt-REMD input and output parameters | | | | Linear regression parameters | | | |
|---|---|---|---|---|---|---|---|---|---|---|---|---|
| | | | $T^{\mathrm{exp}}$ (K) | $\Delta G_{\mathrm{UN}}^{\mathrm{exp}}$ (kcal/mol) | $n_{\mathrm{scal}}$ | $T_{\mathrm{m}}$ (K) | $Q_{\mathrm{eq}}$ | $\Delta G_{\mathrm{UN}}^{\mathrm{sim}}(T^{\mathrm{exp}})$ (kcal/mol) | *m* | *b* | $R^2$ | $n_{\mathrm{scal}}^{*}$ |
| Hpr [a] | 1POH | 85 | 298 | -5.46 [b] | 1.0 | 297.3 | 0.51 | 0.165 | -26.153 | 25.765 | 0.9854 | 1.1940 |
| | | | | | 1.1 | 313.1 | 0.50 | -3.709 | | | | |
| | | | | | 1.2 | 327.6 | 0.50 | -5.864 | | | | |
| | | | | | 1.3 | 342.4 | 0.50 | -7.834 | | | | |
| Urm1 [a] | 2QJL | 99 | 298 | -3.48 [c] | 1.1 | 307.9 | 0.46 | -2.786 | -25.491 | 24.723 | 0.9854 | 1.1064 |
| | | | | | 1.2 | 325.8 | 0.46 | -6.506 | | | | |
| | | | | | 1.3 | 342.9 | 0.47 | -8.729 | | | | |
| | | | | | 1.4 | 359.6 | 0.48 | -10.543 | | | | |
| Src SH2 [a] | 1SPR | 103 | 298 | -7.24 [c] | 1.1 | 307.6 | 0.73 | -2.783 | -16.597 | 15.283 | 0.9910 | 1.3571 |
| | | | | | 1.2 | 324.6 | 0.74 | -4.740 | | | | |
| | | | | | 1.3 | 342.4 | 0.71 | -6.651 | | | | |
| | | | | | 1.4 | 359.0 | 0.71 | -7.678 | | | | |
| Azurin (apo) [a] | 1E65 | 128 | 298 | -5.29 [c] | 1.1 | 306.8 | 0.70 | -3.489 | -19.738 | 17.942 | 0.9795 | 1.1770 |
| | | | | | 1.2 | 321.6 | 0.66 | -6.424 | | | | |
| | | | | | 1.3 | 338.4 | 0.69 | -7.199 | | | | |
| | | | | | 1.4 | 355.6 | 0.65 | -9.810 | | | | |
| CheY [a] | 3CHY | 128 | 298 | -5.20 [d] | 1.0 | 309.0 | 0.50 | -3.906 | -64.410 | 60.542 | 0.9999 | 1.0207 |
| | | | | | 1.1 | 328.1 | 0.45 | -10.233 | | | | |
| | | | | | 1.2 | 347.2 | 0.43 | -16.788 | | | | |
| | | | | | 1.3 | 365.6 | 0.43 | $-\infty$ [f] | | | | |
| Ribonuclease H [a] | 2RN2 | 155 | 298 | -7.20 [e] | 1.0 | 310.5 | 0.23 | -3.924 | -42.080 | 38.167 | 0.9999 | 1.0781 |
| | | | | | 1.1 | 327.7 | 0.23 | -8.150 | | | | |
| | | | | | 1.2 | 344.0 | 0.22 | -12.230 | | | | |
| | | | | | 1.3 | 361.1 | 0.22 | -16.591 | | | | |
| Average | | | | | | | | | | | | 1.1556 |

[a] Insufficiently sampled proteins whose pt-REMD results are the averages of those obtained from pt-REMD simulations starting from their native states and unfolded states. See details in Table 3;

[b] Obtained from Nicholson et al.[13];

[c] Obtained from De Sancho et al.[11];

[d] Obtained from López-Hernéndez & Serrano[14];

[e] Obtained from Leininger et al.[5];

[f] Infinite $\Delta G_{\mathrm{UN}}^{\mathrm{sim}}$ due to the extremely high stability at this $n_{\mathrm{scal}}$ value. It was considered as an outlier and not used in the linear regression.

Supplementary Table 3. Insufficiently sampled training-set proteins and their pt-REMD results

| Protein | PDB ID | REX input and output parameters | | | | | Average | | |
|---|---|---|---|---|---|---|---|---|---|
| | | $n_{\mathrm{scal}}$ | Starting state | $T_{\mathrm{m}}$ (K) | $Q_{\mathrm{eq}}$ | $\Delta G_{\mathrm{UN}}^{\mathrm{sim}}(T^{\mathrm{exp}})$ (kcal/mol) | $T_{\mathrm{m}}$ (K) | $Q_{\mathrm{eq}}$ | $\Delta G_{\mathrm{UN}}^{\mathrm{sim}}(T^{\mathrm{exp}})$ (kcal/mol) |
| bACBP | 2ABD | 1.2 | folded | 314.0 | 0.68 | -3.902 | 313.2 | 0.69 | -3.711 |
| | | | unfolded | 312.3 | 0.70 | -3.521 | | | |
| | | 1.3 | folded | 327.2 | 0.70 | -5.163 | 327.3 | 0.69 | -5.406 |
| | | | unfolded | 327.4 | 0.67 | -5.650 | | | |
| | | 1.4 | folded | 340.9 | 0.67 | -7.368 | 340.3 | 0.65 | -7.505 |
| | | | unfolded | 339.6 | 0.64 | -7.641 | | | |
| | | 1.5 | folded | 355.3 | 0.67 | -8.865 | 354.8 | 0.67 | -8.863 |
| | | | unfolded | 354.3 | 0.67 | -8.861 | | | |
| Cytochrome B562 | 256B | 1.1 | folded | 311.1 | 0.58 | -5.131 | 310.6 | 0.58 | -5.008 |
| | | | unfolded | 310.0 | 0.59 | -4.885 | | | |
| | | 1.2 | folded | 323.4 | 0.59 | -7.521 | 322.8 | 0.59 | -7.402 |
| | | | unfolded | 322.1 | 0.59 | -7.283 | | | |
| | | 1.3 | folded | 335.5 | 0.58 | -10.011 | 335.5 | 0.58 | -10.116 |
| | | | unfolded | 335.4 | 0.57 | -10.222 | | | |
| | | 1.4 | folded | 347.4 | 0.57 | -13.067 | 347.5 | 0.57 | -13.238 |
| | | | unfolded | 347.5 | 0.57 | -13.408 | | | |
| Hpr | 1POH | 1.0 | folded | 297.3 | 0.51 | 0.167 | 297.3 | 0.51 | 0.165 |
| | | | unfolded | 297.3 | 0.51 | 0.163 | | | |
| | | 1.1 | folded | 313.1 | 0.50 | -3.682 | 313.1 | 0.50 | -3.709 |
| | | | unfolded | 313.0 | 0.50 | -3.735 | | | |
| | | 1.2 | folded | 328.0 | 0.50 | -5.992 | 327.6 | 0.50 | -5.864 |
| | | | unfolded | 327.2 | 0.50 | -5.736 | | | |
| | | 1.3 | folded | 342.7 | 0.49 | -8.068 | 342.4 | 0.50 | -7.834 |
| | | | unfolded | 342.0 | 0.50 | -7.600 | | | |
| Urm1 | 2QJL | 1.1 | folded | 307.9 | 0.49 | -2.785 | 307.9 | 0.46 | -2.786 |
| | | | unfolded | 307.8 | 0.44 | -2.788 | | | |
| | | 1.2 | folded | 326.2 | 0.46 | -6.497 | 325.8 | 0.46 | -6.506 |
| | | | unfolded | 325.4 | 0.47 | -6.515 | | | |
| | | 1.3 | folded | 342.5 | 0.48 | -8.519 | 342.9 | 0.47 | -8.729 |
| | | | unfolded | 343.2 | 0.46 | -8.939 | | | |
| | | 1.4 | folded | 359.4 | 0.49 | -10.338 | 359.6 | 0.48 | -10.543 |
| | | | unfolded | 359.7 | 0.48 | -10.747 | | | |
| Src SH2 | 1SPR | 1.1 | folded | 307.5 | 0.74 | -2.676 | 307.6 | 0.73 | -2.783 |
| | | | unfolded | 307.6 | 0.72 | -2.890 | | | |
| | | 1.2 | folded | 325.0 | 0.73 | -4.752 | 324.6 | 0.74 | -4.740 |
| | | | unfolded | 324.1 | 0.74 | -4.728 | | | |
| | | 1.3 | folded | 342.0 | 0.71 | -6.686 | 342.4 | 0.71 | -6.651 |
| | | | unfolded | 342.8 | 0.71 | -6.617 | | | |
| | | 1.4 | folded | 358.8 | 0.72 | -7.635 | 359.0 | 0.71 | -7.678 |
| | | | unfolded | 359.2 | 0.71 | -7.721 | | | |

| | | | | | | | | | |
|---|---|---|---|---|---|---|---|---|---|
| Azurin (apo) | 1E65 | 1.1 | folded | 310.0 | 0.68 | -4.109 | 306.8 | 0.70 | -3.489 |
| | | | unfolded | 303.6 | 0.71 | -2.869 | | | |
| | | 1.2 | folded | 325.0 | 0.68 | -6.002 | 321.6 | 0.66 | -6.424 |
| | | | unfolded | 318.2 | 0.65 | -6.846 | | | |
| | | 1.3 | folded | 341.1 | 0.67 | -7.754 | 338.4 | 0.69 | -7.199 |
| | | | unfolded | 335.6 | 0.70 | -6.643 | | | |
| | | 1.4 | folded | 355.9 | 0.64 | -10.361 | 355.6 | 0.65 | -9.810 |
| | | | unfolded | 355.3 | 0.67 | -9.259 | | | |
| CheY | 3CHY | 1.0 | folded | 308.9 | 0.50 | -3.920 | 309.0 | 0.50 | -3.906 |
| | | | unfolded | 309.0 | 0.50 | -3.892 | | | |
| | | 1.1 | folded | 328.2 | 0.45 | -10.218 | 328.1 | 0.45 | -10.233 |
| | | | unfolded | 328.0 | 0.45 | -10.248 | | | |
| | | 1.2 | folded | 347.0 | 0.43 | -16.858 | 347.2 | 0.43 | -16.788 |
| | | | unfolded | 347.3 | 0.43 | -16.718 | | | |
| | | 1.3 | folded | 365.8 | 0.43 | $-\infty$ | 365.6 | 0.43 | $-\infty$ |
| | | | unfolded | 365.3 | 0.43 | $-\infty$ | | | |
| Ribonuclease H | 2RN2 | 1.0 | folded | 310.8 | 0.23 | -4.050 | 310.5 | 0.23 | -3.924 |
| | | | unfolded | 310.2 | 0.23 | -3.799 | | | |
| | | 1.1 | folded | 328.0 | 0.23 | -8.243 | 327.7 | 0.23 | -8.150 |
| | | | unfolded | 327.3 | 0.23 | -8.058 | | | |
| | | 1.2 | folded | 343.9 | 0.22 | -12.275 | 344.0 | 0.22 | -12.230 |
| | | | unfolded | 344.1 | 0.22 | -12.186 | | | |
| | | 1.3 | folded | 361.2 | 0.22 | -16.411 | 361.1 | 0.22 | -16.591 |
| | | | unfolded | 360.9 | 0.22 | -16.771 | | | |

Supplementary Table 4. The protein name, PDB ID, threshold of native contacts fraction $Q_{kin}$, protein folding rate constant $k_F$, protein folding delay time $t_0$, coefficient of determination $R^2$ for curve fitting, mean *in silico* protein folding time $\langle\tau_F^{sim}\rangle$, mean experimental protein folding time $\langle\tau_F^{exp}\rangle$ and the scaling factor $\alpha = \frac{\langle\tau_F^{exp}\rangle}{\langle\tau_F^{sim}\rangle}$ of training-set proteins

| Protein | PDB ID | $Q_{kin}$ | $k_F$ (ns$^{-1}$) | $t_0$ (ns) | $R^2$ | $\langle\tau_F^{sim}\rangle$ (ns) | $\langle\tau_F^{exp}\rangle$ (s) | $\frac{\langle\tau_F^{exp}\rangle}{\langle\tau_F^{sim}\rangle}$ |
|---|---|---|---|---|---|---|---|---|
| EC298 | 1RYK | 0.83 | 0.151 | 1.873 | 0.99813 | 8.51 | 0.00011 [b] | 12,926 |
| λ-repressor$_{6\text{-}85}$ | 1LMB | 0.68 | 0.098 | 2.601 | 0.99950 | 12.81 | 0.00003 [b] | 2,342 |
| bACBP | 2ABD | 0.80 | 0.172 | 2.639 | 0.99892 | 8.44 | 0.00095 [b] | 112,559 |
| IM7 | 1CEI | 0.64 | 0.055 | 1.844 | 0.99890 | 20.00 | 0.00075 [b] | 37,500 |
| Im9 | 1IMQ | 0.64 | 0.222 | 2.064 | 0.99840 | 6.57 | 0.00066 [b] | 100,457 |
| Cytochrome B562 | 256B | 0.75 | 0.053 | 3.045 | 0.99935 | 21.99 | 0.00125 [c] | 56,844 |
| ABP1 SH3 [a] | 1JO8 | 0.89 | 0.365 (0.527)<br>0.027 (0.473) | 1.187<br>1.635 | 0.99980 | 20.58 | 0.08547 [b] | 4,153,061 |
| Fyn SH3 | 1SHF | 0.75 | 0.218 | 1.871 | 0.99943 | 6.45 | 0.01060 [b] | 1,643,411 |
| CspB Bc | 1C9O | 0.52 | 0.099 | 1.705 | 0.99971 | 11.84 | 0.00073 [b] | 61,655 |
| CspA | 1MJC | 0.64 | 0.084 | 2.603 | 0.99956 | 14.51 | 0.00503 [d] | 346,657 |
| Tenascin [a] | 1TEN | 0.60 | 0.150 (0.940)<br>0.008 (0.060) | 2.262<br>24.845 | 0.99907 | 17.56 | 0.16611 [b] | 9,459,567 |
| Twitchin | 1WIT | 0.60 | 0.063 | 2.698 | 0.99968 | 18.62 | 0.66667 [b] | 35,803,974 |
| Hpr | 1POH | 0.68 | 0.026 | 9.184 | 0.99851 | 47.72 | 0.06711 [e] | 1,406,329 |
| Urm1 | 2QJL | 0.48 | 0.012 | 1.908 | 0.99973 | 82.93 | 0.07576 [b] | 913,542 |
| Src SH2 | 1SPR | 0.83 | 0.217 | 1.643 | 0.99868 | 6.26 | 0.00016 [b] | 25,559 |

| | | | | | | | | |
|---|---|---|---|---|---|---|---|---|
| Azurin (apo) | 1E65 | 0.78 | 0.008 | 12.400 | 0.99955 | 132.08 | 0.00735 [b] | 55,648 |
| CheY | 3CHY | 0.52 | 0.026 | 14.708 | 0.99848 | 53.04 | 0.02653 [f] | 500,189 |
| Ribonuclease H | 2RN2 | 0.40 | 0.022 | 12.110 | 0.99917 | 58.07 | 1.35135 [g] | 23,271,052 |
| | | | | | $\langle Q_{\mathrm{kin}} \rangle = 0.66880,$ | | $\alpha$ = 4,331,293 | |

[a] Insufficiently sampled proteins. Their $k_F$ is shown as $k_i$ ($f_i$) and $t_0$ is shown as $t_i$, where $i$ = 1 or 2;

[b] Obtained from De Sancho et al.[11];

[c] Obtained from Wittung-Stafshede et al.[76];

[d] Obtained from Reid et al.[12];

[e] Obtained from Van Nuland et al.[77];

[f] Obtained from López-Hernéndez & Serrano[14];

[g] Obtained from Raschke & Marqusee[78];

Supplementary Table 5. The levels of $n_{\text{scal}}$ values used to parameterize an arbitrary protein

| Structural Class | Levels | | | | |
|---|---|---|---|---|---|
| | $\langle n^*_{\text{scal}} \rangle_{\text{class}}$ | $1.23 \times \langle n^*_{\text{scal}} \rangle_{\text{class}}$ | $1.46 \times \langle n^*_{\text{scal}} \rangle_{\text{class}}$ | $1.70 \times \langle n^*_{\text{scal}} \rangle_{\text{class}}$ | Overall highest |
| α | 1.1954 | 1.4704 | 1.7453 | 2.0322 | 2.5044 |
| β | 1.4732 | 1.8120 | 2.1508 | 2.5044 | 2.5044 |
| α/β | 1.1556 | 1.4213 | 1.6871 | 1.9644 | 2.5044 |
| Interface | 1.2747 | 1.5679 | 1.8611 | 2.1670 | 2.5044 |

Supplementary Table 6. Interaction site types and nonbonding parameters of the CG ribosome model

| | Chemical group | CG interaction site name | CG interaction site type | $R_i$ (Å) |
|---|---|---|---|---|
| rRNA and tRNA | Phosphate | P | P | 6.447660 |
| | Ribose | R | R | 5.231399 |
| | Adenine, Guanine | PU1 | BR [a] | 5.342436 |
| | | PU2 | | |
| | Uracil, Cytosine | PY | | |
| Ribosomal protein | ALA | SA | SA | 2.862278 |
| | CYS | SC | SC | 3.030648 |
| | ASP | SD | SD | 3.142894 |
| | GLU | SE | SE | 3.367386 |
| | PHE | SF | SF | 3.535755 |
| | GLY | SG | SG | 2.525540 |
| | HIS | SH | SH | 3.423509 |
| | ILE | SI | SI | 3.423509 |
| | LYS | SK | SK | 3.535755 |
| | LEU | SL | SL | 3.423509 |
| | MET | SM | SM | 3.423509 |
| | ASN | SN | SN | 3.199017 |
| | PRO | SP | SP | 3.086771 |
| | GLN | SQ | SQ | 3.423509 |
| | ARG | SR | SR | 3.704125 |
| | SER | SS | SS | 2.918401 |
| | THR | ST | ST | 3.142894 |
| | VAL | SV | SV | 3.311263 |
| | TRP | SW | SW | 3.816371 |
| | TYR | SY | SY | 3.591879 |

[a] Four different basic groups have the same CG interaction site type BR located at the centroid of each conjugated ring.

Supplementary Table 7. Equilibrium bond lengths, angles and dihedrals used for A-site and P-site NC-tRNA (or aa-tRNA)

| PDBid | Resolution (Å) | A-site tRNA | | | | P-site tRNA | | | |
|---|---|---|---|---|---|---|---|---|---|
| | | $b_0$ (CA-R) (Å) | $\theta_0$ (CA-R-PU2) (degree) | $\theta_0$ (CA-R-P) (degree) | $\varphi_0$ (CA-R-P-PU2) (degree) | $b_0$ (CA-R) (Å) | $\theta_0$ (CA-R-PU2) (degree) | $\theta_0$ (CA-R-P) (degree) | $\varphi_0$ (CA-R-P-PU2) (degree) |
| 5nwy | 2.9 | – | – | – | – | 4.66 | 143.32 | 99.51 | -170.62 |
| 5jte | 3.6 | 4.63 | 126.72 | 106.40 | 126.66 | 4.79 | 139.44 | 124.06 | -173.92 |
| 6i0y | 3.2 | – | – | – | – | 4.73 | 111.83 | 133.63 | -152.69 |
| 3jbu | 3.64 | – | – | – | – | 5.02 | 129.39 | 97.48 | -145.81 |
| 4uy8 | 3.8 | – | – | – | – | 4.93 | 111.98 | 134.20 | -154.60 |
| 6enj | 3.7 | 3.91 | 127.52 | 104.99 | 129.18 | – | – | – | – |
| 6enu | 3.1 | – | – | – | – | 4.43 | 136.54 | 109.20 | -163.82 |
| 5ju8 | 3.6 | – | – | – | – | 4.74 | 139.31 | 119.14 | -165.53 |
| Average | | 4.27 | 127 | 106 | 128 | 4.76 | 130 | 117 | -161 |

Supplementary Table 8. Original and rescaled $\langle\tau_{\text{trans}}^{\text{intrinsic}}\rangle_i$, $\langle\tau_{\text{PT}}^{\text{intrinsic}}\rangle$ and $\langle\tau_{\text{TL}}^{\text{intrinsic}}\rangle$ data, as well as corresponding normalized codon usage frequencies $w_i$

| | *E. coli* (at 310 K) | | | | | | | |
|---|---|---|---|---|---|---|---|---|
| | Codon | Fluitt & Viljoen's data[30] (s) | $w_i$ [31] | Rescaled data (s) | Codon | Fluitt & Viljoen's data[30] (s) | $w_i$ [31] | Rescaled data (s) |
| $\langle\tau_{\text{trans}}^{\text{intrinsic}}\rangle_i$ | UUU | 0.136 | 0.022499 | 0.068164 | GUU | 0.026 | 0.018409 | 0.013031 |
| | UUC | 0.195 | 0.016260 | 0.097735 | GUC | 0.208 | 0.015070 | 0.104251 |
| | UUG | 0.050 | 0.013430 | 0.025060 | GUG | 0.042 | 0.025879 | 0.021051 |
| | UUA | 0.157 | 0.013880 | 0.078689 | GUA | 0.073 | 0.010980 | 0.036588 |
| | UCU | 0.055 | 0.008660 | 0.027566 | GCU | 0.039 | 0.015530 | 0.019547 |
| | UCC | 0.246 | 0.008840 | 0.123297 | GCC | 0.415 | 0.025409 | 0.208000 |
| | UCG | 0.096 | 0.008810 | 0.048116 | GCG | 0.044 | 0.032749 | 0.022053 |
| | UCA | 0.106 | 0.007640 | 0.053128 | GCA | 0.083 | 0.020579 | 0.041600 |
| | UGU | 0.075 | 0.005260 | 0.037590 | GGU | 0.035 | 0.024439 | 0.017542 |
| | UGC | 0.109 | 0.006400 | 0.054631 | GGC | 0.049 | 0.028619 | 0.024559 |
| | UGG | 0.168 | 0.015170 | 0.084203 | GGG | 0.081 | 0.011270 | 0.040598 |
| | UGA | 0.012 | 0.001040 | 0.006014 | GGA | 0.324 | 0.008440 | 0.162391 |
| | UAU | 0.053 | 0.016380 | 0.026564 | GAU | 0.077 | 0.032319 | 0.038593 |
| | UAC | 0.077 | 0.012160 | 0.038593 | GAC | 0.116 | 0.019109 | 0.058140 |
| | UAG | 0.019 | 0.000250 | 0.009523 | GAG | 0.036 | 0.018199 | 0.018043 |
| | UAA | 0.011 | 0.002060 | 0.005513 | GAA | 0.057 | 0.039419 | 0.028569 |
| | CUU | 0.260 | 0.011470 | 0.130314 | AUU | 0.097 | 0.030229 | 0.048617 |
| | CUC | 0.204 | 0.010930 | 0.102246 | AUC | 0.128 | 0.024589 | 0.064154 |
| | CUG | 0.035 | 0.051888 | 0.017542 | AUG | 0.266 | 0.027589 | 0.133321 |
| | CUA | 0.286 | 0.003930 | 0.143345 | AUA | 0.128 | 0.004910 | 0.064154 |
| | CCU | 0.143 | 0.007250 | 0.071672 | ACU | 0.055 | 0.009040 | 0.027566 |
| | CCC | 0.197 | 0.005580 | 0.098738 | ACC | 0.153 | 0.022869 | 0.076685 |
| | CCG | 0.134 | 0.022659 | 0.067162 | ACG | 0.129 | 0.014470 | 0.064656 |
| | CCA | 0.237 | 0.008480 | 0.118786 | ACA | 0.178 | 0.007670 | 0.089215 |
| | CGU | 0.028 | 0.020599 | 0.014034 | AGU | 0.085 | 0.009080 | 0.042603 |
| | CGC | 0.035 | 0.021459 | 0.017542 | AGC | 0.127 | 0.015900 | 0.063653 |
| | CGG | 0.397 | 0.005700 | 0.198979 | AGG | 0.461 | 0.001510 | 0.231056 |
| | CGA | 0.034 | 0.003700 | 0.017041 | AGA | 0.190 | 0.002470 | 0.095229 |
| | CAU | 0.296 | 0.012850 | 0.148357 | AAU | 0.109 | 0.018269 | 0.054631 |
| | CAC | 0.222 | 0.009460 | 0.111268 | AAC | 0.161 | 0.021479 | 0.080694 |
| | CAG | 0.231 | 0.029139 | 0.115779 | AAG | 0.102 | 0.010700 | 0.051123 |
| | CAA | 0.179 | 0.015100 | 0.089716 | AAA | 0.076 | 0.033879 | 0.038092 |
| $\langle\tau_{\text{PT}}^{\text{intrinsic}}\rangle$ | | 0.000679 | – | 0.000340 | | | | |
| $\langle\tau_{\text{TL}}^{\text{intrinsic}}\rangle$ | | 0.008381 | – | 0.004201 | | | | |

Supplementary Table 9. The structural information and the parameterization information of the initial 14 *E. coli* enzymes for screening. (* denotes the instable domain or interface that cannot be stabilized using the highest level of $n_{\text{scal}}$ and the median value 1.8611 was used.)

| PDB ID | Name | Length | Domain and Interfaces | Structural Class | Minimum $n^{*}_{\text{scal}}$ required to keep kinetic stability |
|---|---|---|---|---|---|
| 1AKE | Adenylate kinase | 214 | Domain 1: 1 - 214 | α/β | 1.1556 |
| 1AQ2 | Phosphoenolpyruvate carboxykinase | 540 | Domain 1: 1 - 42; 66 - 227 | α/β | 1.1556 |
| | | | Domain 2: 43 - 65; 284 - 342 | β | 1.4732 |
| | | | Domain 3: 228 - 283; 343 - 540 | α/β | 1.1556 |
| | | | 1 \| 2 Interface | – | 1.2747 |
| | | | 1 \| 3 Interface | – | 1.2747 |
| | | | 2 \| 3 Interface | – | 1.2747 |
| 1C2T | Glycinamide ribonucleotide transformylase | 212 | Domain 1: 1 - 212 | α/β | 1.4213 |
| 1FDR | Flavodoxin reductase | 248 | Domain 1: 1 - 96 | β | 1.4732 |
| | | | Domain 1: 97 - 248 | α/β | 1.1556 |
| | | | 1 \| 2 Interface | – | 1.5679 |
| 1PDA | Porphobilinogen deaminase | 313 | Domain 1: 1 - 99; 200 - 220 | α/β | 1.1556 |
| | | | Domain 2: 100 - 199 | α/β | 1.1556 |
| | | | Domain 3: 221 - 313 | α/β | 1.4213 |
| | | | 1 \| 2 Interface | – | 1.5679 |
| | | | 1 \| 3 Interface | – | 1.2747 |
| | | | 2 \| 3 Interface | – | 1.5679 |
| 1VHL | Dephospho-CoA kinase | 206 | Domain 1: 1 - 206 | α/β | 1.1556 |
| 2FMT | Methionyl-tRNA formyltransferase | 315 | Domain 1: 0 - 208 | α/β | 1.4213 |
| | | | Domain 2: 209 - 314 | α/β | 1.6871 |
| | | | 1 \| 2 Interface | – | 1.8611 |
| 2OFP | Ketopantoate reductase | 303 | Domain 1: 1 - 167 | α/β | 1.1556 |
| | | | Domain 2: 168 - 303 | α | 1.1954 |
| | | | 1 \| 2 Interface | – | 1.8611 * |
| 2ZCV | Quinone oxidoreductase 2 | 303 | Domain 1: 1~136; 171~193; 259~267 | α/β | 1.1556 |
| | | | Domain 2: 137~170; 194~258; 268~303 | α/β | 1.9644 |
| | | | 1 \| 2 Interface | – | 2.167 |
| 3CW7 | DNA-3-methyladenine glycosylase 2 | 282 | Domain 1: 1 - 112 | α/β | 1.1556 |
| | | | Domain 2: 113 - 230 | α/β | 1.1556 |
| | | | Domain 3: 231 - 282 | α | 1.1954 |

| | | | | | |
|---|---|---|---|---|---|
| | | | 1 \| 2 Interface | – | 1.8611 * |
| | | | 1 \| 3 Interface | – | 1.2747 |
| | | | 2 \| 3 Interface | – | 1.2747 |
| 3K6L | Peptide deformylase | 169 | Domain 1: 0 - 168 | α/β | 1.4213 |
| 3LBF | Protein-L-isoaspartate O-methyltransferase | 208 | Domain 1: 1 - 208 | α/β | 1.1556 |
| 4C5C | D-alanine--D-alanine ligase B | 306 | Domain 1: 1 - 84 | α/β | 1.1556 |
| | | | Domain 2: 85 - 110; 184 - 306 | α/β | 1.1556 |
| | | | Domain 3: 111 - 183 | α/β | 1.1556 |
| | | | 1 \| 2 Interface | – | 2.1670 |
| | | | 1 \| 3 Interface | – | 1.2747 |
| | | | 2 \| 3 Interface | – | 2.1670 |
| 4KJK | Dihydrofolate reductase | 159 | Domain 1: 1-159 | α/β | 1.4213 |

Supplementary Table 10. Parameterization results for CAT-III

| REX input and output parameters | | | | | Average | Linear regression parameters | | | |
|---|---|---|---|---|---|---|---|---|---|
| $n_{\text{scal}}$ | Starting state | $T_{\text{m}}$ (K) | $Q_{\text{eq}}$ | $\Delta G_{\text{UN}}^{\text{sim}}(T^{\text{exp}})$ (kcal/mol) | $\Delta G_{\text{UN}}^{\text{sim}}(T^{\text{exp}})$ (kcal/mol) | $m$ | $b$ | $R^2$ | $n_{\text{scal}}^{*}$ |
| 1.0 | folded | 323.5 | 0.67 | -9.027 | -6.910 | -14.688 | 8.006 | 0.9658 | 1.2471 |
| | unfolded | 303.5 | 0.55 | -4.792 | | | | | |
| 1.1 | folded | 343.2 | 0.71 | -9.700 | -7.695 | | | | |
| | unfolded | 322.8 | 0.77 | -5.691 | | | | | |
| 1.2 | folded | 361.2 | 0.71 | -12.438 | -9.847 | | | | |
| | unfolded | 340.8 | 0.78 | -7.256 | | | | | |

Supplementary Table 11. Experimentally measured rate constant ($k_{cat}$)[a] and Michaelis constant ($K_M$)[b] of the enzymatic reaction catalyzed by the fast and slow DDLB variants and the corresponding relative specific activity $k_{cat}^{slow}/k_{cat}^{fast}$.

| Biological replicates | $k_{cat}^{slow}$ (min$^{-1}$) | $K_M^{slow}$ (μM) | $k_{cat}^{fast}$ (min$^{-1}$) | $K_M^{fast}$ (μM) | $k_{cat}^{slow}/k_{cat}^{fast}$ |
|---|---|---|---|---|---|
| 1 | 986.05 ± 16.61 | 1181.26 ± 79.81 | 1045.70 ± 18.22 | 1759.32 ± 77.84 | 0.9430 |
| 2 | 895.40 ± 18.58 | 2272.66 ± 114.37 | 1125.37 ± 20.68 | 2414.56 ± 105.02 | 0.7956 |
| 3 | 1083.42 ± 16.94 | 2163.70 ± 80.36 | 1088.12 ± 24.36 | 2570.70 ± 133.06 | 0.9957 |
| 4 | 771.40 ± 9.00 | 1517.97 ± 44.98 | 958.48 ± 16.65 | 1633.22 ± 75.38 | 0.8048 |
| 5 | 601.89 ± 9.84 | 1568.39 ± 78.38 | 698.28 ± 13.87 | 2082.02 ± 118.30 | 0.8619 |
| $\langle k_{cat}^{slow}/k_{cat}^{fast} \rangle$ | 0.8802[c], 95%CI [0.8126, 0.9478] | | | | |

[a] $k_{cat}$ is presented as mean ± standard deviation errors of the curve fitting;

[b] $K_M$ is presented as mean ± standard deviation errors of the curve fitting;

[c] $\langle k_{cat}^{slow}/k_{cat}^{fast} \rangle$ is statistically less than 1 with a one-tailed t-test $p$-value of 0.0186.

#### Supplementary Figures

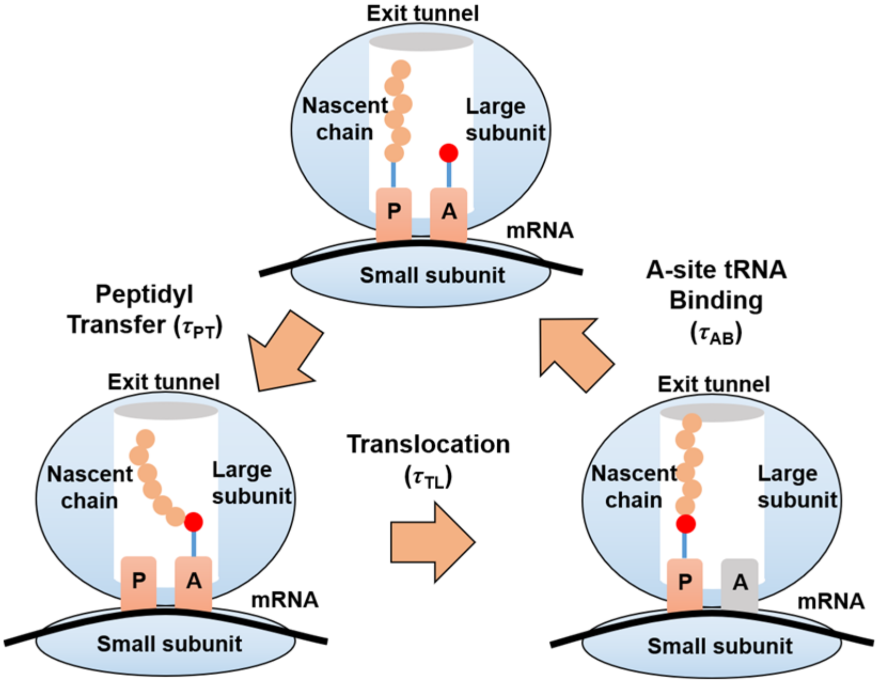

**Supplementary Figure 1. Scheme diagram of the on-ribosome continuous synthesis process.** The on-ribosome continuous synthesis process is simplified into three steps. First, aa-tRNA binds into the A-site. Then, the $NC_l$-tRNA at P-site and the aa-tRNA at A-site undergo the peptidyl transfer reaction catalyzed by the PTC and then forms $NC_{l+1}$-tRNA at A-site. Finally, P-site tRNA is translocated to the E-site and the A-site tRNA is translocated to the P-site, resulting the empty A-site waiting for the next aa-tRNA binding. The ribosome is shown in blue; the $NC_l$-tRNA, $NC_{l+1}$-tRNA and aa-tRNA are shown in orange for NC, pink for tRNA and red for the new amino acid; A-site and P-site are marked as "A" and "P", respectively; the empty A-site is shown in grey; the exit tunnel is shown in white and the mRNA is shown in black. The dwell times before peptidyl transfer, translocation and A-site tRNA binding are denoted as $\tau_{PT}$, $\tau_{TL}$ and $\tau_{AB}$, respectively.

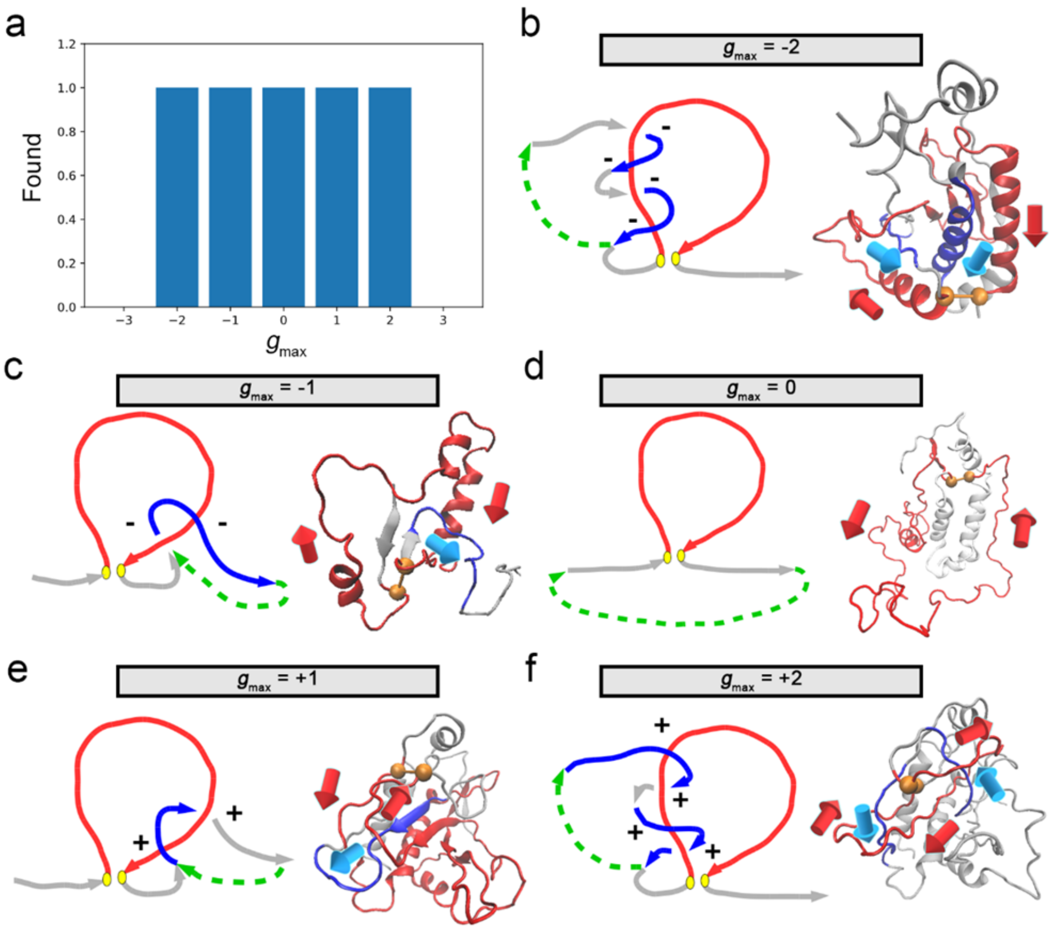

**Supplementary Figure 2. Test case of *g* for detecting topological entanglements.** a, The observed maximum *g* values ($g_{max}$) in the post-translational trajectories of the fast CAT-III variant. The link diagrams (left) and the representative structures (right) are shown for each $g_{max}$ values in panel b, c, d, e and f. The closed loop formed by the native contact (yellow balls) are in red and the tail curves that form entanglements are in blue. The rest regions are shown in gray. The directions of those curves (from N-ter to C-ter) are marked in both the link diagrams and the representative structures. The +/- signs that are used to count the linking numbers are labelled near the curve intersections in the link diagrams. The imaginary curves that close the tail curves are shown in green dashed arrows.

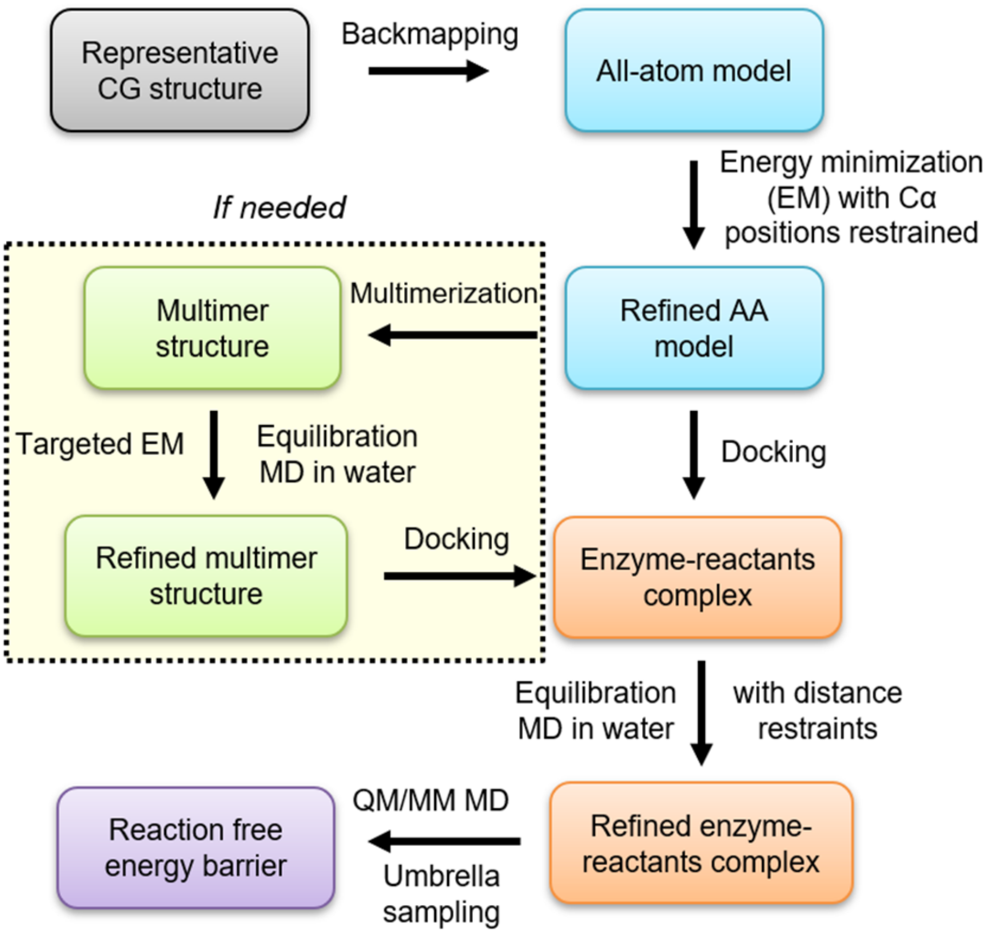

**Supplementary Figure 3. Scheme diagram of the estimation workflow for activation free energy barrier height.** The main workflow for estimating activation free energy barrier height for a representative coarse-grained (CG) structure is shown here. More details are described in Supplementary Methods Section 14.3.

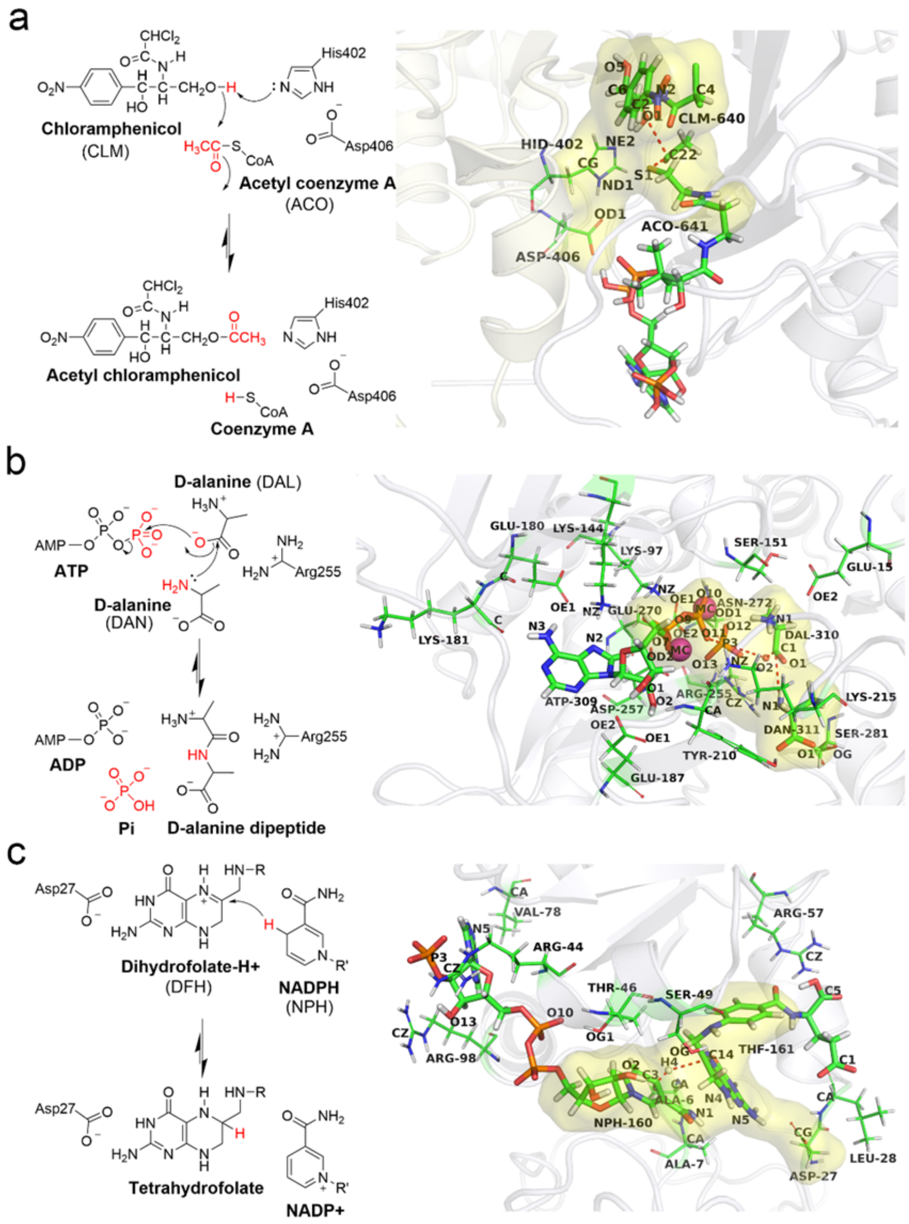

**Supplementary Figure 4. Reaction mechanisms and the ligand binding poses for candidate enzymes used in this study.** a, Reaction mechanism[68] (left) and ligand binding pose (right) of CAT-III; b, Reaction mechanism[70] (left) and ligand binding pose (right) of DDLB; c, Reaction mechanism[71] (left) and ligand binding pose (right) of DHFR. In reaction schemes, the moving groups are highlighted in red. In ligand binding pose diagrams, ligands are shown as bold sticks, whereas the amino acid residues that took part in the distance restraints are shown as thin sticks. The reaction coordinates are presented as red dotted lines. The QM regions are covered by yellow areas. The proteins are shown as Cartoon representation and different chains are shown in different colors.

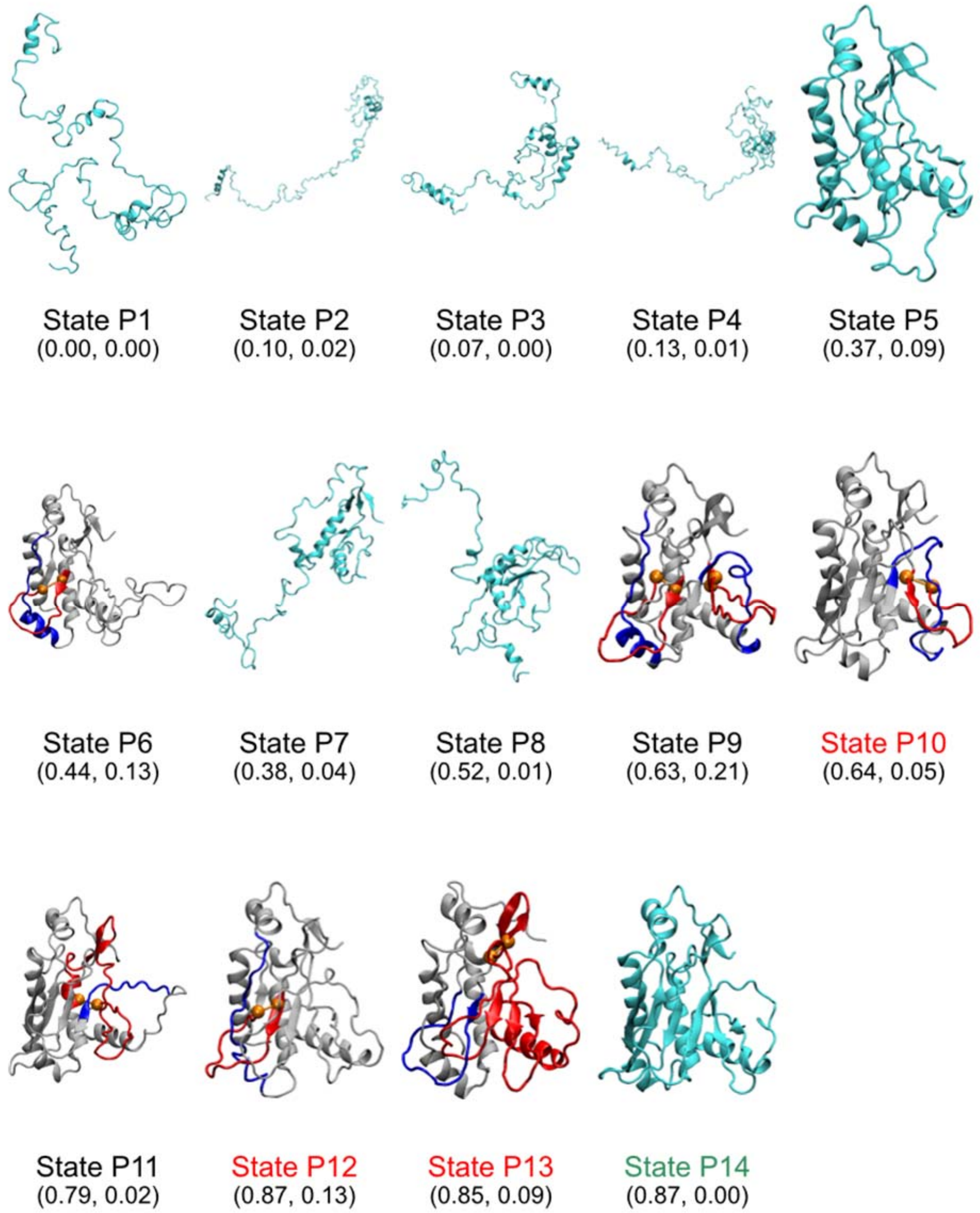

**Supplementary Figure 5. Representative structures of the metastable states of monomer CAT-III post-translational folding process.** The structures are represented by the Cartoon model. The ordinary intermediate structures, as well as the native structure, are shown in cyan. The entangled structures are shown in white, where the closed loop formed by the native contact (shown as two orange balls) is highlighted in red and the segment threading the red loop is highlighted in blue. The state name and the ($Q_{act}$, $G$) coordinates are presented on the bottom of each structure. The name of the near-native-like kinetics traps are highlighted in red and the native state is highlighted in green.

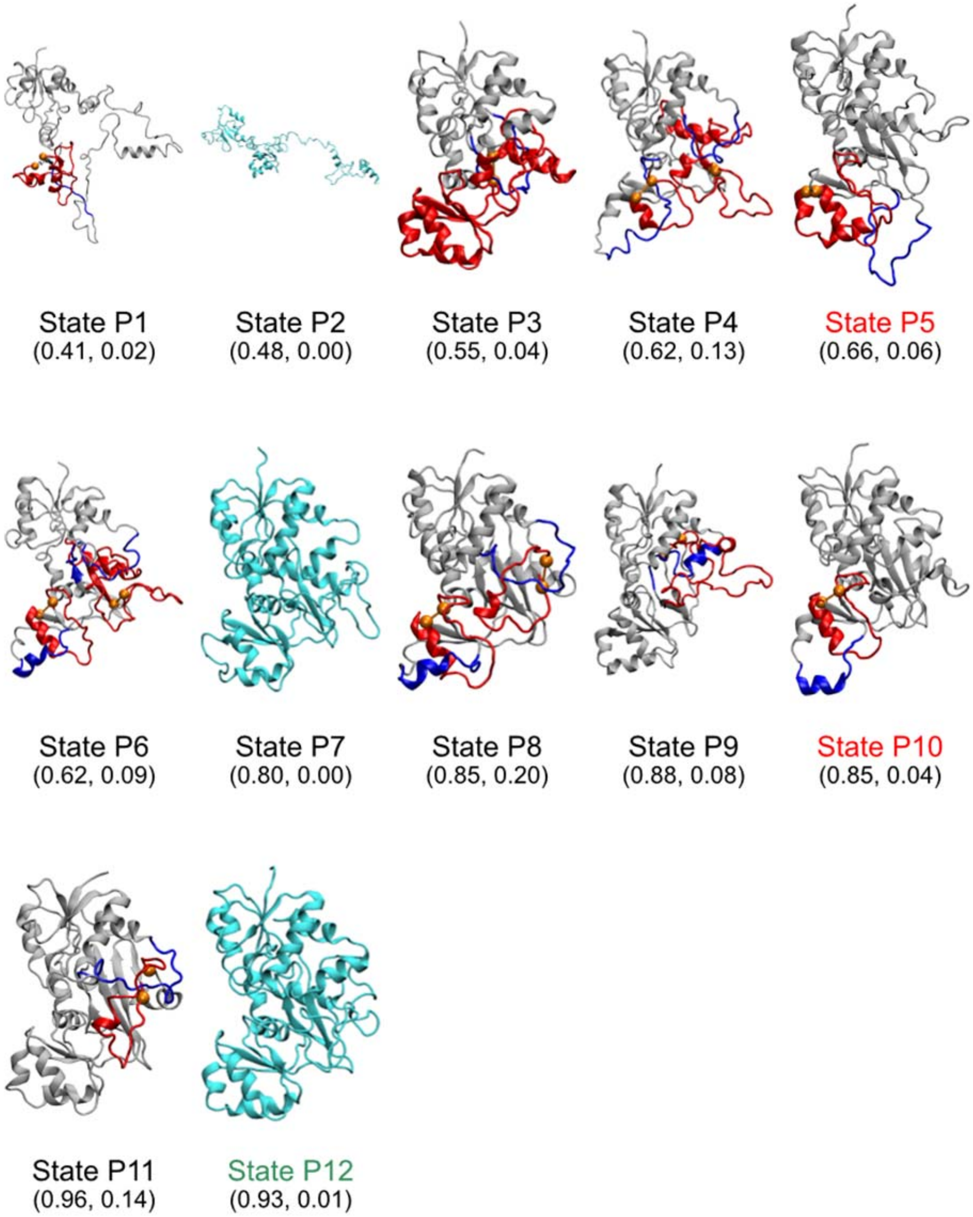

**Supplementary Figure 6. Representative structures of the metastable states of DDLB post-translational folding process.** The structures are represented by the Cartoon model. The ordinary intermediate structures, as well as the native structure, are shown in cyan. The entangled structures are shown in white, where the closed loop formed by the native contact (shown as two orange balls) is highlighted in red and the segment threading the red loop is highlighted in blue. The state name and the ($Q_{act}$, $G$) coordinates are presented on the bottom of each structure. The name of the near-native-like kinetics traps are highlighted in red and the native state is highlighted in green.

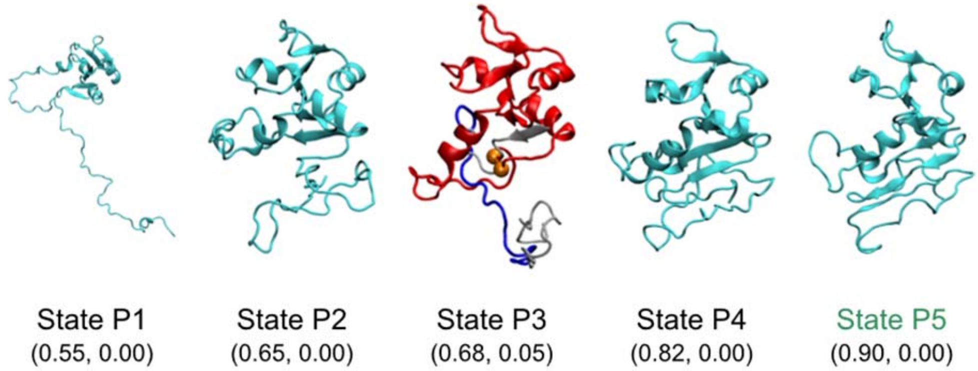

**Supplementary Figure 7. Representative structures of the metastable states of DHFR post-translational folding process.** The structures are represented by the Cartoon model. The ordinary intermediate structures, as well as the native structure, are shown in cyan. The entangled structures are shown in white, where the closed loop formed by the native contact (shown as two orange balls) is highlighted in red and the segment threading the red loop is highlighted in blue. The state name and the ($Q_{act}$, $G$) coordinates are presented on the bottom of each structure. The name of the native state is highlighted in green. No near-native-like kinetic traps were detected for DHFR.

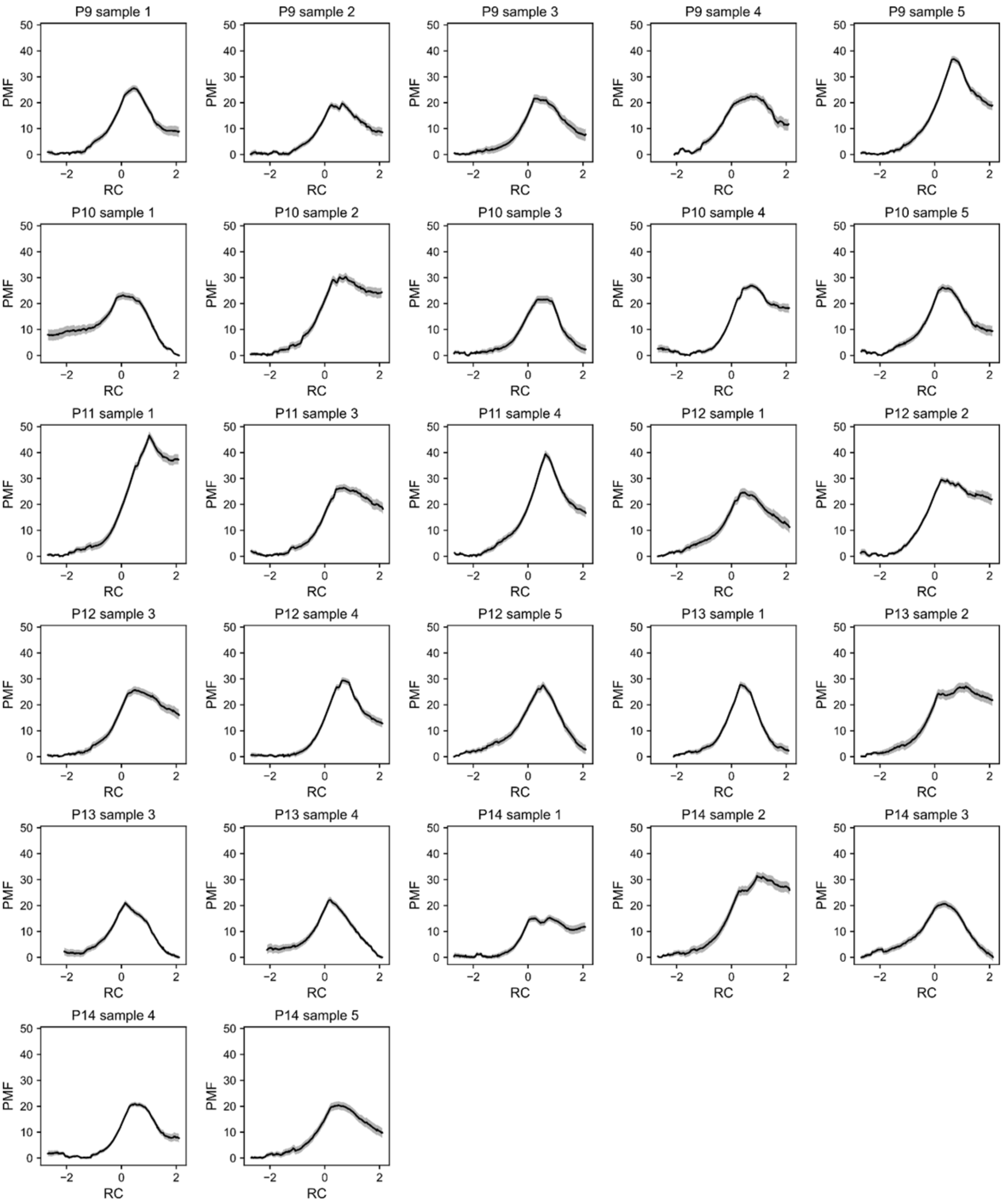

**Supplementary Figure 8. Potential of mean force (PMF) curves as a function of the reaction coordinate (RC) for the samples of CAT-III.** Only the samples that have successful substrates docking and QM/MM simulations are presented in this figure. Error bars (95% CIs) are shown as gray stripes.

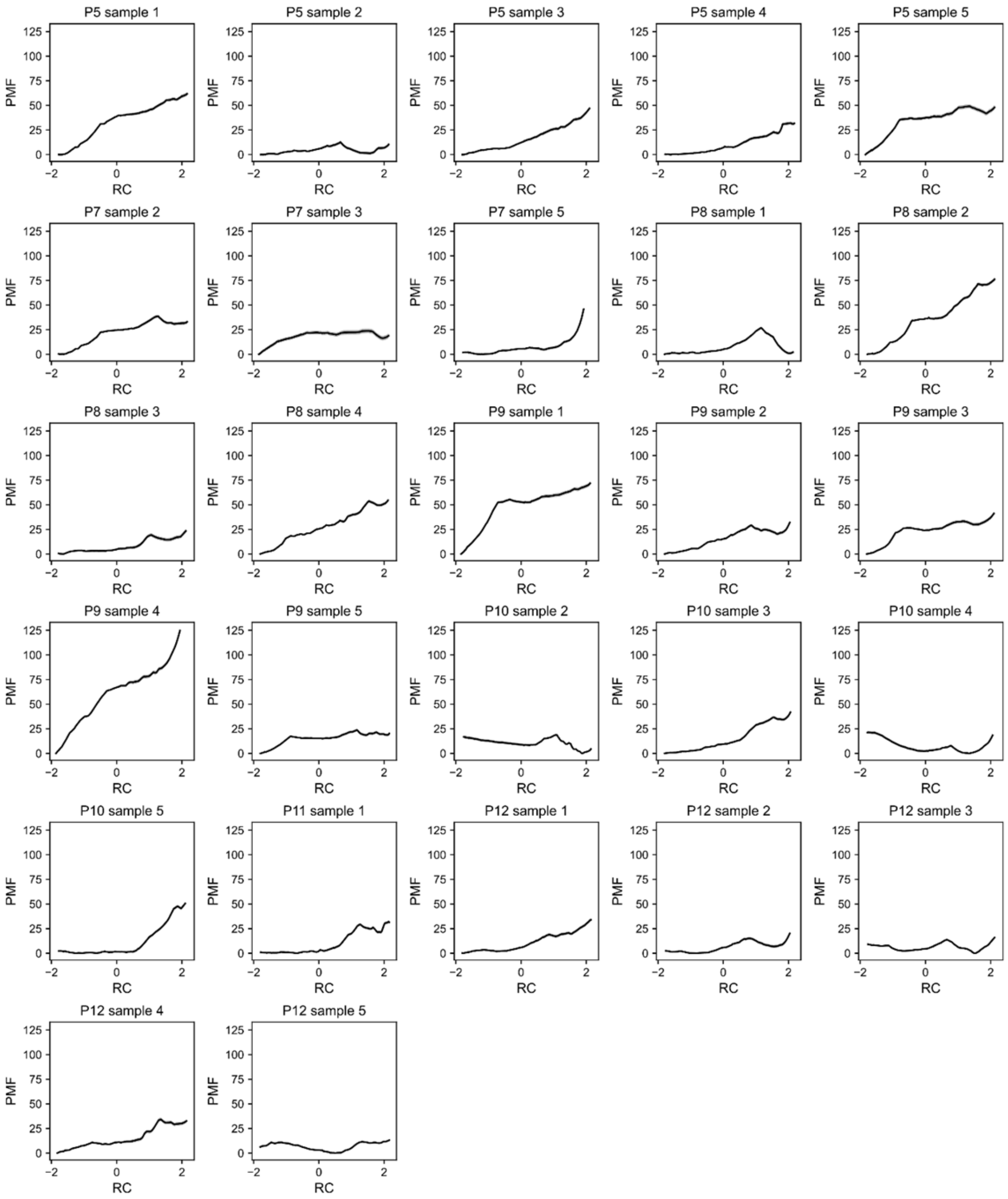

**Supplementary Figure 9. Potential of mean force (PMF) curves as a function of the reaction coordinate (RC) for the samples of DDLB.** Only the samples that have successful substrates docking and QM/MM simulations are presented in this figure. Error bars (95% CIs) are shown as gray stripes.

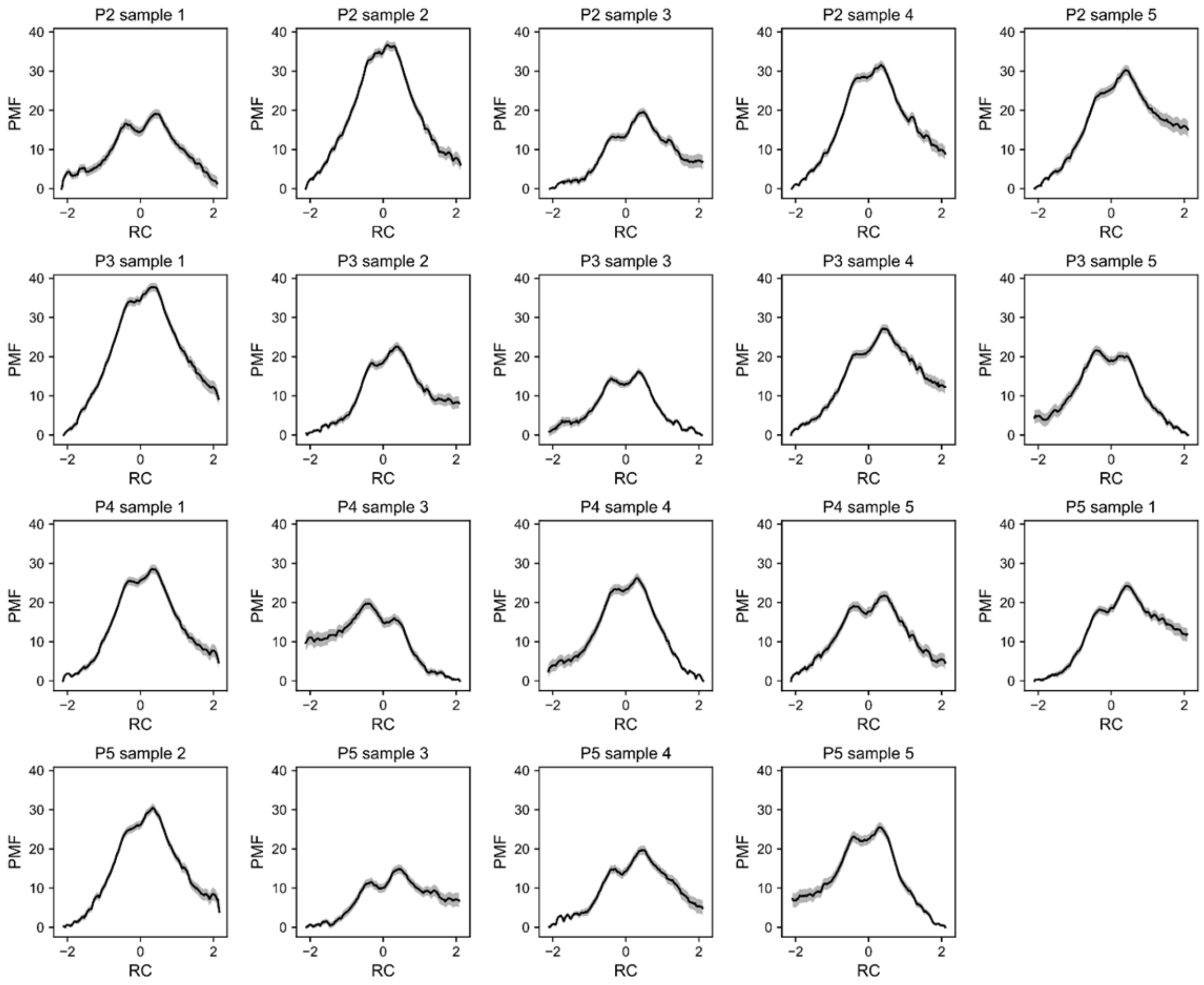

**Supplementary Figure 10. Potential of mean force (PMF) curves as a function of the reaction coordinate (RC) for the samples of DHFR.** Only the samples that have successful substrates docking and QM/MM simulations are presented in this figure. Error bars (95% CIs) are shown as gray stripes.

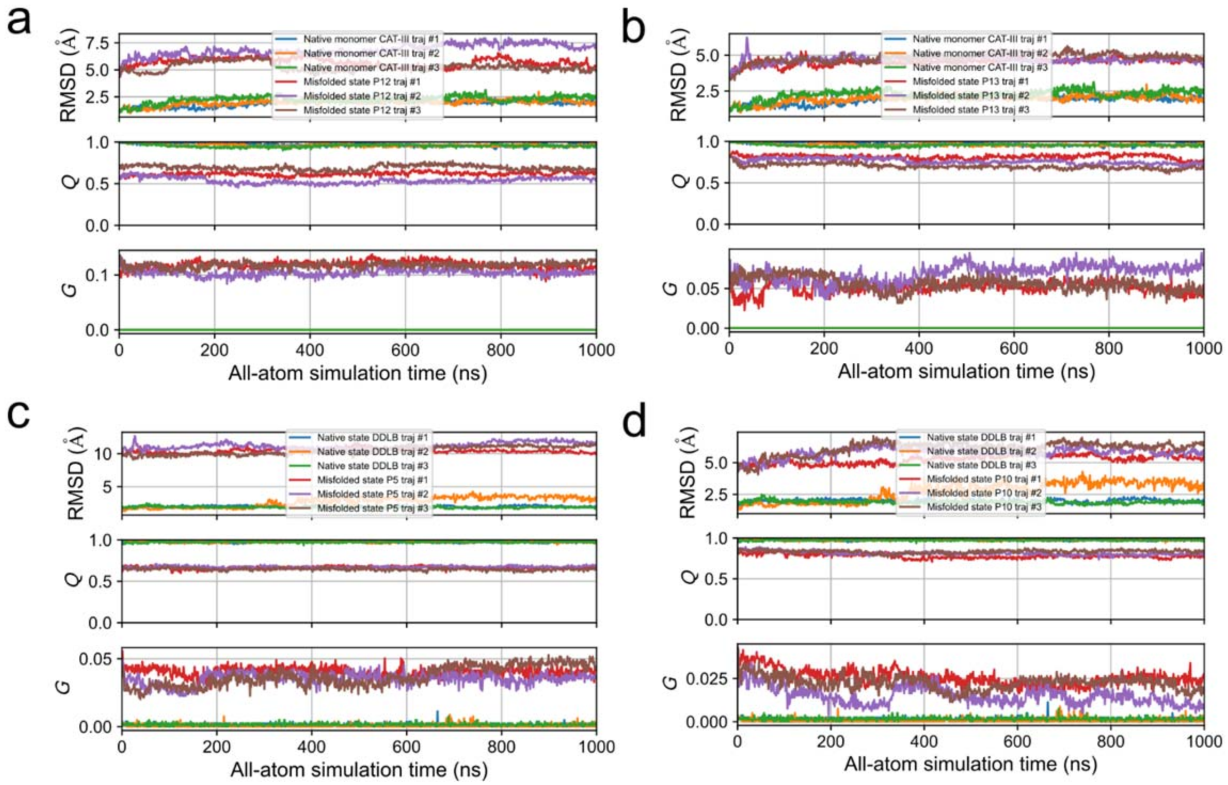

**Supplementary Figure 11. Persistence of the topological entanglements in the all-atom level simulations for (a) state P12 of CAT-III, (b) state P13 of CAT-III, (c) state P5 of DDLB and (d) state P10 of DDLB.** Each test has three replicates. The structural deviation (RMSD), native contact formed (*Q*) and the change of topological entanglements (*G*) of the testing structures are evaluated and compared with these quantities of the native structures (MD simulations starting from the crystal structures), respectively.

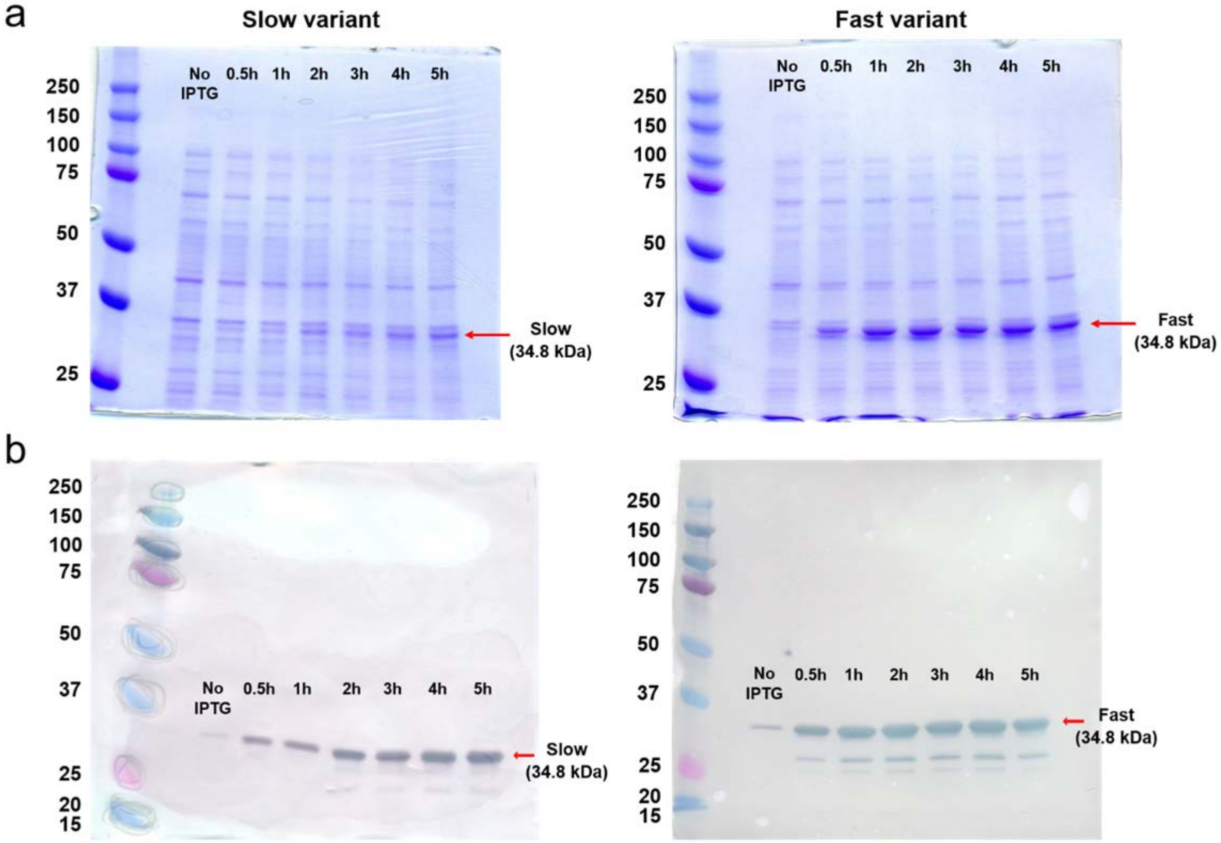

**Supplementary Figure 12. Time-course expression analysis of slow and fast DDLB variants by (a) SDS-PAGE and (b) Western blot ($His_6$-tag), respectively.** Both the gel band intensity in panel a and the $His_6$ band intensity in panel b are higher for the fast variant compared to the slow variant. The fast variant had a higher level of protein expression 5 hours after induction than the slow variant, consistent with the fast mRNA variant translating faster

#### Supplementary Videos

**Supplementary Video I.** Co- and post-translational folding trajectory of a misfolded CAT-III that has a noncovalent lasso entanglement formed after synthesis. The conformation gets trapped in state P12 at the end. The nascent chain is shown in cyan, where the closed loop formed by the native contact is highlighted in red and the segment threading the red loop is highlighted in blue. At each nascent chain length, the conformation is obtained from the last frame of the simulation trajectory. The time of ejection, dissociation and post-translation processes are presented using the experimental timescale on the top left of the scene.

**Supplementary Video II.** Co- and post-translational folding trajectory of a correctly folded CAT-III without any topological entanglements formed. The nascent chain is colored based on secondary structure elements, where alpha-helices are magenta and beta-sheets are yellow. At each nascent chain length, the conformation is obtained from the last frame of the simulation trajectory. The time of ejection, dissociation and post-translation processes are presented using the experimental timescale on the top left of the scene.

**Supplementary Video III.** Co- and post-translational folding trajectory of a misfolded DDLB that has a noncovalent lasso entanglement formed during the synthesis and persisted in the post-translational dynamics. The conformation gets trapped in state P5 at the end. The nascent chain is shown in cyan, where the closed loop formed by the native contact is highlighted in red and the segment threading the red loop is highlighted in blue. At each nascent chain length, the conformation is obtained from the last frame of the simulation trajectory. The time of ejection, dissociation and post-translation processes are presented using the experimental timescale on the top left of the scene.

**Supplementary Video IV.** Co- and post-translational folding trajectory of a correctly folded DDLB without any topological entanglements formed. The nascent chain is colored based on secondary structure elements, where alpha-helices are magenta and beta-sheets are yellow. At each nascent chain length, the conformation is obtained from the last frame of the simulation trajectory. The time of ejection, dissociation and post-translation processes are presented using the experimental timescale on the top left of the scene.
